# Targeting cancer with small molecule pan-KRAS degraders

**DOI:** 10.1101/2023.10.24.563163

**Authors:** Johannes Popow, William Farnaby, Andreas Gollner, Christiane Kofink, Gerhard Fischer, Melanie Wurm, David Zollman, Andre Wijaya, Nikolai Mischerikow, Carina Hasenoehrl, Polina Prokofeva, Heribert Arnhof, Silvia Arce-Solano, Sammy Bell, Georg Boeck, Emelyne Diers, Aileen B. Frost, Jake Goodwin-Tindall, Jale Karolyi-Oezguer, Shakil Khan, Theresa Klawatsch, Manfred Koegl, Roland Kousek, Barbara Kratochvil, Katrin Kropatsch, Arnel A. Lauber, Ross McLennan, Sabine Olt, Daniel Peter, Oliver Petermann, Vanessa Roessler, Peggy Stolt-Bergner, Patrick Strack, Eva Strauss, Nicole Trainor, Vesna Vetma, Claire Whitworth, Siying Zhong, Jens Quant, Harald Weinstabl, Bernhard Kuster, Peter Ettmayer, Alessio Ciulli

**Author notes:** These authors contributed equally to this work (Johannes Popow, William Farnaby, Andreas Gollner, Christiane Kofink).

## Abstract

Despite the high prevalence of cancers driven by KRAS mutations, to date only the G12C mutation has been clinically proven to be druggable via covalent targeting of the mutated cysteine amino acid residue (*1*). However, in many cancer indications other KRAS mutations, such as G12D and -V, are far more prevalent and small molecule concepts that can address a wider variety of oncogenic KRAS alleles are in high clinical demand (*2*). Here we show that a single small molecule can be used to simultaneously and potently degrade 13 out of 17 of the most prevalent oncogenic KRAS alleles, including those not yet tractable by inhibitors. Compared with inhibition, degradation of oncogenic KRAS results in more profound and sustained pathway modulation across a broad range of KRAS mutant cell lines. As a result, KRAS degraders inhibit growth of the majority of cancer cell lines driven by KRAS mutations while sparing models without genetic KRAS aberrations. Finally, we demonstrate that pharmacological degradation of oncogenic KRAS leads to tumour regression in vivo. Together, these findings unveil a new path towards addressing KRAS driven cancers with small molecule degraders.

**One-Sentence Summary:** The most prevalent KRAS variants which drive tumour growth in a major share of cancer patients can be targeted with a single small molecule degrader.

## Introduction

Kirsten rat sarcoma viral oncogene homologue (KRAS) is the most commonly mutated oncogene in human cancers (*3*). Variants, predominantly mutations at Glycine (G) 12 or Glutamine (Q) 61, increase the proportion of activated, GTP-loaded KRAS, enhancing RAF-MEK-ERK (MAPK) signaling and drive tumor growth. To date, clinical advances in drugging oncogenic KRAS variants have relied on specific interactions of small molecules with the mutated amino acid residues. For example, covalent inhibitors rely on a cysteine residue available in KRAS^G12C^ (*1, 4*) while reversible inhibitors rely on interactions between basic moieties and a variant specific aspartate residue in KRAS^G12D^ (*5*). Indeed, beyond these variants, even pre-clinical target validation has been reliant on genetic means that lack dose, kinetic and temporal control. New concepts that can lead to single agents capable of potently and selectively addressing multiple KRAS variants stand to have major clinical impact. Recently, BI-2865 and BI-2493 were disclosed as the first example of KRAS inhibitors capable of engaging a broader spectrum of KRAS alleles than clinically validated inhibitors (*6*).

Small molecule heterobifunctional degraders (proteolysis targeting chimeras - PROTACs) are transforming drug development for oncology, with >25 degrader drugs in clinical trials for several indications (*7, 8*). Early PROTACs recruiting the von Hippel-Lindau (VHL) or cereblon E3 ubiquitin ligases to KRAS^G12C^ have been disclosed based on covalent KRAS binders (*9, 10*). However, as KRAS engagement mediated by covalent target ligands itself leads to irreversible inhibition, this approach inherently lacks key mechanistic advantages of degraders such as substoichiometric and catalytic mode of action (*11*). To engage and degrade a wider range of KRAS variants beyond G12C, PROTAC degraders thus require leveraging of non-covalent KRAS binders, the discovery of which has proven inherently challenging. However, there has been recent progress in this direction, with non-covalent KRAS^G12D^ degraders based on KRAS^G12D^-selective inhibitors currently undergoing early clinical testing (*12*).

Here, we provide pre-clinical validation for a single small molecule degrader, targeting 13 of the 17 most prevalent KRAS mutants, which illuminates a new pan-KRAS degradation concept conferring potential for major clinical benefit. By employing structure-guided design we identify ACBI3, which achieves *in vivo* degradation of oncogenic KRAS, resulting in durable pathway modulation and tumour regressions in *KRAS* mutant xenograft mouse models.

## Identification of VHL-based KRAS Degraders Based on Non-Covalent KRAS Binders

As a starting point, we chose to use a high affinity KRAS switch II pocket ligand we have recently disclosed (*6, 13*). Analysis of co-crystal structures highlighted a solvent exposed sub-pocket formed by the amino acids His95, Glu62, and Asp92, which we deemed to be a promising position to install linkers for PROTAC design (Fig. 1A). Due to the importance of the interaction of the basic center of these substituents with the surrounding amino acids we focused our PROTAC design approach on motifs maintaining the basicity of the molecule in this region. Thorough X-ray crystallographic analysis (PDB accession code 8QUG) of the recently published BI-2865 and close analogs revealed that the homopiperazine compound **1** (dissociation constant for KRAS^G12V^ by surface plasmon resonance – SPR – *K_D_* = 25 nM, table S1), provided a suitable trajectory for linker attachment (Fig. 1A). Applying an initial screening approach based on alkyl and polyethylene glycol (PEG)-based linkers in combination with VHL ligase binders based on VH032 (*14*), we tested the resulting molecules in a biophysical screening assay based on fluorescence polarization (FP). This assay reports affinity for the VHL:EloC:EloB (VCB) complex with or without pre-incubation at saturating concentrations of KRAS^G12D^. Cooperative VCB:PROTAC:KRAS^G12D^ ternary complex formation is indicated by lower competitor concentrations achieving half maximal displacement (*K_D_*) in the presence of KRAS^G12D^ (*15*). This highlighted compound **2** (Fig. 1B) as a highly cooperative (alpha = 479, table S1) and high-affinity ternary complex inducer (*K_D_* = 7 187 nM *vs* 15 nM in the absence or presence of KRAS^G12D^, respectively) (Fig. 1C, table S1). We orthogonally confirmed ternary complex formation via SPR yielding a ternary complex dissociation half-life (*t_1/2_*) of 159 s and an equilibrium dissociation constant (*K_D_*) of 20 nM (Fig. 1D, table S1). Compound **2** dose-dependently degraded KRAS^G12D^ in GP5d cells with a concentration inducing half maximal degradation (*DC_50_*) at 24 hours of 607 nM and a maximal extent of degradation (*D_max_*) of >95% (Fig. 1E, table S2). Similar results were obtained by Western blotting using an antibody detecting both wild type and mutant KRAS expressed by GP5d cells (fig. S1A). Compound **2** also degraded KRAS^G12V^ in SW620 cells (*DC_50_* = 1 203 nM, *D_max_* >95%) indicating that KRAS degradation is not limited to KRAS^G12D^ (fig. S1B, table S2). To enable high throughput characterization of degraders, we set up a bioluminescence-based degradation assay in GP5d cells expressing KRAS^G12D^ with a small luminescence complementation (HiBiT) tag inserted into the endogenous KRAS locus yielding comparable degradation parameters (fig. S1C, table S2) (*16*). Degradation was abolished in the presence of the NEDD8-activating enzyme inhibitor MLN4924 (*17*) or the competing VHL ligand VH298 (*18*) (fig. S1D) supporting that KRAS degradation by compound **2** depends on intracellular recruitment of an active VHL ligase complex. We also detected direct KRAS ubiquitination and intracellular formation of ternary complexes by bioluminescence resonance energy transfer (BRET) based assays (fig. S1E and -F). The cellular target engagement assays, comparing the ability of compound **2** to engage VHL in either permeabilized or live cells (*IC_50_* values 5 µM and >10 µM, respectively) indicated the need to further optimize cellular permeability as well as VHL affinity (Fig. 1F).

**Fig. 1.**
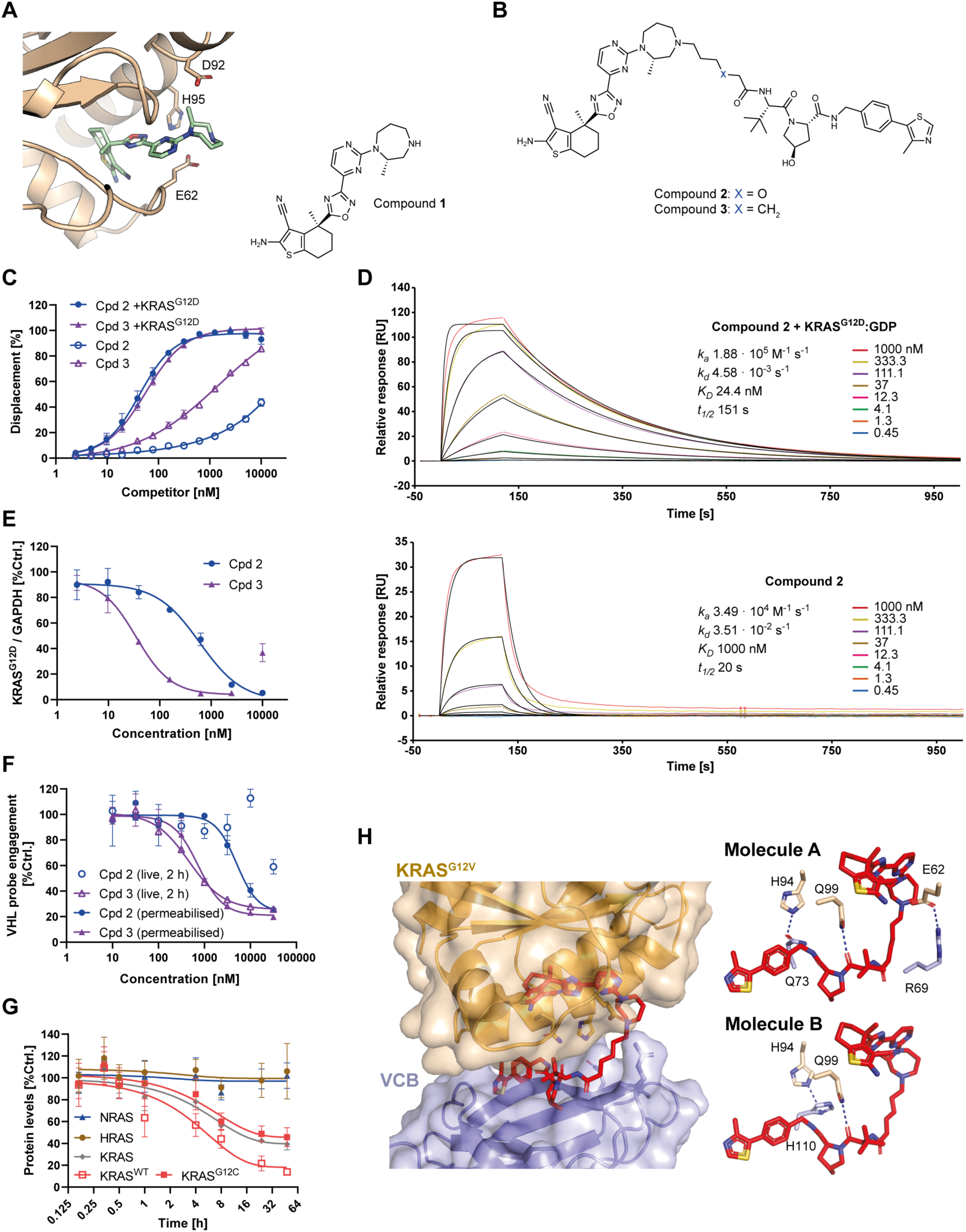
Identification of reversible, KRAS selective degraders. (**A**) Exit vector explored to derivatise KRAS-binders. Three-dimensional crystal structure of KRAS displayed in brown, compound 1 in green. PDB Accession code 8QUG. (**B**) Chemical structures of compound **2** and -**3**. (**C**) VCB FP displacement for compounds **2** and -**3** in presence or absence of saturating KRAS^G12D^ concentrations (N=3, SD). (**D**) SPR characterization of ternary complex for VCB, KRAS^G12D^:GDP and compound **2** (N=3, representative trace). (**E**) Dose-dependent degradation of KRAS^G12D^ in GP5d cells by capillary electrophoresis (24 hours, N=3, SD). (**F**) VHL target engagement by NanoBRET in live or permeabilized HEK293 cells for compounds **2** and -**3** (N=3, SD). (**G**) Degradation time course of total KRAS, KRAS^WT^, KRAS^G12C^, H- and NRAS by 1 µM compound **3** in NCI-H358 cells (N=3, SD) by targeted proteomics. (**H**) Ternary complex co-crystal structure of VCB: compound **3**:KRAS^G12V^.

Having established compound **2** as a VHL-based KRAS degrader, we went on to synthesize a molecular matched pair, replacing the oxygen atom in the linker by a methylene group yielding compound **3** (Fig. 1B). Compound **3** still displayed positive cooperativity and long-lived ternary complex half-life (alpha = 17, *K_D_* = 340 nM by FP, *K_D_* = 80 nM and *t_1/2_* = 103 s by SPR, Fig. 1C, fig. S1G, table S1) albeit reduced compared to compound **2**.

Degradation potency improved by greater than ten-fold for KRAS^G12D^ (GP5d, *DC_50_* = 32 nM, *D_max_* = 99%) (Fig. 1E, table S2) and KRAS^G12V^ (SW620, *DC_50_* = 278 nM, *D_max_* = 88%) (fig. S1B, table S2) with similar results again obtained by <u>Western</u> <u>blotting</u> using an antibody detecting both wild type and mutant KRAS and loss of degradation at high concentrations favoring binary rather than ternary engagement, referred to as “hook effect” (fig. S1A).

Cellular target engagement assays showed that both improved cellular permeability as well as VHL affinity likely contributed to the superior degradation potency of compound **3** (Fig. 1F). Selective targeting of KRAS while sparing H- and NRAS has been linked to the therapeutic window of the KRAS ligand we based our design upon (*6*). We therefore tested selectivity of degradation by targeted proteomics in the cell line NCI-H358. While we detected degradation of both KRAS^G12C^ (*D_max_* ≥ 54%) and KRAS^WT^ (*D_max_* ≥ 86%), which represents roughly one third of the KRAS pool in this cell line, we detected no significant change in H- or NRAS levels (Fig. 1G, fig. S1H).

To understand the binding mode and enable structure-based optimization, we determined the ternary co-crystal structure of compound **3** in complex with VCB and KRAS^G12V^ at 2.2 Å resolution (Fig. 1H) (PDB 8QW6). Compound **3** adopts a “fishhook” conformation, burying the VHL binder section of compound **3** in a *de novo* binding interface formed between KRAS^G12V^ and VHL. The crystal structure contains two molecules of ternary complex in the asymmetric unit, revealing a consistent binding mode (RMSD = 0.54 Å), but a different fingerprint of protein-protein and protein-ligand interactions. This indicates that there is subtle flexibility within the compound **3** ternary complex, allowing the complex to shift between nearby networks of favorable protein-protein interactions. Using a biophysics-led screening approaching we identified compound **3** as a KRAS degrader prototype based on a non-covalent target ligand. However, we noted that compound **3** has a poorer efficiency of degradation for KRAS^G12C^ as opposed to WT (Fig. 1G), motivating us to investigate if further improvements could be made to broaden the range of KRAS mutants we could potently degrade.

## Identification of Pan-KRAS Degraders

With the objective of identifying molecules that potently degrade multiple KRAS mutants we next sought to employ structure-based design using the solved ternary co-crystal structure of compound **3** to improve ternary complex stability and intracellular VHL engagement. Extending ternary complex stability (increased *t_1/2_*) has been shown to improve rate and potency of target protein degradation (*19–21*). Analysis of the VCB:compound **3**:KRAS^G12V^ co-crystal structure (Fig. 1H) highlighted an opportunity to enhance interactions within the ternary complex by improving π-stacking between the exit vector amide of the VHL ligand and Tyr112. We therefore switched the amide for an isoxazole, which has been previously reported to improve VCB affinity in this position in other contexts (*22*), yielding compound **4** (Fig. 2A). A ternary complex co-crystal structure of VCB:compound **4:**KRAS^G12V^ (PDB 8QW7) confirmed a consistent overall binding mode to that observed for compound **3** (RMSD = 0.72 Å), with compound **4** able to engage in π-stacking interaction between the isoxazole and Tyr112 of VCB as designed (Fig. 2B). As with compound **3**, the compound **4** ternary complex revealed different networks of induced protein-protein interactions within the two ternary molecules in the asymmetric unit, indicating complex flexibility. FP, SPR and cellular target engagement studies support a minor improvement in binary VHL engagement for compound **4**, with moderate cooperativity retained (fig. S2A-C, table S1). A larger shift in durability of ternary complex can be observed via SPR for compound **4** (*t_1/2_* = 230 s, table S1) *vs* compound **3** (*t_1/2_* = 103 s, table S1). We then investigated the impact of these structural changes on degradation kinetics in live HiBiT-tagged GP5d cells (fig. S3A-D). While maximal degradation rates (*λ_max_*) did not vary significantly between compound **3** and compound **4** (approximately 0.5 1/hour for both compounds, table S2), the concentration inducing the half maximal cellular degradation rate (*Dmax_50_*) was three-fold lower for compound **4** (*Dmax_50_* = 56 nM for compound **3** *vs Dmax*_50_ = 17 nM for compound **4**) (Fig. 2C, table S2). The improved *Dmax_50_* for compound **4** translated into drastically improved cellular VHL-dependent degradation potencies of compound **4** *vs* compound **3** for endogenous KRAS^G12D^ (GP5d, *DC_50_* = 1 nM, *D_max_* = 99.5%, table S1) and KRAS^G12V^ (SW620, *DC_50_* = 13 nM, *D_max_* = 89%) (Fig. 2D, fig. S3E-G).

**Fig. 2.**
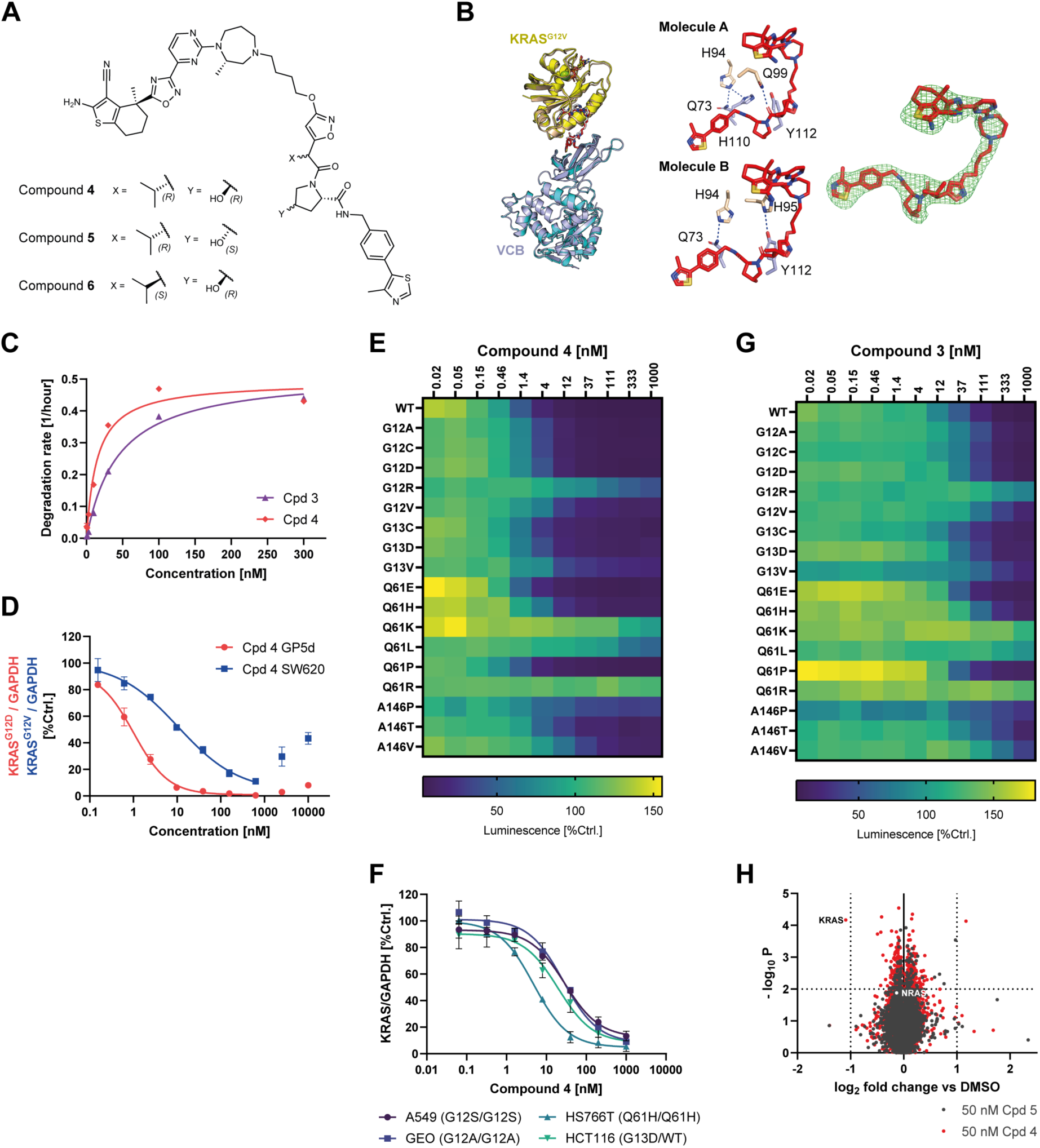
Targeting of all major oncogenic KRAS alleles and sustained pathway engagement with rapid degraders. (**A**) Chemical structures of compound **4** and inactive degrader controls compound **5** ((*S*)-hydroxyproline derivative compound **4**) and -**6** ((*S*)-isopropyl). (**B**) Left: superposition of ternary complex structures of compound **3** (red):VCB (light blue):KRAS^G12V^ (wheat), and compound **4** (white):VCB (teal):KRAS^G12V^ (yellow) displaying conserved ternary binding orientations (RMSD = 0.72 Å). Middle: views from the compound 4 ternary complex PROTAC binding sites (molecules A and B) displaying side chains of residues involved in potential strong protein:protein interactions (blue dotted lines). Right: Fo-Fc omit map contoured to 3 σ for compound **4**. (**C**) Concentration dependency of degradation rates for compound **3** and compound **4** in HiBiT-tagged GP5d cells (N=6, mean±95% CI). (**D**) Dose-dependent degradation of KRAS^G12D^ and KRAS^G12V^ in GP5d and SW260 cells, respectively by capillary electrophoresis (24 hours, N=3, SD). (**E**) Degradation of retrovirally transduced HiBiT-tagged indicated KRAS mutants by compound **4** in GP5d cells (18 hours, N=3) (**F**) Dose-dependent degradation of KRAS^G12S^, KRAS^G12A^, KRAS^G13D^ and KRAS^Q61H^ in A549, GEO, HCT116 and HS766T cells, respectively, by capillary electrophoresis (24 hours, N=3, SD). (**G**) Degradation of retrovirally transduced HiBiT-tagged indicated KRAS mutants by compound **3** in GP5d cells (18 hours, N=2) (**H**) Whole cell proteomics MS analysis of GP2d cells treated with 50 nM compound **4** or inactive stereoisomer compound **5** (8 hours, N=3) data for NRAS and compound **4** highlighted in white.

To gauge KRAS mutation specificity on a broader basis without contribution of potentially confounding factors, we established an isogenic series of cell lines transduced with retroviral constructs expressing the most prevalent KRAS mutants (isoform 4B) fused to a HiBiT-tag in GP5d cells. Analysis of clones isolated from the resulting cell pools revealed that up to 30-fold differences in expression of tagged KRAS did not affect degradation potency of compound **4** in this system (fig. S3H and I). Constructs of both KRAS isoforms (4A and 4B) were degraded with comparable potencies (fig. S3J). Dose titration revealed that compound **4** efficiently degraded 13 of the 17 most prevalent KRAS mutant alleles and KRAS^WT^ with single digit nanomolar potency (Fig. 2E). Supporting this, we obtained comparable results for endogenous KRAS in cell lines expressing KRAS^G12S^, -G12A, -G13D and -Q61H treated with compound **4** (Fig. 2F). KRAS mutants with a more complete loss of GTPase activity, such as KRAS^G12R^ (*DC_50_* = 45 nM, *D_max_* = 59%) and -Q61L/K/R (*DC_50_* > 470 nM *D_max_* < 60%), were degraded less potently, consistent with the relative binary binding affinity to these KRAS mutants of the ligand class employed in this study (*6*). Together, these data are consistent with degradation of KRAS mutants with residual GTPase activity by engagement of the inactive, GDP-bound state. Compound **3** was associated with a similar degradation spectrum albeit with reduced potency for all degradable mutants suggesting that the optimized potency of compound **4** *vs* compound **3** affected a broad spectrum of KRAS alleles (Fig. 2G). We went on to assess the cellular selectivity of compound **4**-induced KRAS degradation by unbiased MS-proteomics in GP2d cells, using compound **5**, a VHL binding deficient stereoisomer of compound **4** as a negative control. KRAS was the only detected protein showing > 2-fold depletion (*p* < 0.01) with NRAS levels not significantly affected (Fig. 2H). Similar results were obtained in KRAS^G12C^ mutant MiaPaCa-2 cells using another VHL binding deficient isomer, compound **6** as a degradation deficient control (fig. S3K). In summary, our structure-guided design approach led us to compound **4**, a highly selective KRAS degrader now acting on a broad spectrum of KRAS mutants with high prevalence in cancer patients.

## KRAS Degradation Potently Suppresses Oncogenic Signaling and Proliferation

Next, we set out to test the ability of a single pan-KRAS degrading molecule, compound **4**, to selectively suppress oncogenic signaling and proliferation in cancer cell lines driven by diverse KRAS mutants. To this end, we first confirmed degradation of KRAS by compound **4** in KRAS-dependent (GP2d, SW620 and NCI-H358 expressing KRAS^G12D^, - G12V or -G12C, respectively) as well as in KRAS independent (A-375, HEK293) cell lines (Fig. 3A, fig. S4A). To compare degradation *vs* inhibition of KRAS, we also profiled compound **5** (VHL-binding deficient stereoisomer) which has the same overall chemical formula and close-to-identical physicochemical properties and engages KRAS non-covalently without inducing KRAS degradation (Fig. 3A, fig. S4A). Alongside, we also profiled the covalent inhibitor of KRAS^G12C^ Sotorasib (*23*) or the non-covalent KRAS^G12D^ inhibitor MRTX-1133 (*24*) as appropriate (Fig. 3A, fig. S4A). Both inhibition and degradation repressed the established markers of MAPK signaling pERK (fig. S4B-D) and *DUSP6* (Fig. 3B, fig. S4E) in KRAS dependent cell lines. Both KRAS independent cell lines did not exhibit suppression of pERK or *DUSP6* exceeding 50% of control (Fig. 3B, fig. S4B and D-E). Compound **4** was > 10-fold more potent in suppressing MAPK signalling in KRAS dependent cell lines compared to its VHL binding deficient stereoisomer compound **5** indicating that E3 ligase engagement and subsequent target degradation as compared to inhibition results in more potent pathway engagement (Fig. 3B, fig. S4B-E, table S2). In the KRAS-dependent cell lines MAPK pathway engagement coincided with prominent inhibition of proliferation, with degradation conferring a greater than ten-fold and up to 100-fold potency gain (Fig. 3C, fig. S4F, table S2). Neither degradation nor inhibition had appreciable antiproliferative effects in the KRAS independent cell lines A-375 and HEK293 (Fig. 3C, fig. S4F). Comparable results albeit with lower potency were obtained for compound **3** (fig. S5, table S2). While long term treatment with compound **4** sustainably suppressed mutant and wild type KRAS expression for up to 72 hours in *KRAS* mutant and *KRAS^WT^* cell lines (fig S6A-D, top row), we noted a pronounced recovery of phospho-protein markers of both the MAPK-and phosphoinositide kinase pathways in *KRAS* mutant cell lines for both KRAS degraders and -inhibitors (fig. S6A-C, second to third row). This observation suggests that reactivation via feedback mechanisms occurs in the presence of minimal levels of mutant KRAS possibly, at least in part mediated by H- and NRAS as suggested by H- and NRAS contributing to MAPK pathway output upon KRAS inhibition (*6*). *DUSP6* modulation and induction of apoptosis confirmed the greater than 10-fold potency gain for KRAS degradation over -inhibition (compare results for compound **4** and 10-fold excess of the degradation inactive stereoisomer compound **5** in fig S6A-C, fourth and fifth row). In agreement with KRAS mediating mitogenic signaling, both KRAS inhibition and degradation induced a prominent G1/G0 cell cycle arrest in *KRAS* mutant but not *KRAS^WT^* cell lines, with a greater than 10-fold potency gain for degradation *vs* inhibition (fig. S6A-D, bottom panels). Analogous to short term proliferation assays, clonogenic growth assays also support the enhanced potency of KRAS degradation over KRAS inhibition in *KRAS* mutant but not *KRAS^WT^*cell lines (fig. S6E-H).

**Fig. 3.**
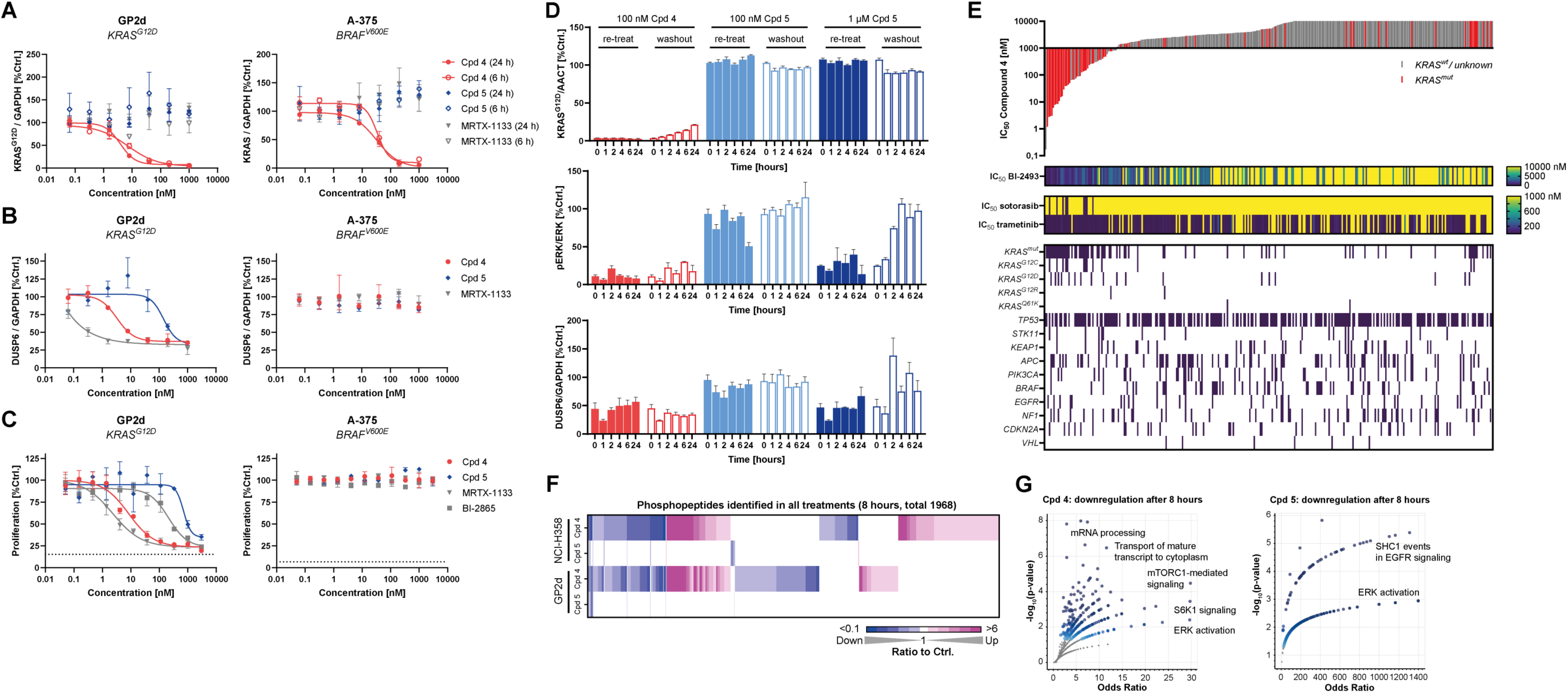
Pan-KRAS degradation impacts MAPK signaling and cancer cell proliferation. (**A**) KRAS degradation (6 and 24 hours, N=2, range), (**B**) *DUSP6* modulation (6 hours, N=3, SD), (**C**) proliferation (5 days, N=3, SD) for compound **4**, compound **5** and MRTX1133 in *KRAS^G12D^* and *BRAF^V600E^* cell lines, respectively. (**D**) KRAS^G12D^, pERK and DUSP6 recovery in GP2d cells upon re-treatment or washout and VHL competition following 18 hours pre-treatment with compound **4** or compound **5** (time after washout, N=3, SD). (**E**) Proliferation inhibition data for 300 cancer cell lines by compound **4** (bars represent *IC_50_* per cell line), BI-2493 (*6*), sotorasib or trametinib (heatmap) (5 days, N=3) including genetic features of tested cell lines. (**F**) Heat map of phosphorylation events upon treatment with compound **4** or compound **5** in NCI-H358 (*KRAS^G12C^*, 500 nM) and GP2d (*KRAS^G12D^*, 100 nM) cells (8 hours, N=3). (**G**) Volcano plots of combined gene set enrichment analyses of phospho-proteome changes induced by compound **4** (upper panel) or compound **5** (lower panel) in NCI-H358 and GP2d cells at 8 hours.

Pharmacological degradation of a target can result in target resynthesis dependent pathway engagement (*25*). To test whether this applies to our KRAS degraders, we pre-treated KRAS^G12D^-dependent GP2d cells with compound **4** to achieve maximal KRAS degradation or the inactive stereoisomer compound **5** at the identical concentration and 10-fold excess to achieve comparable pathway inhibition by non-covalent engagement of KRAS. After pre-incubation, the medium was exchanged to wash out compound **4** or its inactive stereoisomer and replaced by culture medium containing VH298 to compete against VHL binding of any residual compound **4** (Fig 3D, empty bars). Alternatively, we re-exposed the cells to the same treatments as during the pre-incubation (Fig. 3D, filled bars). Upon washout, KRAS^G12D^ levels recovered in a time-dependent fashion reaching 21% of untreated controls 24 hours after washout of compound **4**. In contrast, KRAS^G12D^ levels remained below 5% of controls upon re-addition of compound **4** (Fig. 3D, top panel, compare red empty bars with red solid bars, respectively). Both pERK and DUSP6 rapidly recovered after washout of the inactive stereoisomer compound **5** reaching control levels after 2-4 hours even though the 10-fold excess of compound **5** achieved pERK and DUSP6 suppression comparable to the compound **4** pre-incubation. In contrast, cells pre-treated with compound **4** exhibited long lasting MAPK pathway suppression consistent with the delayed recovery of KRAS^G12D^ after washout (Fig. 3D, middle and bottom panels, compare empty red bars with empty dark blue bars). Of note, re-addition of compound **4** or 1 µM compound **5** appreciably suppressed pERK and DUSP6 for up to 24 hours after media exchange and compound re-addition.

Hence, KRAS degradation results in long-lasting MAPK pathway suppression with delayed recovery compared to non-covalent KRAS inhibition upon compound withdrawal. We also compared the antiproliferative activity of compound **4** to that of KRAS^G12C-^specific covalent-, non-covalent KRAS- and the MEK inhibitor trametinib in a cell line panel comprising 300 cell lines covering a range of KRAS mutations and cancer indications (Fig. 3E). On a global scale, cell lines bearing KRAS mutants had lower concentrations required for half-maximal proliferation inhibition (*IC_50_*) as compared to WT cell lines (geometric mean *IC_50_* = 739 nM *vs* 4 934 nM, respectively) (fig. S6I). Applying a sensitivity cutoff of 1 µM, all sotorasib-sensitive *KRAS^G12C^*-mutant cell lines were also sensitive to compound **4**. In addition, sensitivity to compound **4** was correlated with that of the KRAS inhibitor BI-2493 albeit with higher potency for compound **4**. Notably, most cell lines sensitive to compound **4** as well as BI-2493 bear a KRAS mutation whereas trametinib exhibits less selective antiproliferative effects and potently kills *KRAS^WT^*cell lines (Fig. 3E, heat maps underneath bar plot).

Consistent with the drastically reduced activity of compound **4** on highly hydrolysis impaired KRAS mutants, the two *KRAS^Q61K^* mutant cell lines were insensitive (Fig. 3E, genetic features underneath bar plot). Apart from reasons pertaining to high-throughput antiproliferative activity profiling, some features may explain insensitivity of a subset of *KRAS* mutant cell lines to compound **4**. For instance, lack of antiproliferative activity of compound **4** in cell lines sensitive to KRAS ablation may be linked to its susceptibility to drug efflux (table S3, CaCo-2 assay). Co-treatment with the ABCB1 efflux-pump inhibitor Zosuquidar (*26*) rescued both proliferation and KRAS degradation in LS513 cells expressing high levels of ABCB1 (fig. S6J and K).

To characterize the effects of pan-KRAS degradation more globally on the molecular level, we performed a time-resolved (phospho-)proteomic analysis of NCI-H358 and GP2d cells in response to compound **4** or its VHL-binding deficient isomer compound **5** at the concentration of compound **4** achieving maximal degradation in each cell line (fig. S7A). The results recapitulated KRAS (but not HRAS) degradation in both models (fig. S7B and -C), and quantification of ∼20 000 phosphorylation sites allowed for the comprehensive characterization of affected phosphorylation events (fig. S7D). Phosphorylation events detected in all treatments revealed a pronounced overlap for both compounds in either cell line, with several cell line-specific differences (Fig. 3F). While both compound **4** and compound **5** engaged the MAPK pathway, compound **4** modulated phosphorylation events detected in all treatments with greater effect size than compound **5** (Fig. 3F, fig. S7E). Only a few phosphorylation events displayed more pronounced modulation by compound **5** as compared to compound **4** (Fig. 3F, fig. S7E). Gene ontology enrichment analysis demonstrated reduced activity of multiple pathways, including the cell cycle, in response to compound **4** but not compound **5** (Fig. 3G, fig. S7F and -G). Hence, both inhibition and degradation of KRAS modulate largely overlapping sets of phosphorylation events albeit with distinct effect size and potency. Taken together, as compared to target inhibition achieved by compounds with comparable molecular properties, KRAS degradation enables greater than 10-fold higher potency paired with extended and more pronounced suppression of MAPK signaling. In conclusion, KRAS degradation selectively shuts down oncogenic signaling more potently and more durably than a matched molecular pair inhibitor in KRAS dependent cell lines. This leads to suppression of cell cycle progression, induction of apoptosis and thus inhibition of proliferation in the context of a wide range KRAS mutants *in vitro*.

## *In Vivo* KRAS Degradation leads to regressions in KRAS mutant tumor-bearing mice

Next, we wished to understand possible advantages of KRAS degradation *in vivo*. For instance, degradation has been shown to extend pharmacodynamic (PD) efficacy beyond the detectable pharmacokinetic (PK) presence of a degrader in other settings (*25*). Pharmacokinetic profiling of compound **4** suggested insufficient exposure irrespective of the route of administration. Plasma concentrations achieved via intravenous (*i.v.*) or sub-cutaneous (*s.c.*) dosing did not cover the predicted *in vivo DC_50_* (Fig. 4A, table S3 and S4). The latter was estimated using potency of degradation of HiBiT-labeled KRAS^G12D^ in assays with fetal calf serum (FCS) substituted by serum of NMRI mice or human serum. Whereas the degradation potency of compound **4** was comparable in FCS and human serum (*DC_50_* = 1.4 and 1.9 nM, respectively), we noted a 24-fold potency drop in presence of 10% mouse serum (*DC_50_* = 33 nM) (Fig. 4B). This yields a predicted *in vivo DC_50_* of 851 nM for the GP2d model in NMRI mice (*vs* 3.6 nM in FCS) (table S2).

**Fig. 4.**
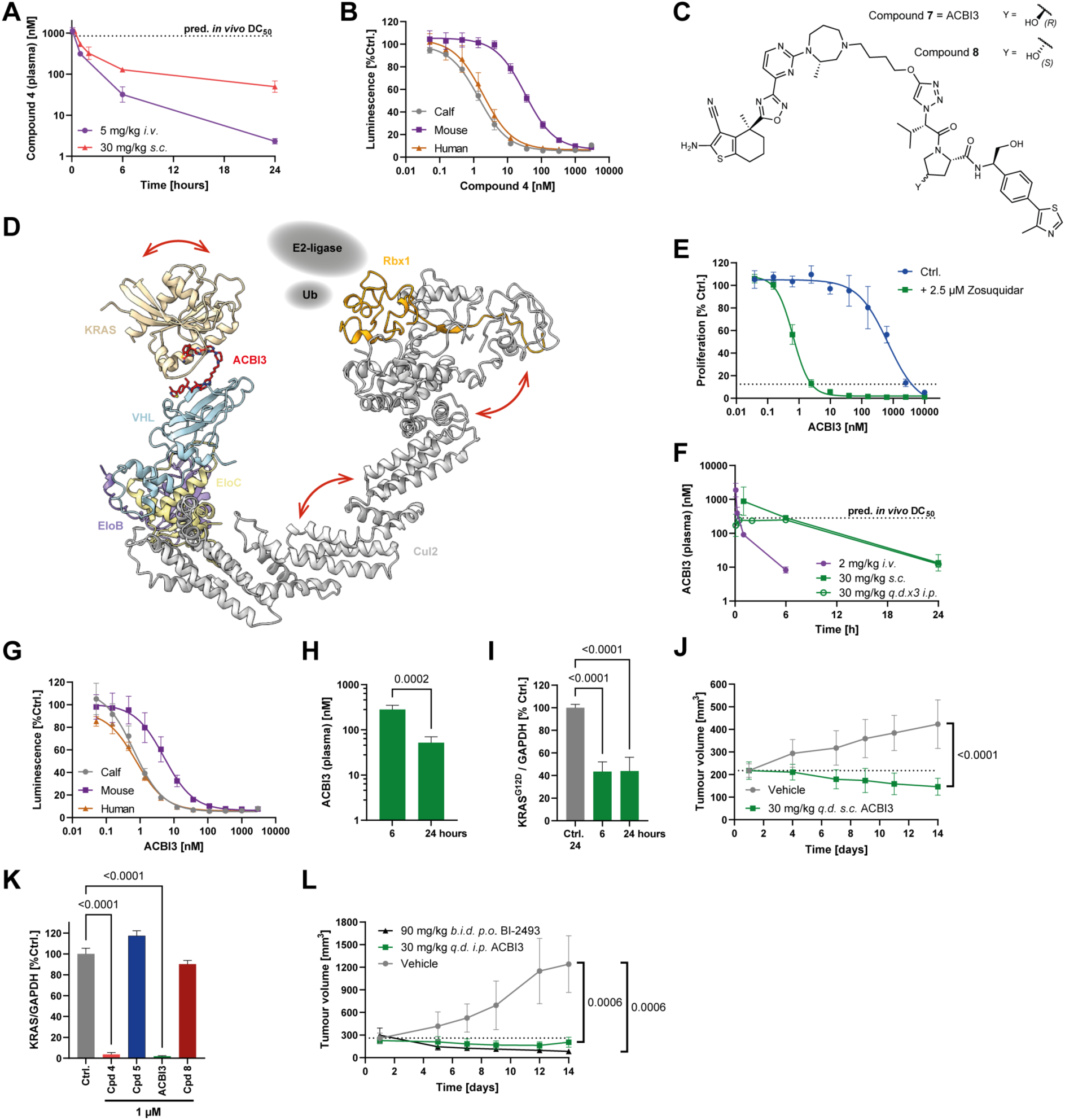
*In vivo* efficacy associated with degradation of KRAS^G12D^ and KRAS^G12V^. (**A**) Pharmacokinetic profile of compound **4** in mice upon *i.v.* or *s.c.* dosing, 2-hydroxypropyl-β-cyclodextrin (HP-β-CD) formulation (N=3, geometric mean and geometric SD). (**B**) Degradation of HiBiT-KRAS^G12D^ (24 hours, N=4, SD) by compound **4** in GP5d cells in presence of 10% fetal calf, mouse or human serum. (**C**) Chemical structure of ACBI3. (**D**) Cryo-EM structure of the KRAS:ACBI3:VCB:Cul2:Rbx1 heptameric complex, major motions indicated as red arrows (Cul2 breathing, KRAS rocking and Rbx1 twisting), expected positions of the E2-ligase and ubiquitin added schematically in dotted ellipses. (**E**) Proliferation of ACBI3 in LS513 cells in presence and absence of 2.5 µM Zosuquidar (5 days, N=3). (**F**) Pharmacokinetic profile of ACBI3 in mice upon *i.v.* (HP-β-CD), *s.c.* or *q.d.x3 i.p.* dosing (N=3, geometric mean and geometric SD). (**G**) Degradation of HiBiT-KRAS^G12D^ (24 hours, N=4, SD) in GP5d cells in presence of 10% fetal calf-, mouse- or human serum. (**H**) ACBI3 plasma levels in subcutaneous GP2d tumor bearing mice dosed with 30 mg/kg ACBI3 *q.d.x3* (N=5, geometric mean and geometric SD, Welch’s t-test). (**I**) *In vivo* degradation of KRAS^G12D^ upon *s.c. q.d.x3* dosing of 30 mg/kg ACBI3. (N=5, mean and SD). (**J**) *In vivo* efficacy of ACBI3 in a GP2d xenograft model (N=10, mean and SD, Wilcoxon test). (**K**) *In vitro* degradation of murine KRAS in B16F10 cells by 1 µM compound **4**, ACBI3 and corresponding inactive stereoisomers compound **5** and -**8** by capillary electrophoresis (24 hours, N=3, mean and SD, One-way ANOVA, Dunnet correction). (**L**) *In vivo* efficacy of ACBI3 in a RKN xenograft model (N=7, mean and SD, Wilcoxon text day 14 (N=4 control-, N=7 treatment groups).

Seeking to achieve *in vivo* active concentrations and to maintain the productive elements of ternary complex molecular recognition observed in compounds **3** and -**4**, we swapped the isoxazole for a triazole and introduced a hydroxymethyl group at the benzylic position of the VHL binder to obtain compound **7,** herein referred to as ACBI3 (Fig. 4C). While slightly improving solubility (4 µg/mL *vs* < 1 µg/mL at pH 4.5, table S3) SPR (fig. S8A) and FP (fig. S8B) suggested remarkable VHL engagement for ACBI3, with high signal detected via SPR beyond >1 000 s resulting in an apparent biophysical half-life of its binary complex with VHL of > 2 000 s (fig. S8A, table S1). Aiming to measure ternary complex stability, we noted that the FP assay reached the limits of quantification due to ternary *K_D_* approaching that of the assay probe (fig. S8B). However, by SPR, ternary complex *K_D_* was quantifiable and 4-fold improved *vs* compound **4** (6 nM for ACBI3 *vs* 26 nM for compound **4**) (fig. S8C, table S1). Single particle cryo-electron microscopic analysis of the KRAS^G12V^:ACBI3:VCB:Cul2:Rbx1 complex (fig. S9 and S10) revealed significant flexibility throughout the entire complex (Fig. 4D), likely necessary for successful ubiquitin transfer.

While high-resolution modelling of ACBI3 binding was limited by a high flexibility of KRAS, we clearly observed the density of the VHL-binding part of ACBI3, as well as the connecting density of the linker (fig. S11A and B). Overlay with a 2.2 Å resolution ternary co-crystal structure of KRAS^G12V^:ACBI3:VCB (PDB 8QVU) supports the same overall binding architecture via both techniques (Fig. 4D, fig. S11C). Details of the PROTAC binding site in the X-ray structure were resolved revealing a flexible binding mode overall consistent with those of compound **3** and compound **4** (fig. S11D). ACBI3 maintains the triazole-Tyr112 π-stack as observed with the isoxazole of compound **4**, while engaging an additional H-bond interaction between the newly incorporated benzylic hydroxy group and Gln99 in KRAS^G12V^ (fig. S11D). Multiple strong protein-protein interactions were apparent in the crystal structure which differed between protomers, indicating that, as with compound **3** and compound **4,** ACBI3 ternary complexes can dynamically transition between different H-bond networks between VHL and KRAS^G12V^.

ACBI3 exhibited potent intracellular VHL engagement, ternary complex formation and ubiquitination translating into potent E3-ligase dependent cellular degradation and proteome-wide selectivity comparable to compound **4** (fig. S12A-H, table S2). Similarly, kinetic degradation parameters of ACBI3 (fig. S12I-L, table S2) and KRAS allele degradation specificity (fig. S13A) were comparable to compound **4**. Testing the antiproliferative activity in a cell line panel revealed that ACBI3 was broadly active on *KRAS* mutant *vs KRAS^WT^* cell lines (geometric mean *IC_50_* = 478 nM *vs* 8.3 µM, respectively) (fig. S13B). Similar to compound **4**, ACBI3 exhibited high efflux in the Caco-2 assay (table S3).

Hence, we also tested the activity of ACBI3 in cell lines in the presence of Zosuquidar (fig S13C). Inhibition of drug efflux resulted in 1 000-fold increased antiproliferative potency of ACBI3 (Fig. 4E) but not the KRAS inhibitor BI-2493 (fig. S13D) in LS513, a cell line with high *ABCB1* expression. The overall sensitivity pattern in the cell line panel was comparable in the presence or absence of Zosuquidar (fig. S13B and C). On a global scale, we observed an average 5-fold shift in potency attributable to efflux transporter expression in KRAS mutant cell lines (fig. S13E). This analysis likely underestimates the full antiproliferative activity of both KRAS inhibition and degradation as several *KRAS* mutant cell lines display more prominent KRAS dependency under anchorage independent (3D) conditions (*6*) not amenable to the applied high-throughput approach employed here (fig S13F-G). We conclude that, in addition to compromising oral bioavailability, ABCB1-mediated efflux creates the need for compensation by higher doses of ACBI3 in ABCB1 expressing tumors.

To establish *in vivo* proof of concept, we formulated ACBI3, which lacks oral bioavailability, in PEG-400 / Transcutol / Kolliphor HS 15 (*27*) to enable multiple *s.c.* daily dosing studies with a delayed absorption profile. *S.c.* administration of 30 mg/kg ACBI3 in this formulation resulted in a plasma concentration-time-profile (Fig. 4F) covering the *in vivo DC_50_* of 281 nM—predicted based on degradation potency shift assays (Fig. 4G) and the degradation potency of ACBI3 in GP2d cells in 10% FCS (*DC_50_* = 3.9 nM) (table S2)—for around 6 hours. Administering ACBI3 to GP2d tumor bearing mice, we observed degradation of KRAS^G12D^ (44% of untreated controls) in tumors consistent with exposures covering the *DC_50_* for at least 6 hours *in vivo* (Fig. 4H and I). While exposures varied 5-fold between the 6 and 24 hours timepoints (Fig. 4H), KRAS^G12D^ levels did not recover to a notable extent consistent with extending the pharmacodynamic efficacy beyond the pharmacokinetic presence of ACBI3 at concentrations covering the predicted *in vivo DC_50_*. In an anti-tumor efficacy study in GP2d tumor bearing mice, 30 mg/kg ACBI3 dosed daily *s.c.* for up to 14 days resulted in pronounced tumor regressions (Fig. 4J) with a tumor growth inhibition of 127% and a significant difference of control *vs* treatment group (*p* < 0.0001). This demonstrates that the PROTAC mediated suppression of oncogenic KRAS observed *in vitro* translates into *in vivo* regressions in KRAS mutant tumor bearing mice. Pleasingly, we also noted no impact on body weight (fig. S13H) suggesting selective degradation of KRAS is systemically tolerated in mice. Of note, ACBI3 degrades murine KRAS *in vitro* and *in vivo* (Fig 4K and fig S13I and -J). While systemically well tolerated, we observed skin lesions in mice of *s.c.* efficacy studies and thus do not recommend further use of this formulation and route of administration for *in vivo* experiments. We therefore moved to intraperitoneal (*i.p.*) delivery of ACBI3 formulated as a nano-milled suspension which upon repeated dosing resulted in exposures covering the predicted *in vivo DC_50_* for six hours and were tolerated (Fig. 4F). Attesting to the antitumor activity of ACBI3 beyond a single KRAS mutant, *i.p.* administered ACBI3 induced regressions *in vivo* in the *KRAS^G12V^ TP53^R175H^* mutant RKN xenograft model (Fig. 4L). Of note, a daily dose of 180 mg/kg of BI-2493 also induced regressions in this model (Fig. 4L). Together these data provide first pre-clinical therapeutic proof-of-concept for pan-KRAS degradation.

## Conclusion

Activating mutations in KRAS are prevalent in patients suffering from solid tumours with unmet medical need. For example, 35% of lung, 45% of colorectal, and up to 90% of pancreatic cancers are associated with KRAS mutations (*2*). This amounts to an incidence of approximately 150 000 new cases in the United States for these three tumor types (*28*). An inherent challenge to targeting KRAS with small molecules is the wide range of mutations leading to oncogenic activation. This in turn creates a challenge to identify small molecules that can broadly address the different mutations with potencies and exposures required for clinical efficacy. Despite intense research investment over decades, so far only allele specific inhibitors have been approved and the need for potent KRAS selective targeting concepts with broad mutation coverage remains high.

Here, we establish degradation of a broad spectrum of oncogenic KRAS variants with a single agent achieving potent and long-lasting suppression of oncogenic signaling *in vitro* and *in vivo*. ACBI3 is a selective, potent and *in vivo* active pan-KRAS degrader discovered via a structure-based design approach guided by optimization of VHL:PROTAC:KRAS ternary complex stability and durability. This optimization allowed us to achieve a degrader that effectively acts on the majority of KRAS mutants with high prevalence in cancer patients, and as a result inhibits proliferation in KRAS mutant cell lines covering a wide range of tumor types. KRAS degradation via ubiquitin ligase recruitment enables a greater than 10-fold higher potency compared to target inhibition and results in prolonged suppression of MAPK signaling. ACBI3 is a first example of a single agent capable of selectively degrading a major share of highly prevalent oncogenic KRAS variants. We also establish that ACBI3 suppresses oncogenic KRAS protein levels *in vivo* beyond its pharmacokinetic presence, ultimately resulting in tumor regression. Identifying approaches to dose ACBI3 safely in mice qualifies this molecule for use in preclinical *in vivo* tumor models. Bifunctional degraders rely on engaging the E3 ligase machinery with a dedicated binding motif which may be associated with ubiquitin proteasome system related resistance mechanisms (*29*) and increased efflux liability (*30*). However, the demonstrated ability of ACBI3 to potently target a wide spectrum of KRAS variants opens up opportunities for degraders to address susceptibility to resistance associated with on-target KRAS mutations.

ACBI3 illustrates many key features specific to degraders made of reversible target binders, such as catalytic mode of action independent of occupancy driven pharmacology. This more broadly suggests fundamental advantages for targeting a spectrum of chemically heterogenous disease-relevant protein variants by targeted protein degradation. We anticipate that pan-KRAS degradation will deliver a new path to address a broad sweep of malignancies with high unmet medical need.

## Acknowledgments

We would like to acknowledge technical and experimental support by Géraldine Garavel, Peter Greb, Karin Stefanie Hofbauer, Christoph Peinsipp, Yvonne Scherbantin, Florian Schiel (chemical syntheses), Sandra Doebel, Michael Galant, and Patrick Werni (molecular cloning, protein production and purification of KRAS proteins), Aldina Crnic, Simone Lieb, Corinna Melichar, Ana Orsolic (cellular biology assays), James Habineza, Sonja Porits, Katrin Gitschtaler (in vivo pharmacology), Nina Braun and Romy Schopper (Cellular assay support), Bernadette Sharps, Thomas Gerstberger and Niklas Baumann (Biochemical assay support), Klaus Rumpel (biophysics assay support), Ines Truebenbach, Ida Dinold, Philipp Toplak (formulation development), Angelika Hörschläger (targeted proteomics) - all from Boehringer-Ingelheim. We acknowledge Abdel Atrih and Douglas Lamont from the University of Dundee FingerPrint Proteomics facility for support with mass spectrometry proteomics. We also acknowledge Dario Alessi from MRC PPU, University of Dundee, for the gift of mouse embryonic fibroblasts.

## Funding

We thank European Synchrotron Radiation Facility and Diamond Light Source for beamtime (BAG proposal MX14980) and for support at beamlines I23-1 and I24 respectively. This work has received funding from Boehringer Ingelheim and the Austrian Promotion Agency (grant numbers 871904 and 900397). Biophysics and drug discovery and proteomics/computing activities at Dundee were partially supported by Wellcome Trust strategic awards (100476/Z/12/Z, 094090/Z/10/Z and 097945/C/11/Z respectively).

## Author contributions

J.P., W.F., C.K. and A.G. contributed equally and will be putting their name first on the citation in their CVs. A.C. and P.E. co-conceived the study, designed and supervised experiments and co-wrote the paper. J.P. and W.F. designed, analysed and supervised experiments, prepared figures and co-wrote the paper. C.K. and A.G. designed compounds and synthetic routes and co-wrote the paper. E.D., A.B.F., J.G.-T., R.M., N.T., H.W. and S.Z. designed compounds and synthetic routes. J. K-O., T.K., O.P. synthesised key compounds. M.K. co-wrote the manuscript. G.F., S.A-S., D.P. and P. S-B. contributed to production and analysis of cryo-EM data. R.K. performed small molecule structure identification and analysis. B.K., P.P., C.H. and A.L. designed and performed phosphoproteomics studies. B.K. produced binary X-ray structures. A.W. and D.Z. crystallised ternary complexes KRAS:PROTAC:VCB and determined their co-crystal structures. K.K. and A.W. designed and performed biophysical assays. S.K., C.W., V.V., S.O. and V.R., designed, interpreted and performed cellular assays. R.M. and V.V. designed, analysed and performed proteomics studies. N.M. designed and interpreted targeted proteomics and co-wrote the manuscript and N.M. and H.A. designed in vivo PK and in vitro ADME studies. M.W., P.S., E.S., S.B. and G.B. designed and interpreted in vivo studies and co-wrote figures. J.Q. measured effects of P-gp substrate properties on cancer cell line panel data.

## Competing interests

J.P., A.G., C.K., G.F., M.W., N.M., C.H., H.A., S.A-S., S.B., G.B., J.K-O., T.K., M.K., R.K., B.K., K.K., A.A.L, S.O., D.P., O.P., V.R., P.S-B., P.S., E.S., J.Q., H.W., P.E. are current or former employees of Boehringer Ingelheim. A.C. is a scientific founder, shareholder and advisor of Amphista Therapeutics, a company that is developing targeted protein degradation therapeutic platforms. The Ciulli laboratory receives or has received sponsored research support from Almirall, Amgen, Amphista Therapeutics, Boehringer Ingelheim, Eisai, Merck KaaG, Nurix Therapeutics, Ono Pharmaceuticals and Tocris-Biotechne.

## Data and materials availability

Atomic coordinates and structure factors for X-ray crystallographic and cryo-electron microscopy structures have been deposited in the Protein Data Bank (and the EMDB) with accession codes 8QUG, 8QU8 (EMD-18657), 8QW6, 8QW7 and 8QVU. All raw data and search results for proteomics studies have been deposited to the ProteomeXchange Consortium with the dataset identifiers PXD046161, PXD045416 and PXD045460, respectively.

## Supplementary Materials

Materials and Methods

Figs. S1 to S8

Tables S1 to S6

Chemical synthesis procedures

ACBI3 proliferation data

## Materials and Methods

### Chemical synthesis

Full details of synthetic procedures and NMR spectra of final compounds are provided as a separate file: Supplementary Information_Synthesis.

### Protein Production

Wild-type and mutant versions of human proteins were used for all protein expression, as follows: VHL (UniProt accession number P40337), ElonginC (Q15369), ElonginB (Q15370) and KRAS G12C, G12D and G12V variants of KRAS C118S (P01116, residues 1-169, hereafter referred to as KRAS^G12C^, KRAS^G12D^ and KRAS^G12V^, respectively).

The VCB complex was expressed and purified as described previously (*31*). Briefly, N-terminally His_6_-tagged VHL (54–213), Elongin C (17–112) and Elongin B (1–104) (Addgene ID 204500 & 204501) were co-expressed in *E. coli* and the complex isolated by Ni-affinity chromatography. The His_6_-tag was removed using TEV protease, and the complex was further purified by anion exchange and size-exclusion chromatography (SEC).

N-terminally His_6_-tagged KRAS^G12D/G12V^ was expressed and purified as described previously (*32*). In brief, the protein was overexpressed in *E. coli* in Terrific Broth (TB) supplemented with 0.2 mM IPTG for expression induction. Expression was performed overnight at 18 °C. The protein was then purified by Ni-affinity chromatography, followed by His_6_-tag removal using TEV protease, another round of Ni-affinity chromatography to remove the His_6_-tag and the uncleaved protein, and SEC.

### Protein Crystallography

VCB, compound **3** and KRAS^G12V^ GDP were mixed in a 1:1.1:1 stoichiometric ratio in 20 mM HEPES, pH 8.0, 100 mM sodium chloride, 1 mM TCEP, 2% DMSO, 1 mM GDP incubated for 30 min. on ice. The complex was concentrated to 8 mg/mL. The drops were prepared by combining 200 nL of the ternary complex with 200 nL of well solution (200 mM lithium sulfate monohydrate, 100 mM BIS-TRIS propane pH 5.5, 25% w/v polyethylene glycol 3350) and crystallized at 4 °C using the sitting-drop vapor diffusion method. Crystals were grown for 27 days then cryoprotected by addition of ∼500 nL well solution supplemented with 20% (v/v) glycerol to the crystallization drop followed by harvesting and flash cooling in liquid nitrogen. VCB, compound **4** and KRAS^G12V^ GDP were mixed in a 1:5:5 stoichiometric ratio in 20 mM HEPES, pH 8.0, 100 mM sodium chloride, 1 mM TCEP, 2% DMSO, 1 mM GDP incubated for 30 min. on ice before purification by size exclusion chromatography (SEC) to isolate the formed ternary complex. The complex was concentrated to 10 mg/mL. The drops were prepared by combining 200 nL of the ternary complex with 200 nL of well solution (200 mM sodium citrate, 100 mM BIS-TRIS propane pH 8.5, 20% w/v polyethylene glycol 3350) and crystallized at 4 °C using the sitting-drop vapor diffusion method. Crystals were grown for 5 days then cryoprotected by addition of ∼500 nL well solution supplemented with 20% (v/v) glycerol to the crystallization drop followed by harvesting and flash cooling in liquid nitrogen.

VCB, ACBI-3 and KRAS^G12V^ GDP were mixed in a 1:1:1.5 stoichiometric ratio in 20 mM HEPES, pH 8.0, 100 mM sodium chloride, 1 mM TCEP, 2% DMSO, 1 mM GDP incubated for 30 min. on ice prior SEC to isolate the formed ternary complex. The eluted complex was concentrated to a final concentration of approximately 5-10 mg/mL. The drops were prepared by combining 200 nL of the ternary complex with 200 nL of well solution and crystallized at 4 °C using the hanging-drop vapor diffusion method. Crystals were grown in 20% w/v PEG 8000, 0.2 M lithium chloride, 0.1 M Tris pH 8.0 for several days until they reached a suitable size. Harvested crystals were flash cooled in liquid nitrogen following gradual equilibration into cryoprotectant solution consisting of 30% (v/v) glycerol in 20% PEG 8000, 0.2 M lithium chloride, 0.1 M Tris, pH 8.0.

Diffraction data for the ternary complex crystals were collected at beamline X10SA at the Swiss Light Source, Switzerland at a wavelength of 1.0 Å. Images were processed using autoPROC (*33*), the phase problem solved using PHASER (*34*) using previously determined structures, the model built using Coot (*35*) and iteratively refined using Phenix (*36*). The structures were deposited in the PDB with codes 8QW6, 8QW7 and 8QVU.

The binary crystal structure of KRAS^G12C^ (UniprotID P01116) with compound **1** was obtained through back-soaking. The hexagonally shaped source crystals were obtained by hanging drop co-crystallization of 0.75 µL of 43 mg/mL KRAS^G12C^ with a mixture of 0.75 µL of 2 mM dimethyl-aminocyanothiophen in 0.1M sodium acetate pH 5.2, 1.55M ammonium sulfate. The crystals were transferred to a 1 μL drop containing 10 mM compound **1** in 0.1 M sodium acetate pH 5.2, 1.55 M ammonium sulfate and incubated for 48h. Subsequently, the crystals were cryocooled in liquid nitrogen, using 25% ethylene glycol as cryoprotectant.

Data were collected at beamline ID23-1 at the European Synchrotron Facility, France at a wavelength of 0.97 Å (compound **1**). Images were processed using autoPROC (*33*), the phase problem solved using PHASER (*34*) using a previously determined structure, the model built using Coot (*35*) and iteratively refined using autoBUSTER, CCP4 (*34*) and Phenix (*36*). The structure has been deposited in the PDB with code 8QUG (compound **1**). Ramachandran analysis showed 95.6% of residues in favored regions, 4.24% in allowed regions and no outliers.

### Cryo-Electron Microscopy

The ternary complex was prepared by mixing 2.15 µL of 48 mg/mL KRAS^G12V^ solution, 5.35 µL of ACBI3 and 318 µL of 1.25 mg/mL VHL-ElonginB-ElonginC-Cullin2-Rbx1-complex obtained from Abcam, UK (catalogue no. ab271787, lot no. GR3409294-5), incubating at room temperature for 10 min., concentrating the sample to 70 μL using a 10MWCO Amicon Ultra concentrator followed by size exclusion chromatography Superdex 200; running buffer 50 mM Tris pH=7.5, 100 mM NaCl, 0.5 mM TCEP, 2 mM MgCl_2_) (fig S9A and B). After elution, 4 μL (0.9 mg/mL) of the heptameric complex peak were immediately applied to a plasma-cleaned (Pelco EasyGlow, 45 s, 15 mA) Quantifoil 1.2/1.3, 200 mesh copper grid and blotted (TedPella 595 filter paper, blotting time 2 s) and vitrified in liquified ethane using an automated plunge freezer (Vitrobot Mk4, Thermo Fisher Scientific; humidity 95%, 4 °C).

Data were collected on a Titan Krios G4 equipped with a Falcon 4 camera (Thermo Fisher Scientific) and a Selectris energy filter (10 eV) at an electron acceleration voltage of 300 kV and a nominal magnification of 165.000x, corresponding to a pixel size of 0.745 Å/px. 7 634 images were acquired in counting mode using EPU data collection software (Thermo Fisher). Data were processed (fig S9C) using the cryoSPARC v4 suite (*38*). Data were motion- and CTF-corrected during online-processing with cryoSPARC live (Patch Motion Correction, Patch CTF Estimation) and bad images rejected based on defocus and motion outliers. 2 414 422 particles were automatically identified (blob-based picking followed by template-based picking) and extracted at a box size of 512 px from 6 907 images. Subsequently, 2 rounds of 2D classification (cryoSPARC , resolution 6 & 3 Å, 40 online-EM iterations, batch size/class 400) carried out, followed by ab-initio reconstruction (3 classes), 3D-heterogenous refinement (3 classes as identified by ab-initio), non-uniform refinement of the most promising class hereof (217 928 particles) and 3D variability analysis (3 modes) (*39*). The modes determined herein were used as initial latent input indices for 3D flex refinement (*40*) with 217 000 particles using a tetrapartite flex mesh mask (KRAS, VHL/EloB/EloC, Cul2-helical repeat region, Rbx1+Cul2-head region), which was followed by DeepEMhancer (*41*) for map sharpening.

Model building and refinement was based on the ternary crystal structure presented in this paper (PDB entry 8QVU) for KRAS, VHL, Elongin B and Elongin C, and PDB entry 5N4W for Cul2 and Rbx1 (*42*). Initial docking and figures were performed using ChimeraX (*43*), while subsequent model building was performed using Coot (*35*). This was followed by real-space refinement with PHENIX (*36*), where the KRAS- and Rbx1-chains were heavily restrained to their input X-ray-models, as densities in this region were low resolution and hence only sufficient to provide the approximate orientation of the proteins. The structure has been deposited in the PDB and EMDB with accession codes 8QU8 and EMD-18657 respectively.

### Surface Plasmon Resonance Experiments

Surface plasmon resonance experiments were performed on Biacore 8K instruments (Cytiva). Streptavidin (Prospec) was immobilized at 25 °C on CM5 Chips (Cytiva) using 10 mM HBS-P+ buffer (pH 7.4) (Cytiva). The surface was activated using 400 mM 1-ethyl-3-(3-dimethylaminopropyl)-carbodiimide and 100 mM N-hydroxysuccinimide (Cytiva) (contact time 420 s, flow rate 10 mL/min). Streptavidin was diluted to a final concentration of 1 mg/mL in 10 mM sodium acetate (pH 5.0) and injected for 600 s. The surface was subsequently deactivated by injecting 1 M ethanolamine for 420 s and conditioned by injecting 50 mM NaOH and 1 M NaCl. Dilution of the biotinylated target proteins and coupling was performed using a running buffer without DMSO. The two KRAS target proteins were prepared at 0.1 mg/mL and coupled to the chip to a density between 200 and 800 response units. All KRAS binary interaction experiments were performed at 25 °C in running buffer (20 mM Tris(hydroxymethyl)aminomethane, 150 mM potassium chloride, 2 mM magnesium chloride,

2 mM Tris(2-carboxyethyl) phosphine hydrochloride, 0.005% Tween20, 40 μM Guanosine 5′-diphosphate, pH 8.0, 1% DMSO). The compounds were diluted in running buffer and injected over the immobilized target proteins (concentration range, 3.33-1 000 nM). Sensorgrams from reference surfaces and blank injections were subtracted from the raw data before data analysis using Biacore Insight software. Affinity and binding kinetic parameters were determined by using a 1/1 interaction model, with a term for mass transport included.

For VCB immobilization, sensor chips pre-coated with streptavidin (Cytiva) were used. Biotinylated Avi-tagged-VCB was prepared as previously described. Excess free biotin was removed by extensive dialysis (3 times) into buffer containing 20 mM HEPES pH 7.5, 200 mM NaCl and 0.25 mM TCEP. All interaction experiments involving VCB were done at 20 °C in running buffer (20 mM HEPES, pH 8.0, 150 mM potassium chloride, 2 mM magnesium chloride, 2 mM TCEP, 0.05% Tween20, 2% DMSO). The compounds were diluted in running buffer and injected over the immobilized target proteins (concentration range, 3.33-1 000 nM). For ternary complex measurements, experiments were run in the presence of 2 µM KRAS^G12D^-GDP during the injection phase. Data analysis was performed as described above.

### Fluorescence Polarization Experiments

All FP measurements were taken using a PHERAstar FS (BMG LABTECH) with fluorescence excitation and emission wavelengths (λ) of 485 nm and 520 nm, respectively. FP competitive binding assays were performed in duplicate on 384-well plates (#3575, Corning) with a total well volume of 15 μL. Each well solution contained 5 nM of FAM-labelled HIF-1α peptide (FAM-DEALAHypYIPMDDDFQLRSF, *K_D_* = 3 nM as measured by a direct FP titration), 15 nM of VCB protein, and decreasing concentrations of PROTACs (13-point serial 2-fold dilutions starting from 10 μM) or PROTAC:KRAS^G12D^-GDP (13-point serial 2-fold dilution starting from 10 μM of the PROTAC and constant concentration of KRAS^G12D^-GDP at 20 μM) in 100 mM HEPES pH 7.5, 100 mM NaCl, 1 mM TCEP with a final DMSO concentration of 2%. To obtain percentage of displacement, control wells containing peptide in the absence of protein (maximum displacement), or VCB and peptide with no compound (zero displacement) were also included. These values were then fitted by nonlinear regression using Prism (GraphPad, version 7.03) to determine average *IC_50_* values and standard error of the mean (SEM) for each titration. A displacement binding model was used to back-calculate inhibition constants (KI) from the measured *IC_50_* values, as described previously (*44*).

### Cell Culture

Cell lines were obtained through ATCC or ECACC, verified for identity by satellite repeat analysis, tested for mycoplasma contamination at regular intervals, and cultured in the specified media in a humidified cell incubator at 37 °C and 5% CO_2_, unless specified otherwise. Eagle’s minimal essential medium (EMEM) was obtained from ATCC (#30-2003), Leibovitz’s L15 medium from Themo Fisher (#11415064), McCoy’s 5A medium from Thermo Fisher (#16600082), Ham’s F-12K medium from Gibco (#21127-022), high glucose Dulbecco’s modified Eagle medium (DMEM) from Sigma (#D6429), RPMI-1640 Glutamax from Gibco (#61870) and RPMI-1640 from Gibco (#A10491). The following cell lines (product codes and culture media / conditions in parentheses) were used for the described experiments: GP2d (ECACC No. 95090714, high glucose DMEM plus 10% FCS), GP5d (ECACC No. 95090715, high glucose DMEM plus 10% FCS), SW620 (ATCC No. CCL-227, Leibovitźs L15 plus 10% FCS, no CO_2_), NCI-H358 (ATCC No. CRL-5807 RPMI-1640 plus 10% FCS), HEK293 (ATCC No. CRL-1573, EMEM plus 10% FCS), MIA PaCa-2 (ATCC No. CRL-1420, high glucose DMEM plus 10% FCS), HCC461 (Cellosaurus CVCL_5135, RPMI-1640 plus 5% FCS), A549 (ATCC No. CCL-185, Ham’s F-12K medium plus 10% FCS), GEO (obtained as a gift from Prof. Stefano Pepe, Universita degli Studi di Napoli Federico II, McCoy’s 5A medium plus 10% FCS plus 20 mM HEPES), HS766T (ATCC No. HTB-134, high glucose DMEM plus 10% FCS), HCT116 (ATCC No. CCL-246, McCoy’s 5A medium plus 10% FCS), LS513 (ATCC No. CRL-2134, RPMI-1640 Glutamax plus 10% FCS), RKN (JCRB No. IFO50317, Ham’s F-12K medium plus 10% FCS), B16F10 (ATCC No. CRL-6475, high glucose DMEM plus 10% FCS), mouse embryonic fibroblasts (generated in Dario Alessi Lab, MRC PPU, University of Dundee; high glucose DMEM plus 10% FCS) and A375 (ATCC No. CRL-1619, high glucose DMEM plus 10% FCS).

### Cell Proliferation

For 2D proliferation assays, cells were seeded at 500 cells per well in 40 μL of growth medium in a white bottom opaque 384-well plate and allowed to grow overnight. For 3D proliferation assays, cells were seeded in 96-well clear round bottom ultra-low attachment microplates (Costar #7007) at a density of 2 000 cells per well in growth media supplemented with 2% FCS and allowed to equilibrate overnight.

To obtain starting densities, a set of cells seeded in parallel were lysed and measured at the time of compound addition using the CellTiter-Glo cell viability reagent (Promega #G7570) per well as per the manufacturer’s recommendation. The compounds were added to the cells at logarithmic dose series using a HP Digital Dispenser D300 (Tecan), normalizing for added DMSO. After compound addition, the cells were incubated for five days (or seven days for 3D growth assays) and the viability measured using the CellTiter-Glo reagent as described above. The results are stated as mean and standard deviation of triplicate experiments. *IC_50_* values including 95% confidence intervals were computed using a 3-parametric logistic model restricting the bottom to be > 0.

### Luminescent caspase activity assay

Cells were seeded at a density of 10 000 cells per well in opaque white flat bottom 96-well plates with a clear bottom (Thermo Fisher #165306) in 100 µL medium and allowed to attach overnight. The next day, compounds were added to the cells at logarithmic dose series using a HP Digital Dispenser D300 (Tecan), normalizing for added DMSO. 48 hours after compound addition, 100 µL Caspase-Glo reagent (Promega #G8090) was added per well and caspase activity was determined by measuring luminescence as per the manufacturer’s recommendation. The results are stated as mean and standard deviation of triplicate experiments.

### Clonogenicity assays

Cells were seeded at 2 000 (A-375) or 5 000 (GP2d, NCI-H358, SW620) cells per well in 12-well plates in 800 µL medium and allowed to attach overnight. The next day, compounds were added as indicated and plates further incubated. One week after compound addition, the medium was changed and compounds re-added. Two weeks after seeding, the plates were carefully washed with PBS and stained for 10 min. at room temperature with 500 µL of a solution of 0.5% crystal violet (Sigma #C0775) in 1% methanol, 1% formaldehyde. Upon removal of the staining solution stained cells were washed with water and plates were scanned. For quantification, stained and dried cells were incubated with 33% acetic acid for 10 min. 100 µL of the supernatant was transferred to a clear bottom 96-well plate and absorption at 595 nm was recorded.

### Cell cycle analysis

Cells were seeded at 200 000 cells per well in 6-well plates in 3 mL medium and allowed to attach overnight. Compounds were added as indicated and cells further incubated for 48 hours. Non-adherent and adherent cells were harvested using TrypLE Express (Gibco #12604013) and washed with 1% BSA in PBS. Cells were fixed in 4% paraformaldehyde (BioLegend Cat. No. 420801, 100 µL per well for 10 min.) and stained in intracellular staining permeabilization wash buffer (BioLegend #421002, 100 µL per well) containing 1:2 000 DAPI (Thermo Scientific #62248, 1 mg/mL stock solution in water) for 10 min. in the dark at room temperature.

Subsequently, cells were sedimented by centrifugation and resuspended in 400 µL per well stain buffer (BD biosciences #554656). Cell cycle profiles were acquired using a BD FACSCanto II device (violet laser 405 nM, excitation 358 nM, emission 457 nM) and BD FACS DIVA 9.0.1 software. Cell cycle phases were assigned applying the Dean-Jett-Fox algorithm and the FlowJo 10.9.0 software.

### High-throughput Proliferation Screen

High throughput proliferation profiling in presence or absence of the ABCB1 inhibitor zosuquidar (2.5 µM) was performed at Horizon Discovery. Briefly, the cells were seeded in 25 μL of growth media in black 384-well tissue culture plates at the density defined for the respective cell line and plates were placed at 37 °C, 5% CO_2_ for 24 hours before treatment. At the time of treatment, a set of assay plates (which did not receive treatment) were collected and ATP concentrations were measured by using CellTiter-Glo v.2.0 (Promega) and luminescence reading on an Envision plate reader (Perkin Elmer). Compound **4** was transferred to assay plates using an Echo acoustic liquid handling system. Assay plates were incubated with the compound for 5 days and were then analyzed by using CellTiter-Glo. All data points were collected by means of automated processes and were subject to quality control and analyzed using Horizon’s proprietary software. Horizon uses growth inhibition as a measure of cell growth. The growth inhibition percentages were calculated by applying the following test and equation: if T < V0 then 100 (1 − (T − V0)/V0) and if T ≥ V0 then 100 (1 − (T − V0)/(V − V0)), where T is the signal measure for a test article, V is the untreated or vehicle-treated control measure and V0 is the untreated or vehicle-treated control measure at time zero (colloquially referred to as T0 plates). This formula was derived from the Growth Inhibition (GI) calculation used in the National Cancer Institute’s NCI-60 high-throughput screen.

### KRAS Degradation Assays via Detection of Split Nanoluciferase (HiBiT)

To assess PROTAC-mediated degradation of HiBiT-tagged KRAS constructs, cells were seeded at 25 000 cells per well in culture medium into white bottom opaque 96-well plates (Perkin Elmer cat no. 5680). Plates were incubated at 37 °C, 5% CO_2_ in a humidified incubator over night to allow the cells to adhere. Test compounds (10 mM stock in DMSO) were added at logarithmic dose series using the HP Digital Dispenser D300 (Tecan), normalizing for added DMSO. Plates were further incubated at 37 °C for 18 hours. Following incubation, 100 µL per well Promega Nano-Glo HiBiT lytic detection reagent mix (Promega Nano-Glo HiBiT Lytic Detection System #N3050), prepared according to the manufactureŕs instructions in the kit, were added. To allow for adequate cell lysis, plates were incubated on an orbital shaker for 15 min. and further incubated 30 min. at room temperature. Upon completion of cell lysis, luminescence was measured using an Envision plate reader using the Ultrasensitive Luminescence Protocol for 96-well plates. Luminescence levels were normalized by the values obtained with DMSO-treated samples and plotted as percent of DMSO control. *DC_50_* values were computed using a three parametric logistic model. *D_max_* values represent the maximal extent of degradation observed and are stated as percent of control treatments.

### GP5d Cells With HiBiT-tagged KRAS Locus

Genetic introduction of a split luciferase tag into the KRAS locus was conducted at Horizon Discovery (https://horizondiscovery.com). Briefly, a HiBiT protein detection tag (amino acid sequence VSGWRLFKKIS) was introduced immediately downstream of the initiating Methionine codon of the endogenous KRAS locus (Ensembl gene ID ENSG00000133703.7) of GP5d cells (ECACC Cat. No. 95090715) by CRISPR-based genome engineering using a KRAS^G12D^ mutant donor construct encoding the HiBiT tag. This resulted in the heterozygous introduction of an N-terminal HiBiT tagged version of KRAS^G12D^ into the KRAS^WT^ allele. Correct modification of the KRAS locus was assessed by PCR-based genotyping and Sanger sequencing of the isolated PCR products. The resulting cell line is referred to as GP5d-HiBiT-KRAS^G12D^.

### Retroviral Transduction of HiBiT-tagged KRAS Mutants

The therapeutically relevant mutant KRAS constructs (WT, A146P, A146T, A146V, G12A, G12C, G12D, G12R, G12V, G13C, G13D, G13V, Q61E, Q61H, Q61K, Q61L, Q61P, Q61R) were obtained by site directed mutagenesis using a KRAS4B WT cDNA construct as a template at Genscript (https://www.genscript.com). GP5d cells (ECACC No. 95090715) expressing transgenic murine Slc7a1 to allow for transduction ecotropic lentiviral particles (Takarabio Cat. No. 631278) were transduced with lentiviral vectors expressing mutant KRAS4B cDNA under control of a CMV promoter. Stably transduced cells were selected using a neomycin selectable marker encoded on the construct.

### HiBiT-based Degradation Assay at 10% Mouse or Human Serum

GP5d cells expressing HiBiT-tagged KRAS^G12D^ were seeded at 25 000 cells per well in 100 µL medium into white bottom opaque 96-well plates. Plates were incubated at 37 °C, 5% CO_2_ in a humidified incubator overnight. The next day, incubation medium was removed and replaced with pre-warmed fresh DMEM medium containing 10% serum from NMRI female mice (without preservatives added, sterile filtered, supplied by Envigo RMS S.r.l. Z.I.) or human serum (Sigma cat. No. S7023). Compounds (10 mM stock in DMSO) were added at logarithmic dose series using a HP Digital Dispenser D300 (Tecan), normalizing for added DMSO. Plates were further incubated at 37 °C for 18 hours. Subsequently, media was removed and replaced with 100 µL PBS at room temperature. The luminescence reaction was started by adding 100 µL of Promega Nano-Glo HiBiT lytic detection reagent mix per well, which was prepared according to the manufactureŕs instructions. Plates were agitated for 15 min. on an orbital shaker and incubated for 30 min. at room temperature. Luminescence was measured on an Envision plate reader using an Ultrasensitive Luminescence Protocol for 96-well plates. Luminescence was normalized to values obtained with DMSO-treated samples and are reported as percentage of DMSO control. *DC_50_* values, *D_max_* values and 95% confidence intervals were computed using a 3-parametric logistic model.

### Kinetic Live Cell HiBiT Detection

LgBiT construct were diluted into Transfection Carrier DNA (Promega) at a mass ratio of 1:4 (w/w), after which FuGENE HD was added at a ratio of 1:3 (μg DNA: μL FuGENE HD). FuGENE HD complexes thus formed were combined with GP5d HiBiT KRAS^G12D^ cells suspended at a density of 2.5 × 10^5^ cells/mL, transferred to a tissue-culture treated flask, and incubated in a humidified, 37 °C/5% CO_2_ incubator for 20 hours. Following transfection, cells were plated in white 96-well tissue culture plates at a density of 5 × 10^4^ cells per well in 100 μL of growth medium and incubated overnight at 37 °C, 5% CO_2_. For continuous live cell detection out to 24 hours, an equal volume of CO_2_-independent medium (Gibco) containing a 2× concentration of Endurazine (Promega) was added to each well. Plates were incubated at 37 °C, 5% CO_2_, for 2.5 hours to allow luminescence to equilibrate before addition of each compound. Plate lid was removed and replaced with a Breathe-Easy sealing membrane. Plates were read for a period of 24 hours on the GloMax Discover (Promega) set to 37 °C. Degradation rate were calculated by fitting only the initial degradation portion of each kinetic concentration curve to the equation: Y = (Y0 – Plateau) * exp(-K * X) + Plateau, where K = degradation rate in units of hour^-1^.

### VHL Nanoluciferase (NanoLuc) Target Engagement (TE) assay

Target engagement and intracellular availability for VHL ligands were assessed using the NanoBRET TE Intracellular E3 ligase assay (Promega). HEK293 cells were transfected with the VHL-NanoLuc fusion constructs using FuGENE HD (Promega) according to the manufacturer’s protocol. Briefly, VHL-NanoLuc constructs were diluted into Transfection Carrier DNA (Promega) at a mass ratio of 1:9 (w/w), after which FuGENE HD was added at a ratio of 1:3 (μg DNA: μL FuGENE HD). FuGENE HD complexes thus formed were combined with HEK293 cells suspended at a density of 2 × 10^5^ cells/mL, transferred to a tissue-culture treated flask, and incubated in a humidified, 37 °C/5% CO_2_ incubator for 20 hours. Following transfection, cells were washed, harvested by trypsinization, and resuspended in Opti-MEM. BRET assays were performed in white, 96-well non-binding surface plates (Corning #3600) at a density of 2 × 10^4^ cells/well. All chemical inhibitors were prepared as concentrated stock solutions in DMSO (Sigma-Aldrich) and diluted in Opti-MEM (unless otherwise noted) to prepare working stocks. For measurement of intracellular affinity for VHL ligands, live cells were equilibrated for 2 hours with VHL tracer and test compound prior to BRET measurements. VHL tracer was prepared as a 100× intermediate stock at 100 μM in pure DMSO, and subsequently diluted to a working concentration of 20× in tracer dilution buffer (12.5 mM HEPES, 31.25% PEG-400, pH 7.5). VHL tracer was added to live cells at a final concentration of 1 μM from the 20× working stock. To measure BRET, NanoBRET NanoGlo Substrate and Extracellular NanoLuc Inhibitor (Promega) were added according to the manufacturer’s recommended protocol, and filtered luminescence was measured on a Pherastar equipped with 450 nm BP filter (donor) and 600 nm LP filter (acceptor), using 1 s integration time. Milli-BRET units (mBU) are calculated by multiplying the raw BRET values by 1 000. Competitive displacement data were then plotted with GraphPad Prism software and *IC_50_* values were determined using the [inhibitor] *vs* response – variable slope (four parameters) equation available in GraphPad Prism (Equation 1): Y = Bottom + (Top – Bottom) / (1 + (*IC_50_* / X)^HillSlope)

For measurement of intrinsic affinity for VHL ligands, the NanoBRET TE assay was applied as described above, with some changes: the cells were permeabilized by adding digitonin to a final concentration of 50 μg/mL, the final VHL tracer concentration was 0.25 μM, the permeabilized cells were incubated with tracer and test compound for 10 min. at room temperature, and the Extracellular NanoLuc Inhibitor was omitted from the NanoGlo substrate solution.

### Cellular NanoBRET Ubiquitylation and Ternary Complex Experiments

GP5d clonal cells (7 × 10^5^) expressing the HiBiT-KRAS were transfected with FuGENE HD (Promega) and 1 μg of HaloTag-VHL, or HaloTag-UBB in 6-well plates. The following day, 5 × 10^4^ transfected cells were replated into white 96-well tissue culture plates in the presence or absence of HaloTag NanoBRET 618 Ligand (Promega) and incubated overnight at 37 °C, 5% CO_2_. The following day, cells were treated with compound for 4 hours. Following treatment time point, NanoBRET NanoGlo reagent (Promega) was added according to the manufacturer’s recommended protocol, and filtered luminescence was measured on a Pherastar with 450 nm BP filter (donor) and 600 nm LP filter (acceptor), using 1 s integration time. Background subtracted NanoBRET ratios expressed in milliBRET units were calculated from the equation: mBRET_ratio = ((acceptor_channel / donor_channel) – (acceptor_channel(no_ligand) / donor_channel(no_ligand)) * 1 000.

### Real-time Quantitative PCR

Cells were seeded at 10 000 cells in 200 µL culture medium per well in a 96-well plate. Cells were allowed to adhere overnight. Compound dilutions were added from DMSO stock solutions using a HP D300 Dispenser (Tecan) (final DMSO concentration 0.1%). Typically, cells were incubated at 37 °C for 6 hours. Treated cells were processed using the FastLane Cell Multiplex Kit (Qiagen #216513). Threshold cycle numbers were determined using an AriaMx Real-time PCR System (Agilent Technologies) and GAPDH (Applied Biosystems Cat. No. 4326317E-1301046, VIC) and DUSP6 (Life Technologies Cat No. Hs00169257_m1, FAM) probes. Results were analyzed by relative quantification and are specified as percent of matched DMSO controls. *IC_50_* values and 95% confidence intervals were computed using a 3-parametric logistic model restricting the bottom to be > 0.

### Protein Detection by Capillary Electrophoresis

Cells (50 000) were seeded the day before treatment into 96-well clear-flat bottom plates and incubated overnight (37 °C, humidified tissue culture incubator, 5% CO_2_). Compounds were added from DMSO stock solution using a digital dispenser and cells are incubated at 37 °C as specified (24 hours for KRAS degradation, 6 hours for determination of pERK levels). At the end of the incubation, medium was removed, cells washed with ice cold PBS, PBS removed and 50 µL lysis buffer (Meso Scale Discovery #R60TX-2) supplemented with 1:100 protease inhibitor solution, 1:100 phosphatase inhibitor I, 1:100 phosphatase inhibitor II (Meso Scale Discovery #R70AA-1), 0.5 µL/mL Benzonase (Novagen #70664-3) and 10 mM dithiothreitol added to each well. Plates with lysis buffer were immediately transferred to -80 °C freezer for at least 2 hours and then thawed on ice. Lysates were transferred to a 96 well V-bottom plate and insoluble debris was pelleted by centrifugation for 20 min. at 4 000 rpm. 40 µL of the supernatant were transferred to a fresh Eppendorf PCR plate. Analysis of protein levels was conducted using the Wes System, ProteinSimple (https://www.bio-techne.com/) as specified by the manufacturer. Briefly, Wes Master Mix (12-230 kDa WES Separation Module, 8 × 25 capillary cartridges, ProteinSimple #SM-W004) and molecular weight markers were prepared as specified in the manufacturer’s recommendations. To detect primary antibodies, anti-rabbit (Protein Simple #DM-001) or anti-mouse (Protein Simple #DM-002) detection modules were used. 7.2 µL lysate + 1.8 µL master mix were combined and heated to 95 °C for 5 min. The following antibodies were used and diluted as recommended in Antibody Diluent II: KRAS^G12D^ (Cell Signaling #14429, 1:100), KRAS^G12V^ (Cell Signaling #14412, 1:100), KRAS (Lifespan C175665, 1:100), Phospho-p44/42 MAPK (Cell Signaling #9101, 1:200), p44/42 MAPK (Cell Signaling #9102, 1:400), GAPDH (Abcam #ab9485, 1:2 000), alpha-actinin (Cell Signaling #3134, 1:1 000), p70 S6 Kinase (Cell Signaling #9202, 1:50), phospho-p70 S6 Kinase (Thr389, Cell Signaling #9205, 1:50). Results were analyzed using the Compass Software. Capillaries with no signal were assumed to be experimental artifacts and were excluded from further analyses as indicated in the raw data. Protein levels were normalized to DMSO controls run in the same cartridge and specified in % of control. *IC_50_* or *DC_50_* and *D_max_* values including 95% confidence intervals were computed using a 3-parametric logistic model restricting the bottom to be > 0.

### Western Blot

#### VHL inhibitor VH298, proteasome inhibitor MG132 and cullin neddylation inhibitor ML4924 co-treatments

5×10^5^ GP5d cells were seeded into 6-well plates 24 hours before treatment. Next day, several wells were treated with 30 nM of Compound **4** or ACBI3. Immediately after, VH298 (Selleckchem #S8449), MG132 (Selleckchem #S2619) and MLN4924 (Tocris #6499) were added to the wells at 10 μM final concentration and incubated for 4 hours. 0.1% (v/v) DMSO was added to the wells with single treatments to achieve the same concentration of DMSO in all wells.

#### Degradation experiments

5 × 10^5^ GP5d cells in 2 mL/well were seeded into 6-well plates 24 hours before treatment. Cells were treated for 4 hours as indicated, washed with PBS and lysed with lysis buffer (1% Triton X-100, 150 mM NaCl, 1 mM EDTA, 50 mM Tris pH 7.4, protease inhibitor cocktail (Roche), 50 units/mL benzonase nuclease (Sigma). Mouse embryonic fibroblasts were seeded at 5 × 10^5^/mL 24 hours before the treatment. They were treated with indicated compounds at 1 µM for 6 and 24 hours, then subsequently washed with PBS and lysed with the same lysis buffer as described above. Both sets of lysates were cleared by centrifugation at 4 °C, at 15 800 × *g* for 10 min. and the supernatants stored at -20 °C. Protein concentration was determined by BCA assay (Pierce) and the absorbance at 562 nm measured by spectrophotometry on a plate reader (BMG Labtech PHERAstar). Samples were separated by SDS-PAGE using 20 μg of protein per well of NuPAGE Novex 4-12% BIS-TRIS gels (Invitrogen) and transferred to 0.2 μm pore nitrocellulose membrane (Amersham) by wet transfer. Western blot images were obtained through detection of mouse anti-pan-KRAS (1:1 000, LsBio-C175665), rabbit anti-β-actin (1:2 500, Cell Signaling Technology #4970S), mouse anti-β-actin (1:2 500, Cell Signaling Technology #3700S), hFAB™ rhodamine anti-actin (1:5 000, Biorad #12004164) and rabbit anti-GAPDH (1:2 500, Abcam ab9485) antibodies with donkey anti-rabbit IRDye 800CW secondary antibody (1:10 000, LI-COR #926-32213) using a ChemiDoc MP imaging system (Bio-Rad). Western blots were quantified using Image Studio Lite (Licor, version 5.2) with normalization to loading control and DMSO and further analyzed using GraphPad Prism (version 9.2.0) and ImageJ (Fiji).

### Pharmacokinetic Analyses

Compound concentrations in plasma aliquots were measured by quantitative HPLC-MS/MS using an internal standard. Calibration and quality control samples were prepared using blank plasma from untreated animals. Samples were precipitated with acetonitrile and injected into a HPLC system (Agilent 1200). Separation was performed by gradients of 5 mmol/L ammonium acetate pH 4.0 and acetonitrile with 0.1% formic acid on a 2.1 mm by 50 mm XBridge BEH C18 reversed-phase column with 2.5 µm particles (Waters). The HPLC was interfaced by ESI operated in positive ionization mode to a triple quadrupole mass spectrometer (5000 or 6500+ Triple Quad System, SCIEX) operated in multiple reaction monitoring mode. Chromatograms were analyzed with Analyst (SCIEX) and pharmacokinetic parameters were calculated by non-compartmental analysis using BI-proprietary software.

### Solubility Testing

Compound solubility was determined by dilution of a 10 mmol/L compound solution in DMSO into buffer to a final concentration of 125 µg/mL. Dilution into a 1:1 mixture of acetonitrile and water was used as reference. After 24 hours, the incubations were filtrated, and the filtrate was analyzed by LC-UV.

### Microsomal Stability

The degradation kinetics of 1 µmol/L compound in 0.5 mg/mL liver microsomes were inferred in 100 mM Tris-HCl pH 7.5, 6.5 mM MgCl_2_ and 1 mM NADPH at 37 °C. Reactions were terminated by addition of acetonitrile and precipitates separated by centrifugation. Compound concentrations in supernatants were measured by HPLC-MS/MS and clearance was calculated from compound half-lives using the well-stirred liver model.

### Plasma Protein Binding

Binding of compound to plasma proteins was determined by equilibrium dialysis of 3 µmol/L compound in plasma or serum against PBS through an 8 kDa molecular-weight cut-off cellulose membrane (RED device, Thermo Fisher) at 37 °C for 5 hours. After incubation, aliquots from donor and acceptor compartments were precipitated and the concentrations in the supernatants were determined by quantitative LC-MS/MS. Calibration and quality control samples were prepared using blank plasma and internal standard. The fraction unbound was calculated as ratio of the compound concentration in the acceptor compartment to the concentration in the donor compartment.

### Bidirectional Permeability in Caco-2 Cells

Bidirectional permeability of test compounds across a Caco-2 cell monolayer was measured as described (*45*). Briefly, Caco-2 cells were seeded onto Transwell inserts (Corning, Wiesbaden, Germany) at a density of 160 000 cells/cm^2^ and cultured in DMEM (high glucose) containing 10% FCS for 14-21 days. Cells were incubated with culture media containing 1 µM compound for 24 hours. After the preincubation period, culture media was removed and fresh transport buffer (128 mM NaCl, 5.4 mM KCl, 1 mM MgSO_4_, 1.8 mM CaCl_2_, 4.2 mM NaHCO_3_, 1.2 mM Na_2_HPO_4_, 0.41 mM NaH_2_PO_4_, 15 mM 2-[4-(2-hydroxyethyl)piperazin-1-yl]ethanesulfonic acid (HEPES), 20 mM glucose, pH 7.4, 0.25% bovine serum albumin) containing 1 µM test compound was added to the apical (apical to basal) or basal (basal to apical) compartment (donor compartment). Transport buffer without test compound was added to the opposite compartment (receiver compartment). Samples were taken at different time points for up to 2 hours. Test compound in the samples was quantified by LC-MS/MS. To elucidate the role of drug transporter P-gp in the transcellular transport, permeability measurement was performed in the absence and presence of 5 µM of the selective P-gp inhibitor zosuquidar. Apparent permeability coefficients (P_ab_, P_ba_) were calculated as P_ab_ = Q_ab_ / (C_0_ × s × t) and P_ba_ = Q_ba_ / (C_0_ × s × t) where Q is the amount of compound recovered in the receiver compartment after the incubation time t, C_0_ the initial compound concentration given to the donor compartment, and s the surface area of the Transwell inserts. Efflux ratio is calculated as the quotient of P_ba_ (mean of duplicate) to P_ab_ (mean of duplicate). The P-gp substrate apafant and one low permeable compound (BI-internal reference, P_ab_ ≈ 3 × 10^−7^ cm/s, no efflux) were included in every assay plate. In addition, transepithelial electrical resistance (TEER) values were measured for each plate before the permeability assay. All three parameters (efflux of the reference substrates, P_ab_ values of the low permeable compound, and TEER values) were used to ensure the quality of the assays.

### Animals, Biomarker and Xenograft experiments

*In vivo* experiments were performed at the AAALAC accredited animal facility of Boehringer Ingelheim RCV GmbH & CoKG. Female BomTac:NMRI-*Foxn1^nu^* mice were obtained from Taconic Denmark at 6-8 weeks of age. After the arrival, mice were allowed to adjust to the housing conditions for at least 5 days before the start of the experiment. Food and water were provided ad libitum. Mice were group housed (3-10 mice per cage) in pathogen-free and controlled environmental conditions (21 ± 1.5 °C temperature, 55 ± 10% humidity, and 12hours light-dark cycle; open cage housing), and handled according to the institutional, governmental and European Union guidelines (Austrian Animal Protection Laws, GV-SOLAS and FELASA guidelines).

Studies were approved by the internal ethics committee of Boehringer Ingelheim RCV GmbH & Co KG in the department of Cancer Pharmacology and Disease Positioning. Furthermore, all protocols were approved by the Austrian governmental committee (MA 60 Veterinary office; approval numbers GZ: MA 58-426431-2019-13, MA 58-458067-2021-17 and MA 58-546776-2022-13).

For pharmacokinetic (PK) studies mice (N=3 per group) were dosed with the respective degrader as an *i.v.* bolus (5 mL/kg) at the indicated dose formulated with HP-ß-CD 25%, as an *s.c.* bolus (5 mL/kg; PEG-400, Kolliphor HS 15 and Transcutol formulation) or as an *i.p.* bolus (5 mL/kg; nanomilled suspension formulation). Five plasma samples were collected from the *vena saphena* at defined timepoints.

To establish subcutaneous tumors for biomarker or xenograft experiments, mice were injected with 5 × 10^6^ GP2d cells with 1:2 Matrigel : PBS with 5% FBS. Tumor diameters were measured with a caliper. The volume of each tumor [in mm^3^] was calculated according to the formula “tumor volume = length * diameter^2^ * π/6.”

For the biomarker study mice were randomized (N=7 for the control and N=5 for each treatment group) when tumor size reached between ∼200 and 500 mm^3^. Mice were treated with ACBI3 three consecutive days before tumors and plasma were collected at 6 and 24 hours after the last treatment. Fresh frozen tumors were lysed in ice-cold Tris lysis buffer (50 mM NaCl, 20 mM Tris-HCl pH 7.5, 1 mM EDTA, 1 mM EGTA, 1% Triton-X-100), supplemented with phosphatase inhibitors (Sigma product codes P0044 and P5726). Lysates were cleared by centrifugation (10 min., 10 000 × g) and analyzed by capillary electrophoresis as described herein.

For analysis of KRAS levels in mouse PBMCs, about 600 µL mouse EDTA blood were diluted 1:4 with PBS supplemented with 2% heat inacivated FCS and 1 mM EDTA. The diluted EDTA blood was then transferred to SepMate tubes (Stemcell Technologies #85415) filled with 3.8 mL of Lymphoprep solution (Abcam #ab286892) and centrifuged for 20 min. at 1200 × g. The upper 1.5 mL were removed by pipetting without disturbing the interphase containing PBMCs. The interphase was collected, transferred into 15 mL Falcon tubes filled up to 15 mL and washed with PBS supplemented with 2% heat inactivated FCS and 1 mM EDTA (400 × g centrifugation, 8 min.). The cell pellet was then lysed in 80 µL lysis buffer (Meso Scale Discovery #R60TX-2) supplemented with 1:100 protease inhibitor solution, 1:100 phosphatase inhibitor I, 1:100 phosphatase inhibitor II (Meso Scale Discovery #R70AA-1), 0.5 µL/mL Benzonase (Novagen #70664-3) and 10 mM dithiothreitol and analyzed by Wes as described herein loading 0.8 µg protein per capillary.

For analysis of KRAS levels in mouse spleens, spleens (stored at -80 °C) were lysed in 79 µL/mg of lysis buffer (Meso Scale Discovery #R60TX-2) supplemented with 1:100 phosphatase inhibitor cocktail #2 (Sigma #P5726), phosphatase inhibitor cocktail #3 (Sigma #P0044), protease inhibitor cocktail (Roche #11836170001), 1 mM phenylmethansulfonylfluoride (Sigma #P7626) and 0.1% sodium dodecylsulfate (Sigma #75746) using Ready Prep Mini grinders and resin tubes (Biorad #163-2146) as recommended by the manufacturer. Lysates were cleared by centrifugation (10 min., 10 000 × g) and analyzed by capillary electrophoresis as described herein loading 0.8 µg protein per capillary.

For the xenograft study mice were randomized into the treatment groups when tumor size reached ∼220-260 mm^3^. Group sizes were calculated based on historical GP2d or RKN xenograft experiments. 10 mice per group for GP2D and 7 mice per group for RKN were used for the study based on the power analysis performed with a sample size calculator. 30 mg/kg ACBI3 was formulated in PEG-400, Kolliphor HS 15 and Transcutol (*s.c.*, 5 mL/kg injection volume) or as a nanomilled suspension in hydroxypropyl cellulose, polysorbate 80 and sodium dodecyl sulfate (*i.p*., 5 mL/kg injection volume). Nanomilling was performed using a ZentriMix 380R (Andreas Hettich GmbH & Co. KG) dual centrifuge with zirconium oxide milling beads for 4 hours at 1 000 rpm and 4 °C. .These formulations were dosed daily with control mice receiving the same formulation at the same dosing schedule without compound added. TGI (tumor growth inhibition was calculated with the formular: TGI =100 × (1-[(treated final day – treated day 1) / (control final day – control day 1)]). In case tumor sizes in control group of RKN tumors reached ethical endpoint (1 500 mm^3^) we carried forward the last measurements for the respective animals for figure 4L. To monitor side effects of treatment, mice were inspected daily for abnormalities and body weight was determined daily during treatment. Two animals out of the GP2d *s.c.* efficacy experiment treatment groups were euthanized at day 7 due to skin lesions and one animal of the *s.c.* efficacy experiment control group was euthanized at day 9 due to skin lesion/irritation. Hence, *s.c.* dosing of ACBI3 formulated in PEG-400 / Transcutol / Kolliphor HS 15 is not recommended for further *in vivo* use. Nanomilled ACBI3 at doses of 30 mg/kg was tolerated upon repeated *i.p*. dosing for up to 14 days. BI-2493 formulated in 0.5% Natrosol and administered in a dosing volume of 10 mL/kg as two doses of 90 mg/kg per day dosed 6 hours apart (daily dose 180 mg/kg).

### Targeted proteomics assay for KRAS^G12C^, KRAS^WT^, HRAS and NRAS

Lysates of 50 000 cells per sample were precipitated using a methanol-chloroform method. Precipitates were digested with 5 µg Lys-C and 10 µg Trypsin in digestion buffer (5 mmol/L tris(2-carboxyethyl)phosphin hydrochloride, 10 mmol/L 2-chloro-acetamide, 1% (w/v) sodium deoxycholate, 50 mmol/L triethylammonium bicarbonate pH 8.5) containing stable isotope-labeled synthetic peptides with the following amino acid sequences: LVVVGAGGVGK(+8), LVVVGAC(carbamidomethyl)GVGK(+8), SFEDIHHYR(+10)EQIK, SFADINLYR(+10)EQIK, SFEDIHQYR(+10)EQIK (SpikeTides TQL, JPT). Digests were acidified and purified by C18 RP solid-phase extraction. Peptides were chromatographically separated on 200 cm µPAK C18 RP columns (PharmaFluidics) using a 60 min. gradient of effectively 2 to 28% acetonitrile in 0.1% formic acid and water at 300 nL/min. on a RSLCnano HPLC system (Thermo Scientific) interfaced by ESI to a Q-Exactive Plus mass spectrometer (Thermo Scientific). Eluting peptide ions were analyzed using parallel reaction monitoring with an isolation window of 0.4 Th, a resolving power of 35 000, an automatic gain control target of 1e5 and a maximum injection time of 120 ms. Included were [M+2H]^2+^ or [M+3H]^3+^ molecular ions of the targeted peptides. Data were analyzed with Skyline and processed with R. The number of wild-type KRAS molecules per cell was calculated as mean of LVV..G minus SFE..H – SFE..Q and SFA minus LVV..C molecules per cell, while the number of molecules per cell of the other RAS species where directly inferred from the individual peptides. KRAS4A and -4B were not discriminated.

### Multiplexed Relative Phospho-proteomics

NCI-H358 cells (4 × 10^6^ per 10 cm dish seeded on the day before treatment) were treated with 500 nM of compound **4** or compound **5**, GP2d (5 × 10^6^ per 10 cm dish seeded on the day before treatment) cells were treated with 100 nM of compound **4** or compound **5** for 5, 10, 20, 30, 40 min., 1, 2, 4, 8, 16, 24, 36 and 48 hours. To correct for any potential treatment-unrelated abundance changes, the cells were treated with DMSO vehicle control for 1 hour, 24 hours and 48 hours (13 treatment time points and 3 controls, 16 conditions in total). After the treatment, cells were washed twice with 5 mL PBS and lysed in 200 μL lysis buffer (2% SDS in 100 mM Tris-HCl, pH 7.5, 95 °C, 10 min.). Lysate was then acidified to 1% trifluoroacetic acid (Sigma Cat. No. 302031) for DNA hydrolysis, followed by neutralization with 4-methylmorpholine (Sigma Cat. No. 407704, 2% final concentration), and stored at -20 °C until further use.

Sample preparation was performed following the previously reported procedures (*46*). The protein yield was determined by Thermo Pierce BCA protein assay. All steps were performed according to the manufacturer’s protocol. 200 μg protein lysate per treatment condition was cleaned by protein aggregation capture on a Bravo Agilent pipetting system following the SP3 Protocol. Briefly, 200 μg protein lysate was mixed with 1500 μg (7.5:1 beads to lysate ratio, 1:1 mix of washed magnetic Sera-Mag™ A & B; Cytiva) for 10 min. 1 000 rpm, room temperature, on a thermoshaker. All volumes were adjusted to 120 μL using 10 mM Tris-HCl (pH 7.6).

Proteins were precipitated by adding 100% ethanol to a final concentration of 70% and shaking for 10 min. at 1250 rpm on the Bravo shaking unit. The supernatant was removed on the magnet. The lysate was washed 3 times with 80% Ethanol, with 5 min. shaking intervals off the magnet and supernatant removal on the magnet. Finally, beads were washed with 100% acetonitrile to remove residual ethanol. Proteins were reduced and alkylated in 100 μL RA buffer (100 mM EPPS/NaOH, pH 8.5, 50 mM CAA, 10 mM TCEP) for 1 hour at 37 °C and 1200 rpm. Next, 2 μg of trypsin was added and digestion was performed overnight at 37 °C and 1 000 rpm. The peptides were recovered on the Bravo magnet. Beads were washed with 110 μL 2% TFA for 3 min. at 1250 rpm on the Bravo shaking unit and the wash supernatant was pooled to the recovered peptides. To further remove magnetic beads from the digest, the recovery plate was further incubated on a magnetic rack for 1 hour at 4 °C.

The digest was desalted using HLB desalting plates (10 mg N-vinylpyrrolidon-divinylbenzol porous particles 30 μm). Plates were cleaned by 500 μL isopropanol, acetonitrile, and solvent B (0.1% TFA 70% acetonitrile), and equilibrated by 1 000 μL solvent A (0.1% TFA in deionized water), each step was followed by a 1 min. centrifugation at 500 rpm. The digest was loaded by gravity. After washing twice with 1 000 μL solvent A, peptides were eluted by gravity in 200 μL Solvent B, and the residual volume was collected via a 1 min. centrifugation at 1 000 rpm. The cleaned peptides were dried down in the speed-vac and stored at -20 °C.

The dried and cleaned peptides were reconstituted in 20 μL of 100 mM EPPS*NaOH (pH 8.5) buffer. TMT-16-plex reagent (Thermo Scientific; LOT: VI310352) was reconstituted in water-free acetonitrile to a working concentration of 20 μg/μL. 5 μL of this TMT reagent solution was transferred to the peptides. The reaction was incubated on the thermoshaker (23 °C, 400 rpm) for 1 hour, followed by quenching with 5 μL of 1.5% hydroxylamine (final concentration of 0.25%). The TMT channels were then pooled together and acidified with formic acid (FA) to a final concentration of 1%. The reaction wells were washed with 25 μL washing solution (2.5% FA in 30% acetonitrile) and added to the TMT pool. The TMT pools were dried down in the speed-vac and stored at -20 °C.

The TMT-pooled peptides were cleaned by solid-phase extraction on 50 mg C18 Sep-PAK cartridges. Briefly, the cartridges were primed with 1 000 μL of acetonitrile, followed by the 1 000 μL washes of buffer B (0.1% FA in 50% acetonitrile), and 1 000 μL of buffer A (0.1% FA in deionized water). The TMT peptides were reconstituted in 700 μL of buffer A (pH adjusted to 2) and loaded on the Sep-PAK cartridges by gravity. The columns were washed twice with 1 000 μL of buffer A. TMT peptides were then eluted with 400 μL of buffer B, dried down in the speed-vac and stored at -20 °C.

TMT peptides were resolved with high pH reversed-phase (RP) fractionation on Thermo Scientific Vanquish HPLC equipped with a Waters Xbridge BEH130 C18 3.5 μm 4.6 × 250 mm column. In brief, dried peptides were reconstituted in 200 μL of 25 mM ammonium bicarbonate pH 8, and fractionated over a 60 min. gradient from 7% to 45% acetonitrile in the presence of 2.5 mM ammonium bicarbonate. The gradient was followed by a 7 min. ramp to 80% acetonitrile of the column wash. Fractions were collected from min. 7 to 55 after injection. Each fraction consisted of 30 s at a flow rate of 1 000 μL/min. The 96 fractions were pooled into 48 and acidified with FA to a final concentration of 0.1%. The 10% of fractionated TMT peptides was separately dried down in the speed-vac for MS analysis of the total proteome.

Phosphorylated peptides were separated from the TMT-pool peptide mix by immobilized metal ion affinity chromatography (IMAC) enrichment on a AssayMAP® Fe(III)-NTA cartridges with the Bravo Agilent pipetting system following the Phosphopeptide Enrichment Protocol, included in the Agilent AssayMAP® Bravo® Protein Sample Prep Workbench v2.0 software suite.

Briefly, the dried 48 TMT peptide fractions were reconstituted in 0.1% TFA 80% acetonitrile and combined into 12 fractions (final volume of 200 μL per fraction). The AssayMAP® Fe(III)-NTA cartridges were primed with 150 μL of 0.1% TFA in acetonitrile at 300 μL/min., and further equilibrated with 150 μL 0.1% TFA in 80% acetonitrile at 10 μL/min. The TMT-pool peptide mix was loaded on the cartridges at 5 μL/min., and the flow through was collected. The cartridges were further washed three times with 150 μL 0.1% TFA in 80% acetonitrile at 50 μL/min. Upon a stringent syringe wash with 200 μL 0.1% TFA in acetonitrile, the retained on the cartridges phosphorylated peptides were eluted with 60 μL of 1% ammonium hydroxide NH_4_OH at 5 μL/min. The eluates were acidified up to 0.1% FA, dried down in the speed-vac, and stored at -20 °C for MS analysis of the phosphoproteome.

Total proteome-TMT peptides were measured with a Fusion Lumos Tribrid mass spectrometer (Thermo Scientific) that was coupled to a Vanquish microflow pump (Thermo Scientific). The sample was reconstituted in 20 μL 1% FA 2% acetonitrile, and 10 μL were directly injected onto the Acclaim PepMap 100 C18 column (2 μm particle size, 1 mm ID × 150 mm). Separation was performed on a 25 min. gradient with a flow rate of 50 μL/min. starting from 4% B, followed by the linear phase to 32% B. The system was finally washed with 95 μL 90% B and re-equilibrated at 1% B. Solvent A consisted of 0.1% FA and 3% DMSO in water. Solvent B consisted of 0.1% FA and 3% DMSO in acetonitrile.

The MS was operated in A fast, data-dependent MS3-mode. The spray voltage was set to 3.5 kV supported by sheath gas (32 units) and aux gas (5 units) with a vaporizer temperature of 125 °C. Every 1.2 s, a full-scan (MS1) was recorded from 360 to 1600 m/z with a resolution of 60k in the Orbitrap in profile mode. The MS1 AGC target was set to 4e5, and the maxIT was set to 50 ms. Based on the full scans, precursors were targeted for MSMS scans if the charge was between 2 and 6, and the intensity exceeded 1e4. The MS2 quadrupole isolation window was set to 0.6 Th. The TMT peptides were HCD fragmented with an NCE of 34%. The MS2 spectra were acquired in the ion trap in rapid mode. The MS2 AGC target was set to 3e4 charges, and the maxIT was set to 40 ms. Precursors that have been targeted for fragmentation were excluded for 50 s for all possible charge stages. TMT reporter ions were measured in a consecutive MS3 scan based on the previous MSMS scan. Thus, a new batch of precursor ions was isolated with an MS3 quadrupole isolation window of 1.2. The isolated precursor was then HCD-fragmented identically to the previous MS2 scan. The top 8 fragment ions of the MS2 scans were isolated in the ion trap in parallel (synchronous precursor selection). Only fragment ions within a range of 400 to 2 000 Th were considered. The selected top10 fragment ions were then HCD fragmented with an NCE of 55%. The MS3 spectrum was acquired with 50k resolution from 100 to 1 000 Th in the Orbitrap in centroid mode. The MS3 AGC target was set to 2e5 charges, and the maxIT was set to 86 ms.

Phospho-TMT peptides were measured with a Fusion Lumos Tribrid mass spectrometer (Thermo Scientific) that was coupled to a Dionex UltiMate 3000 RSLCnano System (Thermo Scientific). Each of the 12 peptide fractions was reconstituted in 20 μL of 50 mM sodium citrate buffer, 10 μL were then injected for the MS analysis. After injection, the sample was transferred onto a trap column (75 μm × 2 cm) that was packed with 5 μm C18 resin (Reprosil PUR AQ – Dr. Maisch). Peptides were washed with the trap washing solvent (5 μL/min., 10 min.) before conveying them to an analytical column (75 μm × 48 cm) that was packed with 3 μm C18 resin (Reprosil PUR AQ – Dr. Maisch). Separation was performed on an 80 min. gradient with a flow rate of 300 nL/min. starting from 4% B, followed by the first linear phase to 22.5% B in 65 min., followed by the second linear phase to 32% B in 15 min. The system was finally washed with 80% B for 2 min. and re-equilibrated at 2% B. Solvent A consisted of 0.1% FA and 5% DMSO in water. Solvent B consisted of 0.1% FA and 5% DMSO in acetonitrile.

The MS was operated in a sensitive, data-dependent MS3-mode. Peptides were ionized using a nano source with 2.1 kV spray voltage. Every 3 s a full-scan (MS1) was recorded from 360 to 1800 m/z with a resolution of 60k in the Orbitrap in profile mode. The MS1 AGC target was set to 4e5, and the maxIT was set to 50 ms. Based on the full scans, precursors were targeted for MSMS scans if the charge was between 2 and 6, the isotope envelop was peptidic (MIPS), and the intensity exceeded 5e4. The MS2 quadrupole isolation window was set to 0.7 Th. Peptide fragmentation occurred in the linear ion trap by CID-targeting the precursor and the precursor-H_2_PO_4_ in parallel (multistage-activation) with a q-value of 0.25, 35% CE, and 10 ms activation time. The MS2 spectrum was acquired with 30k resolution and auto scan range in Orbitrap in centroid mode. The MS2 AGC target was set to 5e4 charges, and the maxIT was set to 60 ms.

The maxIT or AGC target could be dynamically exceeded when the previous scan took longer than the calculated injection time (inject beyond mode). Precursors that have been targeted for fragmentation were excluded for 90 s for all possible charge stages. TMT reporter ions were measured in a consecutive MS3 scan based on the previous MSMS scan. Thus, a new batch of precursor ions was isolated with a charge stage-dependent MS3 quadrupole isolation window of 1.2 Th (z=2), 0.9 Th (z=3), 0.7 Th (z=4-6) to include the first isotope in the isolation while using the most narrow isolation window possible. The isolated precursor was then MSA-fragmented identically to the previous MS2 scan. The top 10 fragment ions of the MS2 scans were isolated in the ion trap in parallel (synchronous precursor selection). Only fragment ions were considered that were within a range of 400 to 2 000 Th and laid outside the precursor exclusion range (precursor -50 Th, precursor + 5 Th). The selected top10 fragment ions were then HCD fragmented with an NCE of 55%. The MS3 spectrum was acquired with 50k resolution from 100 to 1 000 Th in the Orbitrap in centroid mode. The MS3 AGC target was set to 1e5 charges, and the maxIT was set to 120 ms.

MaxQuant (version 2.1.3.0) with its built-in search engine Andromeda was used to identify and quantify phospho and total proteome TMT peptides. MSMS spectra were searched against the human Uniprot database supplemented with common contaminants. Unless stated otherwise, MaxQuant’s default parameters were applied. These included for all searches: Trypsin/P as the proteolytic enzyme with up to three missed cleavage sites allowed; carbamidomethylation of cysteine as fixed modification, oxidation of methionine and N-terminal protein acetylation as variable modifications; precursor tolerance was set to ± 4.5 ppm and fragment ion tolerance to ± 20 ppm (FTMS). Phosphorylation on serine, threonine, and tyrosine was allowed as variable modification specifically for the phosphoproteome enrichment data. Andromeda score and delta score cut-offs for modified peptides were set to 40 and 6, respectively; peptide spectrum match (PSM) and protein false discover rate (FDR) employing a target-decoy approach using reversed protein sequences were set to 1%. Isotope impurities of the TMT batch were specified in the configuration of TMT modifications to allow MaxQuant the automated correction of TMT intensities.

The data plotted in the Fig. 3F and G and fig. S7D-G were extracted from MQ evidence files, where prior to plotting, all impurity-corrected TMT channels were median-based normalized (*46*). Methionine oxidation was removed from the modified sequence, and all duplicated modified sequences were summed up. Next, ratios were calculated against the DMSO controls, where later time points were corrected for their respective DMSO controls of 24 hours and 48 hours. The phospho-responses were categorized for regulation by unsupervised K-means clustering algorithm on Z-score normalized phospho-response ratios (using the Euclidean distance after normalizing the sum of ratios across all TMT channels to 1). The obtained 7 clusters in each treatment were further filtered, and the phosphosites were considered as regulated following the criteria for downregulation: the p-response ratio must be < 0.5 after 8 hours (early) and/or 24 hours (late downregulation); and for upregulation: the ratio > 2 after 8 hours and/or 24 hours.

The protein groups and the evidence files from the total proteome datasets were processed as mentioned above and were used for plotting fig. S7B and -C. fig. S7C was plotted using peptide abundances as mentioned above, whereas fig. S7B represents the abundance of KRAS protein group across the treatment time points.

The gene set enrichment analysis of the median protein phosphorylation responses in both cell lines for Fig. 3F was performed using EnrichR tool (*47*) with integrated NCATS BioPlanet pathway resource. The GO ontology enrichment visualization for fig. S7G was performed using Proteomaps (*48*).

All raw data and search results have been deposited to the ProteomeXchange Consortium (http://www.proteomexchange.org/) via the MassIVE partner repository with the dataset identifier MSV000093122 (ProteomeXchange ID PXD046161). Reviewers can access the dataset at https://massive.ucsd.edu/ProteoSAFe/QueryPXD?id=PXD046161 using the following credentials: Login: MSV000093122_reviewer, Password: revieweronly.

### Multiplexed relative proteomics (Degradation Specificity)

GP2d cells in DMEM were seeded at 5 × 10^6^ cells on a 100 mm plate 24 hours before treatment. Cells were treated in triplicate by addition of compound **4** and compound **5** at 50 nM. After 8 hours, the cells were washed twice with 10 mL of cold PBS and lysed in 4% (w/v) SDS in 100 mM Tris at pH 8.5. MiaPaca-2 cells in RPMI 1640 (Invitrogen) were seeded at 5 × 10^6^ cells on a 100 mm plate 24 hours before treatment. Cells were treated in triplicate by addition of compound **4** and compound **5** at 100 nM for 8 hours. Gp5d cells in DMEM were seeded in triplicate at 5 × 10^6^ cells on a 100 mm plate 24 hours before treatment. Cell were treated with ACBI3 and compound **8** in triplicates at 50 nM and 100 nM for 4 hours. The cells were washed twice with 10 mL of cold PBS and lysed in 500 µL of 100 mM TEAB with 5% (w/v) SDS.

The lysates were pulse sonicated briefly and then centrifuged at 15 000 × g for 10 min. Samples were quantified using a micro-BCA protein assay kit (Thermo Fisher Scientific). 400 µg (Mia Paca-2) or 300 µg (GP2d and GP5d) of each sample was reduced with DTT, alkylated with iodoacetamide and digested with trypsin using the modified S-TRAP mini (ProtiFi) protocol.

Peptide quantification was done using Pierce™ Quantitative Fluorometric Peptide Assay and equal amount from each sample was labelled using TMTproTM 16-plex Label Reagent Set Set (GP2d and GP5d) or TMTproTM 10-plex Label Reagent Set Set (Mia Paca-2) (Thermo Fisher Scientific) as per the manufacturer’s instructions. The samples were then pooled and desalted using a 7 mm, 3 mL C18 SPE cartridge column (Empore, 3M).

LCMS method Q Exactive HF Hybrid Quadrupole-Orbitrap Mass Spectrometer: The pooled and desalted sample was fractionated using high pH reverse-phase chromatography on an XBridge peptide BEH column (130 Å, 3.5 μm, 2.1 × 150 mm, Waters) on an Ultimate 3000 HPLC system (Thermo Scientific/Dionex). Buffers A (10 mM ammonium formate in water, pH 9) and B (10 mM ammonium formate in 90% acetonitrile, pH 9) were used over a linear gradient of 2% to 100% buffer B over 80 min. at a flow rate of 200 μL/min. 80 fractions were collected using a WPS-3000 FC auto-sampler (Thermo Scientific) before concatenation into 20 fractions based on the UV signal of each fraction. All the fractions were dried in a Genevac EZ-2 concentrator and resuspended in 1 % formic acid for MS analysis. The fractions were analyzed sequentially on a Q Exactive HF Hybrid Quadrupole-Orbitrap Mass Spectrometer (Thermo Scientific) coupled to an Dionex Ultimate 3000 RS (Thermo Scientific). Buffers A (0.1% formic acid in water) and B (0.1% formic acid in 80% acetonitrile) were used over a linear gradient from 5% to 35% buffer B over 125 min. and then from 35% buffer B to 98% buffer B in 2 min. at a constant flow rate of 300 nL/min. The column temperature was 50 °C. The mass spectrometer was operated in data dependent mode with a single MS survey scan from 335-1600 m/z followed by 15 sequential m/z dependent MS2 scans. The 15 most intense precursor ions were sequentially fragmented by higher energy collision dissociation (HCD). The MS1 isolation window was set to 0.7 m/z and the resolution set at 120 000. MS2 resolution was set at 60 000. The AGC targets for MS1 and MS2 were set at 3×10^6^ ions and 1×10^5^ ions, respectively. The normalized collision energy was set at 32%. The maximum ion injection times for MS1 and MS2 were set at 50 ms and 200 ms respectively. The mass accuracy was checked before the initiation of sample analysis.

Peptide and protein identification Raw MS data files for all 20 fractions were merged and searched against the Uniprot-sprot-Human-Canonical database by Maxquant software 2.0.3.0 for protein identification and TMT reporter ion quantitation. The Maxquant parameters were set as follows: enzyme used Trypsin/P; maximum number of missed cleavages equal to two; precursor mass tolerance equal to 10 p.p.m.; fragment mass tolerance equal to 20 p.p.m.; variable modifications: oxidation (M), identifier (MW), acetyl (N-term), deamidation (NQ), Gln -> pyro-Glu (Q N-term); fixed modifications: carbamidomethyl (c). The data was filtered by applying a 1% false discovery rate followed by exclusion of proteins with less than two unique peptides.

Quantified proteins were filtered if the absolute fold-change difference between the three DMSO replicates was ≥ 1.5. The mass spectrometry proteomics data generated in cell line Mia Paca-2 have been deposited to the ProteomeXchange Consortium via the PRIDE (*49*) partner repository with the dataset identifier PXD045416. Data generated in cell line GP2d can be found in the same repository with the dataset identifier PXD045460.

LCMS Method Orbitrap Ascend Tribrid Mass Spectrometer: LC separation of the 20 fractions was performed on a Vanquish Neo System (Thermo Scientific) with a Mobile phase A consisting of 0.1% formic acid in HPLC water and Mobile phase B consisting of 0.1% formic acid in 80% acetonitrile. A step gradient was used over 135 min. at constant flow rate of 300 nL/min. consisting of 10% Mobile phase B for 6 min., 27% Mobile phase B for 83 min., 27% Mobile phase B for 25 min., and 35% Mobile phase B for 20 min. Eluted peptides were ionized by electrospray ionization and then analyzed by an Orbitrap Ascend Tribrid spectrometer operating in data dependent mode (Thermo Scientific). The spray voltage was set to 2.2 kV. The full MS1 scan was acquired with a scan range of 350-1 500 m/z at resolution 120 000 with AGC target set as standard. Precursor ions with charge states of 2-5 were isolated by the quadrupole mass filter at 0.7 m/z isolation window and fragmented by collision induced dissociation (CID) at normalized CID collision energy of 30%, activation time of 10 ms and activation Q of 0.25. MS2 analysis was performed in an ion trap detector with a scan range of 400-1200 m/z, the AGC target set as standard, and the maximum injection time 50 ms.

Top 10 most abundant ions were isolated from MS2 at 2 m/z and fragmented on higher energy collision dissociation (HCD) with normalized HCD collision energy of 55% upon synchronous precursor selection (SPS) and after the MS2 passed the real-time search filter (*50*). MS3 scan was performed in the Orbitrap at resolution 60 000 with a scan range of 100-500 m/z and normalized AGC target at 400%. Carbamidomethyl and TMTpro16-plex modification were set as static modifications on cysteine (C) and Lysine-N-terminus (Kn), respectively. Oxidation on Methionine (M) was set as variable modification. Maximum missed cleavages were set to 1.

Maximum variable mode/peptides were set to 2. The maximum search time was set to 100 ms. The scoring threshold was set to 1.4 XCorr, 0.1 dCn, 10 ppm precursor tolerance. MS data were acquired using XCalibur software (Thermo Scientific).

The dataset was analyzed in Perseus 2.0.1.1 software and has been deposited to the ProteomeXchange Consortium via the PRIDE partner repository with the dataset identifier PXD050650.

**Fig. S1.**
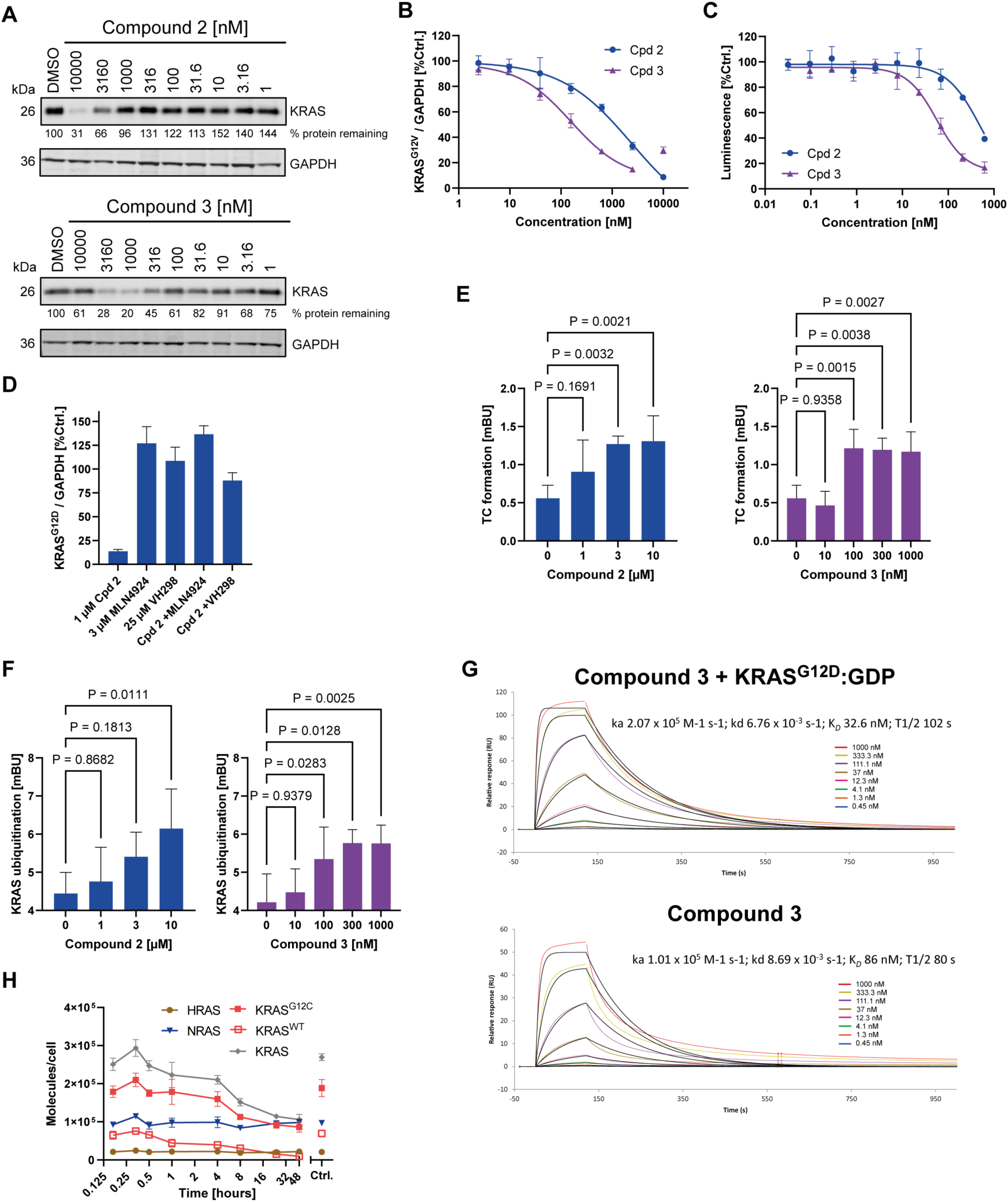
Validation of initial KRAS degraders. (**A**) KRAS (LsBio-C175665) and GAPDH levels by Western blot of GP5d cells treated with compounds **2** or -**3** (4 hours), numbers indicate normalized KRAS levels *vs* controls (N=3). (**B**) Dose-dependent degradation of KRAS^G12V^ in SW620 cells by capillary electrophoresis (24 hours, N=3, SD). (**C**) Dose-dependent degradation of HiBiT-KRAS^G12D^ in GP5d cells for compounds **2** and -**3** (24 hours, N=3, SD). (**D**) KRAS^G12D^ levels in GP5d cells treated with compound **2** in presence or absence of MLN4924 or VH298 by capillary electrophoresis (24 hours, N=3, SD). (**E**) Detection of compound **2** and -**3** induced ternary complex formation for KRAS^G12D^ in Gp5d cells (4 hours, N=3, SD). (**F**) KRAS^G12D^ ubiquitination in GP5d cells treated with compounds **2** or -**3** (4 hours, N=3, SD). (**G**) SPR characterization of ternary and binary complexes for immobilized VCB in the presence of KRAS^G12D^-GDP and compound **3**, or compound **3** alone at 20 °C. (**H**) Data as in Fig. 1G with protein abundance stated as molecules per cell.

**Fig. S2.**
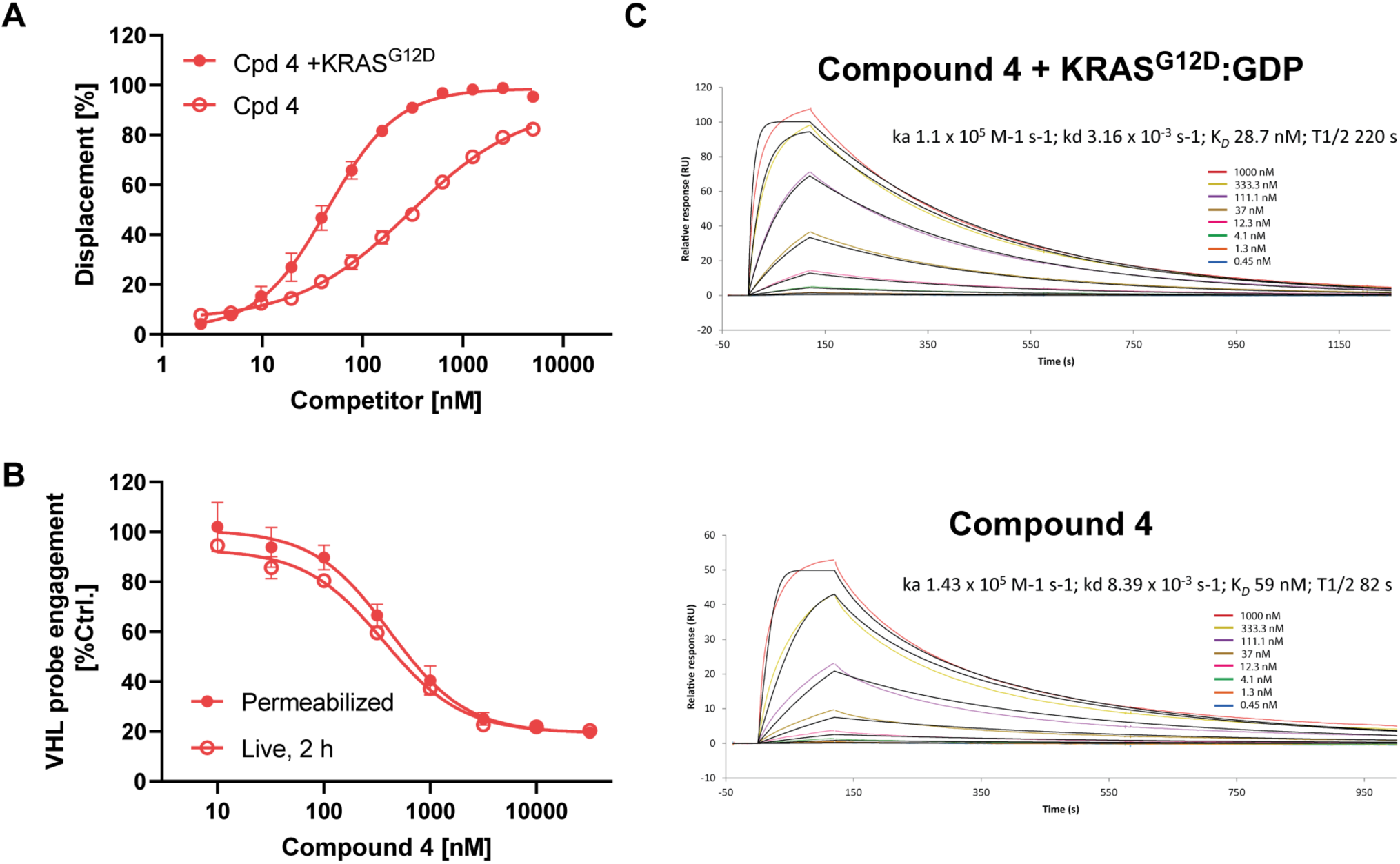
Ternary complex kinetics of compound 4. (**A**) VCB FP displacement for compound **4** in presence or absence of saturating KRAS^G12D^ concentrations (N=3, SD). (**B**) VHL target engagement by nanoBRET in live or permeabilized HEK293 cells for compound **4** (N=3, SD). (**C**) SPR characterization of ternary and binary complexes for immobilized VCB in the presence of KRAS^G12D^-GDP and compound **4**, or compound **4** alone at 20 °C.

**Fig. S3.**
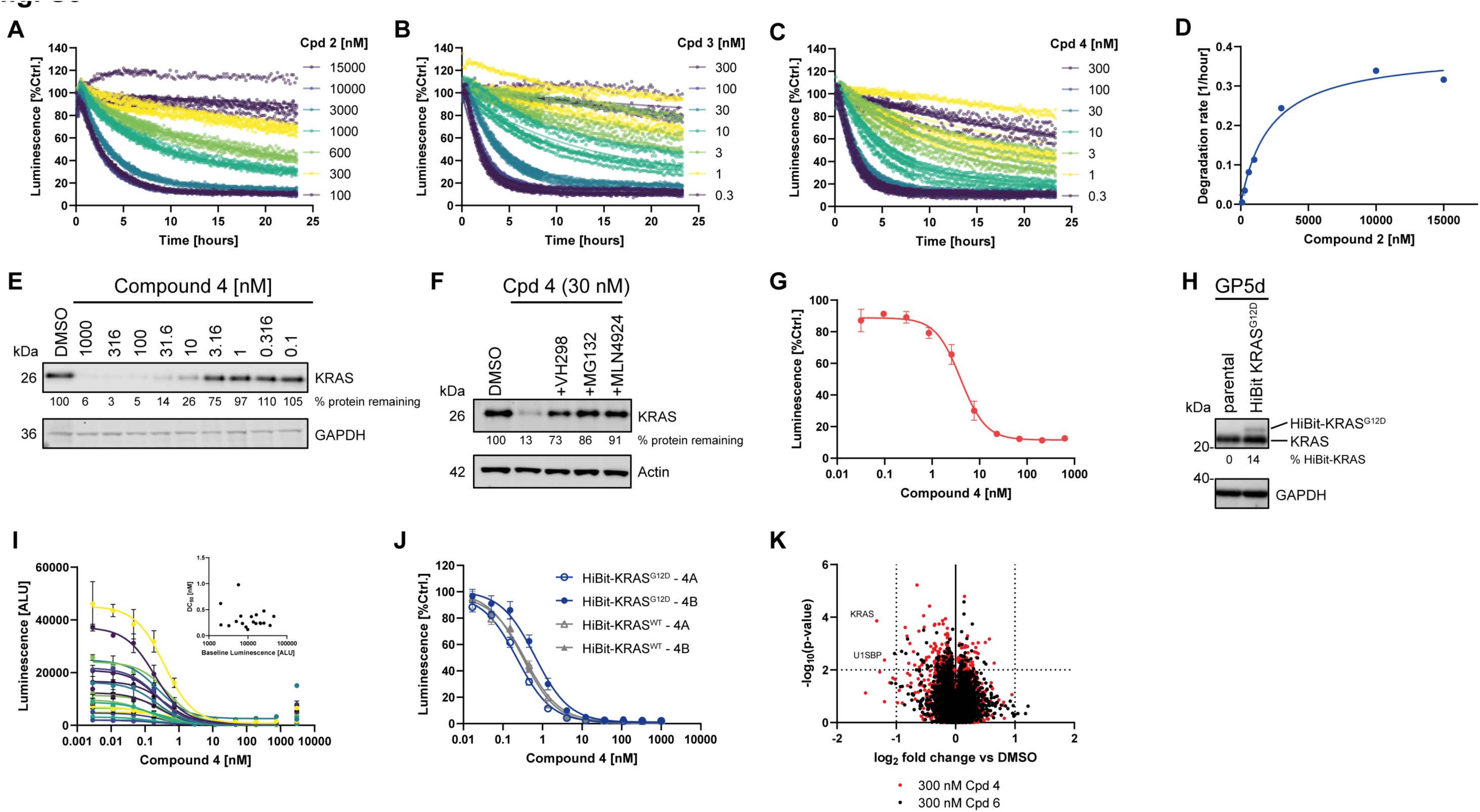
Degradation kinetics of KRAS degraders compound 2, -3 and -4. (**A**) Live cell degradation time course of HiBiT-KRAS^G12D^ in presence of varying concentrations of compound **2**, (**B**) compound **3** and (**C**) compound **4**. (**D**) Degradation rate *vs* concentration for compound **2** (N=6, 95% CI). (**E**) KRAS and GAPDH levels by Western blot of GP5d cells treated with compound **4** (4 hours), numbers indicate normalized KRAS levels *vs* controls (N=3). (F) KRAS levels in GP5d cell treated with compound **4** in presence or absence of MLN4924 or VH298 by Western blot (4 hours), numbers indicate normalized KRAS levels *vs* controls (N=3). (G) Degradation of KRAS^G12D^ by compound **4** in HibiT-tagged GP5d cells. (**H**) KRAS detection by Western blot of GP5d cell pools retrovirally transduced with luminescent HiBiT-tagged KRAS^G12D^, numbers indicate relative HiBiT-KRAS levels (% of endogenous KRAS) *vs* controls (N=3). (**I**) Dose dependent degradation of HiBiT-KRAS in single cell clones of GP5d cells retrovirally transduced with HiBiT-KRAS covering a range of expression levels (N=2, range). Inset plot depicts *DC_50_ vs* baseline expression levels. (**J**) Dose dependent degradation of HiBiT-KRAS^G12D^ or HiBiT-KRAS in GP5d cells retrovirally transduced with the corresponding KRAS4A or -4B isoform constructs. (**K**) Whole cell proteomics MS analysis of Mia PaCa-2 cells treated with 300 nM compound **4** or inactive stereoisomer compound **6** (8 hours, N=3).

**Fig. S4.**
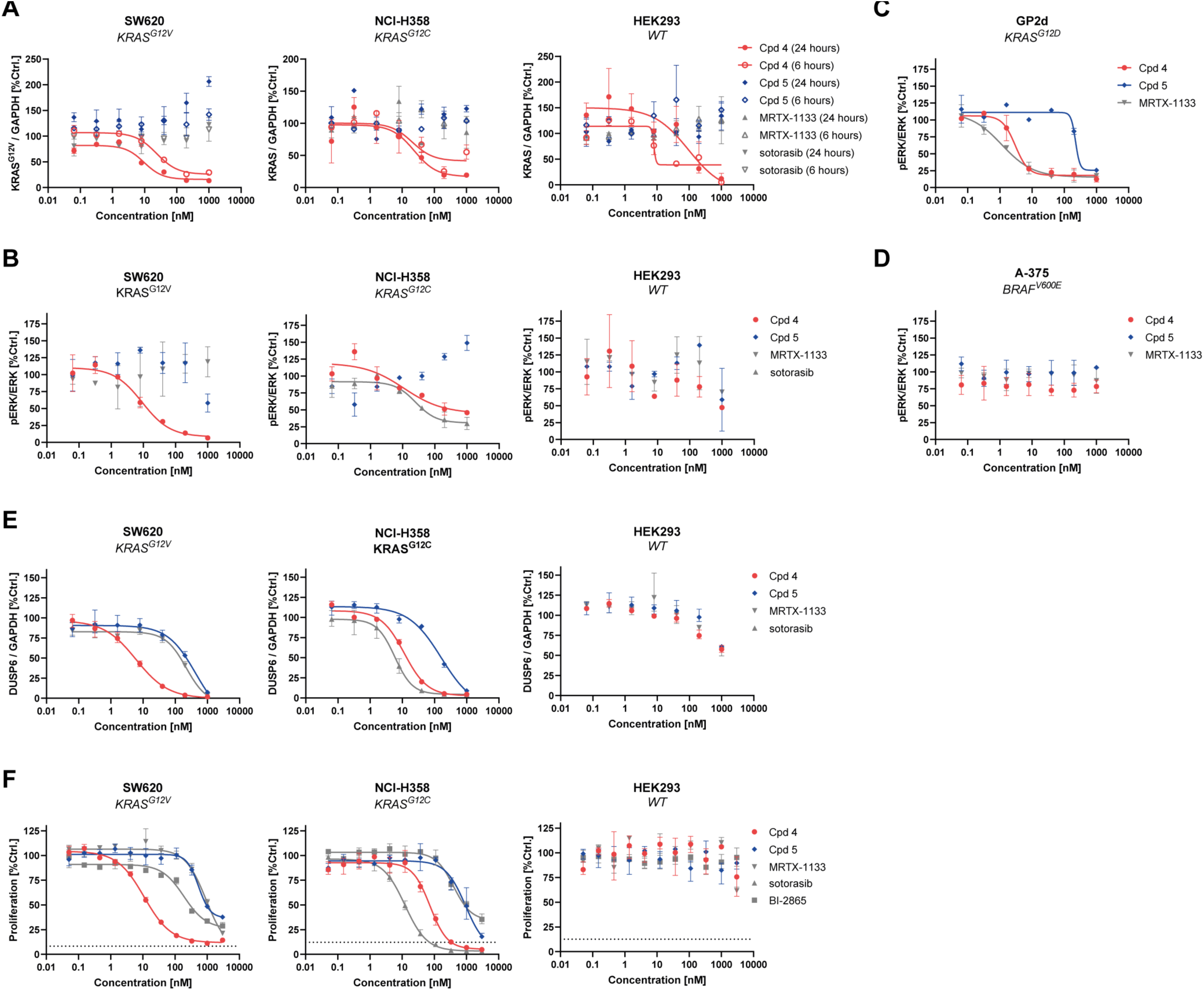
Extended characterization of pan-KRAS degradation on MAPK signaling and cancer cell proliferation. (**A**) KRAS degradation (6 and 24 hours, N=2, range), (**B**) pERK modulation (6 hours, N=2, range) for compound **4**, compound **5**, MRTX1133 or sotorasib in KRAS^G12V^, KRAS^G12C^ and KRAS^WT^ cell lines, respectively. (**C**) pERK modulation (6 hours, N=2, range) for compound **4**, compound **5** and MRTX1133 in GP2d cells and (**D**) A375 cells. (**E**) DUSP6 modulation (6 hours, N=3, SD), (**F**) proliferation (5 days, N=3, SD) for compound **4**, compound **5**, MRTX1133 or sotorasib in KRAS^G12V^, KRAS^G12C^ and KRAS^WT^ cell lines, respectively.

**Fig. S5.**
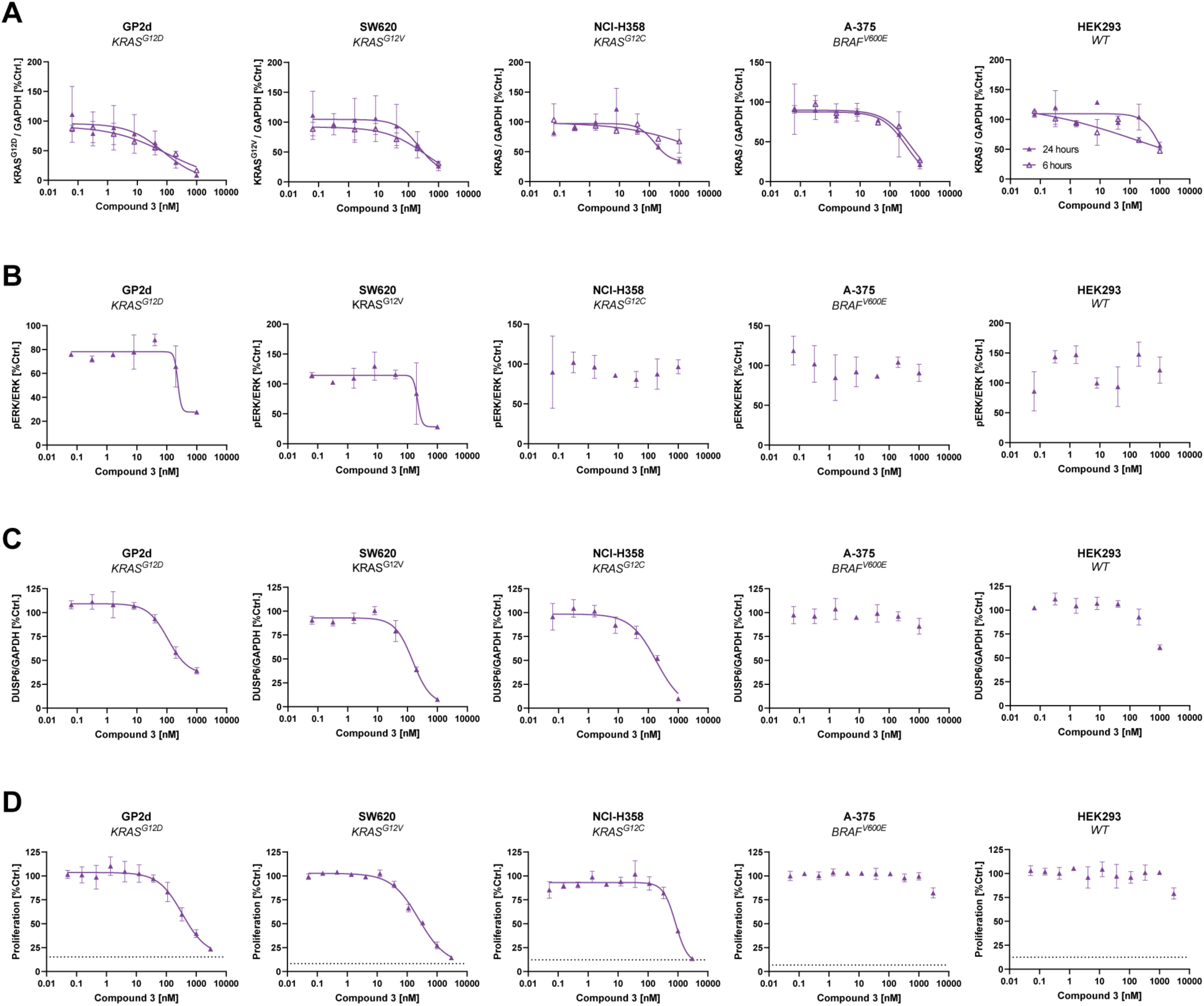
KRAS degradation and inhibition of MAPK pathway and cancer cell proliferation by compound 3. (**A**) KRAS degradation (6 and 24 hours, N=2, range), (**B**) pERK modulation (6 hours, N=2, range), (**C**) DUSP6 modulation (6 h, N=3, SD) (**D**) proliferation (5 days, N=3, SD) for compound **3** in KRAS^G12D^, -V and -C and -WT cell lines, respectively.

**Fig. S6.**
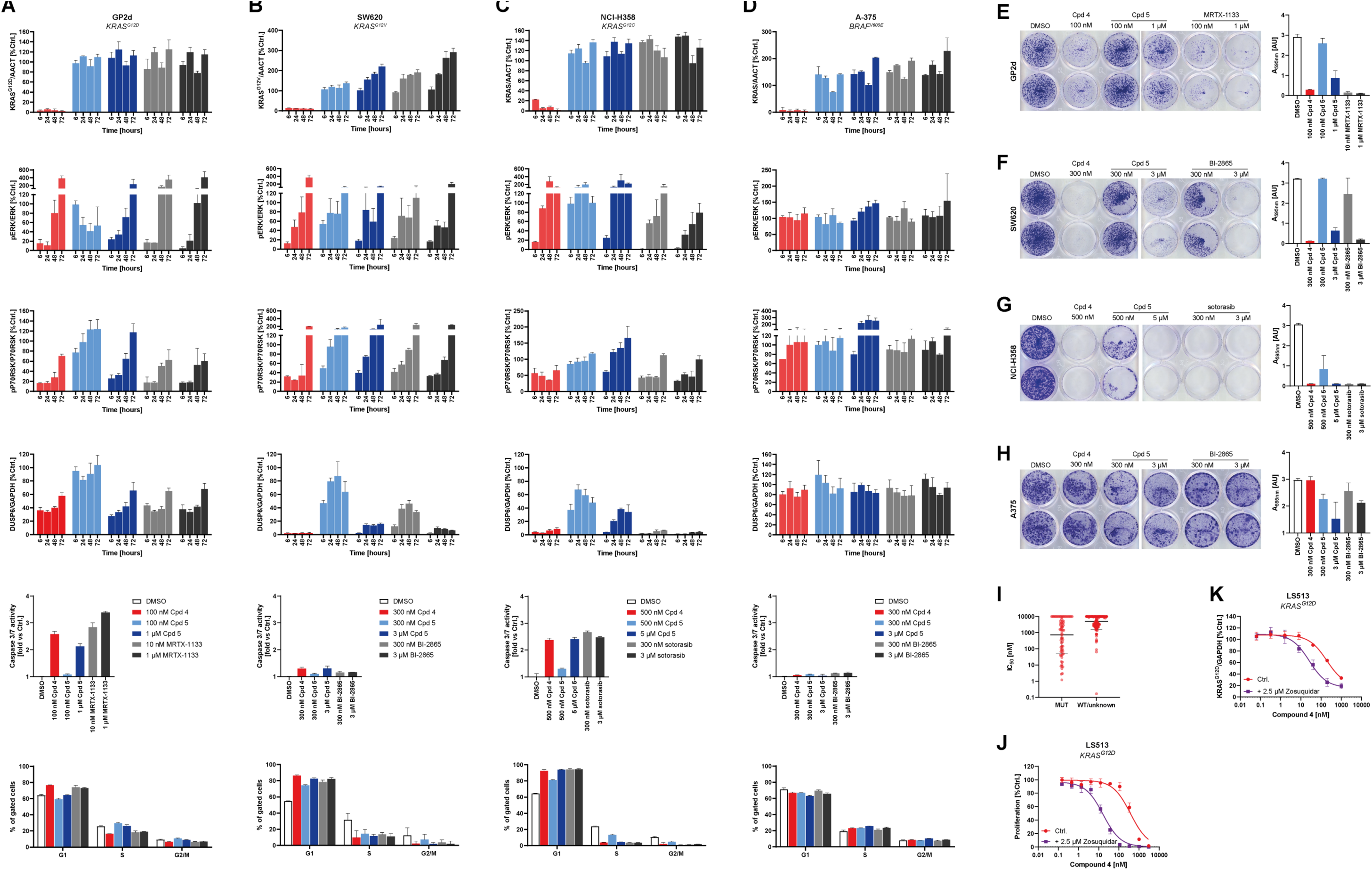
Long-term treatment pharmacodynamic effects and terminal phenotype of KRAS degradation *vs* inhibition in KRAS mutant cancer cell lines. (**A-D**) KRAS degradation (N=3, SD), pERK modulation (N=3, SD), p-p70RSK modulation (N=3, SD), DUSP6 modulation (N=3, SD), Caspase 3/7 induction (48 h, N=3, SD) or cell cycle profile (48 h, N=3, SD) for (**A**) GP2d-, (**B**) SW620-, (**C**) NCI-H358 or (**D**) A-375 cells treated with concentrations of compound **4** inducing maximal degradation, identical concentration or 10-fold excess of the inactive stereoisomer compound **5** or the indicated KRAS inhibitors. (E-H) Inhibition of clonogenic growth of (**E**) GP2d-, (**F**) SW620-, (**G**) NCI-H358 or (**H**) A-375 cells treated with concentrations of compound **4** inducing maximal degradation, identical concentration or 10-fold excess of the inactive stereoisomer compound **5** or the indicated KRAS inhibitors (N=4, images of representative duplicates, accompanying bar graphs depict quantification of all replicates). (**I**) Proliferation (5 days) for compound **4** in 300 cell line panel, analysis by KRAS mutation status (geometric mean, SD). (**J**) Dose dependent inhibition of proliferation of LS513 cells by compound **4** in presence or absence of 2.5 µM of Zosuquidar (5 days, N=3, SD). (**K**) Dose-dependent degradation of KRAS^G12D^ in LS513 cells in presence or absence of 2.5 µM of Zosuquidar (24 hours, N=3, SD).

**Fig. S7.**
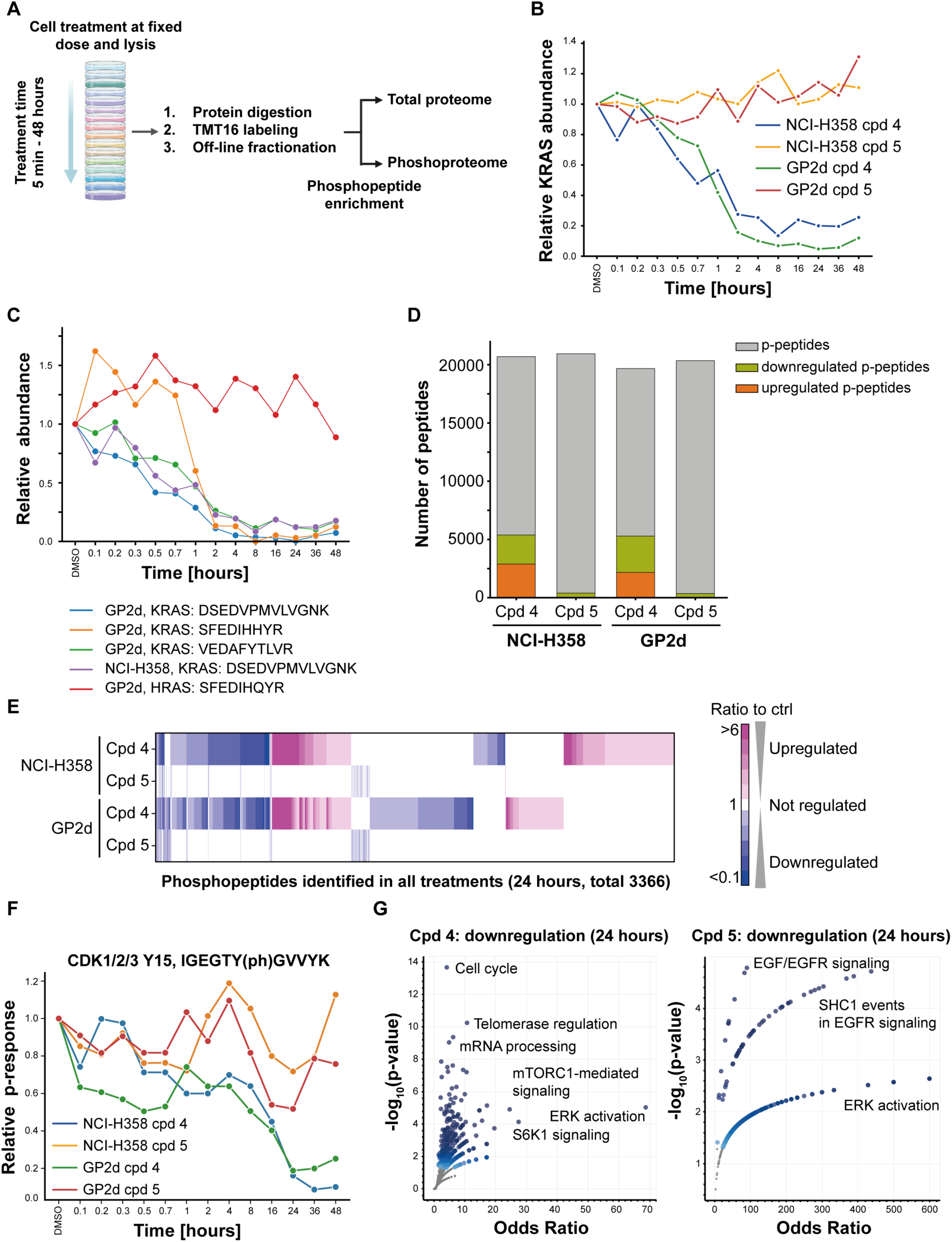
Phospho-proteomics characterization of KRAS degradation *vs* -inhibition. (**A**) Experimental design for the time-course analysis of the proteome and phospho-proteome of NCI-H358 (KRAS^G12C^) and GP2d (KRAS^G12D^) cells in response to compound **4** or compound **5** (500 nM for NCI-H358, 100 nM for GP2d). (**B**) Time course of changes in KRAS protein abundance. (**C**) Time course of changes in individual peptides representing KRAS and HRAS in cell lines treated with compound **4**. (**D**) Number of identified and regulated phosphorylation sites across treatments, categorization of up- or down-regulation of phosphorylation by unsupervised K-means clustering and filtering for the effect size of all identified phospho-sites. (**E**) Heatmap of the regulated phospho-proteome upon 24 hours of compound treatment. (**F**) Time-resolved changes in Tyr15 phosphorylation of CDK1/2/3. (**G**) Volcano plot of gene set enrichment analysis phospho-proteomes downregulated by compound **4** (left) or compound **5** (right) at 24 hours (data for both cell lines combined).

**Fig. S8.**
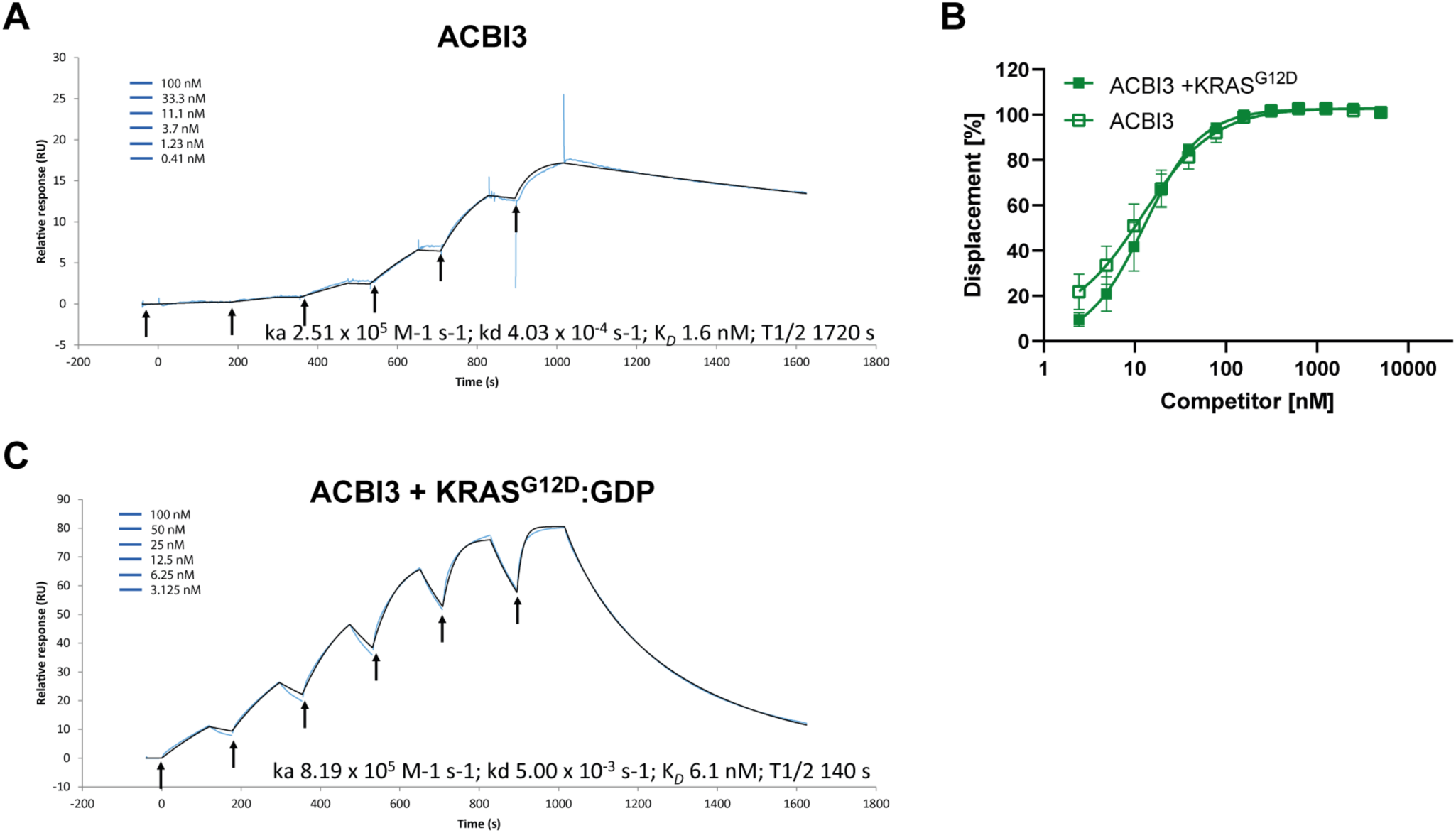
Biophysical characterization of ACBI3 ternary complexes. (**A**) Single-cycle kinetics of interactions between ACBI3 and VCB by SPR at 20 °C. (**B**) VHL FP displacement for ACBI3 in presence or absence of saturating KRAS^G12D^ concentrations. (**C**) Single-cycle kinetics of interactions between ACBI3 and VCB by SPR in presence of saturating KRAS^G12D^:GDP concentrations at 20 °C.

**Fig. S9.**
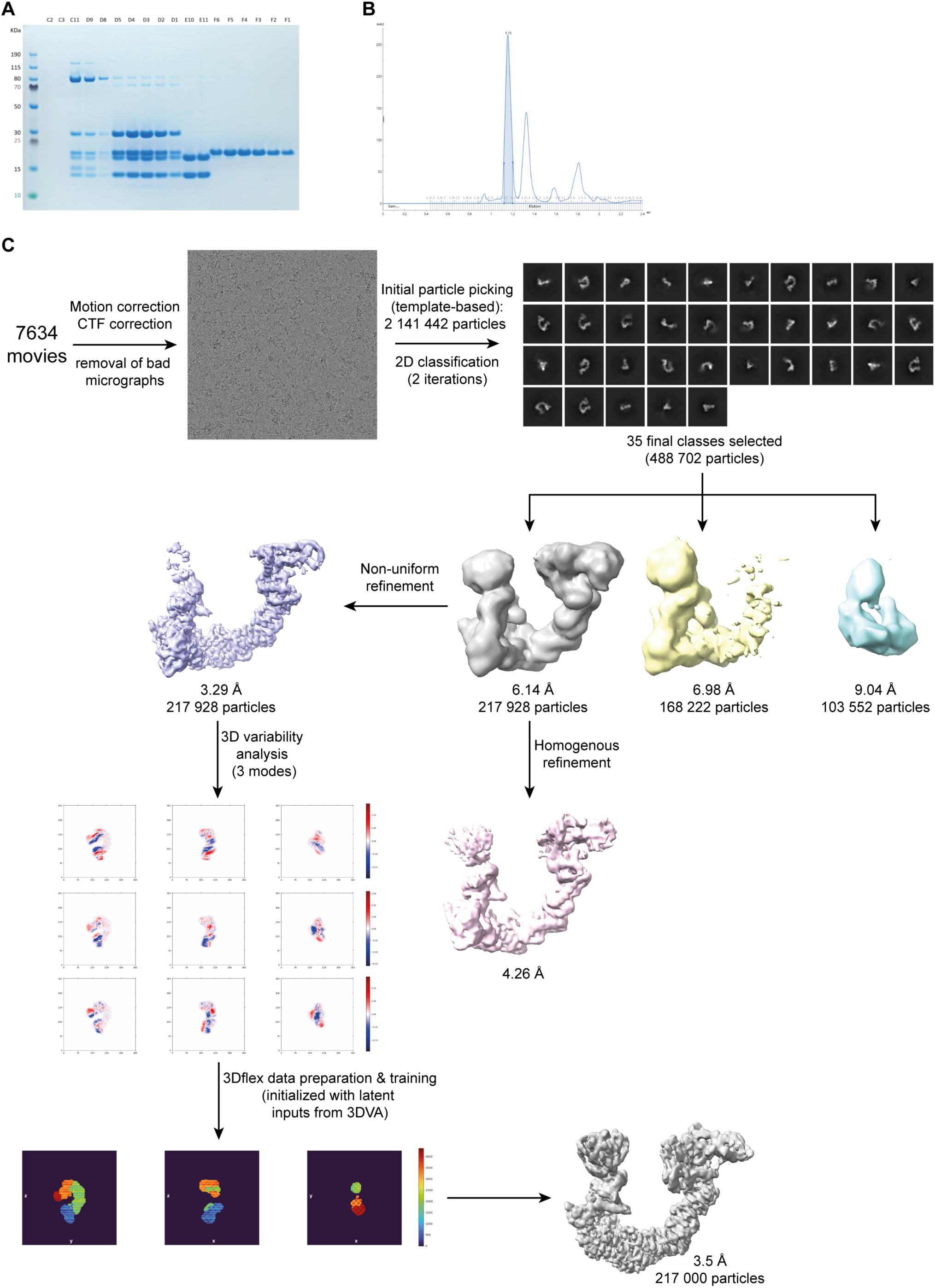
Sample preparation and image processing workflow for determination of the KRAS:ACBI3:VHL:EloB:EloC:Cul2:Rbx1-complex electron microscopy structure. (**A**) SDS-PAGE analysis of fractions eluted from (**B**) Superdex 200 gel filtration of complex assembly reactions. The peak area highlighted in blue was applied to cryo-EM grids for structure determination, as described in the materials and methods section. Lanes C11 and D9 of the SDS-PAGE analysis are the fractions flanking the peak, showing its composition of six proteins. (**C**) Cryo-EM image collection, pre-processing, processing and refinement workflow. Additional details can be found in the materials and methods section.

**Fig. S10.**
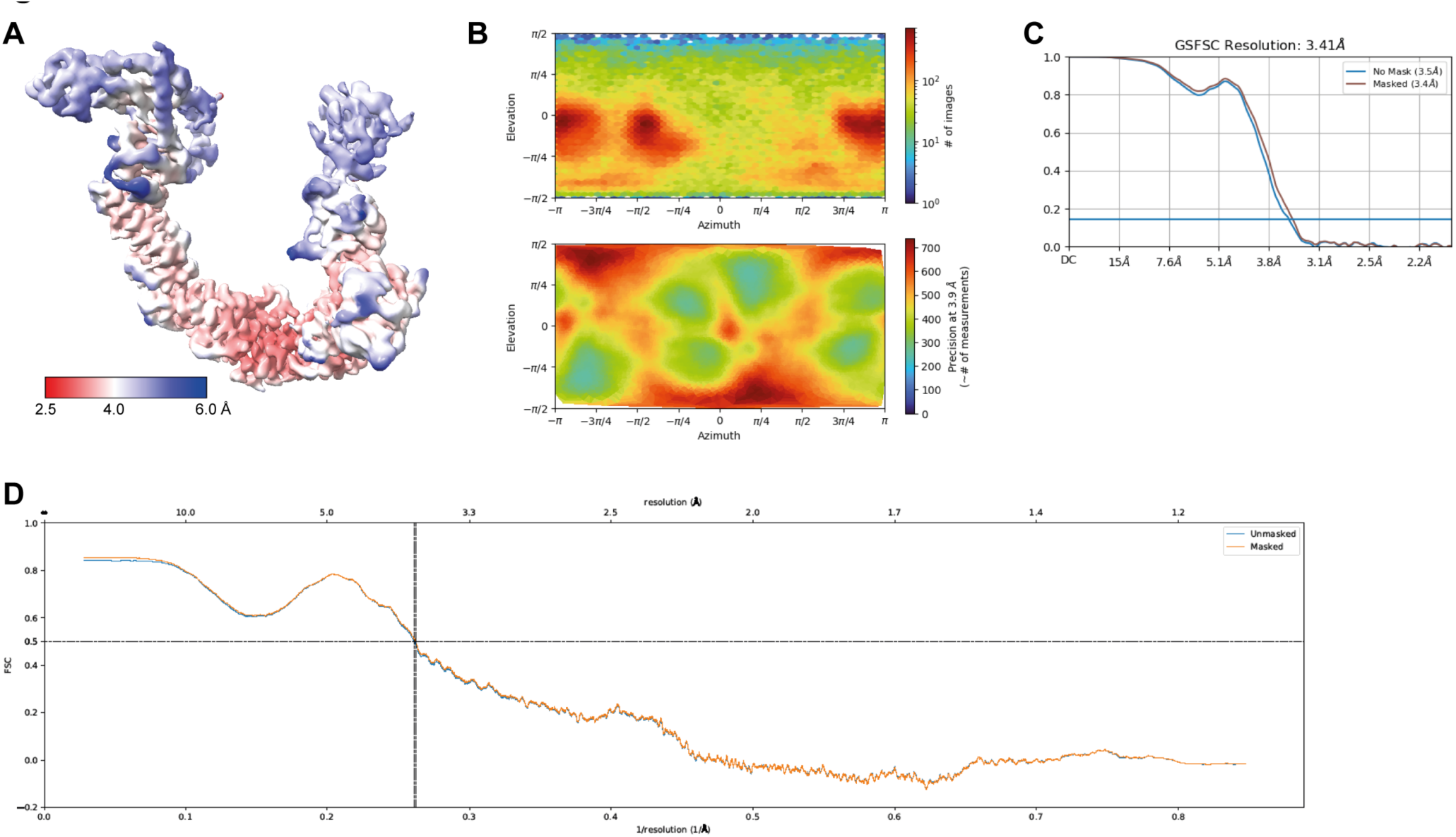
Cryo-EM data analysis of KRAS:ACBI3:VHL:EloB:EloC:Cul2:Rbx1 complexes. (**A**) Local resolution estimation of the complex, volume colored by resolution in Å. (**B**) Angular distribution of particle orientations of non-uniform refinement as described in fig S9. (**C**) Half-map to half-map Fourier shell correlation of 3Dflex refinement. (**D**) Model-to-map Fourier shell correlation of final refinement.

**Fig. S11.**
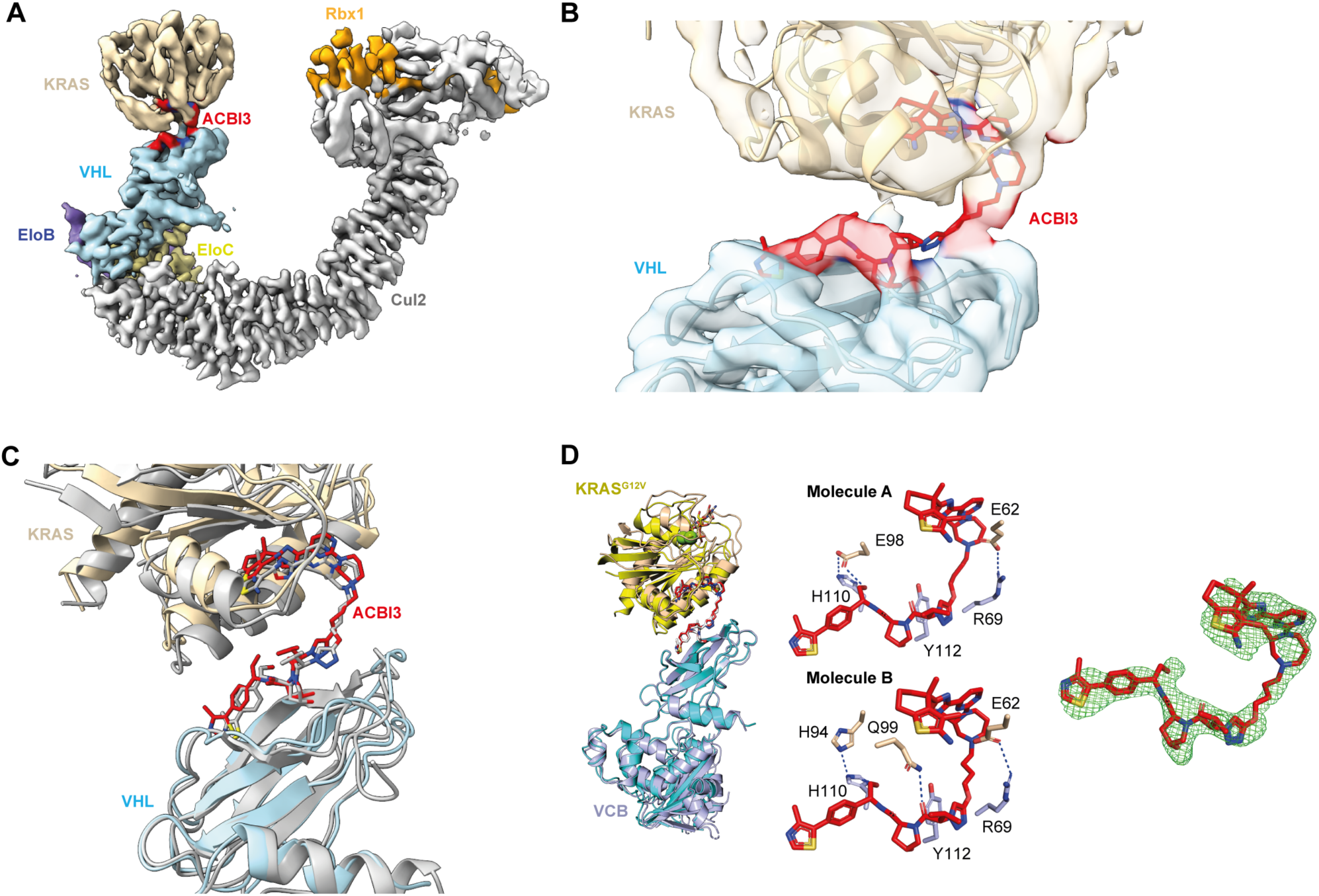
Structural characterization of ACBI3 ternary complexes by X-ray crystallography and cryo-EM microscopy. (**A**) Experimental density of the KRAS^G12V^:ACBI3:VCB:Cul2:Rbx1:complex by cryo-EM. (**B**) Interface between KRAS (sand) and VHL (blue) formed by ACBI3 (red). Shown is cryo-EM density on top of a cartoon representation, with ACBI3 as stick-model. (**C**) Overlay of the ACBI3 complex structures determined by cryo-EM (sand/orange/blue) and X-ray-crystallography (grey). Alignment was performed on VHL. (**D**) Left: superposition of ternary crystal structures of ACBI3 (red):VCB (light blue):KRAS^G12D^ (wheat), and compound **4** (white):VCB (teal):KRAS^G12V^ (yellow) displaying conserved ternary binding orientations (RMSD = 1.21 Å). Middle: Views of the ACBI3 ternary complex PROTAC binding sites (molecules A and B) displaying side chains of residues involved in potential strong protein:protein interactions (blue dotted lines). Right: Fo-Fc omit map contoured to 3 σ for ACBI3.

**Fig. S12.**
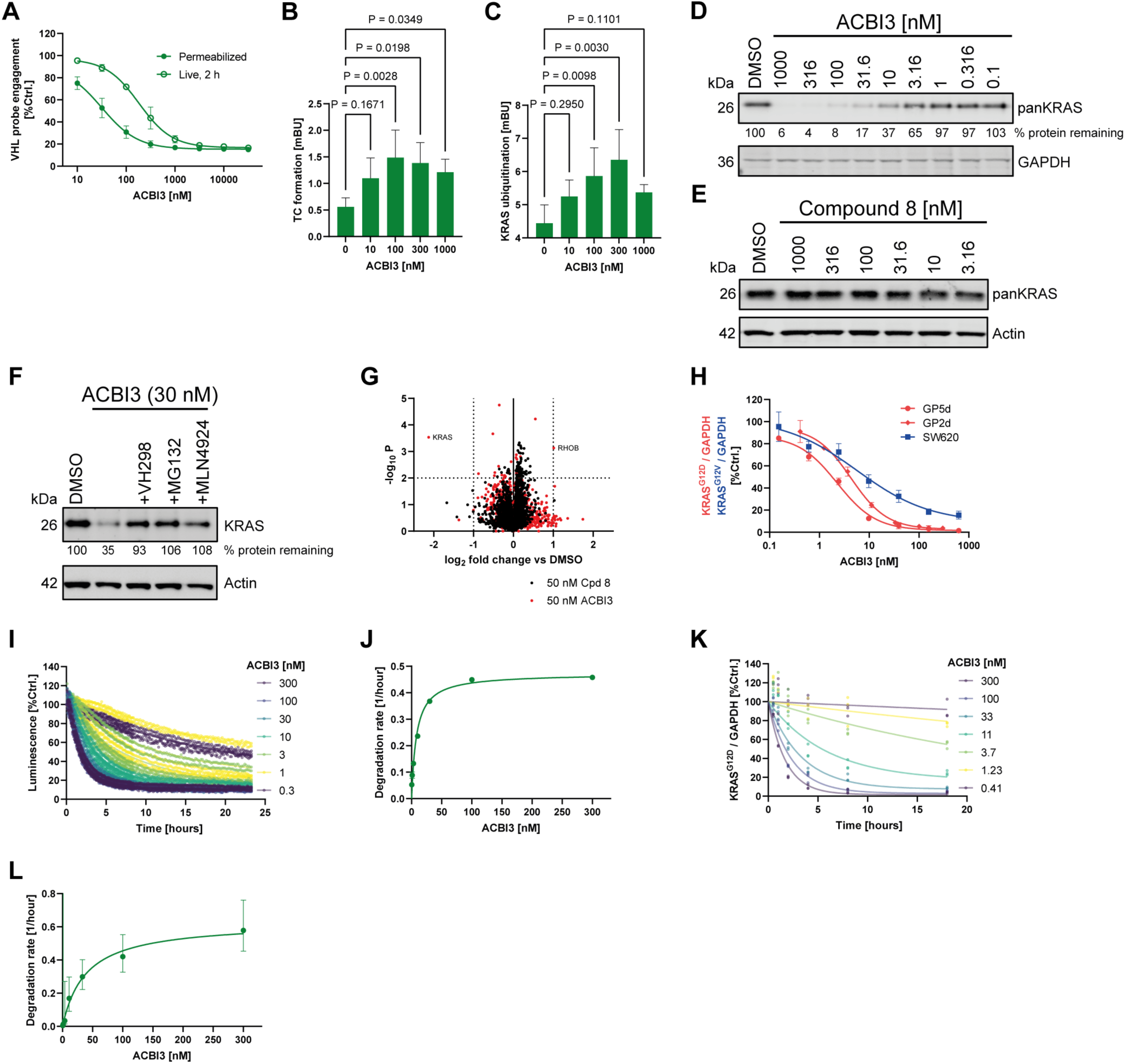
Degradation parameters, intracellular target engagement and selectivity of ACBI3. (**A**) VHL engagement by NanoBRET in live or permeabilized HEK293 cells for ACBI3 (N=3, SD). (**B**) Detection of VCB:ACBI3:KRAS^G12D^ complexes in GP5d cells (4 hours, N=3, SD, one-way ANOVA, Dunnet correction). (**C**) KRAS^G12D^ ubiquitination in GP5d cells treated with ACBI3 (4 hours, N=3, SD, one-way ANOVA, Dunnet correction). (**D**) KRAS and GAPDH levels by Western blot of GP5d cells treated with ACBI3 or (**E**) inactive stereoisomer compound **8** (4 hours), numbers indicate normalized KRAS levels *vs* controls (N=3). (**F**) KRAS levels in GP5d cells treated with ACBI3 in presence or absence of MLN4924 or VH298 by Western blot (4 hours), numbers indicate normalized KRAS levels *vs* controls (N=3). (**G**) Whole cell proteomics MS analysis of GP2d cells treated with 50 nM ACBI3 or inactive stereoisomer compound **8** (8 hours, N=3). HRAS (log_2_ fold change -0.0006, -log P 0.001) and NRAS (log_2_ fold change -0.12, -log P 0.52) levels were not significantly affected. (**H**) Degradation of KRAS^G12D^ or KRAS^G12V^ in GP5d, GP2d or SW620 cells by ACBI3, respectively, by capillary electrophoresis (24 hours – 18 hours for GP2d, N=3, SD). (**I**) Live cell degradation time course of HiBiT-KRAS^G12D^ in presence of varying concentrations of ACBI3 (N=6). (**J**) Degradation rate by live cell degradation *vs* concentration for ACBI3 (mean and 95% CI). (**K**) Degradation timecourse of endogenous KRAS^G12D^ in GP2d cells at varying concentrations of ACBI3 by capillary electrophoresis (N=3). (**L**) Degradation rate for endogenous KRAS^G12D^ *vs* concentration for ACBI3 (mean and 95% CI).

**Fig. S13.**
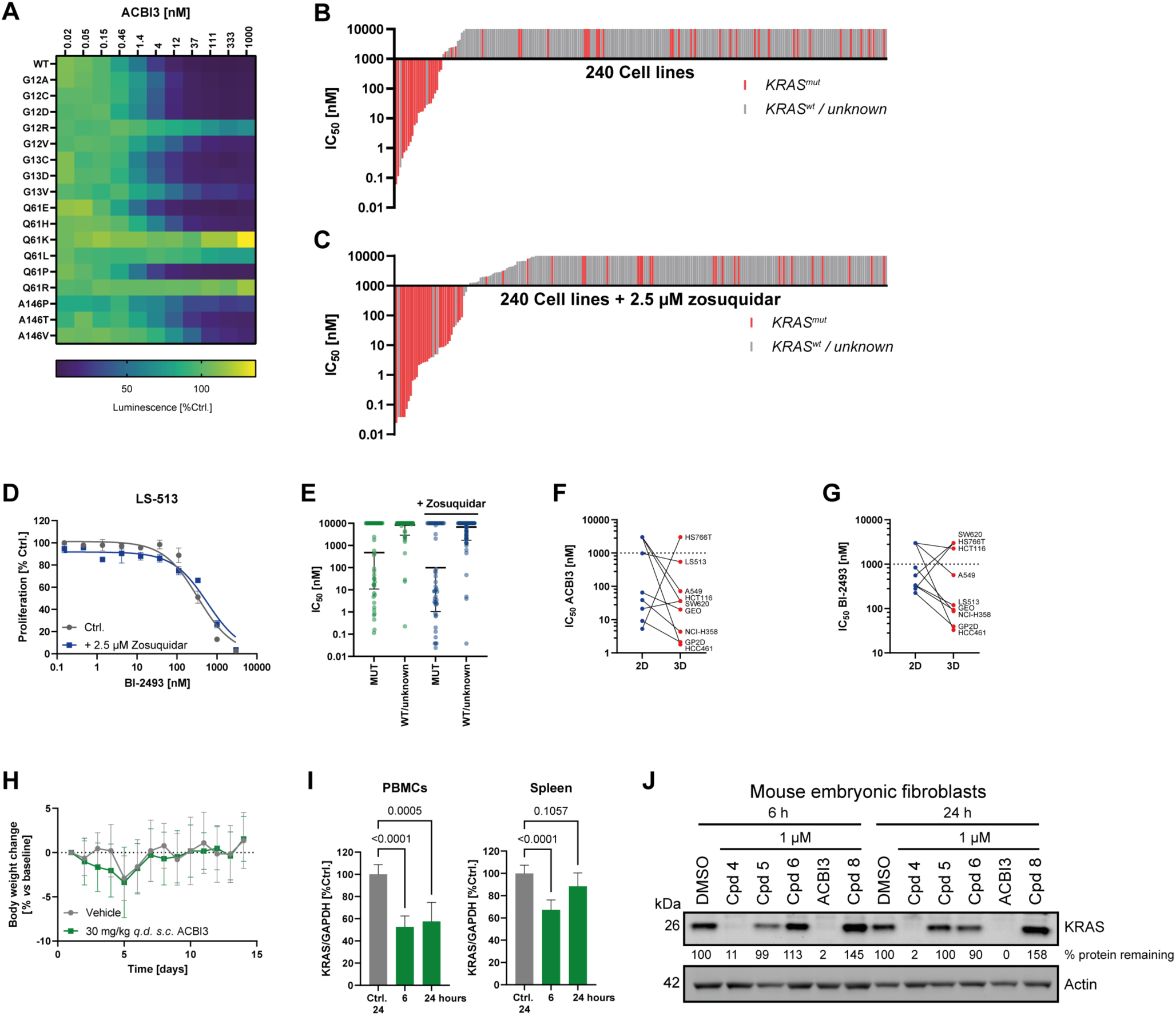
Extended cellular profiling and supplemental data of *in vivo* experiments for ACBI3. (**A**) Degradation of indicated retrovirally transduced HiBiT-KRAS mutants by ACBI3 (24 hours, N=3). (**B**) Proliferation inhibition data for 240 cancer cell lines for ACBI3, bars represent *IC_50_* per cell line (5 days, N=3). (**C**) Proliferation inhibition data for 240 cancer cell lines for ACBI3 in presence of 2.5 µM Zosuquidar, bars represent *IC_50_* per cell line (5 days, N=3). (**D**) Dose dependent inhibition of proliferation of LS-513 cells by BI-2493 in presence or absence of 2.5 µM of Zosuquidar (5 days, N=3, SD). (**E**) Cumulative proliferation data for ACBI3 in 240 cell line panel in presence or absence of 2.5 µM Zosuquidar for KRAS^WT^ *vs* KRAS mutant cell lines (geometric mean, geometric SD). (**F-G**) *IC_50_* of dose dependent 2D (N=3, 5 days) *vs* 3D (N=3, 10 days) proliferation assays of indicated cell lines for (**F**) ACBI3 or (**G**) BI-2493. (**H**) Effect of ACBI3 or vehicle treatment on body weight of GP2d tumor bearing mice (N=10, mean and SD). (**I**) *In vivo* degradation of KRAS in mouse PBMCs or spleens upon *s.c. q.d.x3* dosing of 30 mg/kg ACBI3. (N=5, mean and SD). (**J**) *In vitro* degradation of murine KRAS in mouse embryonic fibroblasts cells by 1 µM compound **4**, ACBI3 and corresponding inactive stereoisomers compound **5**, -**6** and -**8** by Western blot, numbers indicate normalized KRAS levels *vs* controls (N=3).

**Table S1.** Data summary biophysical characterization.

|  |  | Cpd 1 | Cpd 2 | Cpd 3 | Cpd 4 | Cpd 5 | ACBI3 |
| --- | --- | --- | --- | --- | --- | --- | --- |
| SPR | VCB $K_D$ [nM] | | 737 ± 388 (N=3) | 80 ± 24 (N=3) | 50 ± 9 (N=3) | | 11.1 ± 0.3 * (N=2) |
| | VCB $t_{1/2}$ [s] | | n.d. | 67 | 111 | | 2 100 * |
| | VCB $k_{on}$ [1/(M s)] | | n.d. | 1.36E+05 | 1.41E+05 | | 3.36E+05 * |
| | VCB+KRAS<sup>G12D</sup> $K_D$ [nM] | | 20 ± 5 (N=3) | 30 ± 6 (N=3) | 26 ± 5 (N=3) | | 6 ± 1 * (N=2) |
| | VCB+KRAS<sup>G12D</sup> $t_{1/2}$ [s] | | 159 | 103 | 230 | | 138 * |
| | VCB+KRAS<sup>G12D</sup> $k_{on}$ [1/(M s)] | | 2.24E+05 | 2.42E+05 | 1.26E+05 | | 8.19E+05 * |
|  | Cooperativity alpha [-] |  | 36.9 | 2.7 | 1.9 |  | 0.2 |
| | KRAS<sup>G12D</sup> GDP $K_D$ [nM] | N.D. | 240 ± 31 (N=3) | 13 ± 2 (N=3) | 11 ± 1 (N=3) | 25 ± 5 (N=3) | 5 ± 1 (N=3) |
| | KRAS<sup>G12V</sup> GDP $K_D$ [nM] | 25 ± 12 (N=4) | 136 ± 1 (N=3) | 9 ± 1 (N=3) | 8 ± 0 (N=3) | 13 ± 5 (N=3) | 4 ± 1 (N=3) |
| FP | VCB $K_D$ [nM] | | 7 187 ± 2 552 (N=3) | 340 ± 128 (N=3) | 110 ± 38 (N=3) | 1 350 ± 370 (N=3) | 5 ± 2.5 (N=3) |
| | VCB+KRAS<sup>G12D</sup> $K_D$ [nM] | | 15 ± 2 (N=3) | 20 ± 3 (N=3) | 16 ± 1 (N=3) | 670 ± 120 (N=3) | 4 ± 1 (N=3) |
|  | Cooperativity alpha [-] |  | 479.1 | 17.0 | 6.9 | 2.0 | 1.3 |

SPR, surface plasmon resonance; FP, fluorescence polarization \* Values are derived from single cycle experiment kinetic fitting. Binding affinities towards KRAS<sup>G12D</sup>, KRAS<sup>G12V</sup>, and VHL-ElonginC-ElonginB complex (VCB), as well as VCB ± KRAS<sup>G12D</sup> measuring cooperativities (alpha), ternary complex affinity and half-lives. Errors are ± standard deviation with repeats (N) specified in brackets. Ternary complex cooperativity (alpha) is calculated as ratio between VCB $K_D$ / VCB + KRAS<sup>G12D</sup> $K_D$ values measured in the same run, respectively.

**Table S2.** Data summary *in vitro* degradation, pathway engagement and proliferation.

|  | Cell line / condition | Compound 2 | Compound 3 | Compound 4 | Compound 5 | MRTX1133 | sotorasib | ACBI3 | BI-2865 |
| --- | --- | --- | --- | --- | --- | --- | --- | --- | --- |
| ***DC<sub>50</sub>* 6 hours (95% CI) [nM]** | **GP2d (KRAS<sup>G12D</sup>)** | n.d. | 53.4<br>(4.1-812) | 7.3<br>(3.7-14.1) | >1 000 | >1 000 | n.d. | n.d. | n.d. |
|  | **SW620 (KRAS<sup>G12V</sup>)** | n.d. | 159<br>(40.9-725.6) | 24<br>(13.1-43.6) | >1 000 | >1 000 | n.d. | n.d. | n.d. |
|  | **NCI-H358** | n.d. | 158.9<br>(0-infinity) | 19.6<br>(3.3-99.1) | >1 000 | n.d. | >1 000 | n.d. | n.d. |
|  | **A375** | n.d. | 750.6<br>(162.3-infinity) | 36.7<br>(17.7-75.2) | >1 000 | >1 000 | n.d. | n.d. | n.d. |
|  | **HEK293** | n.d. | >1 000 | 12.8<br>(1.9-212.6) | >1 000 | >1 000 | n.d. | n.d. | n.d. |
| ***DC<sub>50</sub>* 24 hours (95% CI) [nM]** | **GP5d (KRAS<sup>G12D</sup>)** | 607.1<br>(387.3-764.3) | 31.8<br>(22.7-43.4) | 0.9<br>(0.7-1.1) | n.d. | n.d. | n.d. | 2.1<br>(1.7-2.6) | n.d. |
|  | **GP5d (HiBiT-KRAS<sup>G12D</sup>)** | 493.6<br>(295.7-613.5) | 68.1<br>(51.6-90) | 1.4<br>(1.3-1.6) | n.d. | n.d. | n.d. | 0.7<br>(0.5-0.8) | n.d. |
|  | **GP5d (HiBiT-KRAS<sup>G12D</sup>)**<br>**10% mouse serum** | n.d. | n.d. | 33.1<br>(28.2-38.8) | n.d. | n.d. | n.d. | 5.1<br>(3.8-6.9) | n.d. |
|  | **GP5d (HiBiT-KRAS<sup>G12D</sup>)**<br>**10% human serum** | n.d. | n.d. | 1.9<br>(1.5-2.4) | n.d. | n.d. | n.d. | 0.8<br>(0.6-0.9) | n.d. |
|  | **GP2d (KRAS<sup>G12D</sup>)** | n.d. | 76.3<br>(4.3-1 985) | 3.6<br>(1.7-7.5) | >1 000 | >1 000 | n.d. | 3.9<br>(3-4.9) | n.d. |
|  | **SW620 (KRAS<sup>G12V</sup>)** | 1 203<br>(757.1-1 569) | 278.1<br>(17.4-infinity) | 12.6<br>(5.8-29) | >1 000 | >1 000 | n.d. | 6.9<br>(3.7-12.9) | n.d. |
|  | **NCI-H358** | n.d. | 240.9<br>(34.7-infinity) | 25.1<br>(2.4-295.2) | >1 000 | n.d. | >1 000 | n.d. | n.d. |
|  | **A375** | n.d. | 455.1<br>(61.6-infinity) | 33.4<br>(15.1-58.4) | >1 000 | >1 000 | n.d. | n.d. | n.d. |
|  | **HEK293** | n.d. | >1 000 | 86.7<br>(6.1-977.2) | >1 000 | >1 000 | n.d. | n.d. | n.d. |

Table S2 (continued).
|  |  |  |  |  |  |  |  |  |  |
| --- | --- | --- | --- | --- | --- | --- | --- | --- | --- |
| **$D_{max}$ 24 hours (95% CI) [%]** | **GP5d (KRAS<sup>G12D</sup>)** | 100 (93-100) | 99 (94-100) | 100 (98-100) | n.d. | n.d. | n.d. | 99 (97-100) | n.d. |
|  | **GP5d (HiBiT-KRAS<sup>G12D</sup>)** | 100 (80-100) | 95 (88-100) | 94 (91-96) | n.d. | n.d. | n.d. | 95 (92-97) | n.d. |
|  | **GP2d (KRAS<sup>G12D</sup>)** | n.d. | 96 (57-100) | 96 (85-100) | <15 | <15 | n.d. | 99 (96-100) | n.d. |
|  | **SW620 (KRAS<sup>G12V</sup>)** | 100 (90-100) | 88 (83-96) | 89 (79-100) | <15 | <15 | n.d. | 83 (77-100) | n.d. |
|  | **NCI-H358** | n.d. | 81 (43-100) | 86 (53-100) | <25 | n.d. | <15 | n.d. | n.d. |
|  | **A375** | n.d. | 76 (71-80) | 100 (82-100) | <15 | <15 | n.d. | n.d. | n.d. |
|  | **HEK293** | n.d. | 48 (42-54) | 100 (39-100) | <20 | <15 | n.d. | n.d. | n.d. |
| **$\lambda_{max}$ (95% CI) [1/hour]** | **GP5d (HiBiT-KRAS<sup>G12D</sup>, live mode)** | 0.4 (0.3-0.6) | 0.6 (0.5-0.6) | 0.5 (0.4-0.6) | n.d. | n.d. | n.d. | 0.5 (0.4-0.5) | n.d. |
| **$D_{max50}$ (95% CI) [nM]** |  | 3 447 (1 480-9658) | 56 (46-69) | 17 (9-33) | n.d. | n.d. | n.d. | 8 (5-14) | n.d. |
| **$\lambda_{max}/D_{max50}$ [1/(μM hour)]** |  | 0.1 | 11 | 29 | n.d. | n.d. | n.d. | 58 | n.d. |
| **$IC_{50}$ pERK (95% CI) [nM]** | **GP2d** | n.d. | 771.6 (210.9-1554) | 3.1 (1.4-6.9) | 1 091.0 (206.0-infinity) | 1.3 (0.5-2.9) | n.d. | n.d. | n.d. |
|  | **SW620** | n.d. | 708.4 (48.8-2225.0) | 9.2 (3.4-25.7) | >1 000 | >1 000 | n.d. | n.d. | n.d. |
|  | **NCI-H358** | n.d. | >1 000 | 8.2 (1.0-112.1) | >1 000 | n.d. | 29 (8.5-107.7) | n.d. | n.d. |
|  | **A375** | n.d. | >1 000 | >1 000 | >1 000 | >1 000 | n.d. | n.d. | n.d. |
|  | **HEK293** | n.d. | >1 000 | >1 000 | >1 000 | >1 000 | n.d. | n.d. | n.d. |

**Table S2 (continued).**
|  |  |  |  |  |  |  |  |  |  |
| --- | --- | --- | --- | --- | --- | --- | --- | --- | --- |
| ***IC*<sub>50</sub> DUSP6**<br>**(95 % CI)**<br>**[nM]** | **GP2d** | n.d. | 128.9<br>(76.1-221) | 3.3<br>(2-5.4) | 245.3<br>(61.2-3760) | 0.1<br>(0-0.3) | n.d. | n.d. | n.d. |
|  | **SW620** | n.d. | 205.3<br>(127.5-341.7) | 6.2<br>(4.4-8.6) | 428.7<br>(212.3-1 110.0) | 309.8<br>(195.0-522.0) | n.d. | n.d. | n.d. |
|  | **NCI-H358** | n.d. | 248.3<br>(124.2-527.9) | 11.4<br>(8.9-14.6) | 138.4<br>(97.1-197.7) | n.d. | 5.6<br>(3.6-8.8) | n.d. | n.d. |
|  | **A375** | n.d. | >1 000 | >1 000 | >1 000 | >1 000 | n.d. | n.d. | n.d. |
|  | **HEK293** | n.d. | >1 000 | >1 000 | >1 000 | >1 000 | n.d. | n.d. | n.d. |
| ***IC*<sub>50</sub>**<br>**Proliferation**<br>**(95 % CI)**<br>**[nM]** | **GP2d** | n.d. | 373.7<br>(233.6-607.3) | 8.3<br>(5.5-12.8) | 1738<br>(561.8-19 428) | 2.4<br>(1.7-3.4) | n.d. | n.d. | 250.7<br>(141.7-449.1) |
|  | **SW620** | n.d. | 231.7<br>(187-287.5) | 11.0<br>(9.6-12.6) | 993.0<br>(602.2-1 732.0) | 1 523.0<br>(1 004.0-2 458.0) | n.d. | n.d. | 207.7<br>(144.6-297.7) |
|  | **NCI-H358** | n.d. | 983.2<br>(723.9-1331) | 85.4<br>(61.7-118.4) | 1 755.0<br>(945.9-3 931.0) | n.d. | 12.3<br>(10.1-14.9) | n.d. | 613.1<br>(416.1-922.8) |
|  | **A375** | n.d. | >1 000 | >1 000 | >1 000 | >1 000 | n.d. | n.d. | >1 000 |
|  | **HEK293** | n.d. | >1 000 | >1 000 | >1 000 | >1 000 | n.d. | n.d. | >1 000 |

n.d. not determined

**Table S3.**
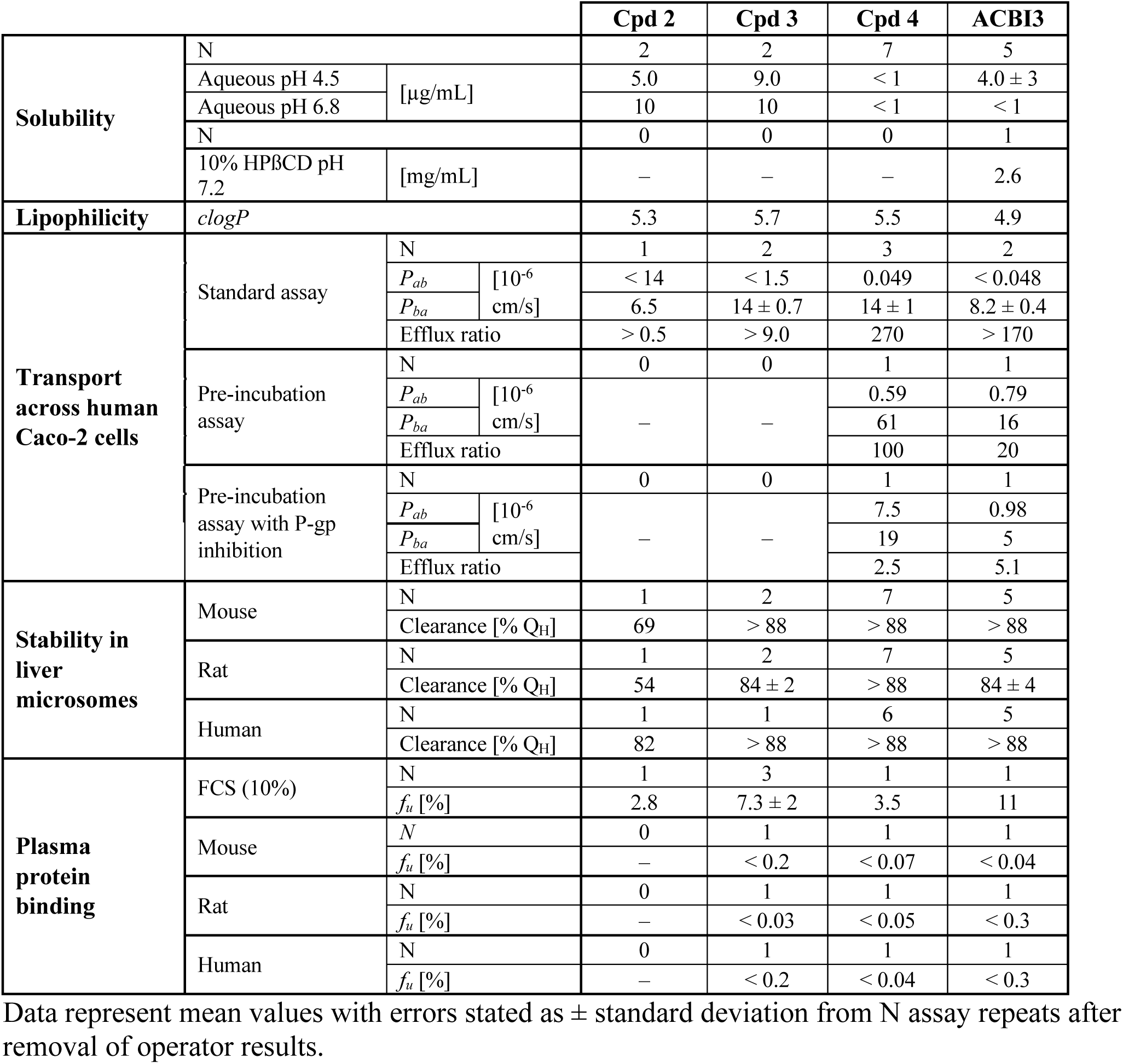
Physicochemical properties and in vitro DMPK parameters.

**Table S4.** Compound pharmacokinetics in NMRI mice.

|  | **Cpd 4** |  | **ACBI3** |  |
| --- | --- | --- | --- | --- |
| **Administration route** | *i.v.* | *s.c.* | *i.v.* | *s.c.* |
| **Formulation** | HP-β-CD | HP-β-CD,<br>Ringer's<br>solution | HP-β-CD | PEG-400,<br>Transcutol,<br>Kolliphor HS 15 |
| **Dose** [mg/kg] | 5 | 30 | 2 | 30 |
| **AUC<sub>0-inf</sub>** [nmol/(L hour)] | 1 890 | 5 330 | 857 | 9 040 |
| **Clearance** [mL/(min kg)] | 45 | – | 39 | – |
| ***V<sub>ss</sub>*** [L/kg] | 5.3 | – | 1.6 | – |
| **Mean residence time**<br>[hours] | 2.0 | 6.8 | 0.67 | 6.8 |
| **Bioavailability** [%] | – | 47 | – | 70 |

All data represent mean of N=3 animals.

**Table S5.** Crystallographic data collection and refinement statistics.

|  | KRAS<sup>G12C</sup>-<br>compound **1**<br>(PDB: 8QUG) | KRAS<sup>G12V</sup>:compound<br>**3**:VCB<br>(PDB: 8QW6) | KRAS<sup>G12V</sup>:compound<br>**4**:VCB<br>(PDB: 8QW7) | KRAS<sup>G12D</sup>:ACBI3:VCB<br>(PDB: 8QVU) |
| --- | --- | --- | --- | --- |
| **Data collection** |  |  |  |  |
| Space group | P 4<sub>3</sub> 2<sub>1</sub> 2 | P 1 | P 1 2<sub>1</sub> 1 | P 1 2<sub>1</sub> 1 |
| Cell dimensions |  |  |  |  |
| *a*, *b*, *c* (Å) | 91.0, 91.0, 52.7 | 70.2, 72.3, 80.8 | 72.5, 125.9, 81.1 | 72.2, 120.9, 81.0 |
| $\alpha$, $\beta$, $\gamma$ (°) | 90, 90, 90 | 112.3, 100.6, 100.6 | 90, 111.6, 90 | 90, 111.6, 90 |
| Resolution (Å) | 64.4-1.6 (1.654-1.562) * | 66.3-2.21 (2.46-2.21) | 67.4-2.4 (2.70-2.41) | 75.2-2.2 (2.54-2.2) |
| *R*<sub>meas</sub> | 0.043 (1.696)* | 0.065 (0.577) | 0.035 | 0.153 (1.417) |
| *I*/σ*I* | 26.2 (1.4)* | 7.4 (1.5) | 13.2 (1.6) | 8.0 (1.6) |
| Completeness (%) | 95.3 (67.5)* | 82.3 (54.3) | 92.8 (64.1) | 90.6 (60.6) |
| Redundancy | 12.6 | 2.5 | 7.0 | 7.1 |
| **Refinement** |  |  |  |  |
| Resolution (Å) | 40-1.6 | 46.4-2.2 | 64.67-2.4 | 55.86-2.7 |
| No. reflections | 26 177 | 39 189 | 30 834 | 32 178 |
| *R*<sub>work</sub> / *R*<sub>free</sub> | 19.1/22.2 | 25.50/29.49 | 24.59/27.26 | 24.16/28.04 |
| No. atoms (non-hydrogen) |  |  |  |  |
| Protein | 1 371 | 7 869 | 7 566 | 7 720 |
| Ligand | 32 | 140 | 132 | 151 |
| Water | 204 | 48 | 24 | 66 |
| *B*-factors |  |  |  |  |
| Protein | 35.7 | 47.78 | 56.85 | 61.32 |
| Ligand | 33.7 | 33.7 | 46.02 | 40.3 |
| Water | 50.0 | 35.3 | 46.97 | 35.64 |
| R.m.s. deviations |  |  |  |  |
| Bond lengths (Å) | 0.010 | 0.025 | 0.016 | 0.008 |
| Bond angles (°) | 1.00 | 2.06 | 1.82 | 1.16 |

\*Values in parentheses are for highest-resolution shell.

**Table S6.** Cryo-EM data collection, refinement and validation statistics.

|  |  |
| --- | --- |
|  | KRAS<sup>G12V</sup>-ACBI3-VHL-EloB-EloC-Cul2-Rbx1 complex<br>(EMD-18657)<br>(PDB 8QU8) |
| Data collection and processing |  |
| Magnification | 165.000× |
| Voltage (kV) | 300 |
| Electron exposure (e<sup>-</sup>/Å<sup>2</sup>) | 40 |
| Defocus range (μm) | -0.8 to -2.0 |
| Pixel size (Å) | 0.745 |
| Symmetry imposed | C1 |
| Initial particle images (no.) | 2 414 422 |
| Final particle images (no.) | 217 000 |
| Map resolution (Å) | 3.5 |
| FSC threshold | 0.143 |
| Refinement |  |
| Initial model used (PDB code) | 8QUG (KRAS),<br>5N4W (VHL/EloB/EloC/Cul2/Rbx1) |
| Model-to-map resolution (global, Å) |  |
| FSC threshold 0.143 | 2.3 |
| FSC threshold 0.5 | 3.8 |
| Correlation coefficient (mask) | 0.73 |
| Map sharpening | DeepEMhancer |
| Model composition |  |
| Non-hydrogen atoms | 10 855 |
| Protein residues | 1 325 |
| Ligand atoms | 72 |
| B factors (Å<sup>2</sup>) |  |
| Protein | 133 |
| Ligand | 173 |
| R.m.s. deviations |  |
| Bond lengths (Å) | 0.003 |
| Bond angles (°) | 0.663 |
| Validation |  |
| MolProbity score | 1.91 |
| Clashscore | 9.40 |
| Poor rotamers (%) | 0.17 |
| Ramachandran plot |  |
| Favored (%) | 93.75 |
| Allowed (%) | 6.10 |
| Disallowed (%) | 0.15 |

**Data S1.**

Chemical synthesis procedures are provided as a separate file.

**Data S2.**

Cell line panel proliferation data for ACBI3 are provided as a separate file.

## General information

Commercially available dry solvents were used from Sigma Aldrich. All reagents unless otherwise noted were commercially available and purchased from Sigma Aldrich, Combi Blocks, ABCR, Fluorochem, Activate or Enamine, at least 95% pure and used without further purification. All reactions were carried out in oven-or flame dried glassware under nitrogen atmosphere. Normal phase TLC was carried out on pre-coated silica plates (Kieselgel 60 F254, BDH) with visualization via UV light (UV 254 and/or 365 nm) and/or basic potassium permanganate solution. Isolute® phase separator columns from Biotage were used. Flash column chromatography was performed using either a Teledyne Isco Combiflash Rf or Rf200i or a Biotage Isolera One with prepacked Redisep RF normal phase disposable columns. Reverse phase chromatography was carried out using Biotage SNAP-C18 columns. Strong cation exchange (SCX) chromatography was carried out using Biotage Isolute SCX-2 columns. NMR Spectra were recorded on Bruker 400 MHz, 500 MHz or 600MHz spectrometers as specified. Chemical shifts are quoted in ppm and referenced to the residual solvent signals: ^1^H NMR *δ* (ppm) = 7.26 (CDCl_3_-*d*), ^13^C NMR *δ* (ppm) = 77.2 (CDCl_3_-*d*), ^1^H NMR *δ* (ppm) = 2.50 (DMSO-*d_6_*), ^13^C NMR *δ* (ppm) = 39.5 (DMSO-*d_6_*). Signal splitting patterns are described as singlet (s), doublet (d), triplet (t), quartet (q), quintet (quin.), multiplet (m), broad (br) or a combination thereof. Coupling constants (*J*) are measured in Hertz (Hz). Diastereomeric ratios (dr) were calculated using the ratios of NMR integrals.

### Analytical MS Methods and instrumentation

**Method 1:** HRMS data was recorded using a LTQ Orbitrap XL (Thermo Scientific) coupled with a Triversa Nanomate Nanospray ion source (ADVION Bioscience Inc.) The mass calibration was performed using the Pierce LTQ Velos ESI positive ion calibration solution from Thermo Scientific (Product Nr. 88323).

MS parameters: The scan window was set to 50–400 amu with a maximum injection time of 500 ms and 1 microscan. Resolution of the Orbitrap was 60000 with a mass accuracy ≤ 5 ppm. The ion mode set to positive with a capillary temperature 200 °C and voltage of 60 eV. The tube lens potential was set to 110 eV. 12 NanoESI voltage was 1.45 kV and the N_2_ gas pressure set to 0.45 psi. Total sample volume was 5 µL and the acquisition time was 0.4 sec, with 10 scans of averaging per spectrum Sample dilution: 10 mM DMSO stock solution was diluted 1:200 in 50% MeOH +0.01% formic acid.

**Method 2**: Agilent Technologies 1200 series HPLC connected to an Agilent Technologies 6130 quadrupole LC/MS with an Agilent diode array detector. **Method 2a** was run under the following conditions: Waters Xbridge C18 column, 2.5 µm particle size, 2.1 x 20 mm. Run time 2.1 minutes, flow 1 mL/min, column temperature 60 °C and 5 µL injections. Solvent A (20mM NH_4_HCO_3_/NH_4_OH pH 9), solvent B (MS grade acetonitrile). Start 10% B, gradient 10% - 95% B from 0.0 - 1.5 min, 95% B from 1.5 - 2.0 min, gradient 95% - 10% B from 2.0 – 2.1 min. Purity was determined via UV detection with a bandwidth of 170 nm in the range from 230-400 nm. **Method 2b** used a Waters XBridge column (50 mm × 2.1 mm, 3.5 μm particle size) and the compounds were eluted with a gradient 5−95% acetonitrile/water + 0.1% formic acid (“acidic method”).

### Preparative purification methods and instrumentation

**Method 3**: Preparative HPLC was performed on a Gilson system with Waters C18 columns (50 mm x 150 mm; 10 µm particle size) and a gradient of 10% to 95% acetonitrile in water over 8 minutes and a flow rate of 150 mL/min, with a SunFire column and 0.1% formic acid (**Method 3a**) or a XBridge column and 0.5% ammonium bicarbonate/ammonia buffer pH 9 (**Method 3b**) in the aqueous phase.

**Method 4**: Preparative HPLC was performed on a Waters Prep 150 LC system with a Waters XBridge C18 column (100 mm x 19 mm; 5 μm particle size) and a gradient of 5% to 95% acetonitrile in water over 10 minutes and a flow rate of 25 mL/min, with 0.1% formic acid (**Method 4a**) or ammonia (**Method 4b**) in the aqueous phase.

### Method A: Chrial purification

Chiral separations were performed on a Agilent 1100/1200 system using a Chiralpak; Part. No. 85394; IE, 5 µm; 150 x 2.1 mm column. Solvents used were n-Heptan / EtOH + 0,1% DEA, 0-70% isocratic flow. Detection signal was UV 315 nm, (bandwidth 170, reference off) and the spectrum range: 190 – 400 nm; slit: 4 nm. Peak width > 0.0031 min (0.063 s response time, 80Hz), 0,5 µL standard injection and 1.2 mL/min flow at 45°C column temperature.

Stop time: 5 min

**Abbreviations used:** aq. for aqueous, BH_3_-DMS for borane-dimethylsulfide complex, Boc for *N-tert*-butyloxycarbonyl, Boc_2_O for di-*tert*-butyl dicarbonate, DCE for 1,2-dichloroethane, DCM for dichloromethane, DEA for diethylamine, DMA for *N*,*N*-dimethylacetamide, DMF for *N*,*N*-dimethylformamide, DIPEA for *N*,*N*-diisopropylethylamine, DMSO for dimethylsulfoxide, Et_3_N for triethylamine, EtOAc for ethyl acetate, HATU for 1-[bis(dimethylamino)methylene]-1*H*-1,2,3-triazolo[4,5-*b*]pyridinium 3-oxideHexafluorophosphate, KOAc for potassium acetate, MeOH for methanol, MsCl for methanesulfonyl chloride, PTSA for *p*-toluene sulfonic acid monohydrate, sat. for saturated, SOCl_2_ for thionyl chloride, THF for tetrahydrofuran.

## Synthesis of pan KRAS binders

### Synthesis of KRAS binder Compound 1

#### Intermediate Int-6

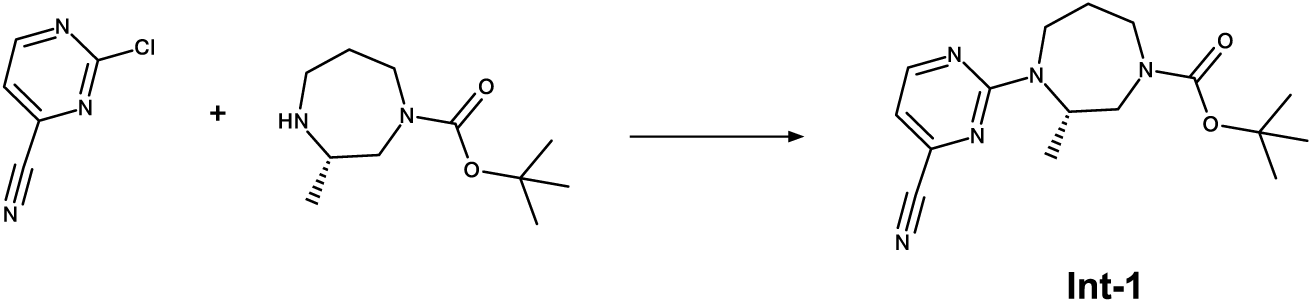

To a solution of (S)-tert-butyl-3-methyl-1,4-diazepane-1-carboxylate (846 mg, 3.948 mmol, 1.0 eq.) and 2-chloropyrimidine-4-carbonitrile (529 mg, 3.791 mmol, 1.0 eq) in DMSO (4 ml) is added TEA (1.1 ml, 7.896 mmol, 2.0 eq.) at rt. The reaction mixture is stirred at 80 °C for 1 h. After complete conversion the reaction mixture is cooled to rt and water and EtOAc is added. The phases are separated. The organic layer is washed with water, dried over sodium sulfate, then filtered and concentrated under reduced pressure to the get crude product which is purified by NP chromatography (0-20%, EtOAc/Heptane) to obtain intermediate Int-1 (754 mg, 2.376 mmol, 63 % yield).

^1^H NMR (DMSO-d_6_) δ: 8.63 (br d, J=4.6 Hz, 1H), 7.08-7.13 (m, 1H), 5.07-5.17 (m, 0.19H rotamer), 5.01 (br d, J=5.3 Hz, 0.84H rotamer), 4.13 (br d, J=14.4 Hz, 0.56H rotamer), 4.05 (br d, J=14.2 Hz, 0.49H rotamer), 3.64-3.94 (m, 2H), 2.75-3.12 (m, 2H), 1.46-1.80 (m, 2H), 1.27 (br s, 5.71H rotamer), 1.16 (s, 3.18H rotamer), 1.05 (s, 3H) Integrals may not add up to whole numbers due to overlapping rotamers or rotamers resonating under residual DMSO-d6 or HDO peaks

^13^C NMR (DMSO-d6) δ: 160.8, 160.6, 160.4, 153.9, 153.7, 140.5, 116.3, 112.1, 112.0, 78.5, 78.4, 52.2, 48.6, 48.3, 47.7, 47.6, 27.8, 27.7, 27.6, 26.1, 26.0, 15.0

More carbon peaks detected than present in the structure due to presence of rotamers

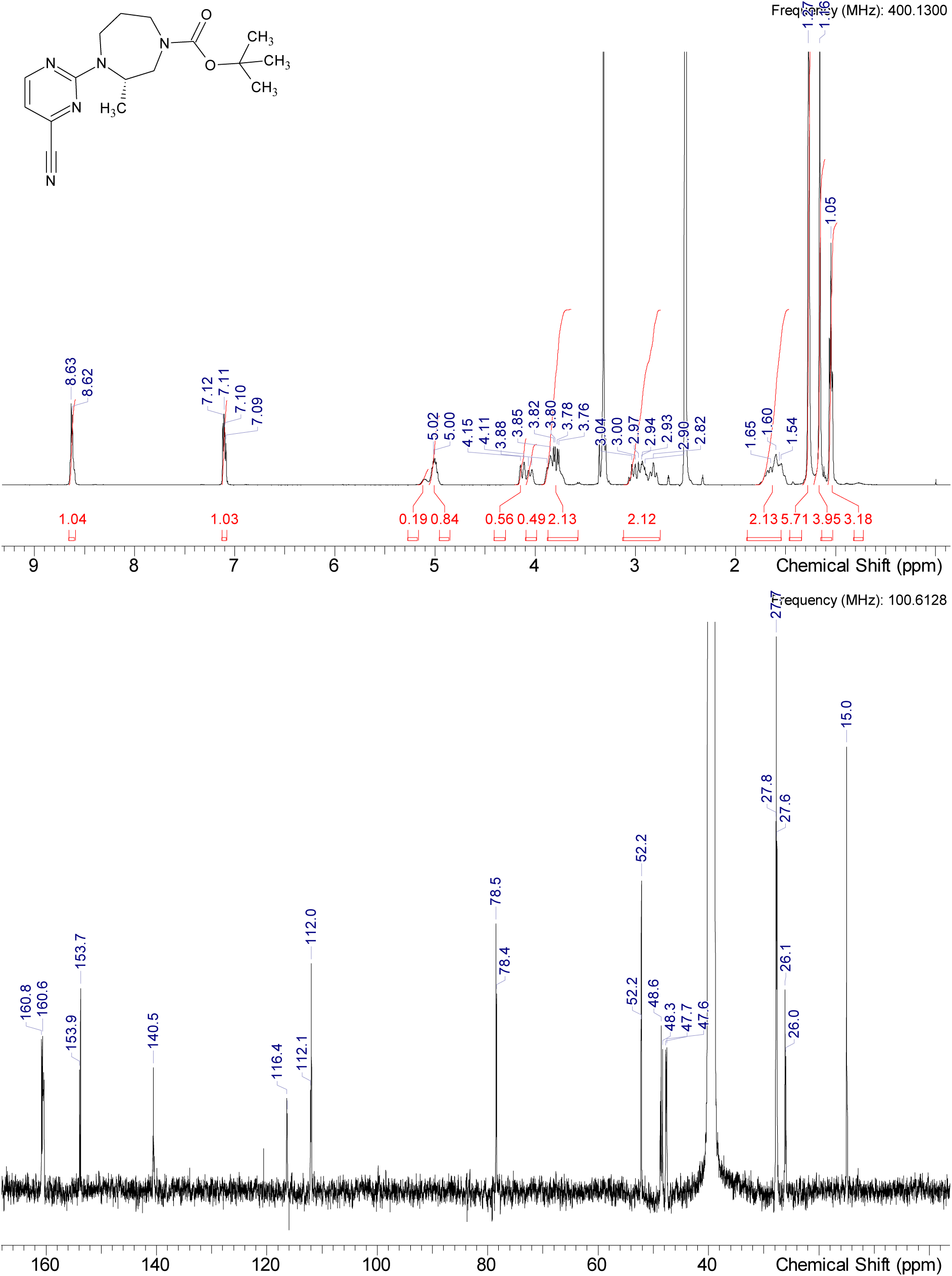

HRMS (m/z): [M+H]+ calcd. for C16H23N5O2 , 318.19245; found, 318.19245

#### Intermediate Int-2

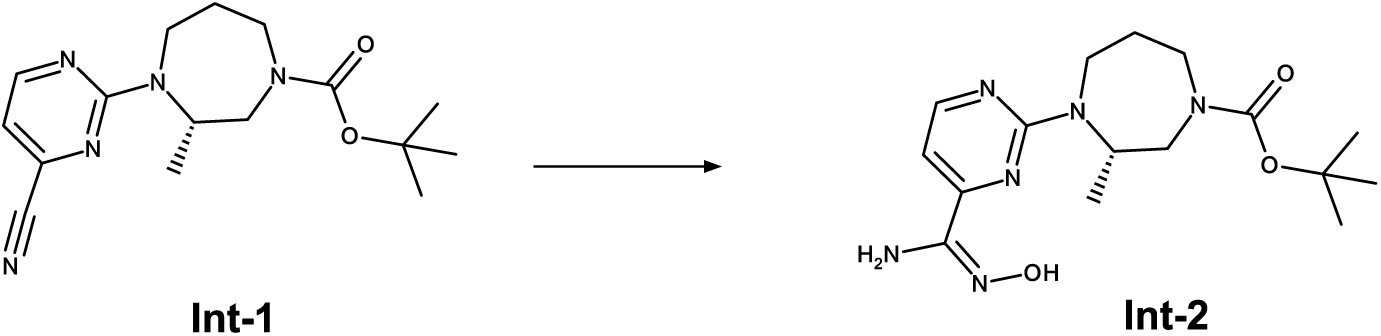

To a solution of Int-1 (33.85 g, 106.65 mmol, 1.0 eq) in EtOH (270 ml) is added hydroxylamine solution 50 % in water (13.05 ml, 213.30 mmol, 2.0 eq) at rt. The reaction mixture is stirred at 60 °C for 1 h. After complete conversion the reaction mixture is concentrated under reduced pressure to afford Int-2 (33.8 g, 102.7 mmol, 96 % yield) which is used for the next step without further purification.

^1^H NMR (DMSO-d_6_) δ: 10.06-10.21 (m, 1H), 8.29 (d, J=5.1 Hz, 1H), 6.90 (br dd, J=12.9, 4.8 Hz, 1H), 5.75 (br d, J=9.9 Hz, 2H), 5.10 (br d, J=14.2 Hz, 1H), 4.25 (br dd, J=57.4, 13.6 Hz, 1H), 3.63-3.92 (m, 2H), 3.25 (br t, J=13.3 Hz, 1H), 2.75-3.08 (m, 2H), 1.44-1.84 (m, 2H), 1.25 (br d, J=8.6 Hz, 5H), 1.15 (s, 3H), 1.10-1.31 (m, 1H), 1.05 (br d, J=3.3 Hz, 3H)

^13^C NMR (DMSO-d_6_) δ: 160.3, 157.8, 157.5, 154.0, 153.8, 148.7, 148.4, 103.5, 78.4, 52.6, 48.0, 27.7, 26.5, 15.3

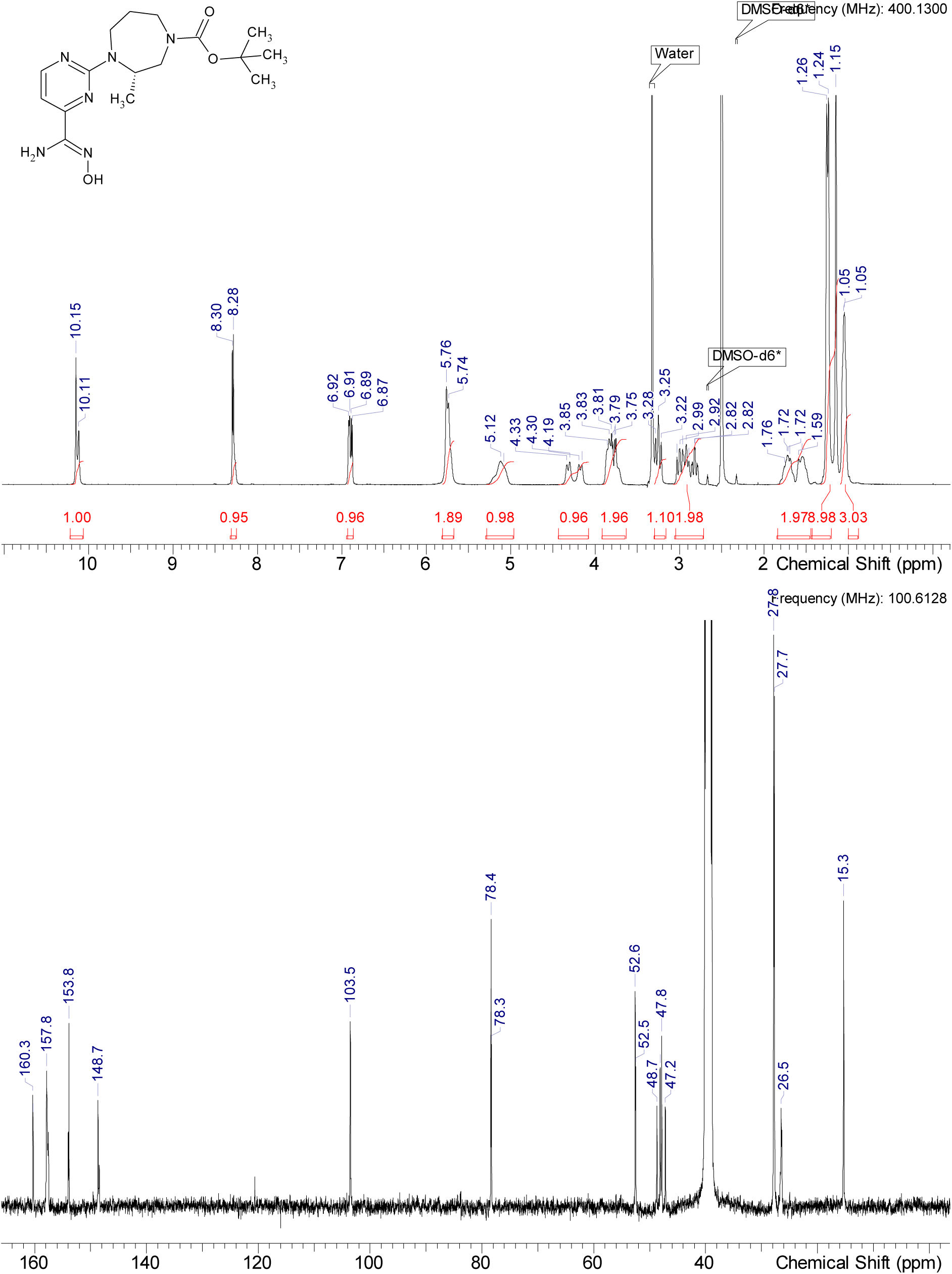

HRMS (m/z): [M+H]+ calcd. for C16H26N6O3 , 351.21371; found, 351.21371

#### Intermediate Int-4

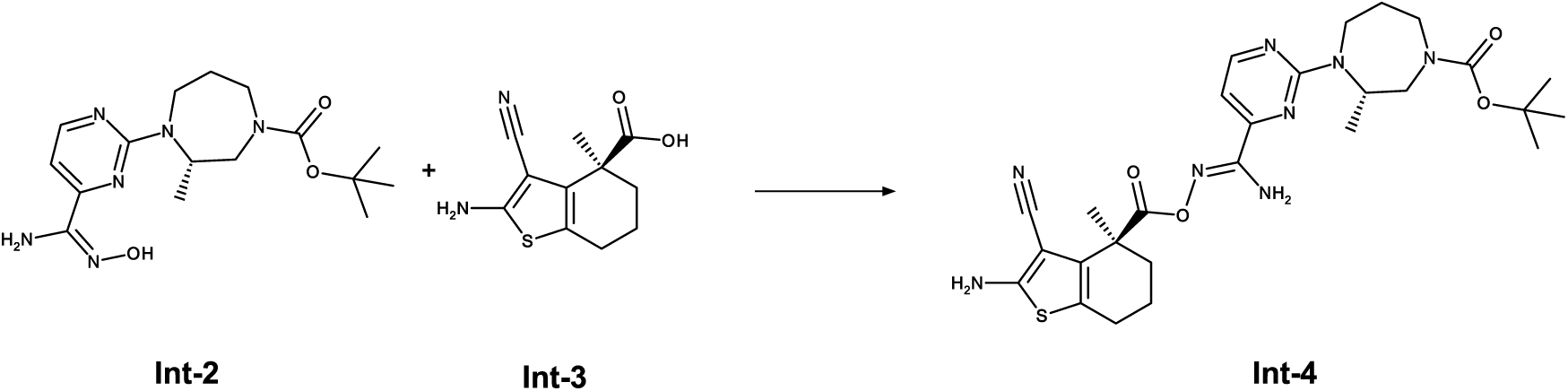

To a stirred solution of Int-2 (2.53 g, 10.70 mmol, 1.0 eq.) in DMSO (10 ml) are added TEA (2.17 g, 21.40 mmol, 2.0 eq.) and HATU (4.27 g, 11.24 mmol, 1.10 eq.) at rt. The mixture is stirred for 15 min at rt. Int-3 (3.75 g, 10.70 mmol, 1.0 eq) is added at rt and stirred overnight. After complete conversion the reaction mixture is diluted with water and EtOAc. The phases are separated. The organic layer is washed with water, dried over sodium sulfate, filtered, and concentrated under reduced pressure to get the crude product. This crude material is purified by normal phase column chromatography (0-5%, DCM/MeOH, 15 ml 7N NH3 / l solvent) to afford Int-4 (4.59 g, 8.071 mmol, 75 % yield).

^1^H NMR (DMSO-d_6_) δ: 8.30-8.52 (m, 1H), 6.89-7.08 (m, 3H), 5.41-6.84 (m, 2H), 5.00-5.26 (m, 1H), 4.28-4.50 (m, 0.48H rotamer), 4.18 (br d, J=14.7 Hz, 0.36H rotamer), 3.60-3.94 (m, 2H), 3.15-3.31 (m, 2H), 2.97 (br dd, J=22.8, 12.9 Hz, 2H), 2.41-2.47 (m, 1H), 2.12-2.21 (m, 1H), 1.74 (br d, J=8.1 Hz, 4H), 1.60 (s, 4H), 1.26 (br s, 5H rotamer), 1.15 (s, 4H rotamer), 1.05 (br s, 3H) Integrals may not add up to whole numbers due to overlapping rotamers or rotamers resonating under residual DMSO-d6 or HDO peaks

^13^C NMR (DMSO-d_6_) δ: 171.9, 163.9, 160.9, 159.5, 156.4, 156.2, 154.3, 154.0, 153.6, 132.9, 132.9, 121.1, 119.6, 117.6, 117.5, 105.4, 105.3, 82.7, 78.9, 78.8, 53.0, 52.9, 49.2, 48.8, 48.3, 47.8, 44.8, 36.1, 28.3, 28.2, 24.4, 24.3, 20.4, 15.9, 15.7

More carbon peaks detected than present in the structure due to presence of rotamers

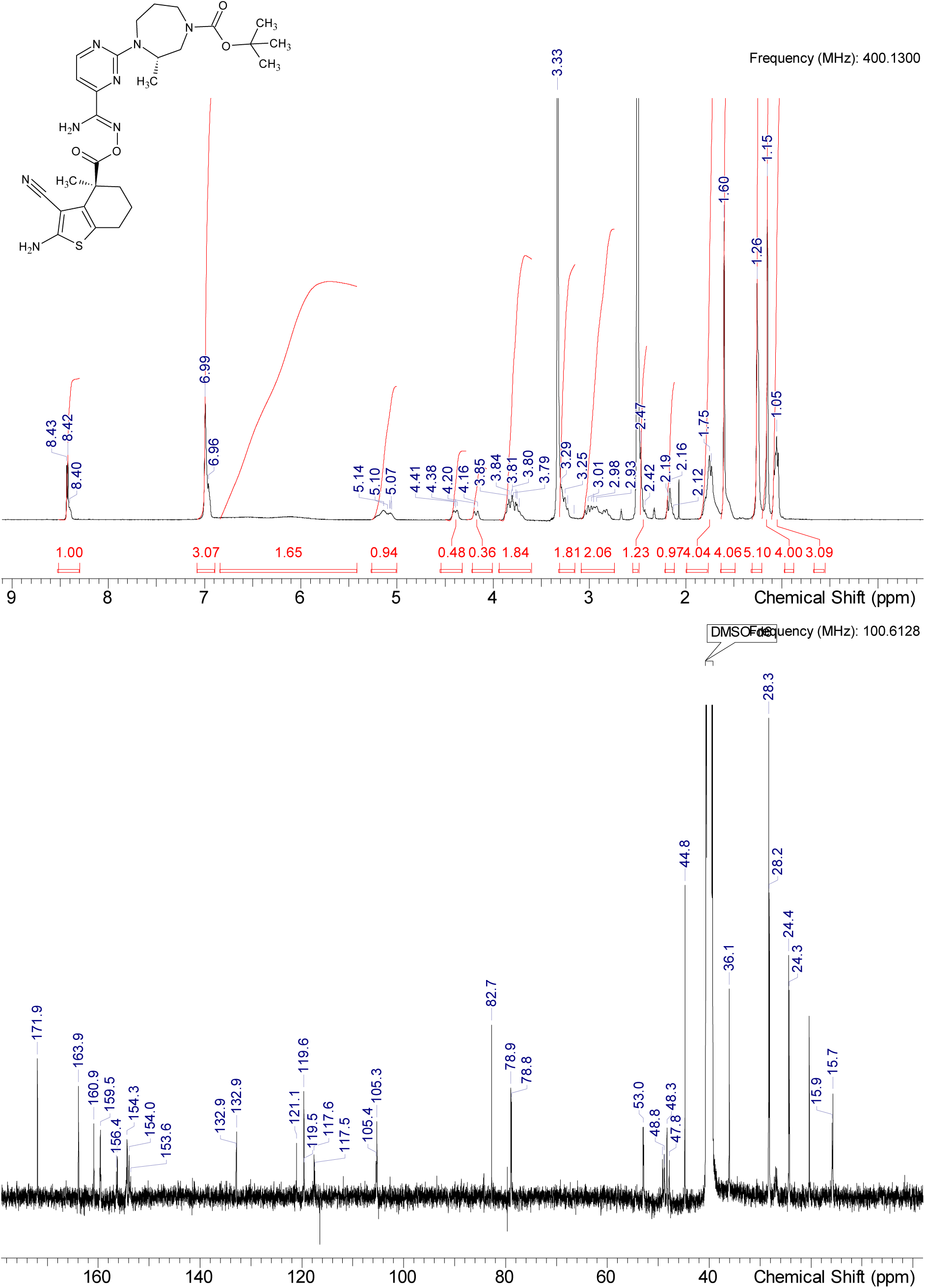

HRMS (m/z): [M+H]+ calcd. for C27H36N8O4S , 569.26530; found, 469.21271 (-Boc)

#### Intermediate Int-5

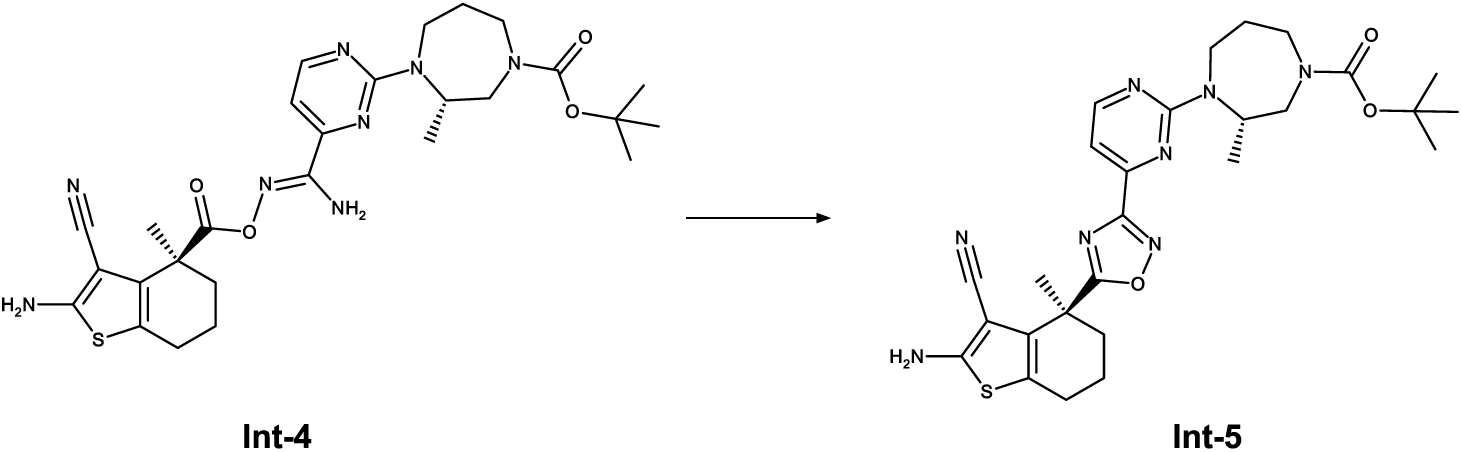

To a stirred solution of Int-4 (300 mg, 0.528 mmol, 1.0 eq) in THF (3 mL) is added DBU (160 mg, 1.06 mmol, 2.0 eq) at rt. The reaction mixture is stirred at 70 °C overnight. After complete conversion the reaction mixture is concentrated under reduced pressure to get the crude product. The crude product is purified by normal phase column chromatography (0-2%, DCM/MeOH, 15 ml 7N NH3 / l solvent ) to afford Int-5 (231 mg, 0.419 mmol, 79 % yield.

^1^H NMR (DMSO-d_6_) δ: 8.55 (br d, J=4.4 Hz, 1H), 7.12 (br d, J=5.0 Hz, 1H), 7.08 (s, 2H), 5.22-5.38 (m, 0.45H rotamer), 5.11 (br d, J=5.0 Hz, 0.52H rotamer), 4.07-4.36 (m, 1H), 3.65-3.92 (m, 2H), 2.69-3.11 (m, 2H), 2.52-2.57 (m, 2H), 2.01-2.25 (m, 2H), 1.84 (br s, 3H), 1.79 (s, 3H), 1.63-1.76 (m, 1H), 1.47-1.63 (m, 1H), 1.27 (br s, 2H rotamer), 1.17 (br d, J=11.3 Hz, 7H rotamer), 1.06 (br d, J=6.3 Hz, 3H)

Integrals may not add up to whole numbers due to overlapping rotamers or rotamers resonating under residual DMSO-d6 or HDO peaks

^13^C NMR (DMSO-d_6_) δ: 184.9, 167.6, 167.5, 164.2, 161.3, 160.5, 154.5, 154.3, 154.0, 131.8, 120.1, 120.0, 116.2, 116.1, 108.1, 107.9, 81.8, 79.0, 79.0, 78.9, 53.1, 52.8, 49.0, 48.4, 48.2, 37.4, 37.2, 28.3, 28.2, 28.0, 26.8, 26.6, 25.2, 24.0, 19.9, 15.9, 15.7

More carbon peaks detected than present in the structure due to presence of rotamers

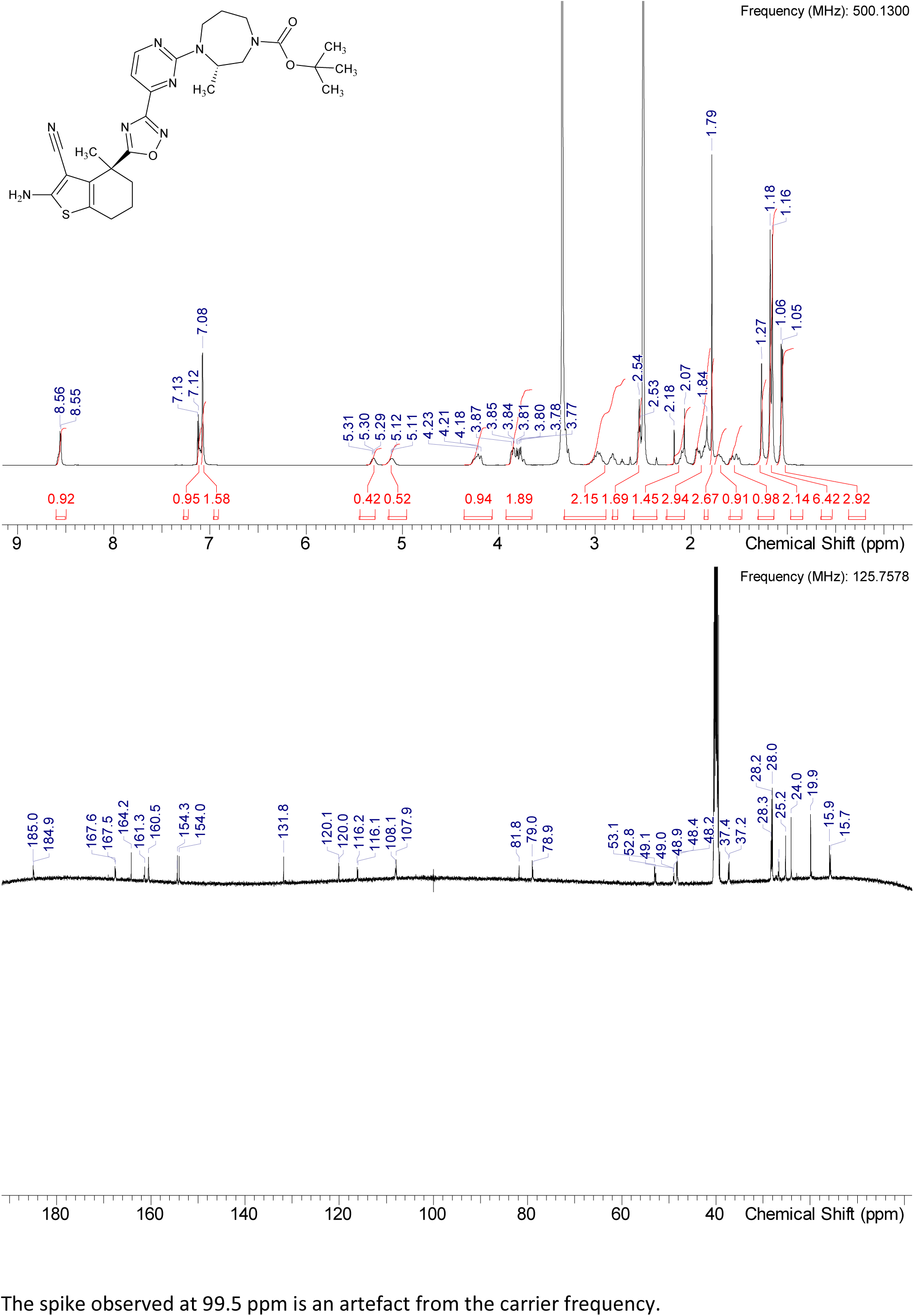

HRMS (m/z): [M+H]+ calcd. for C27H34N8O3S , 551.25473; found, 551.25421

#### Compound 1

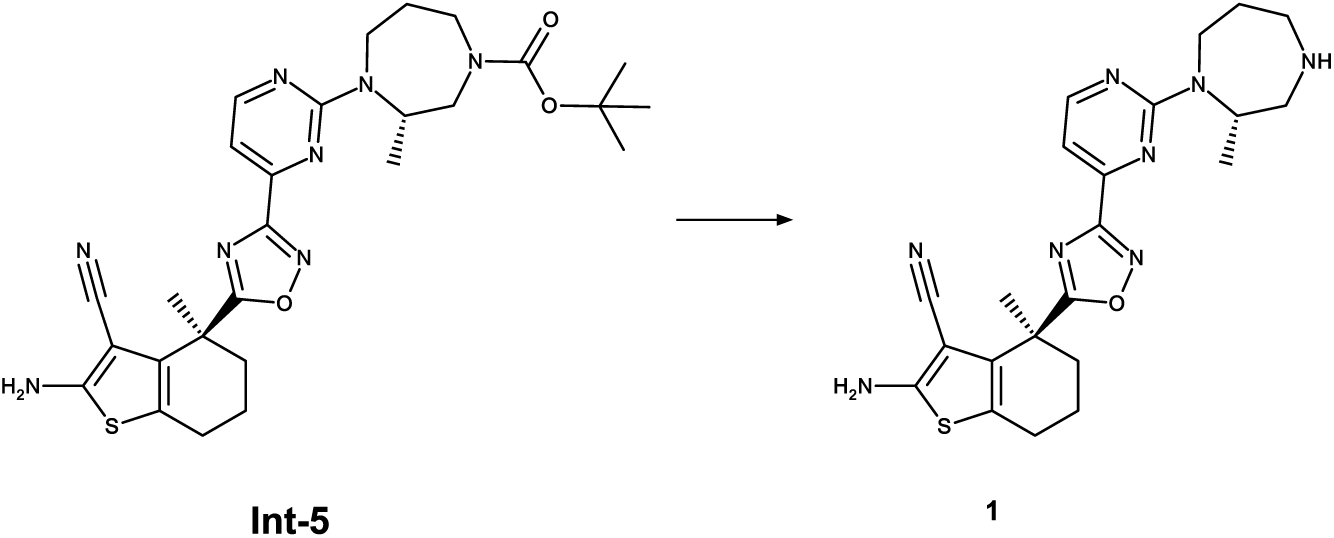

To a stirred solution of Int-5 (19.01 g, 34.52 mmol, 1.0 eq.) in MeOH (350 mL) is added conc. HCl (32.88 mL, 345.21 mmol, 10.0 eq.) at rt. The reaction mixture is stirred at 50 °C for 2 h. After complete conversion the reaction mixture is concentrated under reduced pressure and diluted with water. The aqueous phase is extracted with DCM. The combined organic layers are dried over sodium sulfate, filtered and concentrated under reduced pressure to afford crude 1 (14.54 g, 32.27 mmol, 94 % yield) which can be used for the next step without further purification.

^1^H NMR (DMSO-d_6_) δ: 8.56 (d, J=4.7 Hz, 1H), 7.11 (br d, J=4.1 Hz, 1H), 7.07 (s, 2H), 4.57-4.78 (m, 1H), 4.17-4.39 (m, 1H), 3.12-3.29 (m, 3H), 2.98 (br d, J=11.7 Hz, 1H), 2.54 (br d, J=5.0 Hz, 4H), 2.03-2.13 (m, 1H), 1.80-1.99 (m, 3H), 1.78 (s, 3H), 1.62 (br s, 2H), 0.97-1.08 (m, 3H) 1H under DMSO

^13^C NMR (DMSO-d_6_) δ: 184.5, 167.2, 163.8, 161.1, 160.1, 153.5, 131.4, 119.8, 115.8, 107.4, 81.4, 54.9, 54.6, 51.7, 50.1, 36.8, 29.3, 24.8, 23.6, 19.5, 15.7, 1C under DMSO

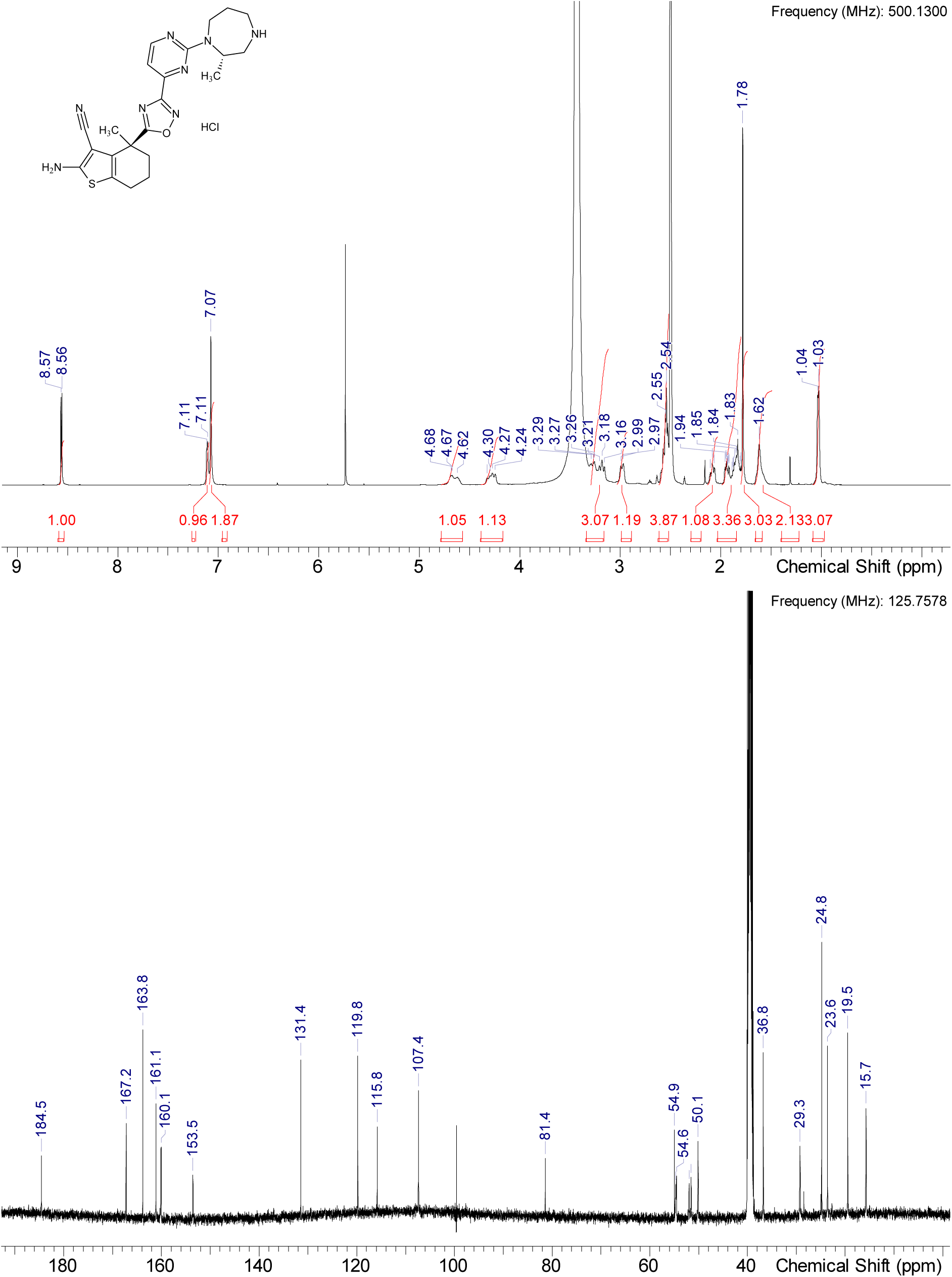

HRMS (m/z): [M+H]+ calcd. for C22H26N8OS , 451.20231; found, 451.20224

#### Compound S-1

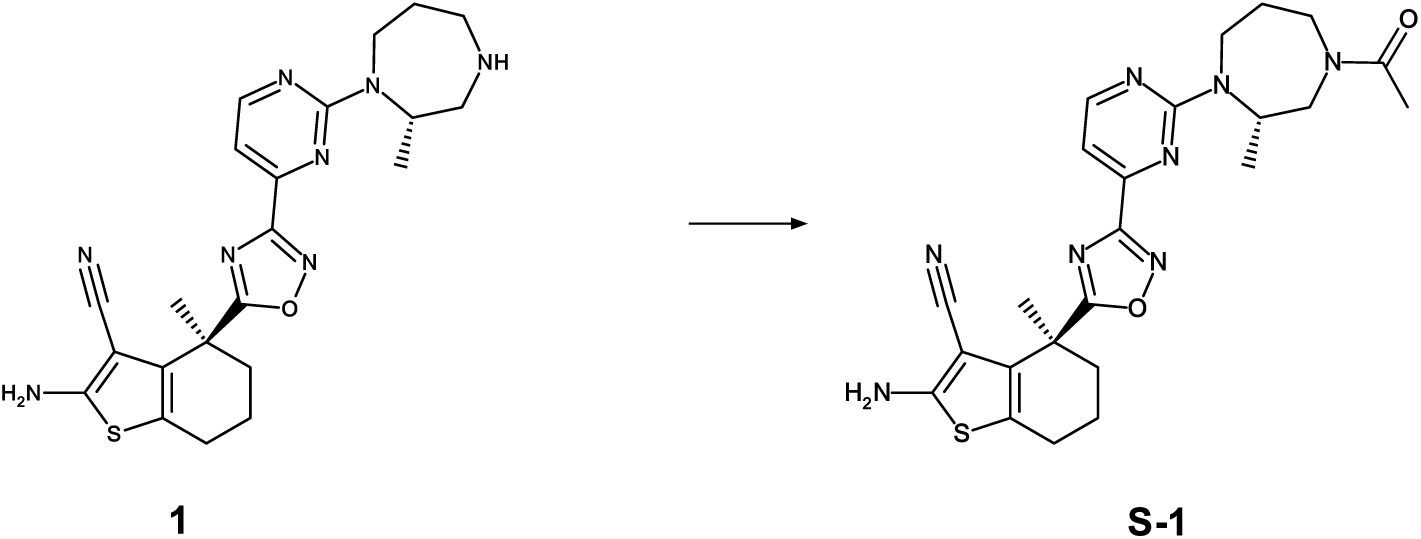

To a stirred solution of 1 (30 mg, 0.067 mmol, 1.0 eq.) in NMP (1 mL) was added acetic acid (0.011 mL, 0.200 mmol, 3.0 eq.) at 0°C was added HATU (37 mg, 0.093 mmol, 1.4 eq.) and DIPEA (0.058 mL, 0.333 mmol, 5.0 eq.). The mixture was stirred for 15 min and afterwards diluted with water and acetonitrile, purified by RP chromatography (Method 3b) to obtain S-1 (28 mg, 0.057 mmol, 85 % yield).

^1^H NMR (DMSO-d_6_) δ: 8.57 (d, J=5.0 Hz, 0.41H rotamer), 8.56 (d, J=4.7 Hz, 0.61H rotamer), 7.14 (d, J=5.0 Hz, 0.52H rotamer), 7.13 (br d, J=5.0 Hz, 0.53H rotamer), 7.08 (s, 2H), 5.00-5.25 (m, 0.44H rotamer), 4.81-4.99 (m, 0.6H rotamer), 4.10-4.45 (m, 2H), 3.96-4.09 (m, 0.44H rotamer), 3.79 (br d, J=13.9 Hz, 0.59H rotamer), 3.35-3.41 (m, 1H), 2.95 (br dd, J=13.4, 12.1 Hz, 0.61H rotamer), 2.73 (br s, 0.47H rotamer), 2.52-2.60 (m, 2H), 2.04-2.14 (m, 1H), 1.75-1.99 (range, 9H rotamer), 1.71 (br s, 2H), 1.13 (br s, 1.3H rotamer), 1.06 (br s, 1.77H rotamer)

Integrals may not add up to whole numbers due to overlapping rotamers or rotamers resonating under residual DMSO-d6 or HDO peaks

^13^C NMR (DMSO-d_6_) δ: 184.5, 168.9, 168.8, 168.5, 167.0, 167.0, 163.7, 163.7, 160.9, 160.3, 160.0, 153.5, 131.3, 119.6, 115.7, 107.8, 107.6, 81.3, 81.3, 53.4, 51.0, 49.9, 49.0, 48.3, 48.1, 46.4, 36.7, 28.5, 28.2, 26.0, 25.7, 24.7, 23.5, 21.0, 20.5, 19.4, 15.6, 15.4, 15.3, 15.1

More carbon peaks detected than present in the structure due to presence of rotamers

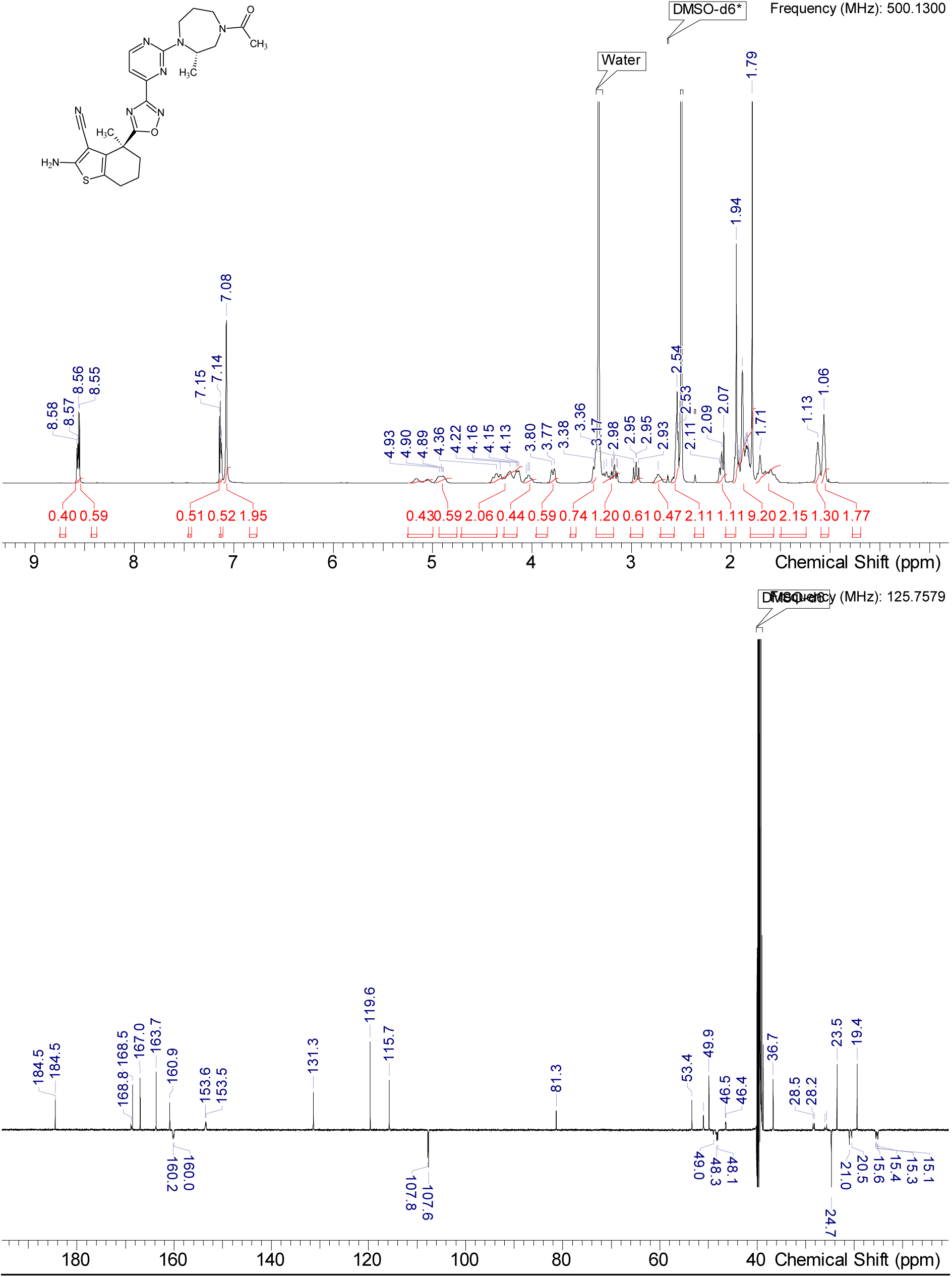

HRMS (m/z): [M+H]+ calcd. for C24H28N8O2S , 493.21287; found, 493.21304

## Synthesis of PROTAC compounds

### Synthesis of compound 2

#### Intermediate Int-6

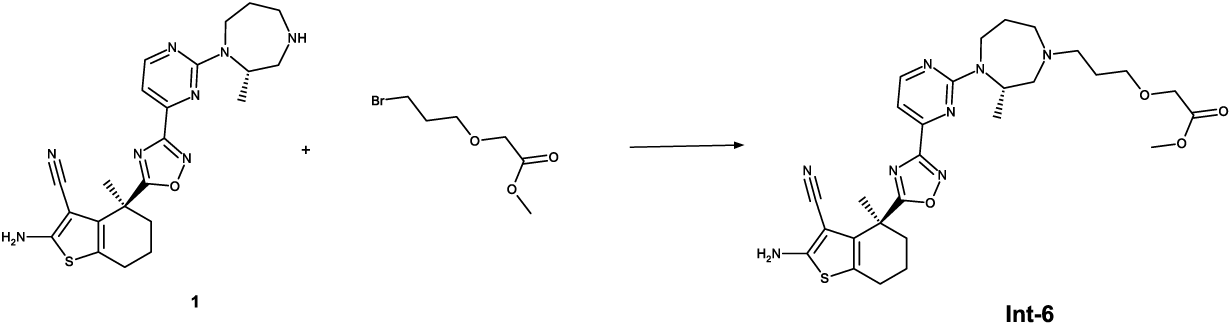

To a solution of 1 (450 mg, 0.899 mmol, 90%, 1.0 eq.) in acetonitrile (5 mL) was added 2-(3-bromopropoxy)acetate (399 mg, 1.798 mmol, 95%, 2.0 eq.) and DIPEA (408 mg, 3.146 mmol, 3.5 eq.). The mixture was heated to 50°C and stirred at this temperature for 16 h. The mixture is cooled to RT, filtered and purified by RP HPLC (Method 3b) and lyophilized to obtain Int-6 (338 mg, 0.582 mmol, 65% yield).

^1^H NMR (DMSO-d_6_) δ: 8.56 (d, J=4.6 Hz, 1H), 7.04-7.13 (m, 3H), 4.70 (dt, J=11.3, 5.9 Hz, 1H), 4.29 (br t, J=16.3 Hz, 1H), 4.02 (br d, J=7.1 Hz, 2H), 3.60 (br d, J=6.3 Hz, 3H), 3.41 (t, J=6.3 Hz, 2H), 3.13-3.28 (m, 1H), 3.00 (br d, J=14.2 Hz, 1H), 2.81 (br d, J=11.9 Hz, 1H), 2.52-2.59 (m, 3H), 2.39-2.48 (m, 2H), 2.03-2.16 (m, 1H), 1.75-2.00 (m, 6H), 1.49-1.75 (m, 4H), 1.02 (br s, 3H) 1H under DMSO

^13^C NMR (DMSO-d_6_) δ: 184.4, 170.7, 167.1, 163.7, 160.7, 160.1, 160.0, 153.5, 153.5, 131.3, 119.6, 115.7, 107.2, 81.4, 69.0, 67.4, 59.7, 59.3, 56.3, 52.6, 52.0, 51.2, 50.0, 36.7, 27.7, 27.5, 27.3, 24.7, 23.6, 19.4, 15.7, 15.6 1C under DMSO, 1 carbon only observed in HSQC

More carbon peaks detected than present in the structure due to presence of rotamers

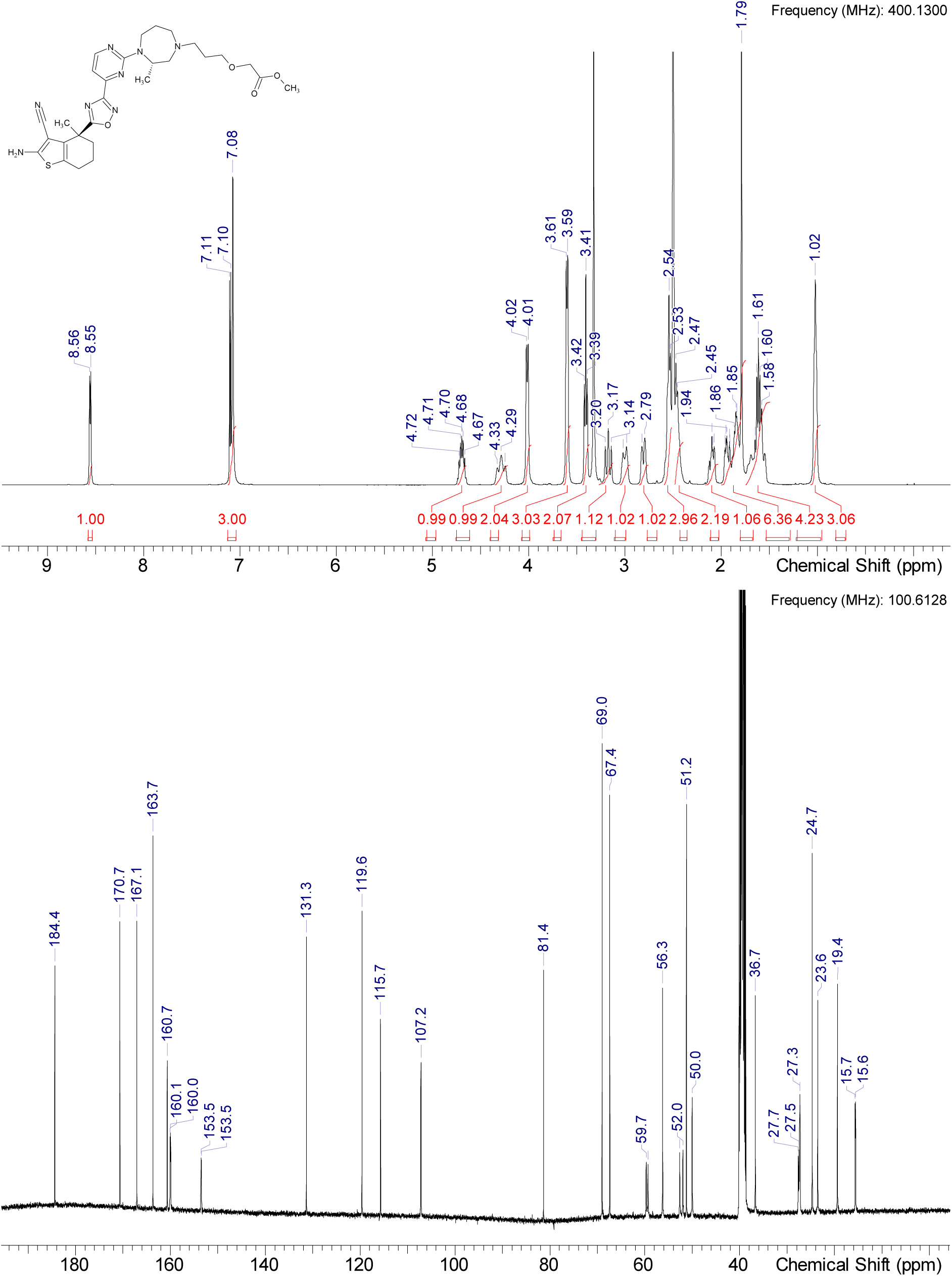

HRMS (m/z): [M+H]+ calcd. for C28H36N8O4S , 581.26530; found, 581.26593

#### Intermediate Int-7

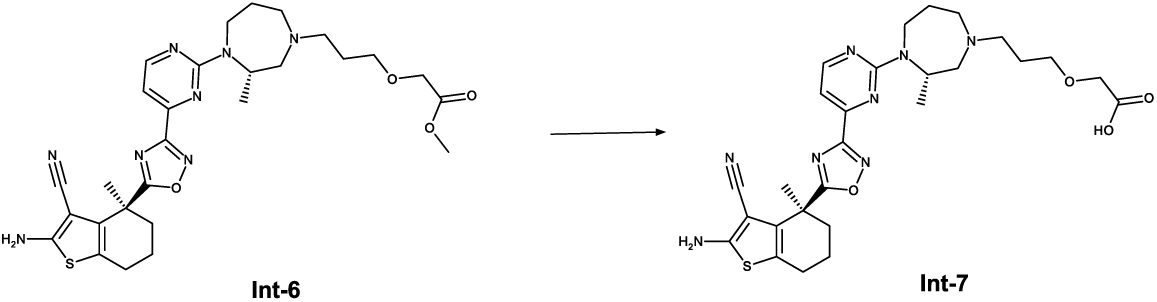

To a solution of Int-6 (300 mg, 0.517 mmol, 1.0 eq.) in MeOH (5 mL) is added aqueous NaOH solution (2 mL, 2 M) and the mixture is stirred at 50°C for 1 h. The mixture is cooled to RT and the volatiles are removed under reduced pressure. The residue is taken up in a mixture of acetonitrile and water, purified by RP HPLC (Method 3b) and lyophilized to obtain Int-7 (251 mg, 0.443 mmol, 86% yield).

^1^H NMR (DMSO-d_6_) δ: 8.55 (d, J=4.8 Hz, 1H), 7.10 (br s, 1H), 7.08 (s, 2H), 4.86 (br s, 1H), 4.34 (br s, 1H), 3.42-3.96 (m, 4H), 3.30 (br d, J=11.7 Hz, 3H), 3.01 (br d, J=11.2 Hz, 1H), 2.65-2.93 (m, 4H), 2.52-2.60 (m, 2H), 2.02-2.24 (m, 1H), 1.81-2.00 (m, 5H), 1.79 (s, 3H), 1.57-1.74 (m, 2H), 1.00-1.13 (m, 3H)

^13^C NMR (DMSO-d_6_) δ: 184.4, 172.5, 167.1, 163.7, 160.5, 160.0, 153.4, 131.3, 119.7, 115.8, 107.6, 81.4, 68.7, 57.3, 56.9, 56.5, 54.2, 53.3, 48.2, 38.7, 25.7, 25.6, 24.7, 23.6, 19.4, 1C under DMSO

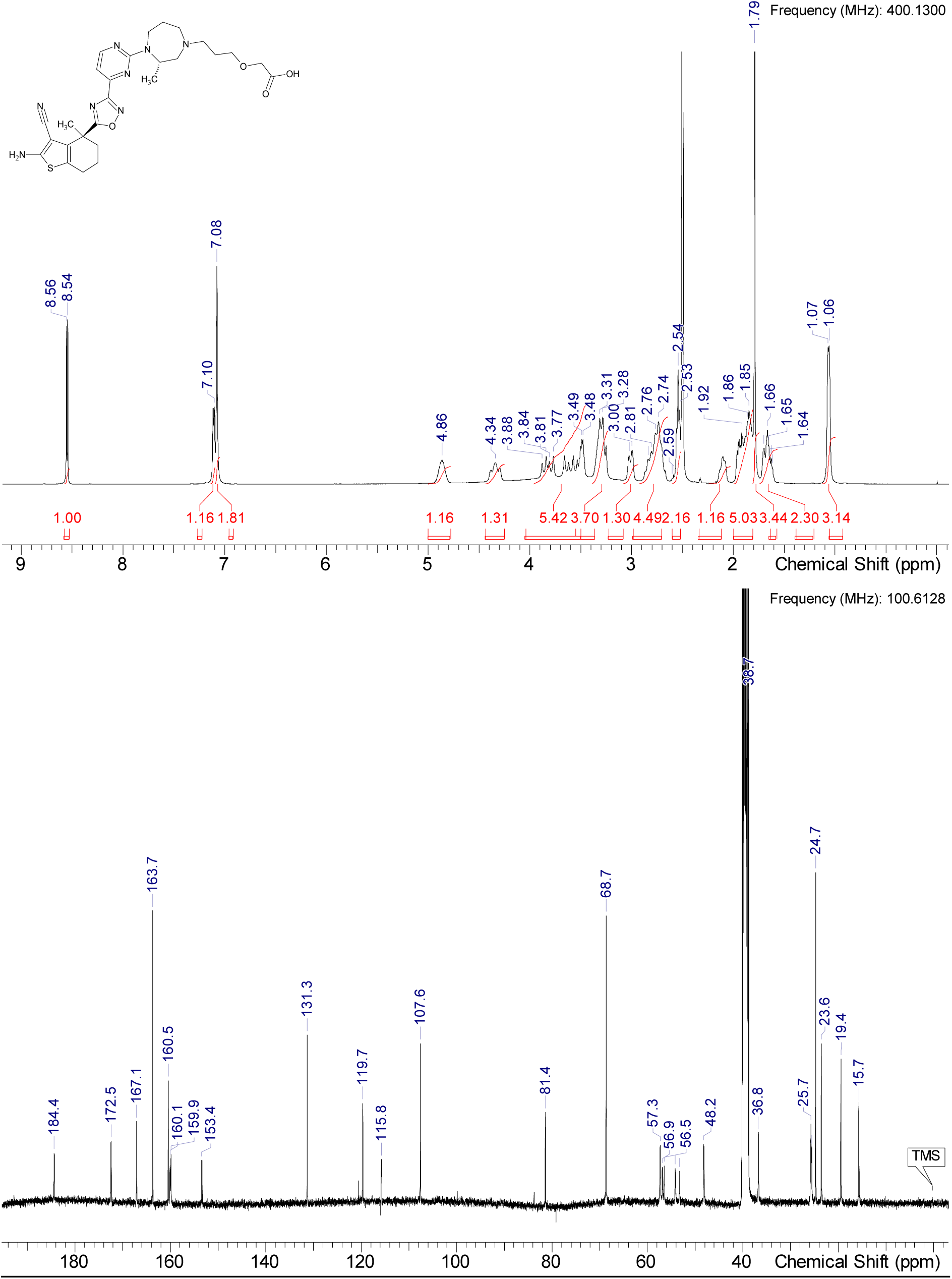

HRMS (m/z): [M+H]+ calcd. for C27H34N8O4S , 567.24965; found, 567.24951

#### Compound 2

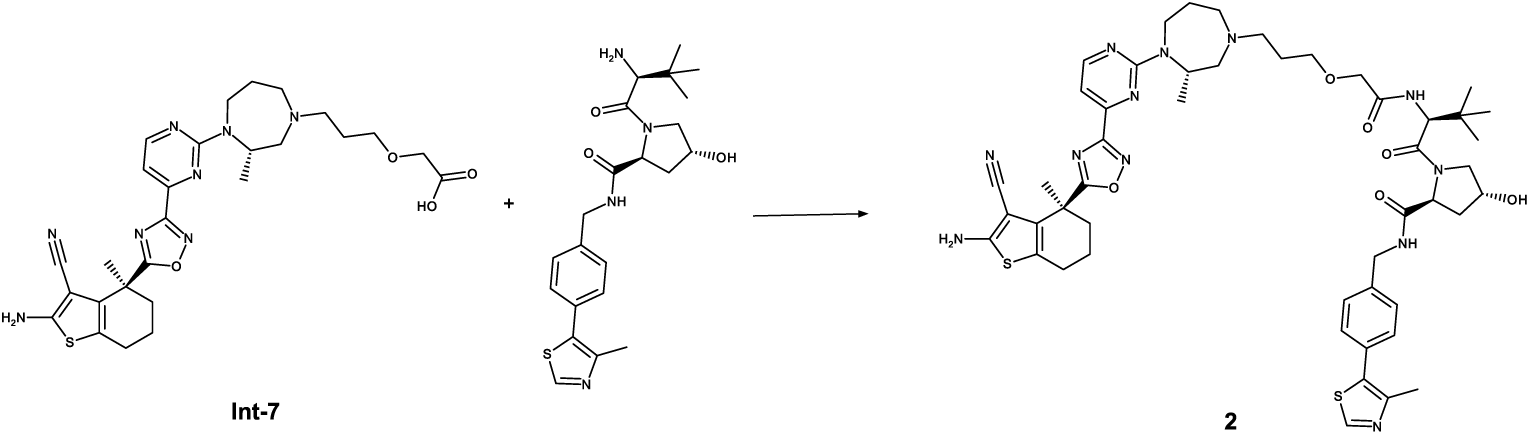

To a solution of Int-7 in DMF (3.0 mL) is added HATU (227 mg, 0.598 mmol, 1.4 eq.), (2S,4R)-1-[(2S)-2-amino-3,3-dimethylbutanoyl]-4-hydroxy-N-{[4-(4-methyl-1,3-thiazol-5-yl)phenyl]methyl}pyrrolidine -2-carboxamide hydrochloride (248 mg, 0.532 mmol, 1.2 eq.) and DIPEA (172 mg, 1.33 mmol, 3.0 eq.). The reaction mixture is stirred at room temperature for 2 h. The reaction mixture is concentrated under reduced pressure. The residue is purified by chromatography to give 2. The mixture is filtered, diluted with water and acetonitrile, purified by RP HPLC (Method 3b) and lyophilized to obtain 2 (347 mg, 0.354 mmol, 80% yield).

^1^H NMR (DMSO-d_6_) δ: 8.95-8.99 (m, 1H), 8.67 (br s, 0.14H rotamer), 8.60 (t, J=6.0 Hz, 1H), 8.52-8.56 (m, 1H), 7.39 (d, J=1.5 Hz, 4H), 7.32 (br d, J=7.9 Hz, 1H), 7.09 (d, J=4.8 Hz, 1H), 7.07 (s, 2H), 5.15 (d, J=3.3 Hz, 1H), 5.00 (d, J=2.3 Hz, 0.07H rotamer), 4.62-4.73 (m, 1H), 4.54 (br d, J=9.6 Hz, 1H), 4.15-4.48 (m, 6H), 3.86 (br s, 1.84H rotamer), 3.83 (br s, 0.18H rotamer), 3.56-3.70 (m, 2H), 3.44 (br t, J=6.2 Hz, 3H), 3.13 (br t, J=12.7 Hz, 1H), 3.00 (br d, J=10.1 Hz, 1H), 2.81 (br d, J=10.9 Hz, 1H), 2.52-2.60 (m, 4H), 2.45 (br s, 0.61H rotamer), 2.43 (s, 2.97H rotamer), 2.01-2.16 (m, 2H), 1.90 (br s, 4H), 1.78 (s, 3H), 1.66 (br s, 3H), 1.47-1.56 (m, 1H), 1.00 (br s, 3H), 0.88-0.97 (m, 9H) 1.77H rotamer)

^13^C NMR (DMSO-d6) δ: 184.4, 171.8, 169.1, 168.5, 167.1, 163.7, 160.7, 160.1, 160.0, 153.5, 153.5, 151.4, 147.7, 139.4, 131.3, 131.1, 129.7, 128.9, 128.7, 128.1, 127.5, 119.6, 115.7, 107.2, 81.4, 69.4, 69.1, 68.9, 59.8, 59.4, 58.7, 56.6, 56.3, 55.6, 52.9, 52.1, 50.0, 41.7, 37.9, 36.7, 35.8, 27.7, 27.5, 27.3, 27.2, 26.2, 26.1, 24.7, 23.6, 19.4, 15.9, 15.6, 15.6

More carbon peaks detected than present in the structure due to presence of rotamers

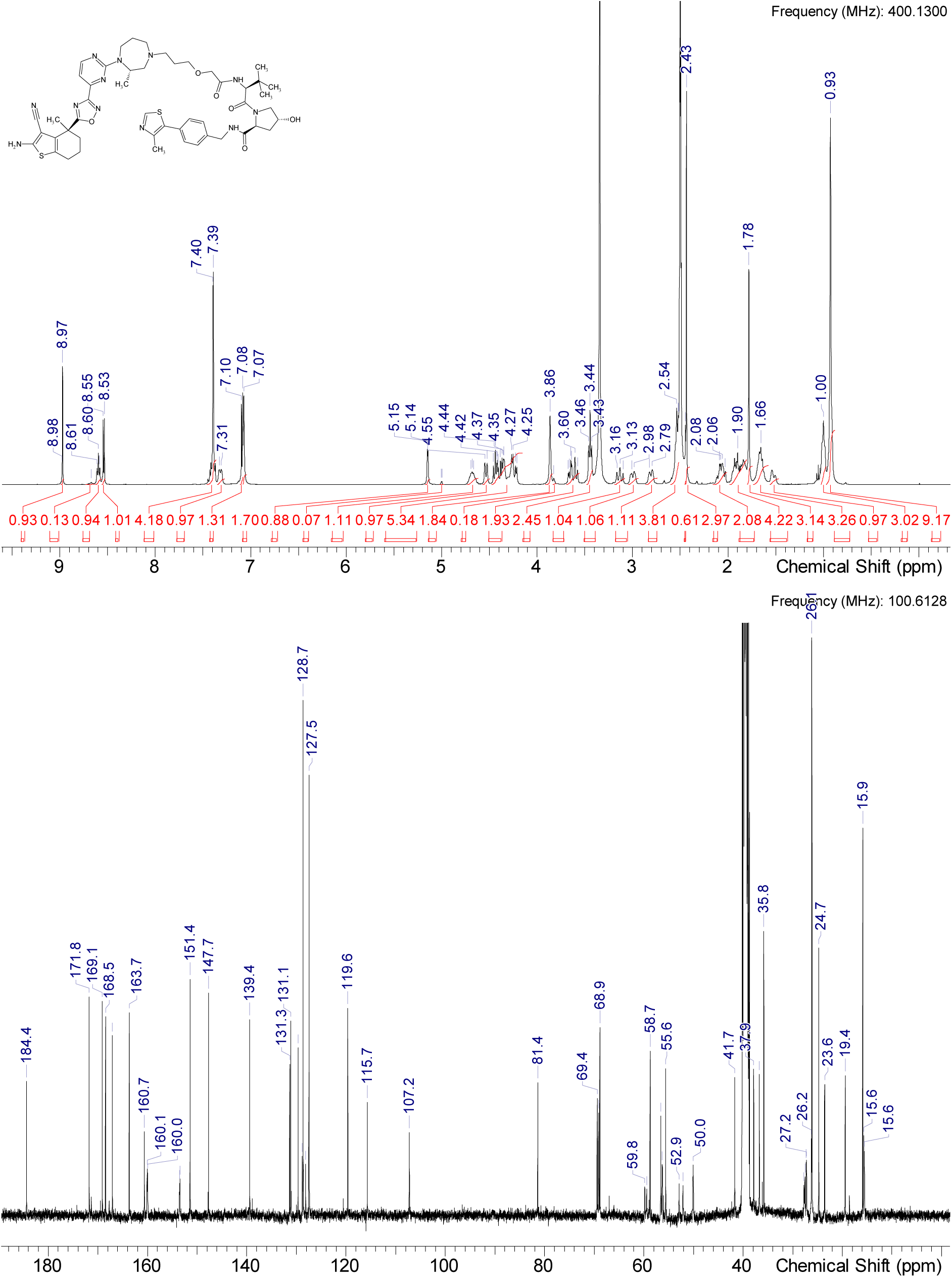

HRMS (m/z): [M+H]+ calcd. for C49H62N12O6S2 , 979.44295; found, 979.44348

### Synthesis of compound 3

#### Intermediate Int-8

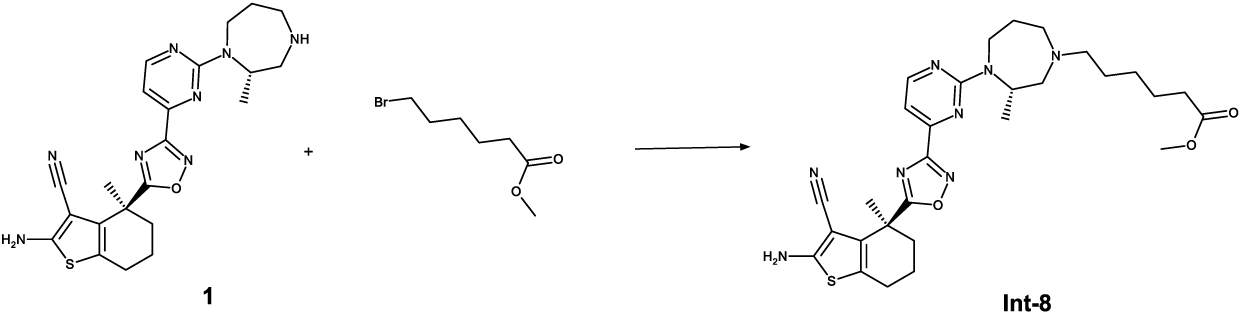

To a solution of 1 (450 mg, 0.899 mmol, 1.0 eq.) is added methyl 6-bromohexanoate (383 mg, 1.798 mmol, 2.0 eq.) and DIPEA (408 mg, 3.146 mmol, 3.5 eq.) and the mixture was stirred at 50 °C over night. The mixture is cooled to RT, filtered and purified by RP HPLC (Method 3b) and lyophilized to obtain Int-8 (332 mg, 0.574 mmol, 64% yield).

^1^H NMR (DMSO-d_6_) δ: 8.55 (d, J=4.8 Hz, 1H), 7.10 (d, J=4.8 Hz, 1H), 7.08 (s, 2H), 4.69 (dt, J=11.3, 5.9 Hz, 1H), 4.28 (br t, J=16.7 Hz, 1H), 3.54 (br d, J=5.6 Hz, 3H), 3.16 (br t, J=12.7 Hz, 1H), 2.99 (br d, J=13.9 Hz, 1H), 2.80 (br d, J=11.7 Hz, 1H), 2.52-2.59 (m, 3H), 2.01-2.46 (m, 6H), 1.80-2.01 (m, 3H), 1.79 (s, 3H), 1.32-1.74 (m, 6H), 1.19 (br d, J=6.8 Hz, 2H), 1.02 (br s, 3H)

^13^C NMR (DMSO-d_6_) δ: 184.4, 173.3, 167.1, 163.7, 160.7, 160.0, 159.9, 153.5, 153.5, 131.3, 119.6, 115.7, 107.2, 81.4, 59.7, 59.3, 56.3, 55.7, 54.9, 51.1, 50.1, 50.0, 38.7, 36.7, 33.2, 27.7, 27.5, 26.7, 26.6, 26.2, 24.7, 24.3, 23.6, 19.4, 15.7, 15.6

More carbon peaks detected than present in the structure due to presence of rotamers

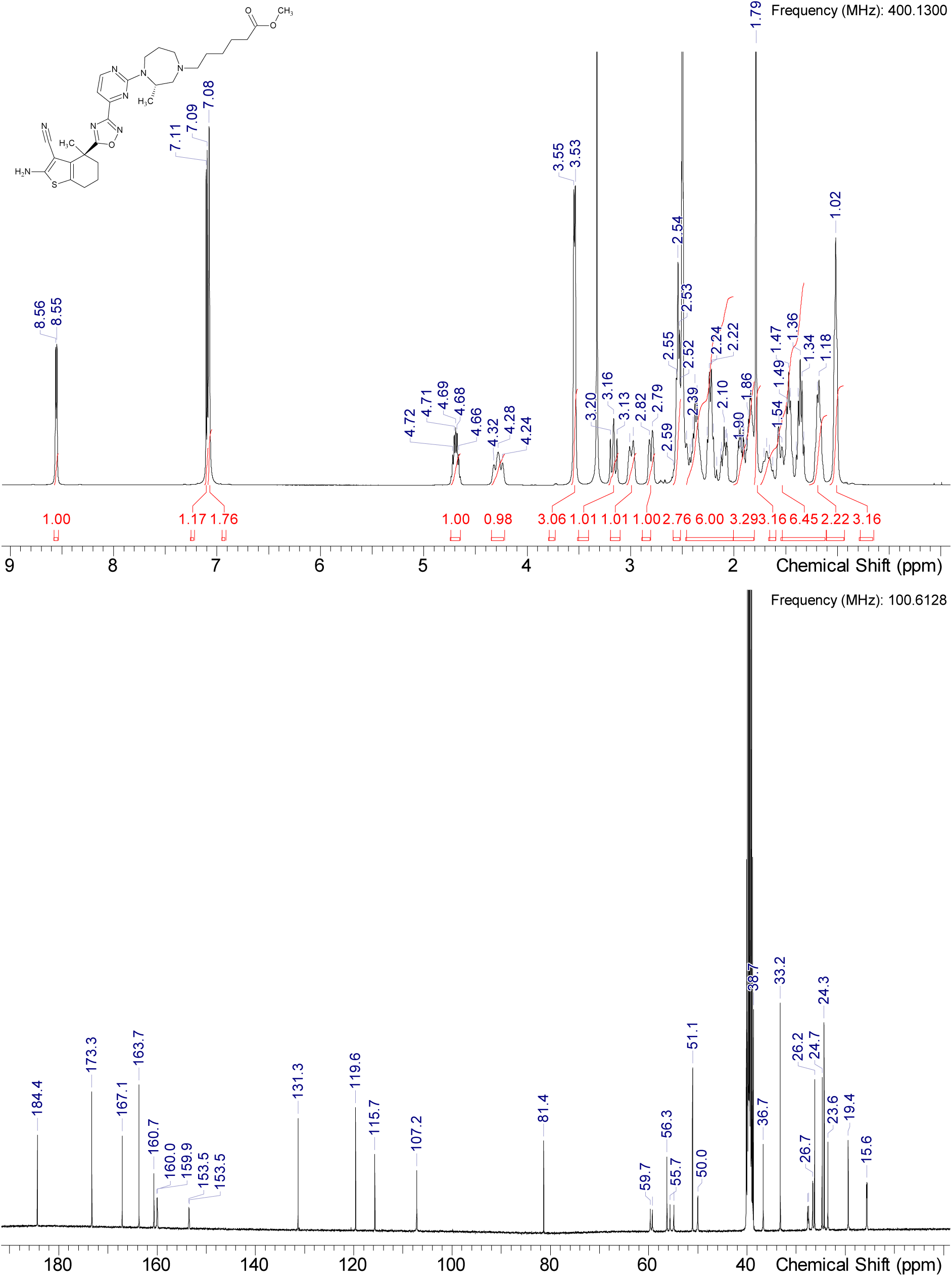

HRMS (m/z): [M+H]+ calcd. for C29H38N8O3S , 579.28603; found, 579.28595

#### Compound 3

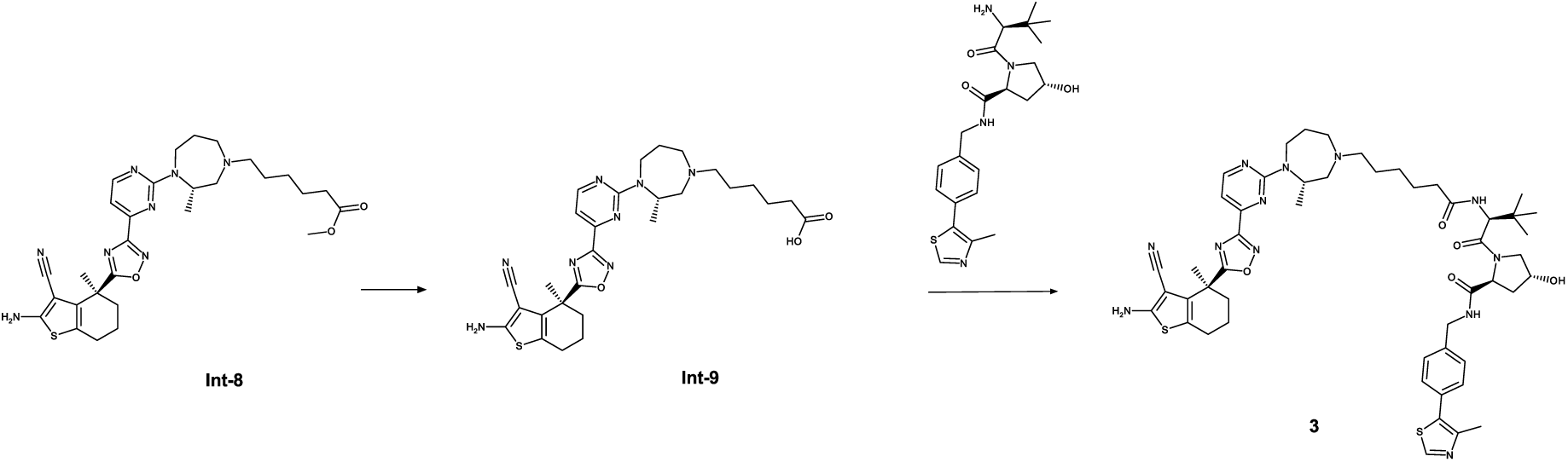

To a solution of Int-8 (43 mg, 0.073 mmol, 1.0 eq.) in THF (2.0 mL) and water (2 mL) is added LiOH H_2_O (15 mg, 0.36 mmol) and the reaction mixture is stirred at room temperature overnight. The reaction mixture is neutralized using 4M HCl in water, and concentrated under reduced pressure. The crude acid intermediate Int-9 is taken up in DMF (2.0 mL). HATU (50 mg, 0.13 mmol, 1.8 eq.), HOAt (18 mg, 0.13 mmol, 1.8 qe.), (2S,4R)-1-[(2S)-2-amino-3,3-dimethylbutanoyl]-4-hydroxy-N-{[4-(4-methyl-1,3-thiazol-5-yl)phenyl]methyl}pyrrolidine -2-carboxamide hydrochloride (43 mg, 0.073 mmol, 1.8 eq.) and DIPEA (35 μL, 0.21 mmol, 2.9 eq.) are added. The reaction mixture is stirred at room temperature overnight. The reaction mixture is concentrated under reduced pressure. The residue is purified by RP chromatography (Method 3b) to give 3.

^1^H NMR (DMSO-d_6_) δ: 8.56 (br d, J=4.1 Hz, 1H), 7.10 (d, J=4.8 Hz, 1H), 7.08 (s, 2H), 4.70 (dt, J=11.3, 5.9 Hz, 1H), 4.29 (br t, J=16.7 Hz, 1H), 3.11-3.23 (m, 2H), 2.95-3.06 (m, 1H), 2.81 (br d, J=11.7 Hz, 1H), 2.54 (br d, J=6.1 Hz, 3H), 2.38 (br d, J=6.8 Hz, 3H), 2.04-2.23 (m, 3H), 1.85 (br d, J=4.1 Hz, 3H), 1.79 (s, 3H), 1.41 (dt, J=33.8, 7.1 Hz, 6H), 1.13-1.26 (m, 2H), 1.02 (br s, 3H)

^13^C NMR (DMSO-d_6_) δ: 184.4, 174.6, 167.1, 163.7, 160.7, 160.1, 160.0, 153.5, 153.5, 131.3, 119.6, 115.7, 107.2, 81.3, 59.7, 59.2, 56.3, 55.8, 54.9, 50.1, 50.0, 38.7, 36.7, 33.8, 27.7, 27.5, 26.7, 26.6, 26.4, 24.7, 24.4, 23.5, 19.4, 15.7, 15.6, 1C under DMSO

More carbon peaks detected than present in the structure due to presence of rotamers

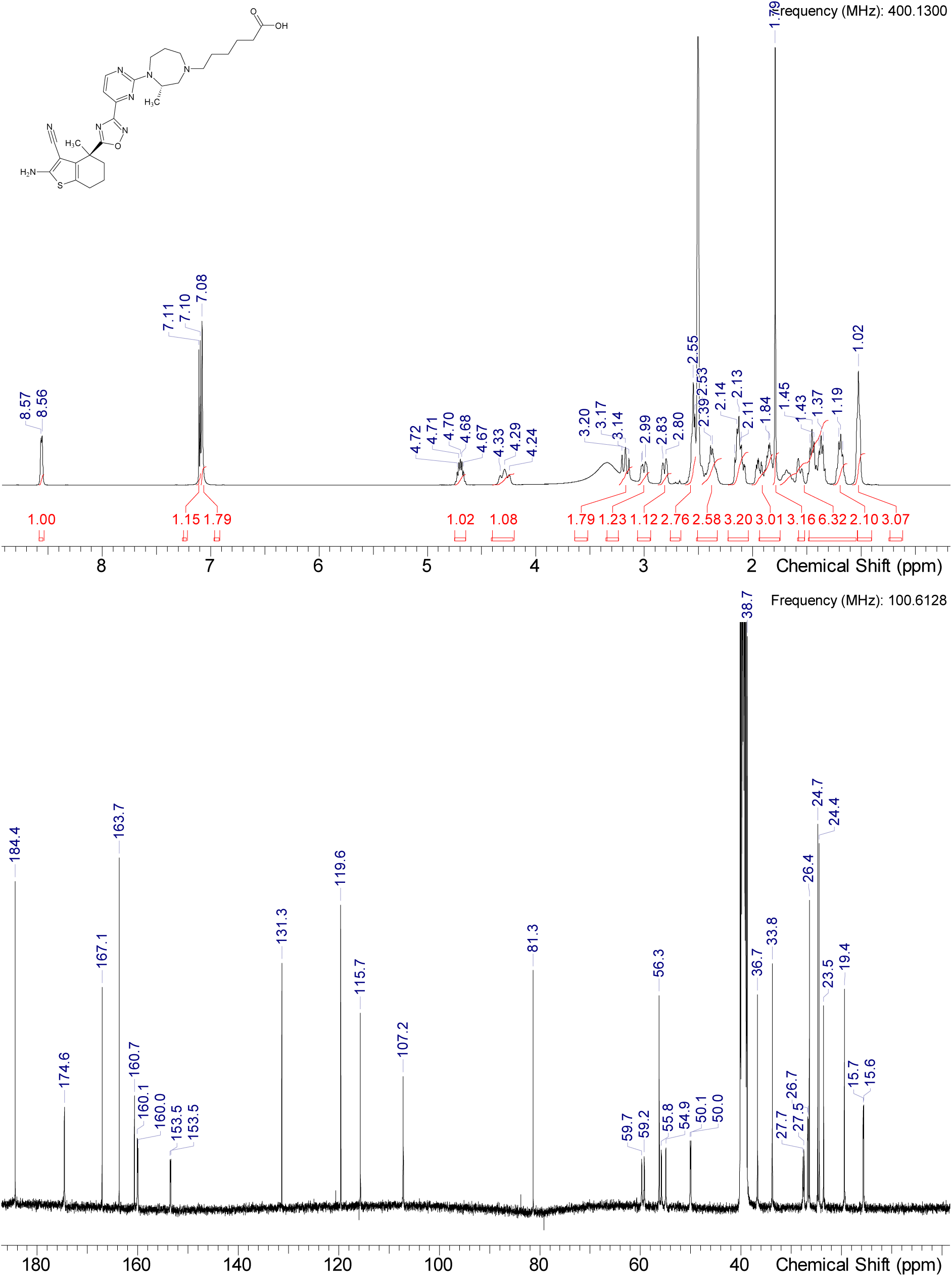

HRMS (m/z): [M+H]+ calcd. for C28H36N8O3S , 565.27038; found, 565.27045

^1^H NMR (DMSO-d_6_) δ: 8.98 (s, 1H), 8.50-8.61 (m, 2H), 7.80 (br t, J=7.2 Hz, 1H), 7.40-7.43 (m, 2H), 7.36-7.39 (m, 2H), 7.10 (d, J=4.8 Hz, 1H), 7.08 (s, 2H), 5.11 (d, J=3.3 Hz, 1H), 4.62-4.74 (m, 1H), 4.52 (br d, J=9.4 Hz, 1H), 4.38-4.48 (m, 2H), 4.34 (br s, 1H), 4.17-4.31 (m, 2H), 3.58-3.70 (m, 2H), 3.16 (br t, J=12.5 Hz, 1H), 2.99 (dd, J=14.2, 5.6 Hz, 1H), 2.81 (br d, J=11.7 Hz, 1H), 2.52-2.60 (m, 3H), 2.44 (s, 3H), 2.30-2.41 (m, 2H), 2.22 (br s, 1H), 1.97-2.16 (range, 3H), 1.79-1.96 (range, 4H), 1.78 (s, 3H), 1.61-1.72 (m, 1H), 1.52-1.60 (range, 1H), 1.30-1.52 (range, 4H), 1.10-1.24 (m, 2H), 1.02 (br t, J=7.0 Hz, 3H), 0.91 (s, 9H), 1H under DMSO

^13^C NMR (DMSO-d_6_) δ: 184.4, 172.1, 171.9, 169.7, 167.1, 163.7, 160.7, 160.1, 160.0, 153.5, 153.5, 151.4, 147.7, 139.5, 131.3, 131.1, 129.6, 128.6, 128.0, 127.4, 119.6, 115.7, 107.2, 81.3, 68.8, 59.9, 59.4, 58.7, 56.3, 56.3, 56.1, 55.2, 50.2, 50.0, 41.6, 37.9, 36.7, 35.2, 34.8, 27.7, 27.5, 26.7, 26.6, 26.4, 26.3, 25.3, 24.7, 23.6, 19.4, 15.9, 15.7, 15.6;

More carbon peaks detected than present in the structure due to presence of rotamers, 1C under DMSO

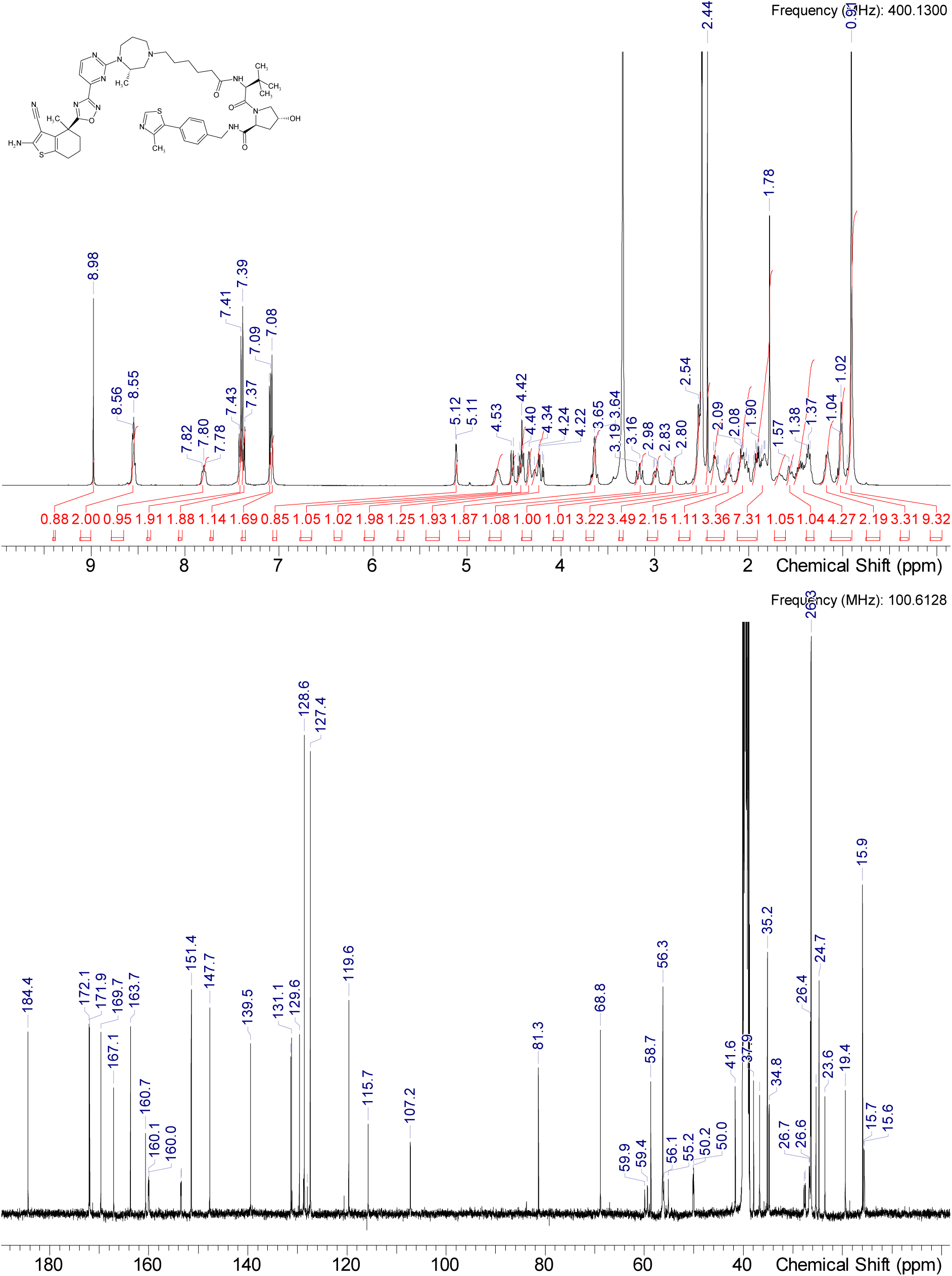

HRMS (m/z): [M+H]+ calcd. for C50H64N12O5S2 , 977.46368; found, 977.46423

### Synthesis of compound 4 and 6

#### Intermediate Int-10

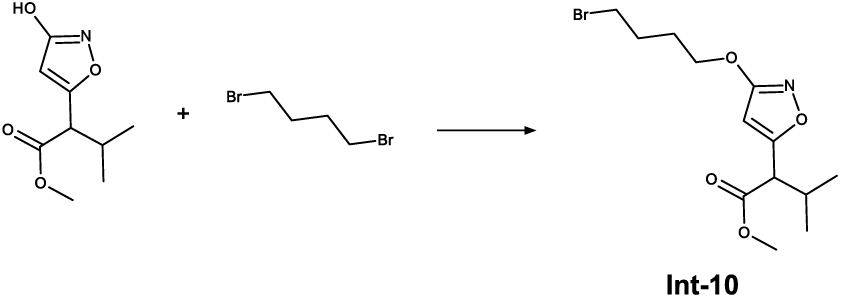

To a stirred solution of methyl 2-(3-hydroxy-1,2-oxazol-5-yl)-3-methylbutanoate (11.00 g, 0.055 mol, 1.0 eq.) in DMF (75.0 mL) is added potassium carbonate (22.86 g, 0.166 mol, 3.0 eq.) at 0 °C. 1,3-Dibromobutane (11.92 g, 0.055 mol, 1.0 eq.) is added dropwise and the reaction mixture is stirred at 0 °C for 9 hours. After complete conversion the reaction mixture is quenched with water and extracted with EtOAc. The organic layer is washed with ice water, dried over sodium sulfate and concentrated under reduced pressure to give the crude product. The obtained crude compound is purified by column chromatography (hexane/EtOAc, 0-10%) to yield Int-10 (5.01 g, 0.015 mmol, 27% yield) as a racemic mixture.

^1^H NMR (DMSO-d_6_) δ: 6.19 (s, 1H), 4.18 (t, J=6.3 Hz, 2H), 3.68 (d, J=8.8 Hz, 1H), 3.65-3.66 (m, 3H), 3.58 (t, J=6.6 Hz, 2H), 2.28 (s, 1H), 1.87-1.96 (m, 2H), 1.79-1.87 (m, 2H), 0.92 (d, J=6.6 Hz, 3H), 0.84 (d, J=6.6 Hz, 3H)

^13^C NMR (DMSO-d_6_) δ: 171.3, 169.9, 169.7, 94.3, 68.8, 52.3, 51.0, 34.7, 29.9, 28.8, 27.2, 20.3, 19.6

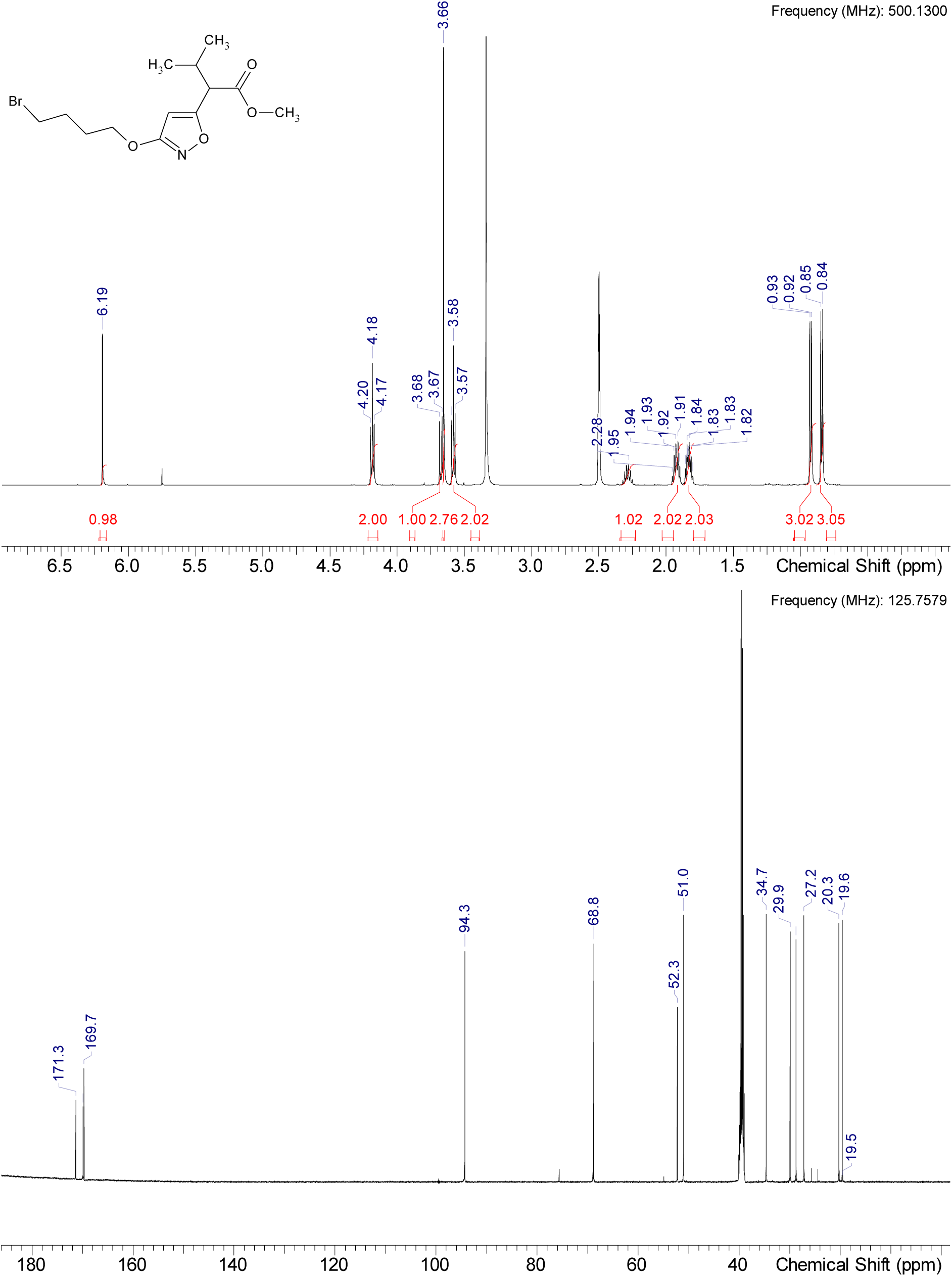

HRMS (m/z): [M+H]+ calcd. for C13H20BrNO4 , 334.06485; found, 334.06482

#### Intermediate Int-11

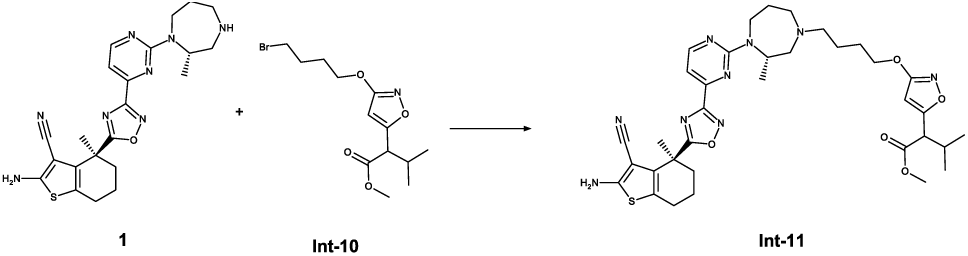

To a stirred solution of 1 (1.50 g, 3.22 mmol, 1.0 eq.) and Int-10 (1.16 g, 3.48 mmol, 1.1 eq) in acetonitrile (15.0 mL) is added potassium carbonate (0.89 g, 6.45 mmol, 2.0 eq) and the mixture is stirred at 60 °C under argon for 22 h. The reaction mixture is allowed to cool to rt, filtered and the solid is washed with acetonitrile. The combined solution is concentrated under reduced pressure and purified by normal phase chromatography (DCM / MeOH, 0-10%) to give Int-11 (1.73, 2.46 mmol, 76 % yield) as a mixture of two diastereomers.

^1^H NMR (DMSO-d_6_) δ: 8.56 (d, J=5.0 Hz, 1H), 7.03-7.14 (m, 3H), 6.16 (br d, J=4.4 Hz, 1H), 4.71 (dt, J=11.3, 5.9 Hz, 1H), 4.22-4.39 (m, 1H), 4.12 (br s, 2H), 3.66 (s, 4H), 3.19 (br t, J=12.8 Hz, 1H), 2.95-3.09 (m, 1H), 2.83 (br d, J=11.7 Hz, 1H), 2.52-2.62 (m, 3H), 2.38 (br s, 3H), 2.28 (br d, J=8.2 Hz, 1H), 2.06-2.16 (m, 1H), 1.82-2.00 (m, 3H), 1.79 (s, 3H), 1.42-1.76 (m, 6H), 1.03 (br t, J=6.9 Hz, 3H), 0.93 (d, J=6.6 Hz, 1H), 0.84 (d, J=6.6 Hz, 3H), 2 H under DMSO

^13^C NMR (DMSO-d_6_) δ: 184.4, 171.4, 169.9, 169.6, 167.1, 163.7, 160.7, 160.1, 160.0, 153.5, 153.5, 131.3, 119.6, 115.7, 107.2, 94.3, 81.3, 69.6, 59.7, 59.2, 56.2, 55.3, 54.5, 52.2, 51.0, 50.0, 38.7, 36.7, 29.9, 27.7, 27.5, 26.3, 24.7, 23.5, 23.3, 23.2, 20.3, 19.6, 19.4, 15.7, 15.6

More carbon peaks detected than present in the structure due to presence of rotamers, 1C under DMSO

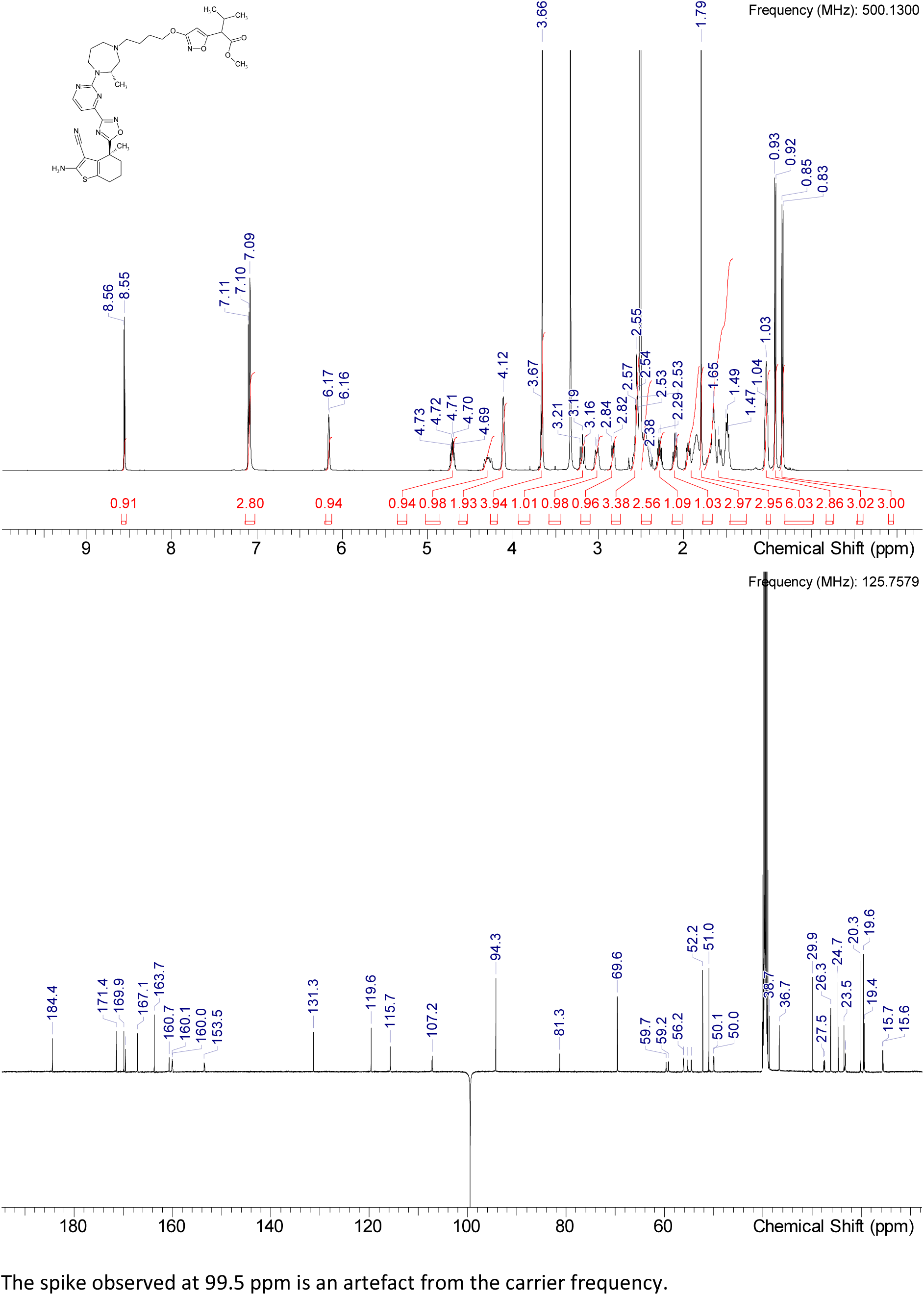

HRMS (m/z): [M+H]+ calcd. for C35H45N9O5S , 704.33371; found, 704.33331

#### Intermediate Int-12

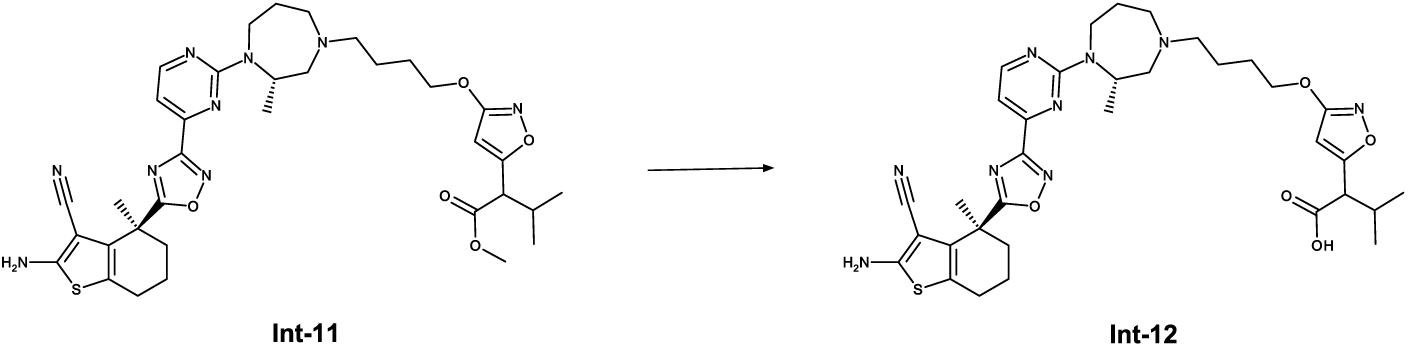

To a stirred solution of Int-11 (1.30 g, 1.85 mmol, 1.0 eq.) in methanol (13.0 ml) is added sodium hydroxide solution (2 M in water, 1.85 ml, 3.70 mmol, 2.0 eq) and the reaction mixture is stirred at 45°C for 1 h. After complete conversion the reaction mixture is concentrated under reduced pressure. The crude product is purified by RP chromatography (Method 3a) yielding Int-12 (1.10 g, 1.60 mmol, 86%) as a mixture of two diastereomers.

^1^H NMR (DMSO-d_6_) δ: 11.47 (br s, 0.31H rotamer), 9.93 (br d, J=2.8 Hz, 0.25H rotamer), 9.72 (br s, 0.39H rotamer), 8.64 (br d, J=4.7 Hz, 1H), 7.19-7.30 (m, 1.04H rotamer), 6.14 (s, 0.66H rotamer), 5.98-6.11 (m, 0.3H rotamer), 4.97 (br s, 0.29H rotamer), 4.89 (br d, J=5.7 Hz, 0.75H rotamer), 4.30-4.51 (range, 1.32H rotamer), 4.19 (br t, J=6.0 Hz, 2H), 4.08 (br d, J=3.2 Hz, 1H), 3.51 (br d, J=8.8 Hz, 6H), 3.17-3.31 (m, 3H), 2.79-2.94 (m, 0.68H rotamer), 2.67-2.68 (m, 1H), 2.52-2.57 (m, 2H), 2.19-2.28 (m, 1H), 2.18 (s, 1H), 2.01-2.14 (m, 1H), 1.80-2.01 (range, 7H), 1.78 (s, 3H), 1.74 (br s, 1H), 1.58-1.69 (m, 1H), 1.13 (br s, 3H rotamer), 0.95 (br d, J=6.6 Hz, 1H), 0.78-0.87 (m, 3H)

^13^C NMR (DMSO-d_6_) δ: 184.6, 171.3, 170.8, 170.6, 167.0, 163.8, 160.5, 160.1, 153.8, 153.6, 131.3, 131.3, 119.7, 119.6, 115.8, 115.7, 108.4, 94.0, 69.0, 57.8, 55.4, 53.1, 51.6, 47.6, 38.8, 38.2, 36.8, 36.7, 29.7, 25.7, 24.7, 23.6, 20.5, 19.6, 19.5, 19.4, 15.5, 15.4, 15.2

More carbon peaks detected than present in the structure due to presence of rotamers, 1C under DMSO

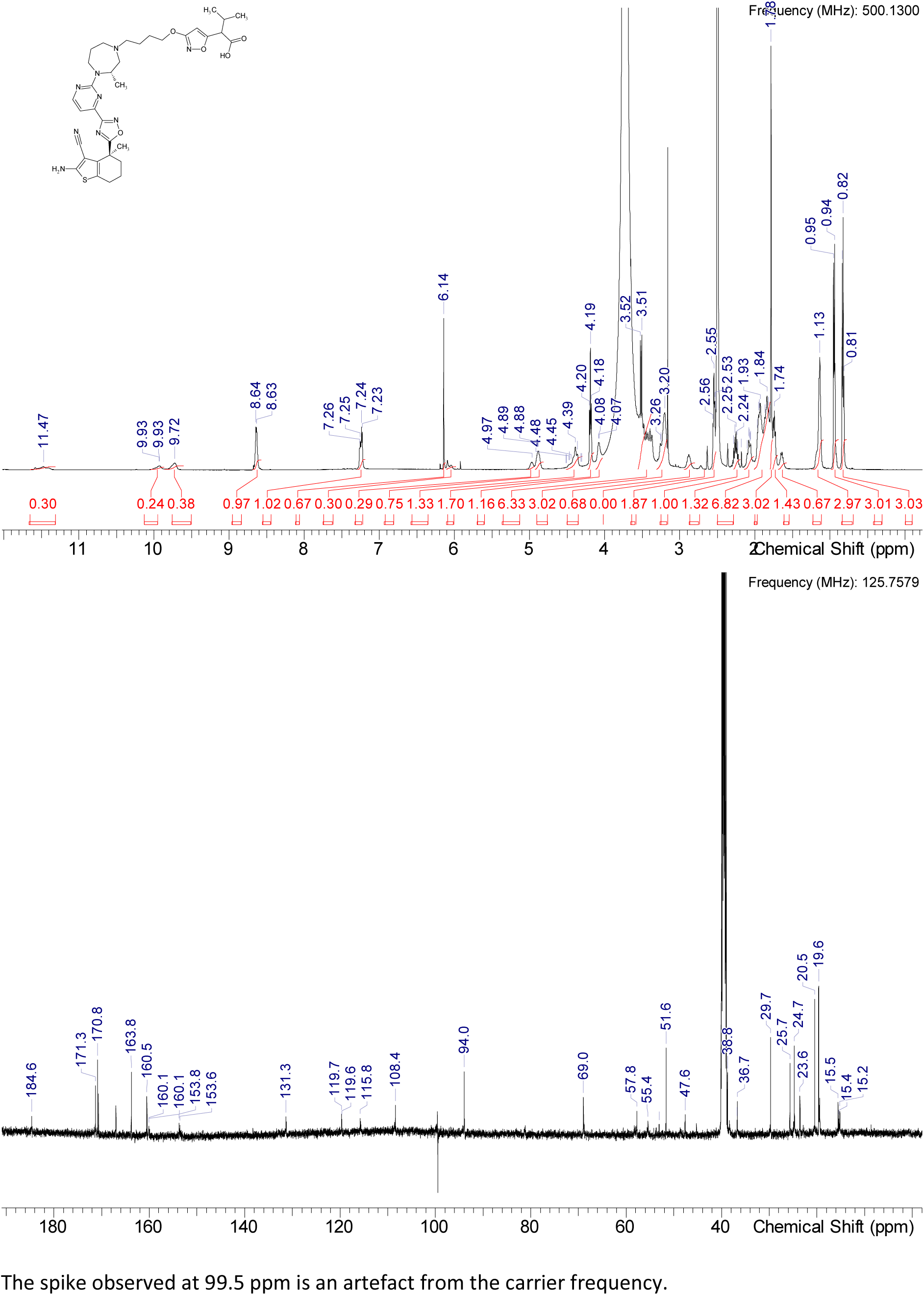

HRMS (m/z): [M+H]+ calcd. for C34H43N9O5S , 690.31806; found, 690.31806

#### Compound 4 and 6

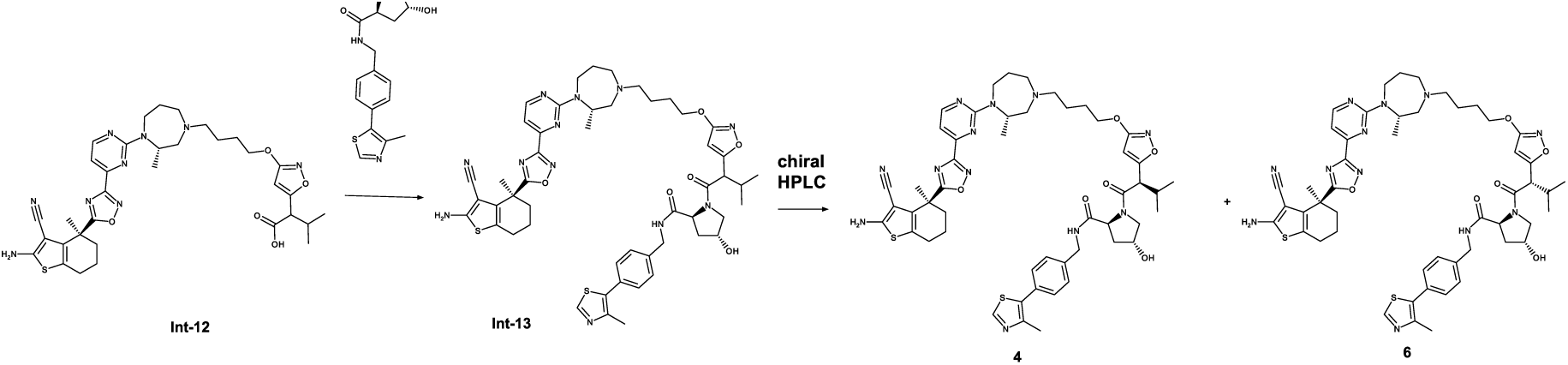

To a stirred solution of Int-12 (490 mg, 0.71 mmol, 1.0 eq.), (2S,4R)-4-hydroxy-N-{[4-(4-methyl-1,3-thiazol-5-yl)phenyl]methyl}pyrrolidine-2-carboxamide (326 mg, 0.923 mmol, 1.1 eq.) and HATU (330 mg, 0.85 mmol, 1.2 eq.) in DMSO (4.9 ml) is added DIPEA (1.24 mL, 7.10 mmol, 10.0 eq) and the reaction mixture is stirred at rt for 15 min. After complete conversion the reaction mixture is quenched with water, diluted with acetonitrile and purified by RP chromatography (Method 3b) to obtain Int-13 (426 mg, 0.431, 61 %) as a mixture of two diastereomers. The mixture Int-13 (330 mg, 0.334 mmol) is separated by preparative chiral HPLC (Method A) to obtain 4 (136 mg, 0.137 mmol) and less active diastereomer 6 (146 mg, 0.148 mmol).

^1^H NMR (DMSO-d_6_) δ: 8.98 (s, 0.5H rotamer), 8.95-8.97 (m, 0.5H rotamer), 8.82 (s, 0.1H rotamer), 8.54 (s, 1.5H rotamer), 8.40-8.48 (m, 0.5H rotamer), 7.35-7.48 (m, 3H), 7.31 (s, 1H), 7.03-7.13 (m, 3H), 6.04 (br dd, J=15.9, 4.6 Hz, 1H), 5.65 (br d, J=4.4 Hz, 0.1H rotamer), 5.12 (dd, J=10.7, 3.5 Hz, 1H), 5.04 (d, J=2.8 Hz, 0.1H rotamer), 4.97 (d, J=3.2 Hz, 0.1H rotamer), 4.61-4.79 (range, 1H), 4.42 (br t, J=7.7 Hz, 0.5H rotamer), 4.17-4.39 (m, 4.5H rotamer), 3.99-4.14 (m, 2H), 3.71-3.83 (m, 0.9H rotamer), 3.65 (br d, J=9.5 Hz, 0.4H rotamer), 3.51-3.62 (m, J=5.0 Hz, 1H), 3.39-3.50 (m, 1H), 3.09-3.24 (m, 1H), 2.99 (br d, J=3.8 Hz, 1H), 2.80 (br s, 1H), 2.52-2.60 (range, 3H), 2.45 (s, 1H rotamer), 2.44 (s, 1.5H rotamer), 2.43 (br s, 0.8H rotamer), 2.41 (s, 0.7H rotamer), 2.39 (br d, J=7.9 Hz, 1H rotamer), 2.35-2.38 (m, 0.54H rotamer), 2.18-2.29 (m, 1H), 1.98-2.14 (m, 2H), 1.80-1.97 (m, 4H), 1.78 (s, 3H), 1.52-1.73 (m, 4H), 1.35-1.51 (m, 2H), 0.97-1.05 (m, 3H), 0.96 (d, J=6.6 Hz, 1.5H rotamer), 0.94 (br d, J=6.6 Hz, 1.5H rotamer), 0.82 (d, J=6.9 Hz, 1.5H rotamer), 0.78 (br d, J=6.6 Hz, 1.5H rotamer), 0.65 (d, J=6.6 Hz, 0.5H rotamer), 0.56 (d, J=6.6 Hz, 0.5H rotamer)

^13^C NMR (DMSO-d_6_) δ: 184.4, 171.6, 171.6, 171.4, 171.3, 171.3, 167.7, 167.4, 167.1, 163.7, 160.7, 160.1, 160.0, 151.5, 151.4, 147.8, 147.8, 139.4, 139.3, 131.4, 131.1, 129.8, 129.7, 129.0, 128.9, 128.8, 128.7, 128.1, 127.9, 127.5, 127.4, 119.6, 115.7, 107.2, 93.6, 93.5, 93.2, 81.3, 69.5, 69.4, 68.7, 68.6, 59.7, 58.9, 56.2, 55.8, 55.4, 55.3, 54.5, 50.0, 49.3, 41.6, 41.5, 38.0, 37.9, 36.7, 31.1, 30.8, 27.7, 27.5, 26.3, 24.7, 23.6, 23.2, 20.7, 20.5, 19.9, 19.6, 19.4, 15.9, 15.7, 15.6

More carbon peaks detected than present in the structure due to presence of rotamers

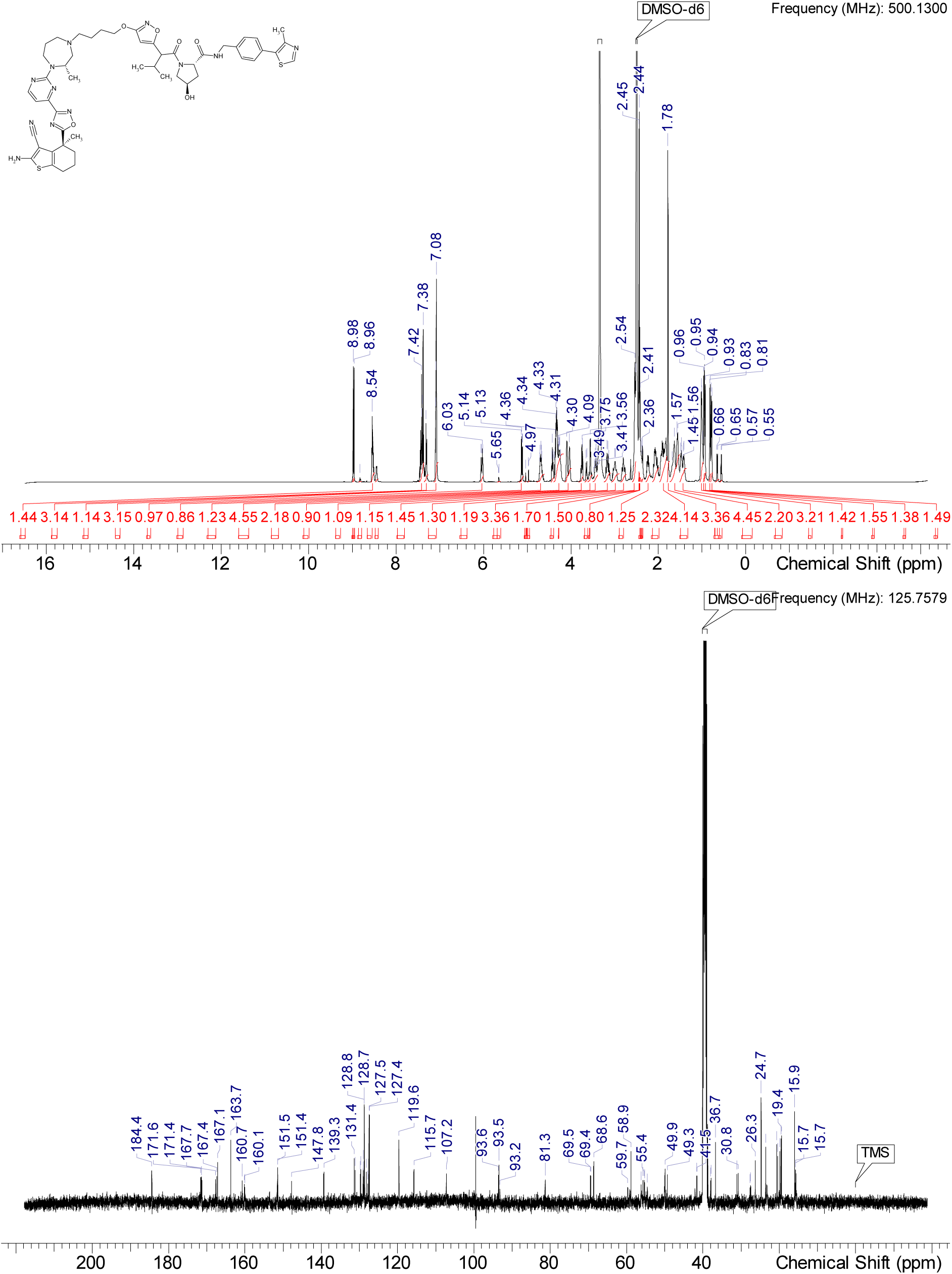

HRMS (m/z): [M+H]+ calcd. for C50H60N12O6S2 , 989.42730; found, 989.4267

^1^H NMR (DMSO-d_6_) δ: 8.95-9.01 (m, 1H), 8.52-8.56 (m, 1H), 8.44 (br d, J=5.0 Hz, 0.8H rotamer), 7.45 (s, 0.4H rotamer), 7.37-7.43 (range, 2H rotamer), 7.31 (d, J=7.9 Hz, 1.6H rotamer), 7.02-7.14 (m, 3H), 6.02 (br d, J=5.4 Hz, 1H), 4.68 (br s, 2H), 4.43 (br t, J=7.7 Hz, 1H), 4.20-4.37 (m, 4H), 4.09 (br s, 0.4H rotamer), 4.04 (br d, J=3.5 Hz, 1.6H rotamer), 3.74 (d, J=8.5 Hz, 0.8H rotamer), 3.55 (br d, J=4.7 Hz, 2H), 3.09-3.21 (m, 1H), 2.92-3.06 (m, 1H), 2.74-2.86 (m, 1H), 2.52-2.59 (m, 3H), 2.42-2.47 (m, 3H rotamer), 2.32-2.42 (m, 2H), 1.97-2.29 (range, 4H), 1.81-1.96 (range, 4H), 1.78 (s, 3H), 1.36-1.74 (range, 6H), 0.98-1.05 (m, 3H), 0.96 (d, J=6.6 Hz, 2.4H rotamer), 0.82 (d, J=6.6 Hz, 2.4H rotamer), 0.63-0.69 (m, 0.6H rotamer), 0.52-0.59 (m, 0.6H rotamer)

^13^C NMR (DMSO-d_6_) δ: 184.4, 171.8, 171.5, 171.3, 171.2, 170.6, 167.6, 167.3, 167.1, 163.7, 160.6, 160.1, 159.9, 153.5, 153.4, 151.6, 151.4, 147.8, 147.7, 139.3, 139.0, 131.3, 131.1, 131.0, 130.2, 129.7, 129.2, 129.0, 128.7, 128.2, 128.1, 127.3, 119.6, 115.7, 107.2, 93.6, 93.5, 81.3, 69.5, 69.4, 68.7, 67.0, 59.7, 59.2, 58.8, 58.6, 56.2, 56.0, 55.4, 55.3, 54.8, 54.6, 50.1, 50.0, 49.6, 49.3, 42.0, 41.5, 38.7, 37.9, 36.7, 30.8, 29.8, 27.7, 27.5, 26.3, 24.7, 23.5, 23.3, 23.1, 20.7, 20.7, 19.5, 19.5, 19.4, 15.9, 15.8, 15.7, 15.6

More carbon peaks detected than present in the structure due to presence of rotamers

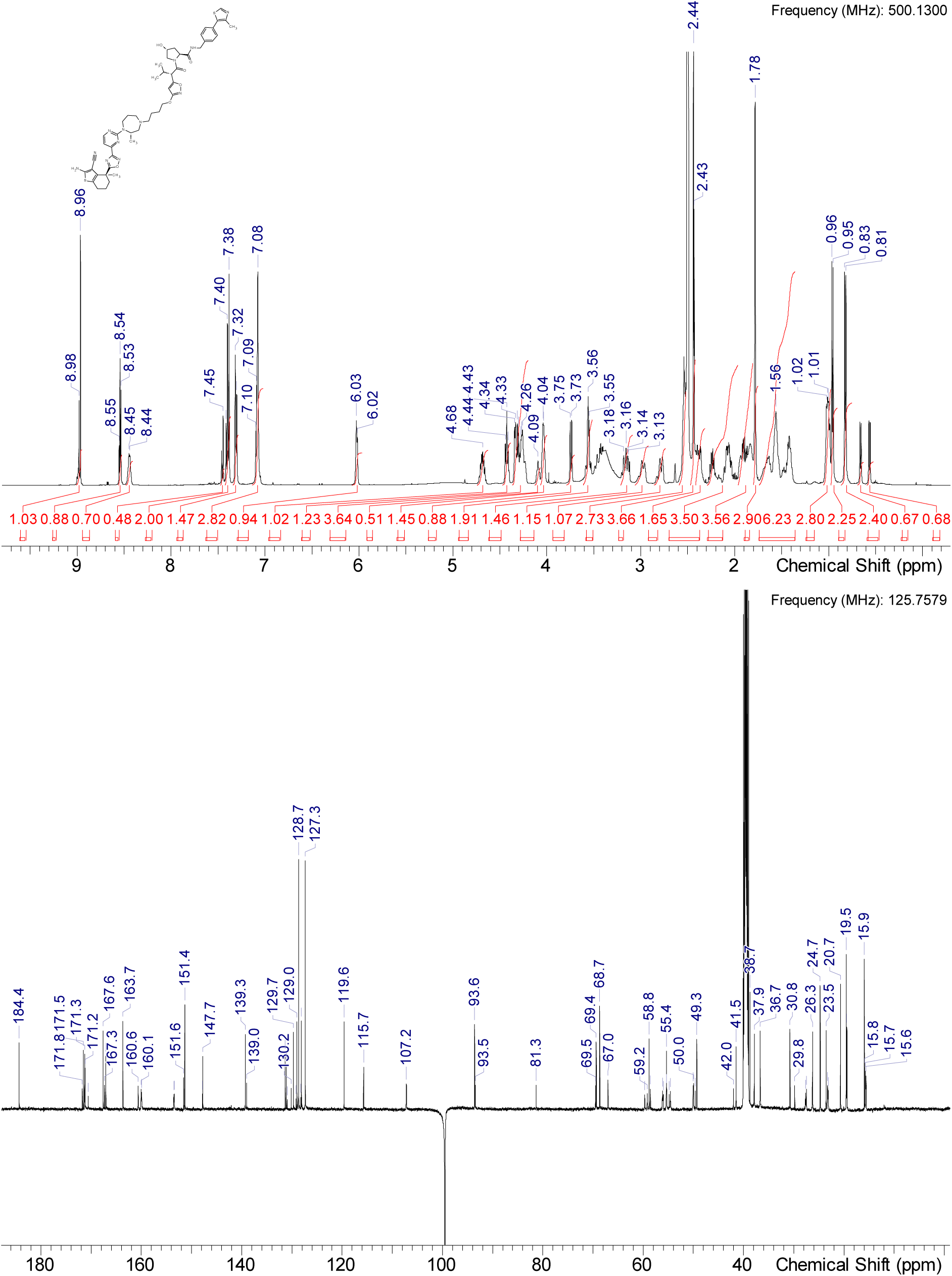

^1^H NMR (DMSO-d_6_) δ: 8.98 (s, 0.85H rotamer), 8.97 (s, 0.15H rotamer), 8.82 (s, 0.15H rotamer), 8.55 (d, J=4.4 Hz, 1H), 8.51 (t, J=6.0 Hz, 1H), 7.40-7.46 (m, 2H), 7.36-7.39 (m, 1.7H rotamer), 7.34 (d, J=8.2 Hz, 0.3H rotamer), 7.05-7.12 (m, 3H), 6.03-6.08 (m, 0.85H rotamer), 5.66 (br s, 0.15H rotamer), 5.10 (d, J=3.8 Hz, 0.85H rotamer), 5.02 (d, J=2.8 Hz, 0.15H rotamer), 4.67-4.75 (m, 0.85H rotamer), 4.64 (s, 0.15H rotamer), 4.18-4.40 (m, 5H), 4.10 (br s, 1.7H rotamer), 3.99-4.06 (m, 0.3H rotamer), 3.76 (dd, J=10.6, 4.3 Hz, 0.85H rotamer), 3.65 (br d, J=9.8 Hz, 0.85H rotamer), 3.54-3.61 (m, 0.3H rotamer), 3.44 (br d, J=10.7 Hz, 0.85H rotamer), 3.35-3.41 (m, 0.3H rotamer), 3.18 (br t, J=12.6 Hz, 1H), 2.94-3.07 (m, 1H), 2.82 (br d, J=12.0 Hz, 1H), 2.54 (br s, 4H), 2.41-2.46 (m, 4H), 2.16-2.30 (m, 1H), 1.81-2.14 (range, 6H), 1.78 (s, 3H), 1.37-1.74 (range, 7H), 1.02 (br s, 3H), 0.94 (d, J=6.3 Hz, 3H), 0.79 (br d, J=6.6 Hz, 3H)

^13^C NMR (DMSO-d_6_) δ: 184.4, 171.6, 171.5, 171.4, 171.2, 171.1, 171.0, 168.3, 167.4, 167.1, 163.7, 160.6, 160.1, 160.0, 153.5, 153.4, 151.4, 147.8, 147.8, 139.4, 138.9, 131.3, 131.1, 131.0, 130.1, 129.8, 128.9, 128.8, 127.8, 127.5, 119.6, 115.7, 107.2, 93.5, 93.1, 81.3, 69.5, 69.3, 68.6, 67.0, 59.7, 59.2, 59.1, 58.8, 56.2, 56.1, 55.7, 55.3, 54.9, 54.5, 50.0, 50.0, 49.9, 49.3, 42.1, 41.6, 38.7, 38.0, 36.7, 32.0, 31.1, 27.7, 27.5, 26.3, 24.7, 23.5, 23.3, 23.2, 20.4, 20.1, 20.0, 19.8, 19.4, 15.9, 15.9, 15.7, 15.6

More carbon peaks detected than present in the structure due to presence of rotamers

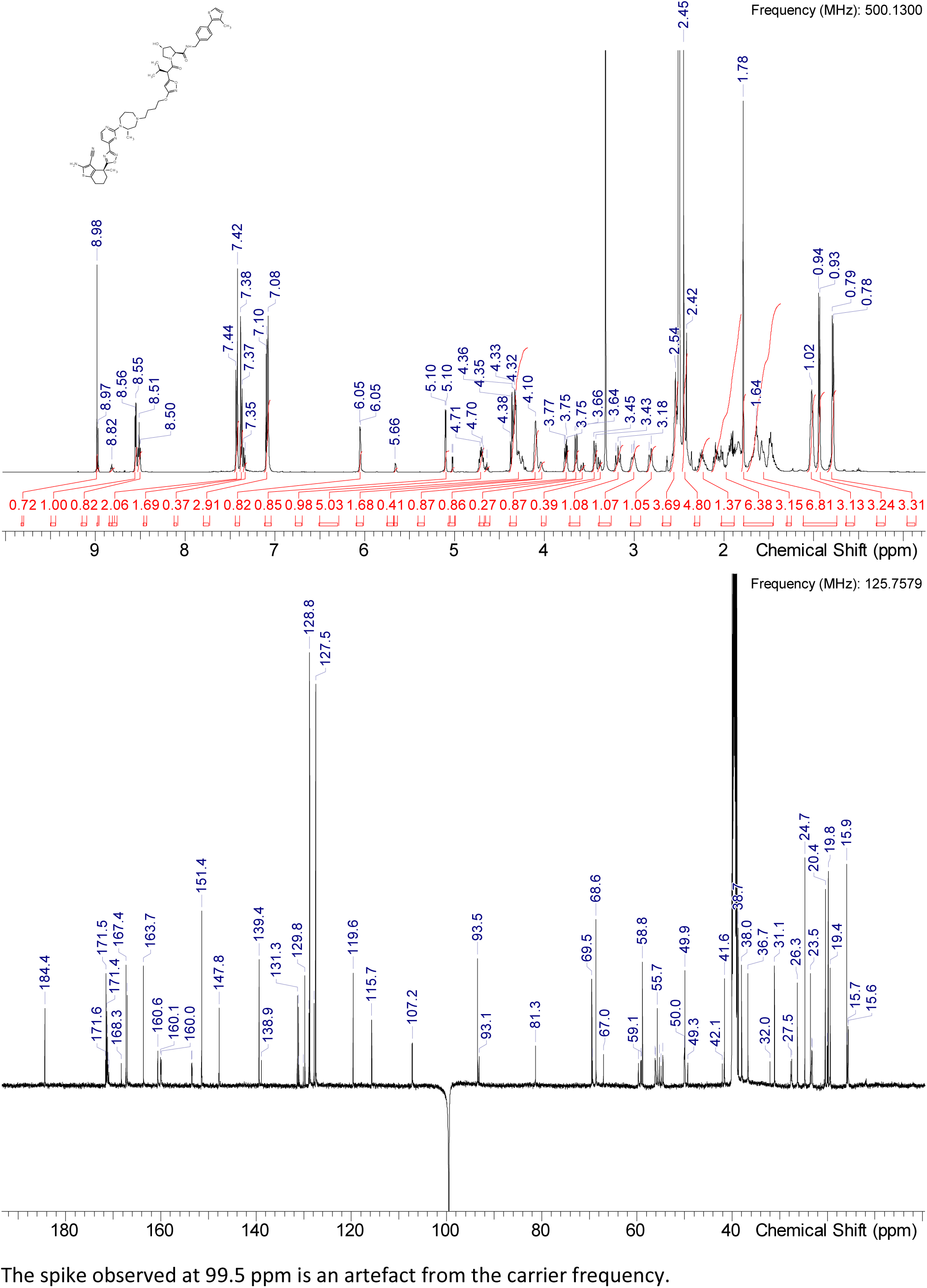

HRMS (m/z): [M+H]+ calcd. for C50H60N12O6S2 , 989.42730; found, 979.42767

### Synthesis of cis VHL negative control compound 5

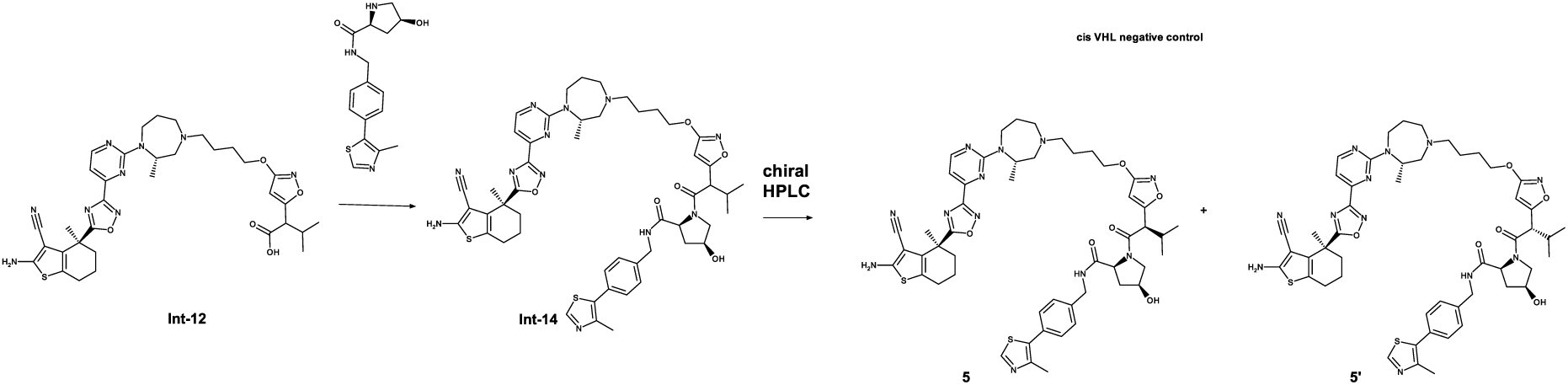

To a stirred solution of Int-12 (166 mg, 0.24 mmol, 1.0 eq.), (2S,4S)-4-hydroxy-N-{[4-(4-methyl-1,3-thiazol-5-yl)phenyl]methyl}pyrrolidine-2-carboxamide (115 mg, 0.361 mmol, 1.5 eq.) and HATU (112 mg, 0.29 mmol, 1.2 eq.) in a 1:1 mixture of DMSO and acetonitrile (3 ml) is added TEA (174 mL, 1.20 mmol, 5.0 eq) and the reaction mixture is stirred at rt for 30 min. After complete conversion the reaction mixture is quenched with water, diluted with acetonitrile and purified by RP chromatography (Method 3b) to obtain a mixture of diastereomers Int-14 (160 mg, 0.182, combined 67 % yield). The mixture is separated by preparative chiral HPLC (Method A) to obtain negative control compound 5 (46 mg, 0.045 mmol) and 5’ (61 mg, 0.062 mmol).

^1^H NMR (DMSO-d_6_) δ: 8.98 (s, 0.85H rotamer), 8.96 (s, 0.15H rotamer), 8.68-8.74 (m, 0.15H rotamer), 8.66-8.68 (m, 0.15H rotamer), 8.62 (br t, J=5.8 Hz, 0.85H rotamer), 8.50-8.57 (m, 1H rotamer), 7.29-7.47 (m, 4H), 7.01-7.15 (m, 3H), 6.06 (br d, J=5.7 Hz, 0.85H rotamer), 5.74 (br d, J=3.8 Hz, 0.15H rotamer), 5.43 (br s, 1H), 5.13-5.28 (m, 0.15H rotamer), 4.85-5.03 (m, 0.15H rotamer), 4.65-4.76 (m, 1H), 4.59-4.65 (m, 0.15H rotamer), 4.15-4.46 (range, 5H), 3.98-4.13 (range, 2H), 3.63-3.80 (m, 2H), 3.56 (br dd, J=10.1, 4.1 Hz, 1H), 3.45-3.50 (m, 0.15H rotamer), 3.17 (br t, J=12.6 Hz, 1H), 2.92-3.07 (m, 1H), 2.81 (br d, J=11.7 Hz, 1H), 2.53 (br s, 2H), 2.41-2.46 (m, 4H), 2.22-2.37 (m, 2H), 2.02-2.13 (m, 1H), 1.93 (br s, 1H), 1.83 (br d, J=3.2 Hz, 2H), 1.78 (s, 3H), 1.71-1.76 (m, 1H), 1.51-1.70 (range, 4H), 1.48 (br d, J=7.6 Hz, 2H), 1.19-1.27 (m, 2H), 1.01 (br d, J=6.3 Hz, 3H), 0.94 (br d, J=6.3 Hz, 2.55H rotamer), 0.87-0.91 (m, 0.45H rotamer), 0.81-0.87 (m, 0.45H rotamer), 0.79 (br d, J=6.6 Hz, 2.45H rotamer)

^13^C NMR (DMSO-d_6_) δ: 184.4, 172.3, 171.8, 171.4, 171.2, 171.0, 168.4, 167.6, 167.1, 163.7, 160.7, 160.1, 160.0, 153.5, 153.5, 151.5, 147.8, 139.1, 138.8, 131.3, 131.1, 131.0, 130.0, 129.8, 128.8, 128.8, 128.7, 127.8, 127.5, 127.4, 119.6, 115.7, 107.2, 107.2, 93.6, 93.2, 81.3, 69.5, 69.3, 69.0, 67.5, 59.7, 59.3, 59.2, 58.9, 56.2, 56.1, 55.4, 55.3, 54.8, 54.5, 50.0, 50.0, 49.7, 49.2, 42.2, 41.8, 38.7, 37.2, 36.7, 32.0, 31.5, 31.3, 30.8, 30.3, 29.4, 29.0, 28.7, 27.7, 27.5, 26.3, 24.7, 23.5, 23.3, 23.2, 22.1, 21.4, 20.4, 20.1, 19.9, 19.8, 19.7, 19.5, 19.4, 15.9, 15.7, 15.6,

More carbon peaks detected than present in the structure due to presence of rotamers

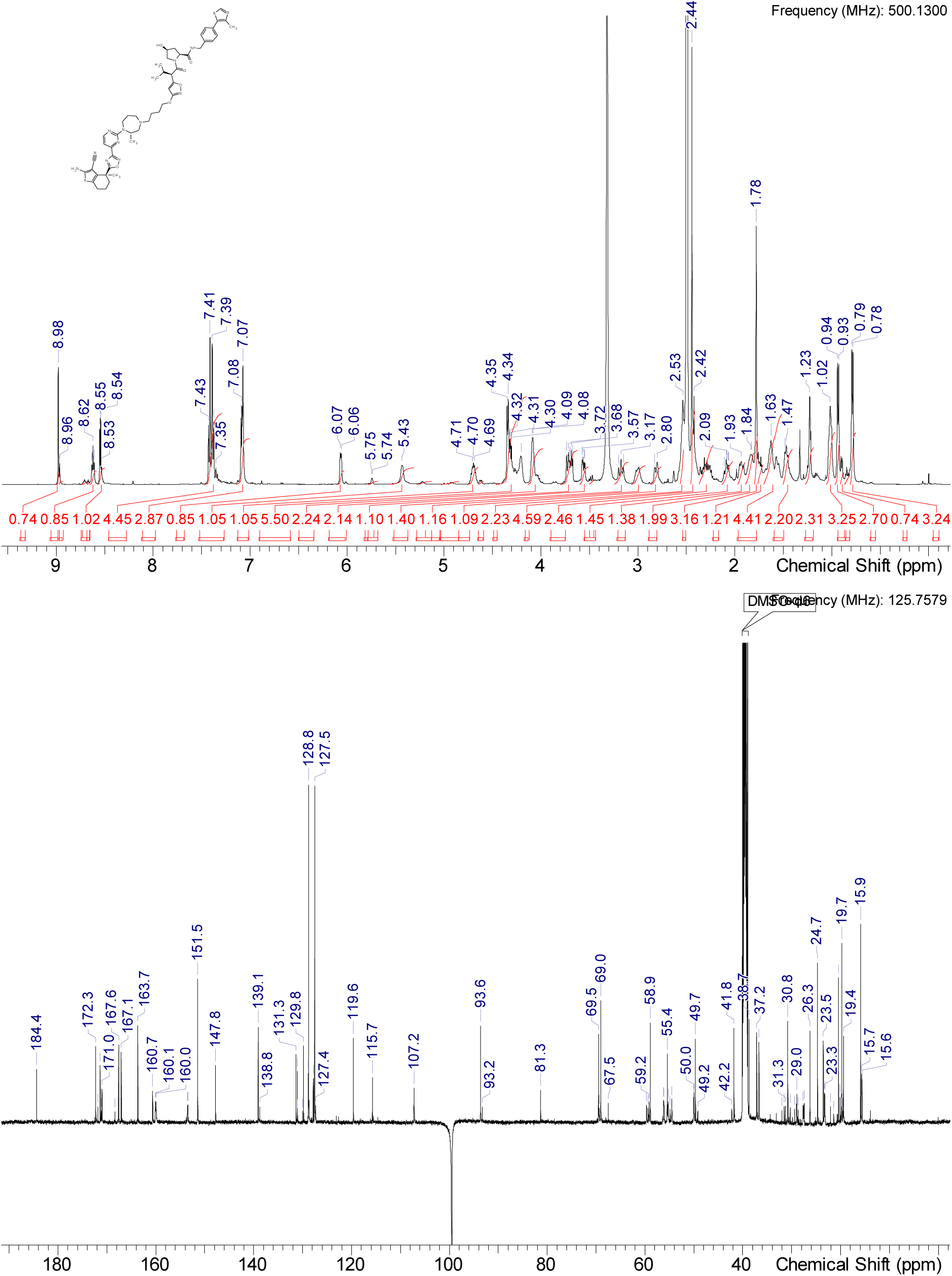

^1^H NMR (DMSO-d_6_) δ: 8.98 (s, 0.25H rotamer), 8.96 (s, 0.75H rotamer), 8.53 (d, J=4.7 Hz, 1H), 8.48 (br d, J=5.7 Hz, 0.75H rotamer), 7.26-7.51 (m, 4H), 7.02-7.15 (m, 3H), 6.05 (br d, J=6.0 Hz, 0.25H rotamer), 6.01 (br d, J=3.2 Hz, 0.75H rotamer), 5.37 (br d, J=6.6 Hz, 0.75H rotamer), 5.20 (br d, J=5.4 Hz, 0.25H rotamer), 4.68 (br s, 1H), 4.19-4.43 (range, 5H), 3.96-4.18 (m, 2H), 3.85 (br dd, J=10.4, 5.4 Hz, 1H), 3.76 (d, J=8.8 Hz, 0.75H rotamer), 3.61 (br dd, J=12.0, 5.0 Hz, 0.25H rotamer), 3.10-3.23 (m, 1H), 2.97 (br d, J=14.5 Hz, 1H), 2.74-2.85 (m, 1H), 2.51-2.57 (m, 3H), 2.43 (s, 4H), 2.36 (br s, 3H), 2.23 (br d, J=8.2 Hz, 1H), 2.07 (br d, J=10.1 Hz, 1H), 1.94 (br d, J=9.1 Hz, 1H), 1.83 (br s, 2H), 1.76-1.80 (m, 3H), 1.36-1.73 (range, 7H), 1.20-1.25 (m, 1H), 0.96-1.05 (m, 3H), 0.93 (d, J=6.6 Hz, 2.25H rotamer), 0.81 (d, J=6.6 Hz, 2.25H rotamer), 0.69 (br d, J=6.6 Hz, 0.75H rotamer), 0.60 (br d, J=6.9 Hz, 0.75H rotamer)

^13^C NMR (DMSO-d_6_) δ: 184.4, 172.1, 172.1, 171.4, 171.3, 171.1, 167.8, 167.5, 167.1, 163.7, 160.7, 160.6, 160.1, 159.9, 153.5, 153.4, 151.5, 151.4, 147.8, 147.7, 139.0, 138.9, 131.3, 131.1, 131.0, 130.1, 129.7, 128.9, 128.9, 128.8, 128.7, 128.1, 127.4, 127.3, 119.6, 115.7, 107.2, 93.5, 81.3, 69.5, 69.4, 69.1, 67.4, 59.7, 59.2, 58.9, 56.2, 56.0, 55.3, 55.1, 54.6, 50.1, 50.0, 49.7, 49.3, 42.2, 41.7, 38.7, 37.0, 36.7, 30.6, 29.9, 27.7, 27.5, 26.3, 24.7, 23.5, 23.2, 23.1, 20.7, 20.6, 19.5, 19.4, 15.9, 15.9, 15.7, 15.6,

More carbon peaks detected than present in the structure due to presence of rotamers

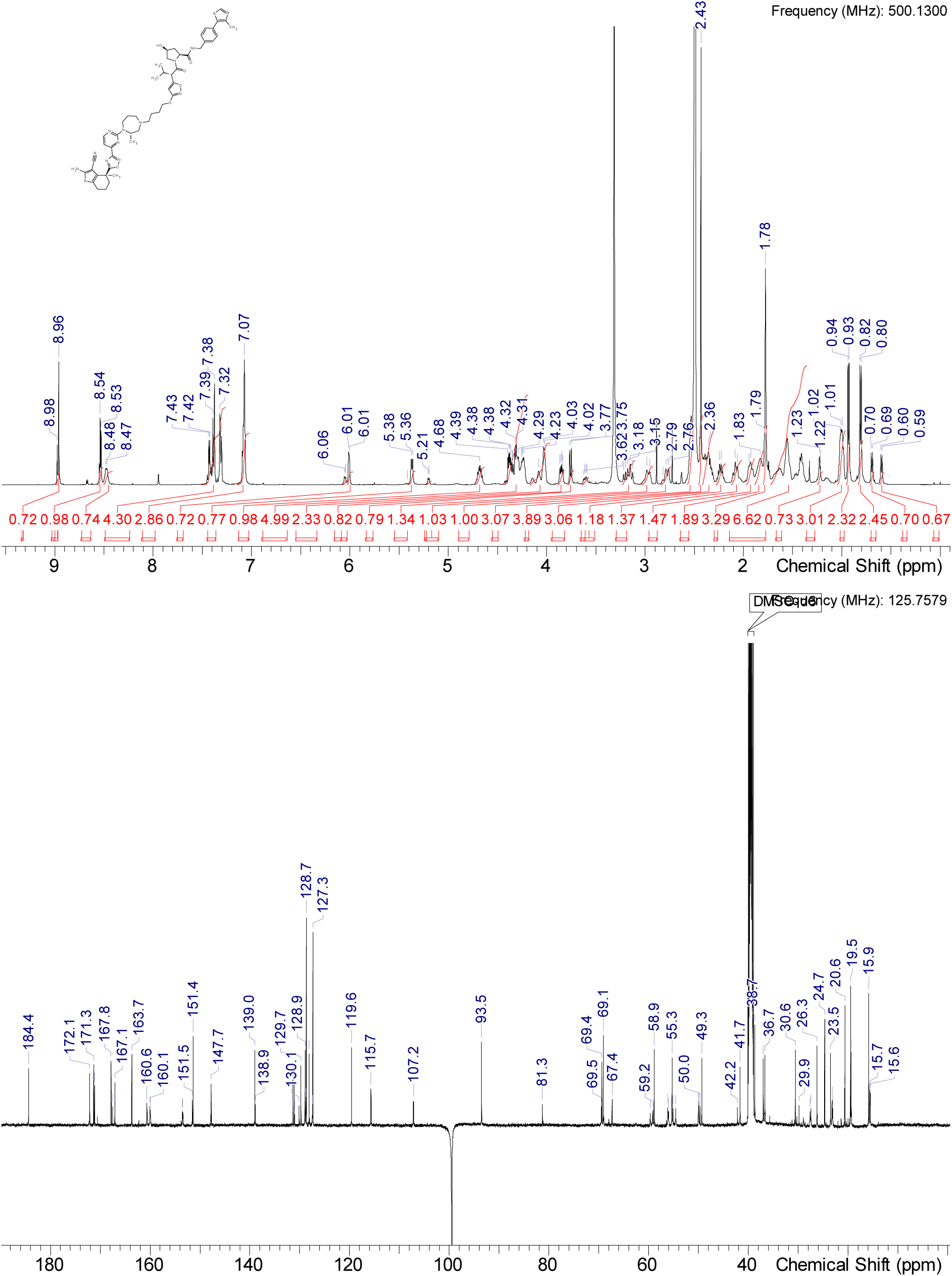

### Synthesis of Compound 7 (ACBI3)

#### Intermediate Int-15

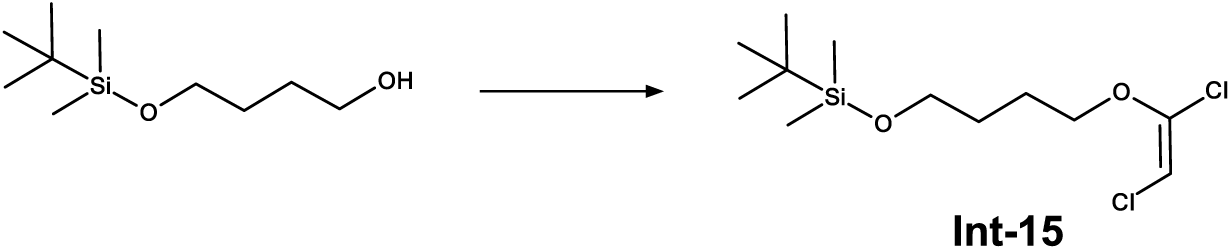

To a stirred solution of 4-[(tert-butyldimethylsilyl)oxy]butan-1-ol (20.00 g, 0.10 mol, 1.0 eq.) in THF (200 ml) is added sodium hydride (2.81 g, 0.12 mol, 1.2 eq.) at 0 °C and stirred at rt for 1 h. To the reaction mixture is cooled to 0 °C and trichloroethylene (15.38 g, 0.12 mol, 1.2 eq.) is added at 0 °C. The reaction mixture is warmed to rt and stirred for 48 h. The reaction mixture is quenched with ice cold water and the aqueous solution extracted with EtOAc (3 times). The combined organic layer is washed with water and brine, dried over sodium sulfate, filtered and concentrated under reduced pressure to afford the crude product which is purified by normal phase chromatography yielding Int-15 (20.00 g, 0.067 mol, 68% yield).

^1^H NMR (DMSO-d_6_) δ: 6.07 (s, 1H), 4.04 (t, J=6.3 Hz, 2H), 3.61 (t, J=6.2 Hz, 2H), 1.62-1.72 (m, 2H), 1.57 (dd, J=8.6, 5.6 Hz, 2H), 0.86 (s, 9H), 0.01-0.05 (m, 6H)

^13^C NMR (DMSO-d_6_) δ: 142.9, 97.8, 71.8, 62.0, 28.4, 25.8, 25.1, 17.9, -5.3

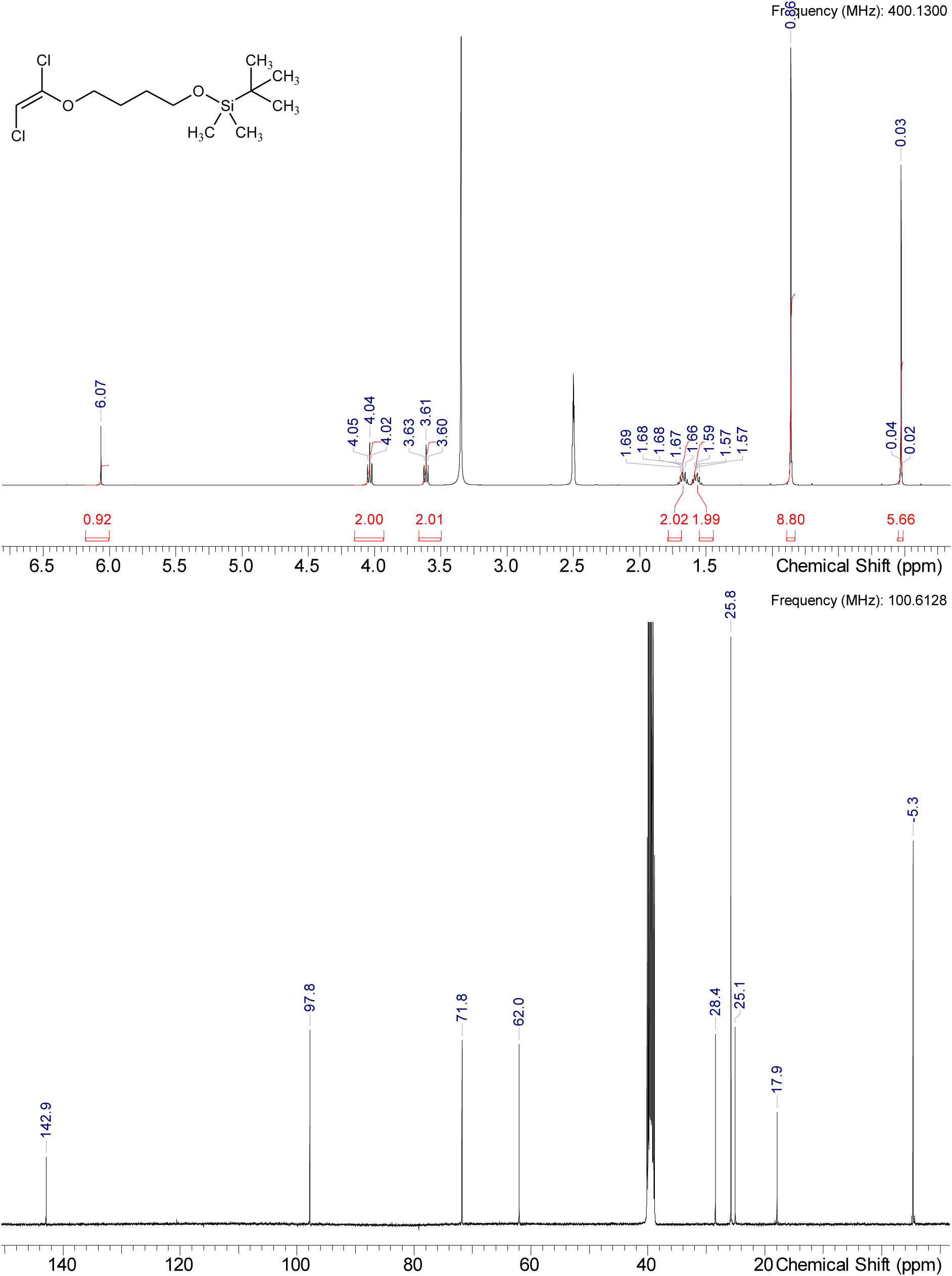

#### Intermediate Int-16

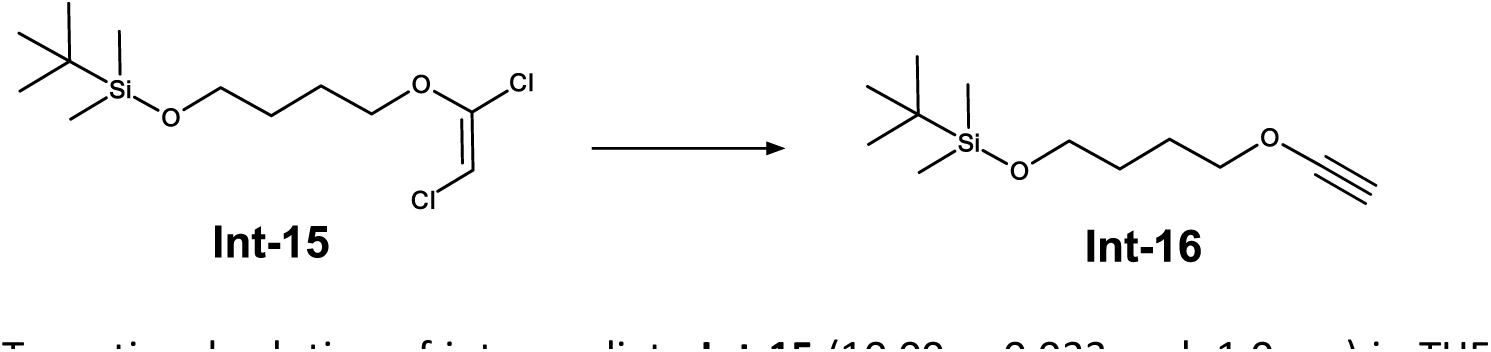

To a stirred solution of intermediate Int-15 (10.00 g, 0.033 mol, 1.0 eq.) in THF (200 ml) is added n-butyllithium solution (33 mL, 0. mol, 2.5 eq, 2.5 M in hexanes) at -78 °C and stirred at -40 °C for 2 h. After complete conversion the mixture is quenched with saturated ammonium chloride solution and diluted with ice cold water. The aqueous solution extracted with EtOAc (3 x). The combined organic layer is washed with water and brine, dried over sodium sulfate, filtered, and concentrated under reduced pressure to afford the crude product. The crude product is purified by column chromatography yielding intermediate Int-16 (6.98 g, 0.031 mmol, 92% yield).

^1^H NMR (DMSO-d_6_) δ: 4.10 (t, J=6.6 Hz, 2H), 3.60 (t, J=6.2 Hz, 2H), 2.33 (s, 1H), 1.72 (dd, J=8.6, 6.3 Hz, 2H), 1.43-1.57 (m, 2H), 0.83-0.88 (m, 9H), 0.01-0.04 (m, 6H) ^13^C NMR (DMSO-d_6_) δ: 90.9, 78.4, 61.9, 28.4, 28.0, 25.8, 24.8, 17.9, -5.4

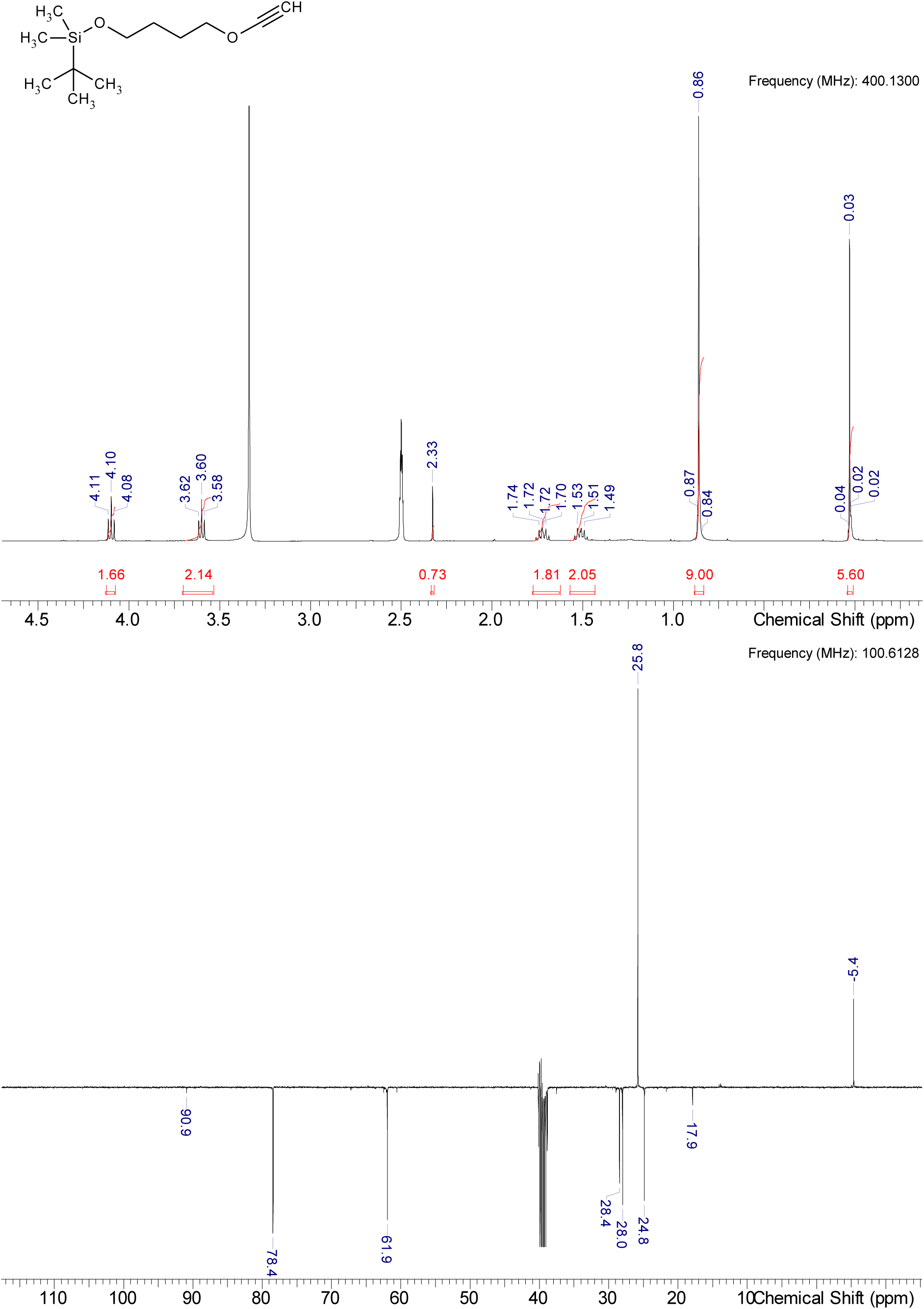

#### Intermediate Int-18

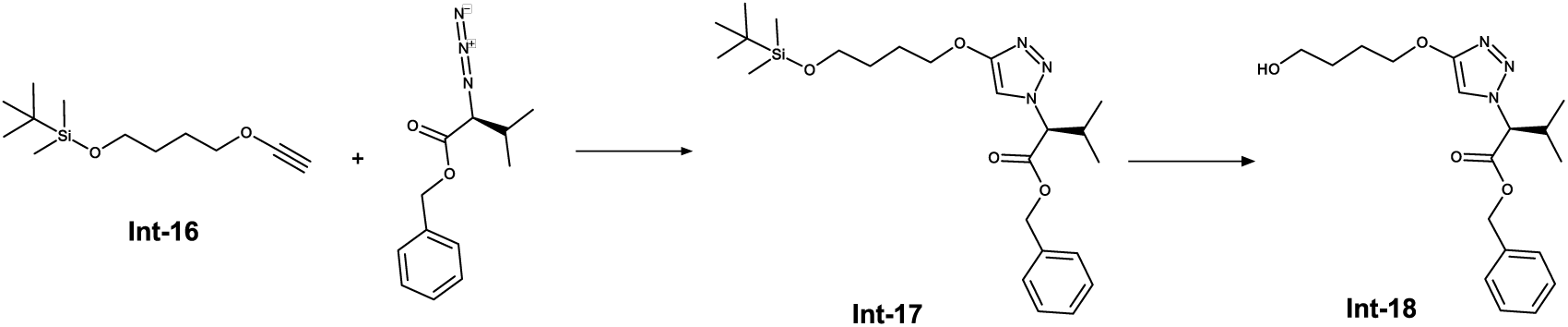

To a stirred solution of intermediate Int-16 (39.95 g, 174.91 mmol, 2.0 eq.) in EtOAc (200 ml) and water (200 ml) is added methyl (2R)-2-azido-3-methylbutanoate (20.40 g, 87.45 mmol, 1.0 eq.), sodium L-ascorbate (17.3 g, 87.37 mmol, 1 eq) and copper(II) sulfate pentahydrate (6.99 g, 43.96 mmol, 0.5 eq) at 20 °C. The reaction is stirred for 3 h at 20 °C. To the reaction mixture is added water (200 ml) and EtOAc the phases are separated. The organic layer is washed with brine and dried over sodium sulfate, filtered, and concentrated under reduced pressure to afford the crude product Int-17 as yellow oil which is directly used in the next step without further purification.

^1^H NMR (DMSO-d_6_) δ: 7.76 (s, 1H), 7.30-7.40 (m, 5H), 5.12-5.26 (m, 3H), 4.08 (td, J=6.6, 1.6 Hz, 2H), 3.62 (t, J=6.3 Hz, 2H), 2.52-2.58 (m, 1H), 1.73 (dd, J=8.8, 6.3 Hz, 2H), 1.52-1.62 (m, 2H), 0.92 (d, J=6.6 Hz, 3H), 0.86 (s, 9H), 0.77 (d, J=6.6 Hz, 3H), 0.03 (s, 6H)

^13^C NMR (DMSO-d_6_) δ: 168.0, 160.3, 135.2, 128.4, 128.3, 128.1, 106.8, 70.1, 68.5, 66.9, 62.1, 30.4, 28.6, 25.8, 25.3, 18.7, 18.3, -5.3, 1 carbons only observed in HSQC

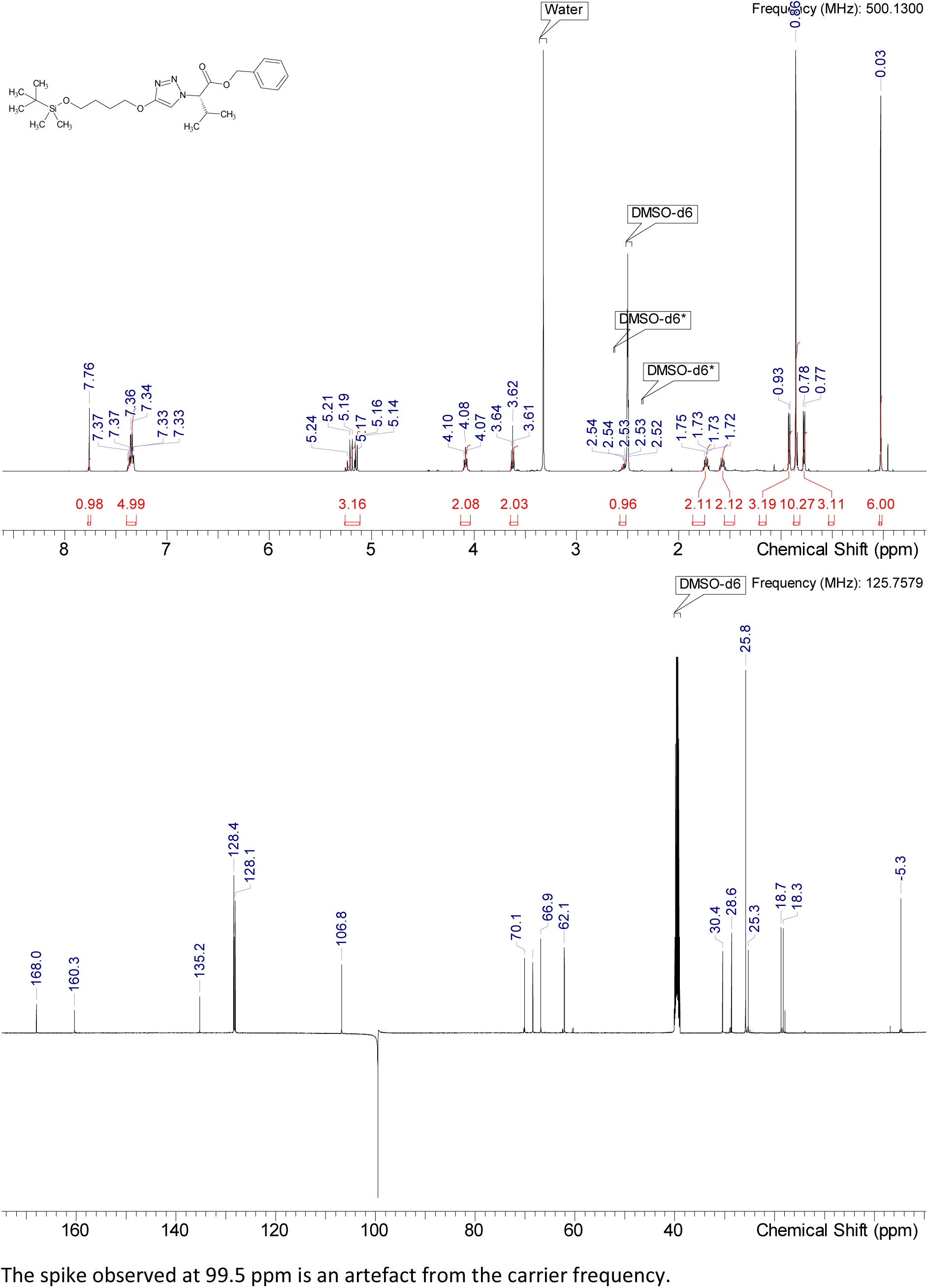

HRMS (m/z): [M+H]+ calcd. for C24H39N3O4Si , 348.19153; found, 347.18425 (-C6H14Si)

To the stirred solution of the obtained crude mixture in in THF (150 mL) at 10 °C is added HF-pyridine (38.6 g, 389.88 mmol) and the mixture is stirred at 10-20 °C for 2 h. To the mixture is added aqueous NaHCO_3_ solution (1 M, 1.5 L) and the pH is adjusted to pH 7-8. The phases are separated, and the aqueous phase is extracted with ethyl acetate and the combined organic layers are dried aver Na_2_SO_4_. The crude product is purified by column chromatography (Petrol ether/EtOAC, 0-10%) to obtain intermediate Int-18 (23.0 g, 66.20 mmol, 76 % yield over two steps).

^1^H NMR (DMSO-d_6_) δ: 7.77 (s, 1H), 7.28-7.42 (m, 5H), 5.20 (d, J=11.7 Hz, 2H), 5.15 (d, J=8.5 Hz, 1H), 4.45 (br s, 1H), 3.99-4.11 (m, 2H), 3.43 (t, J=6.5 Hz, 2H), 2.54 (m, 1H), 1.68-1.76 (m, 2H), 1.53 (dd, J=8.5, 6.3 Hz, 2H), 0.92 (d, J=6.6 Hz, 3H), 0.78 (d, J=6.6 Hz, 3H)

^13^C NMR (DMSO-d_6_) δ: 168.0, 160.4, 135.3, 128.5, 128.3, 128.1, 106.8, 70.2, 68.5, 66.9, 60.3, 30.4, 28.8, 25.4, 18.7, 18.3

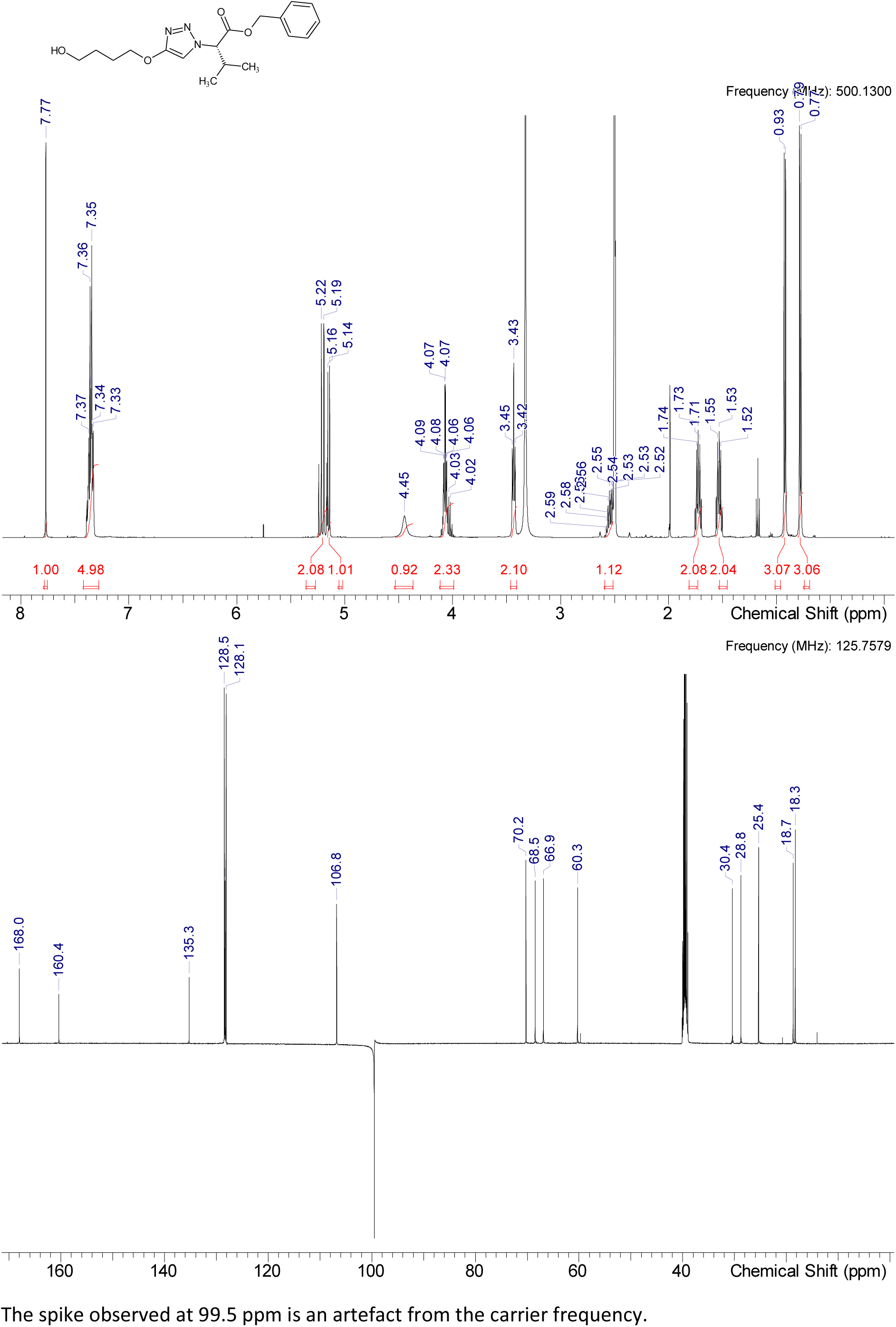

HRMS (*m/z*): [M+H]^+^ calcd. for C18H25N3O4 , 348.19178; found, 348.1911

#### Intermediate Int-19

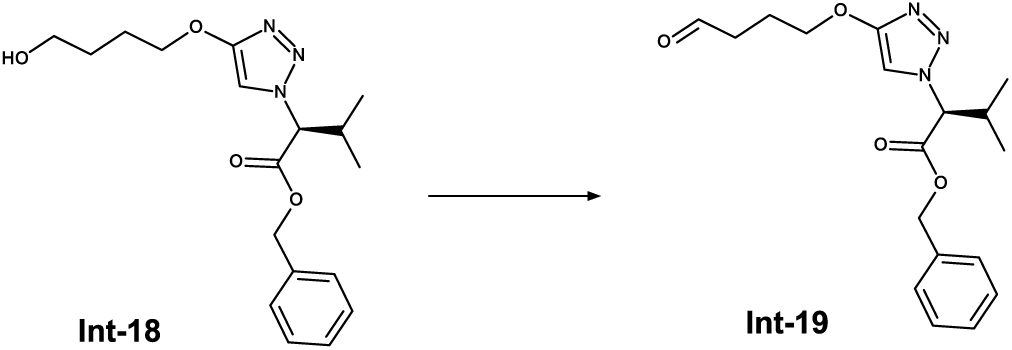

To a stirred solution of intermediate Int-18 (1.50 g, 4.272 mmol, 1.0 eq) in DCM (30 mL) is added Dess-Martin periodinane (1.81 g, 4.272 mmol, 1.0 eq) and stirred at rt for 2.5 h. To the mixture is added saturated aqueous NaHCO3 solution and the mixture is stirred for 10 min. The mixture is diluted with DCM and the phases are separated. The organic phase is concentrated under reduced pressure and the crude product is purified by column chromatography (Hexane/EtOAc, 0-40%) to obtain Int-19 (1.2 g, 3.47 mmol, 81 % yield).

^1^H NMR (DMSO-d_6_) δ: 9.70 (t, J=1.1 Hz, 1H), 7.78 (s, 1H), 7.31-7.40 (m, 5H), 5.10-5.30 (m, 3H), 4.08 (t, J=6.6 Hz, 2H), 2.52-2.62 (m, 3H), 1.96 (quin, J=6.8 Hz, 2H), 0.92 (d, J=6.8 Hz, 3H), 0.78 (d, J=6.6 Hz, 3H)

^13^C NMR (DMSO-d_6_) δ: 202.9, 168.0, 160.2, 135.3, 128.5, 128.4, 128.2, 107.0, 69.5, 68.5, 66.9, 30.4, 21.5, 18.7, 18.3, 1C under DMSO

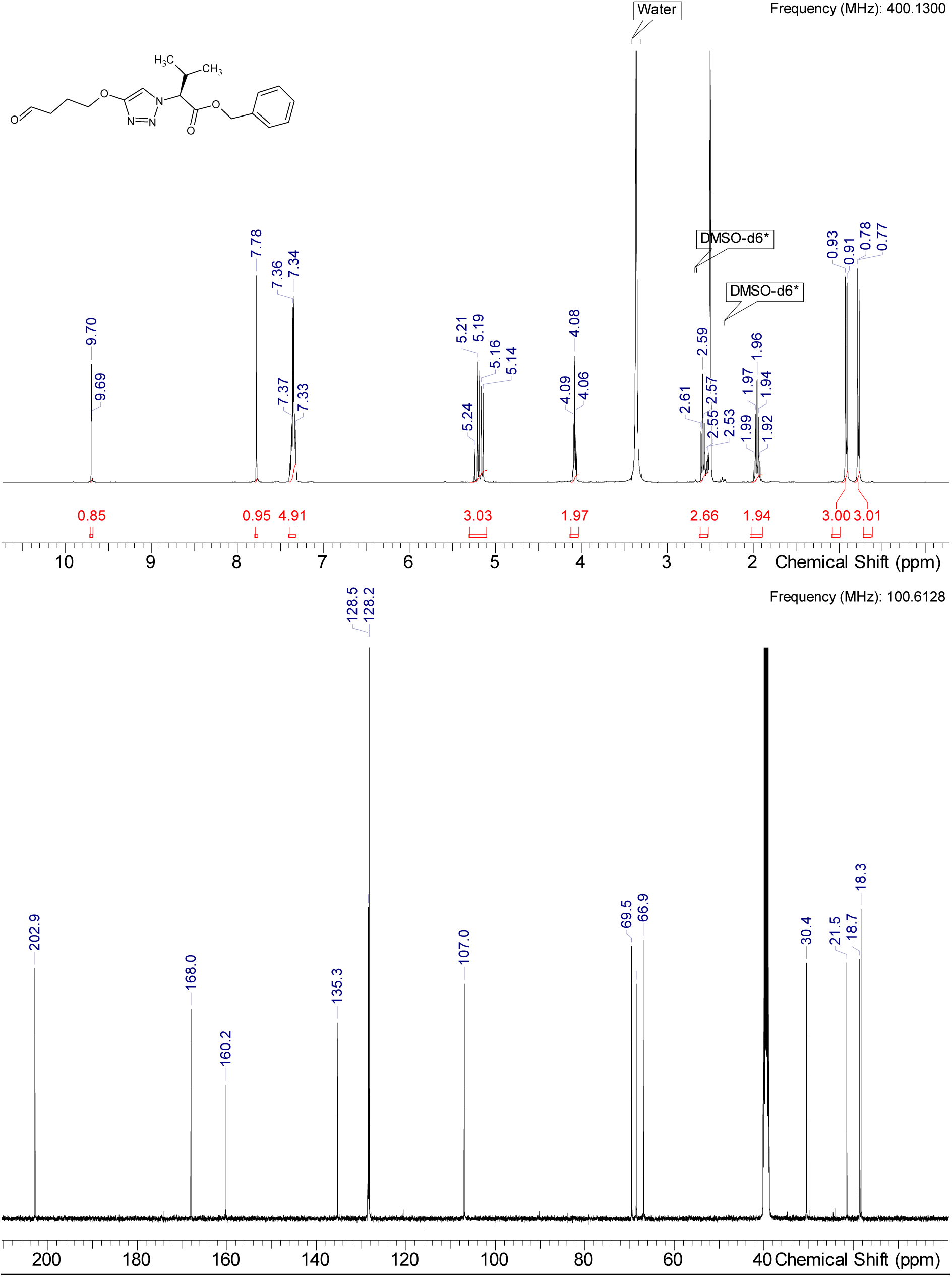

HRMS (m/z): [M+H]+ calcd. for C18H23N3O4 , 346.17613; found, 346.17587

#### Intermediate Int-20

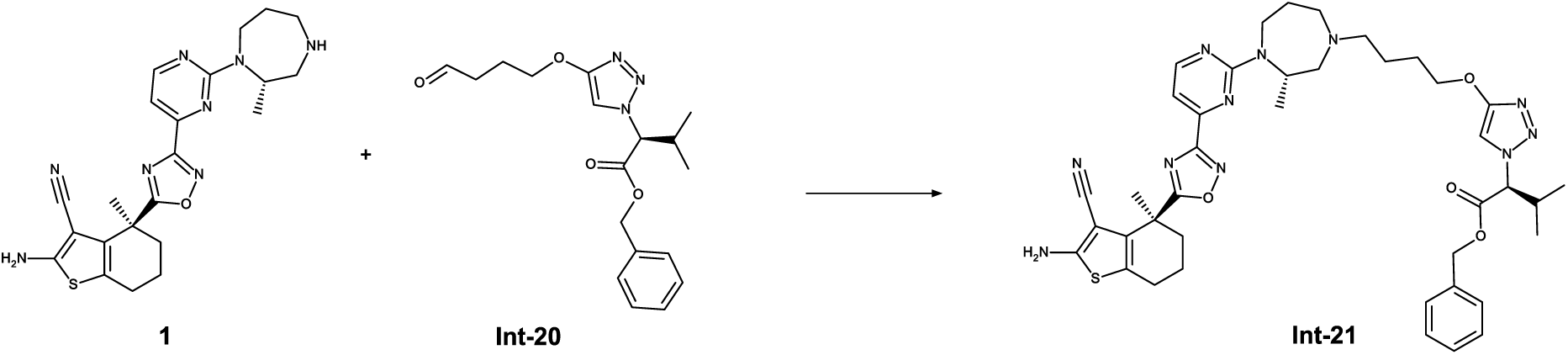

To a stirred solution of 1 (200 mg, 0.444 mmol, 1.0 eq.) and sodium triacetoxy borohydride (235 mg, 1,110 mmol, 2.5 eq.) in DMF (3 mL) at RT is slowly added a solution of Int-20 (153 mg, 0.444 mmol, 1.0 eq.) in DMF (1 mL). The reaction is stirred for 30 min before it is quenched by addition of water. The solvents are removed under reduced pressure and the crude product is purified by chromatography (Method 3a) to obtain Int-21 (1.33 g, 1.71 mmol, 84 % yield).

^1^H NMR (DMSO-d_6_) δ: 8.55 (d, J=4.7 Hz, 1H), 7.74 (s, 1H), 7.30-7.39 (m, 5H), 7.09 (s, 1H), 7.08 (s, 2H), 5.19 (d, J=11.0 Hz, 2H), 5.13 (br d, J=8.8 Hz, 1H), 4.65-4.75 (m, 1H), 4.21-4.37 (m, 1H), 4.03 (br s, 2H), 3.18 (br t, J=12.6 Hz, 1H), 3.01 (br d, J=13.9 Hz, 1H), 2.82 (br d, J=11.7 Hz, 1H), 2.51-2.59 (m, 4H), 2.37-2.46 (m, 2H), 2.01-2.19 (m, 1H), 1.80-1.98 (m, 3H), 1.78 (s, 3H), 1.44-1.74 (m, 6H), 1.02 (br s, 3H), 0.91 (d, J=6.6 Hz, 3H), 0.77 (q, J=6.6 Hz, 3H), 1H under DMSO

^13^C NMR (DMSO-d_6_) δ: 184.4, 168.0, 167.1, 163.7, 160.4, 160.1, 160.0, 153.6, 153.5, 135.2, 131.3, 128.5, 128.3, 128.1, 119.6, 115.7, 107.2, 106.8, 81.3, 70.2, 68.5, 66.9, 59.7, 59.2, 56.2, 56.1, 55.3, 54.6, 50.1, 50.0, 38.7, 36.7, 30.4, 27.7, 27.5, 26.4, 24.7, 23.6, 23.4, 23.3, 19.4, 18.7, 18.3, 15.7, 15.6

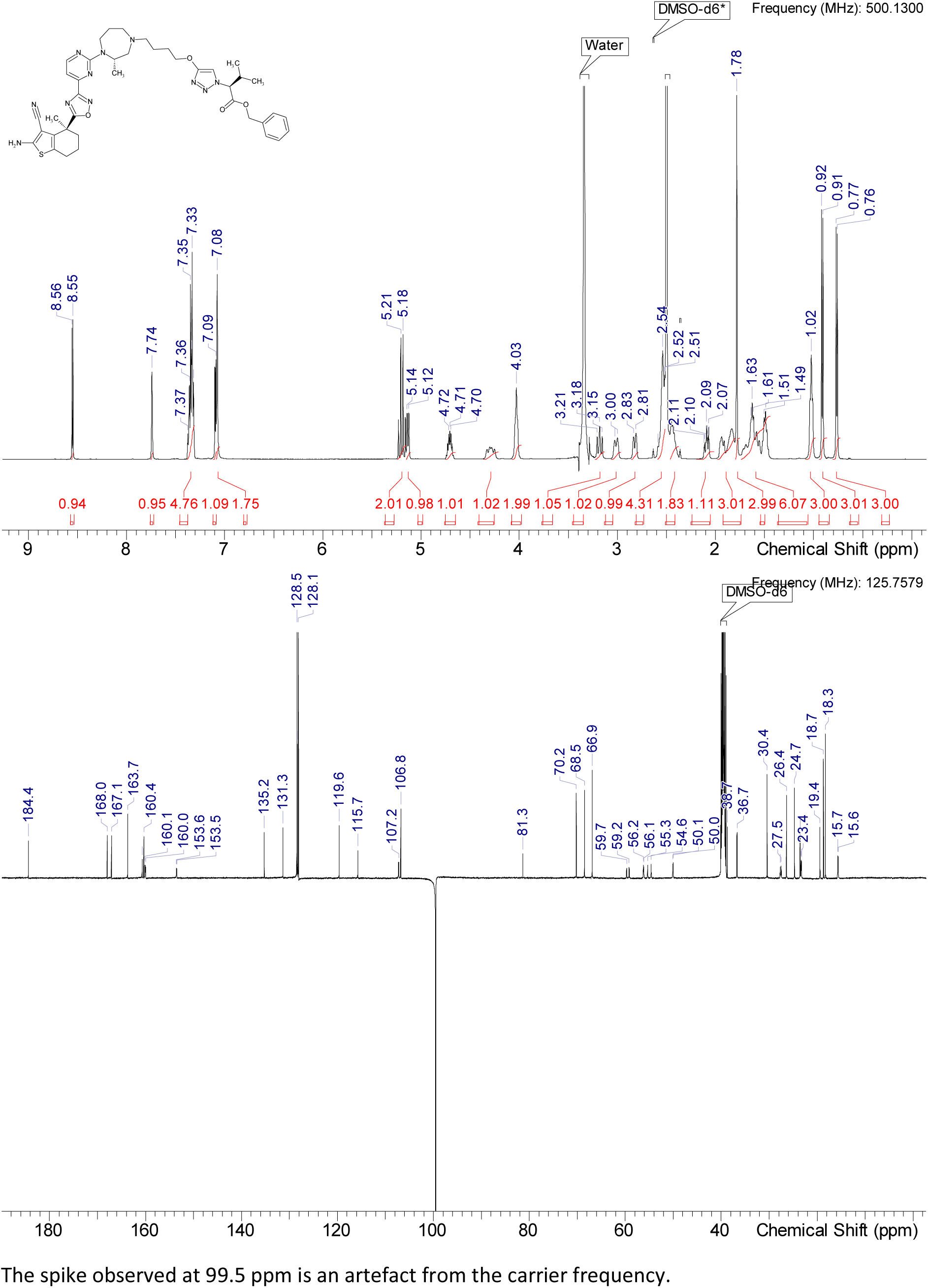

HRMS (m/z): [M+H]+ calcd. for C40H49N11O4S , 780.37625; found, 780.37671

#### Intermediate Int-22

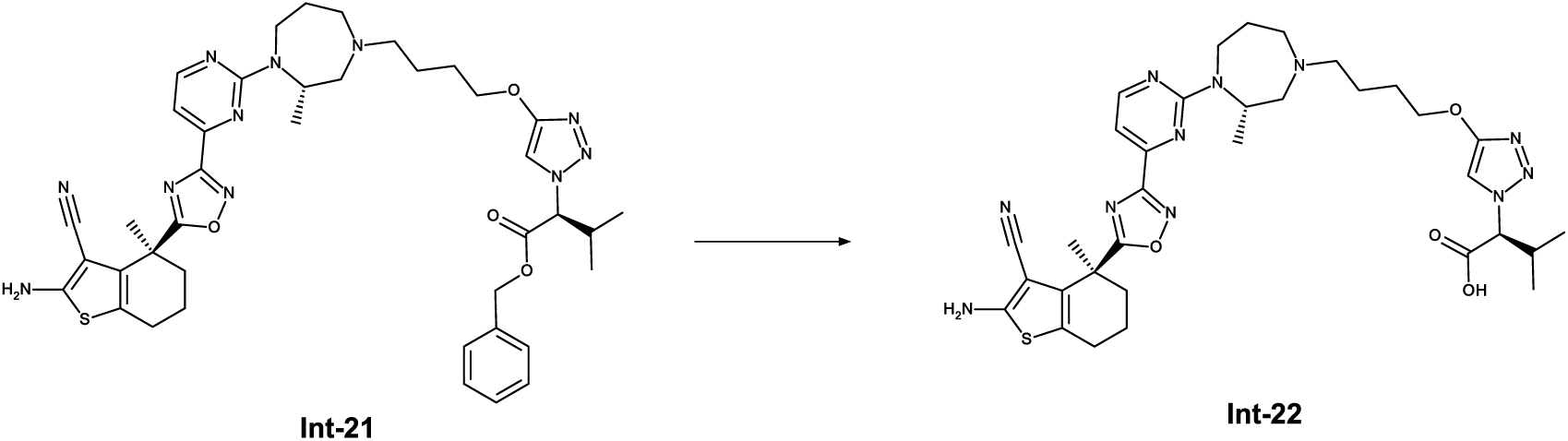

To a solution of intermediate Int-21 (1.33 g, 1.705 mmol, 1.0 eq.) in MeOH (25 mL) in a hydrogenation reactor is added Pd/C (10%, 350.00 mg). The reaction mixture stirred under a pressure of 8 bar H_2_ for 8 h. After complete conversion the reaction mixture is filtered, and the solvent is removed under reduced pressure to give intermediate Int-22 (1.12 g, 1.624 mmol, 95 % yield) which was used without further purification.

^1^H NMR (DMSO-d_6_) δ: 8.51 (d, J=4.8 Hz, 1H), 7.57 (s, 1H), 6.96-7.19 (m, 3H), 4.71 (br d, J=7.1 Hz, 2H), 4.17-4.39 (m, 1H), 3.97 (br t, J=6.0 Hz, 3H), 2.66-2.96 (m, 3H), 2.50-2.65 (m, 3H), 2.23-2.38 (m, 2H), 1.95-2.08 (m, 1H), 1.45-1.95 (m, 14H), 0.92-1.06 (m, 3H), 0.84 (d, J=6.6 Hz, 3H), 0.67 (d, J=6.8 Hz, 3H)

^13^C NMR (DMSO-d_6_) δ: 184.4, 169.6, 167.1, 163.7, 160.6, 160.1, 153.5, 131.3, 120.6, 119.6, 115.7, 107.5, 106.5, 81.3, 70.1, 69.9, 58.7, 56.0, 55.8, 49.3, 38.7, 36.7, 30.5, 26.8, 26.3, 24.7, 23.5, 19.4, 19.1, 18.4, 15.6, 15.5, 1C under DMSO

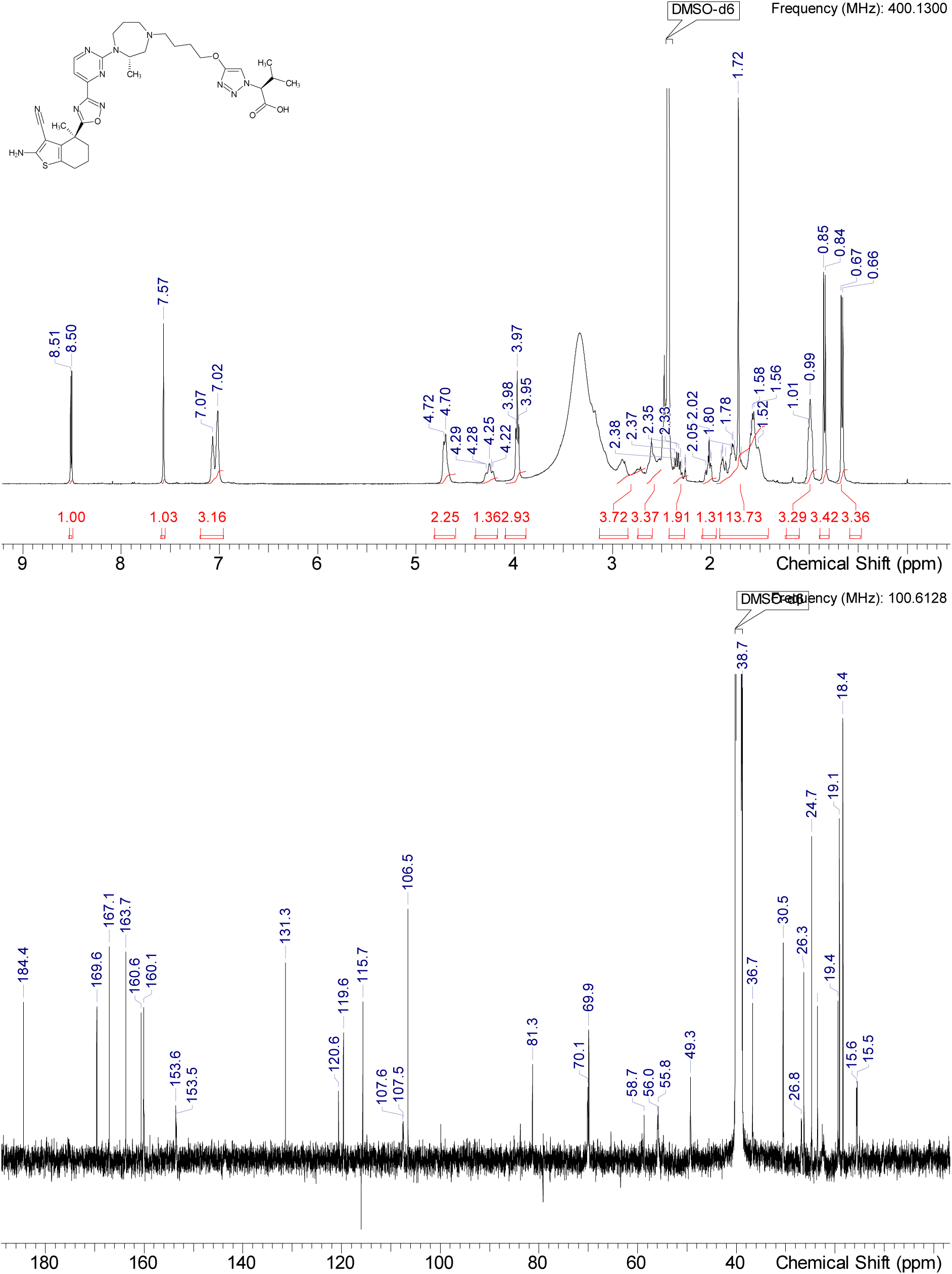

HRMS (m/z): [M+H]+ calcd. for C33H43N11O4S , 689.32202; found, 689.32904

#### ACBI3

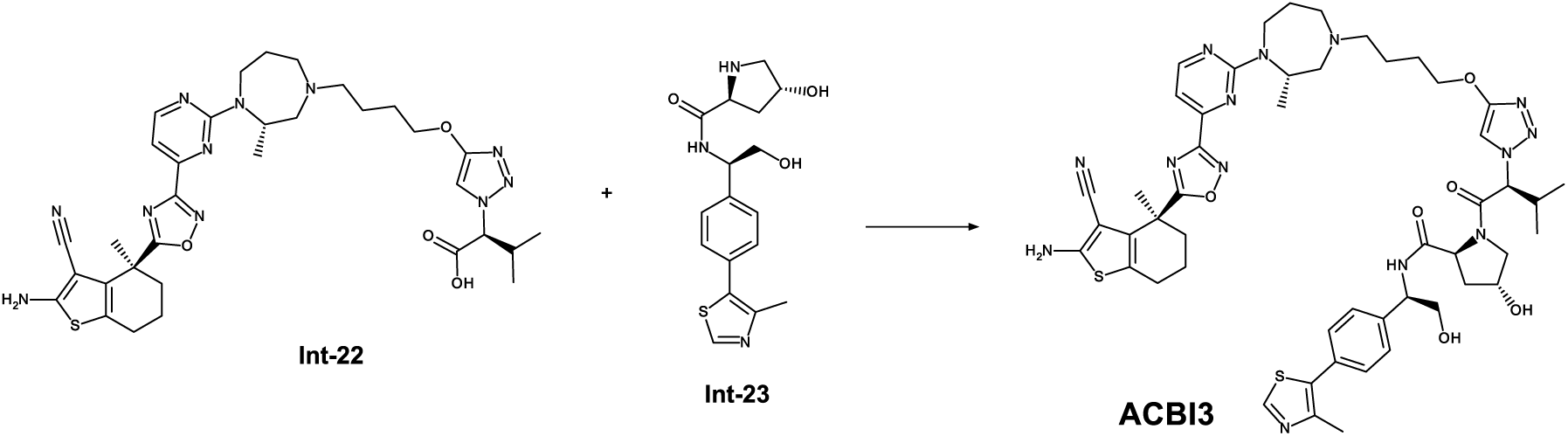

To a stirred solution of intermediate Int-22 (69 mg, 0.10 mmol, 1.0 eq.), amine Int-23 (55 mg, 0.13 mmol, 1.3 eq.) in a mixture of DCM (1 mL) and acetonitrile (0.35 mL) is added triethylamine (83.8 µl, 0.60 mmol, 6.0 eq.) at rt and the mixture is stirred for 5 min before propane phosphonic acid anhydride (119.0 µl, 50% solution in ethyl acetate, 0.200 mmol, 2.0 eq.) is added and the mixture is stirred for another 20 min. The mixture is diluted with DCM and saturated aqueous NaHCO_3_ solution is added before it is stirred for 5 min. The phased are separated and the organic layer is concentrated under reduced pressure and the residue is purified by RP HPLC (Method 3b) and lyophilized to obtain ACBI3 (63 mg, 0.062 mmol, 62 % yield).

^1^H NMR (DMSO-d_6_) δ: 8.98 (s, 1H), 8.52-8.59 (m, 1H), 8.47 (d, J=7.9 Hz, 1H), 7.65 (s, 1H), 7.41-7.49 (m, 2H), 7.37 (d, J=8.2 Hz, 2H), 7.04-7.14 (m, 3H), 5.14 (br s, 2H), 4.85 (br s, 2H), 4.65-4.76 (m, 1H), 4.44 (s, 1H), 4.21-4.37 (m, 2H), 3.97-4.10 (m, 2H), 3.75 (br d, J=6.6 Hz, 1H), 3.53-3.67 (m, 3H), 3.12-3.24 (m, 1H), 2.95-3.08 (m, 1H), 2.79-2.89 (m, 1H), 2.52-2.58 (m, 3H), 2.45-2.47 (m, 3H), 2.35-2.43 (m, 2H), 2.01-2.13 (m, 2H), 1.82-1.99 (m, 3H), 1.79 (s, 5H), 1.44-1.74 (m, 7H), 1.02 (br d, J=6.6 Hz, 6H), 0.66 (br d, J=6.3 Hz, 3H)

^13^C NMR (DMSO-d_6_) δ: 184.4, 170.5, 167.1, 166.1, 163.7, 160.7, 160.5, 160.1, 160.0, 153.6, 153.5, 151.5, 147.7, 141.2, 131.4, 131.1, 129.9, 128.7, 128.6, 127.5, 127.4, 119.6, 115.7, 107.2, 105.6, 81.3, 70.1, 68.6, 67.1, 64.5, 58.8, 56.1, 55.4, 54.7, 50.1, 50.0, 37.8, 36.7, 31.6, 27.7, 27.5, 26.4, 24.7, 23.6, 23.4, 23.2, 19.4, 18.3, 18.1, 16.0, 15.7, 15.6, 1C under DMSO

More carbon peaks detected than present in the structure due to presence of rotamers

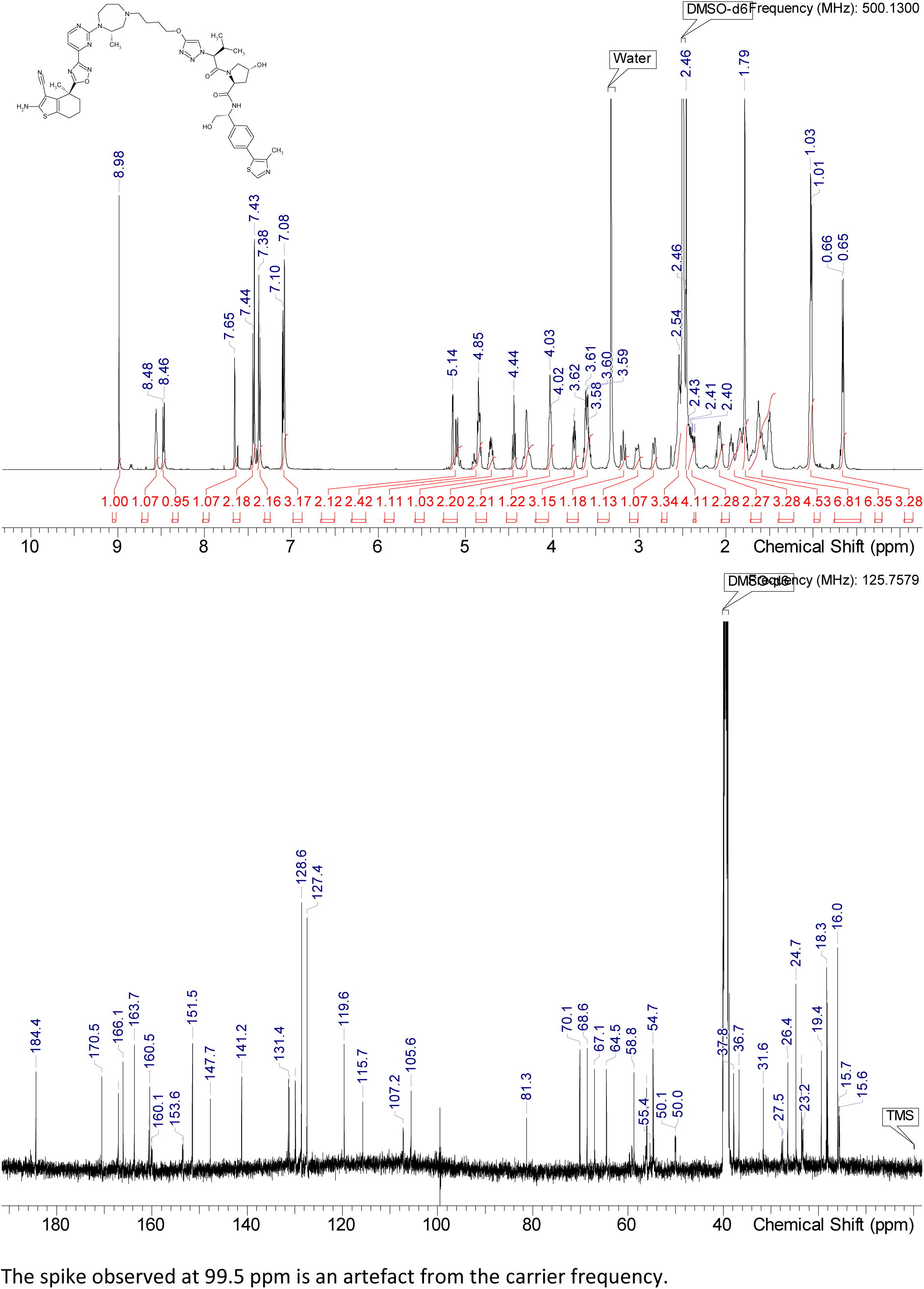

HRMS (m/z): [M+H]+ calcd. for C50H62N14O6S2 , 1019.44909; found, 1019.44867

### Synthesis of the negative control compound 8 (cis-ACBI3)

#### Intermediate Int-24

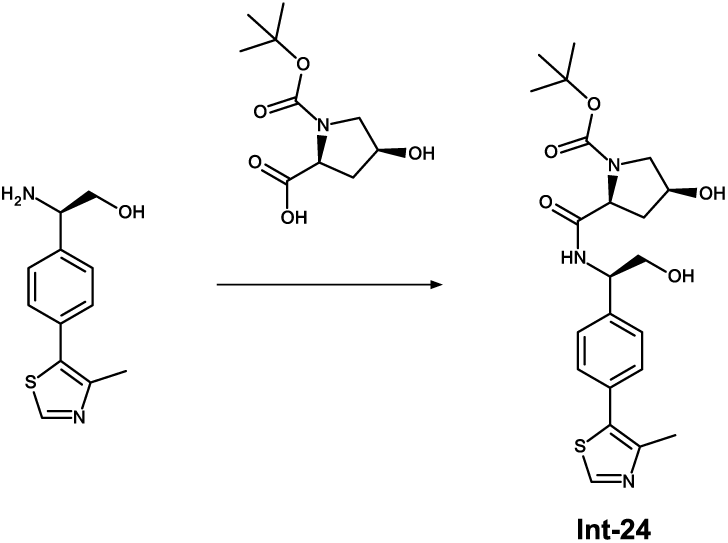

To a stirred solution of (2S,4S)-1-[(tert-butoxy)carbonyl]-4-hydroxypyrrolidine-2-carboxylic acid (70 mg, 0.284 mmol, 1.0 eq.) in DMSO ( 1 mL) is added (2R)-2-amino-2-[4-(4-methyl-1,3-thiazol-5-yl)phenyl]ethan-1-ol (93 mg, 0.303 mmol, 81%, 1.1 eq.), HATU (173 mg, 0.440 mmol, 1.5 eq. and DIPEA (0.256 mL, 1.468 mmol, 5.0 eq.) and the mixture is stirred at RT for 20 min. The mixture is diluted with water and acetonitrile and purified by RP HPLC (Method 3a) and lyophilized to obtain Int-24 (84 mg, 0.188 mmol, 63% yield).

^1^H NMR (DMSO-d_6_) δ: 8.98 (s, 1H), 8.30 (br d, J=7.9 Hz, 0.66H rotamer), 8.24 (br d, J=7.6 Hz, 0.35H rotamer), 7.38-7.46 (m, 4H), 5.19 (br d, J=5.6 Hz, 0.34H rotamer), 5.16 (br d, J=6.1 Hz, 0.67H rotamer), 4.77-4.92 (m, 2H), 4.11-4.21 (m, 2H), 3.64 (br d, J=3.5 Hz, 2H), 3.49 (dd, J=10.6, 5.3 Hz, 1H), 3.14-3.26 (m, 1H), 2.45 (s, 3H), 2.24-2.39 (m, 1H), 1.63-1.77 (m, 1H), 1.42 (br s, 3.17H rotamer), 1.32 (s, 5.84H rotamer)

^13^C NMR (DMSO-d6) δ: 172.5, 172.2, 153.9, 153.4, 151.5, 147.7, 141.0, 131.1, 129.9, 128.6, 127.5, 127.4, 78.9, 78.8, 68.8, 68.0, 64.4, 64.1, 58.9, 54.9, 54.4, 37.7, 28.0, 28.0, 15.9

More carbon peaks detected than present in the structure due to presence of rotamers

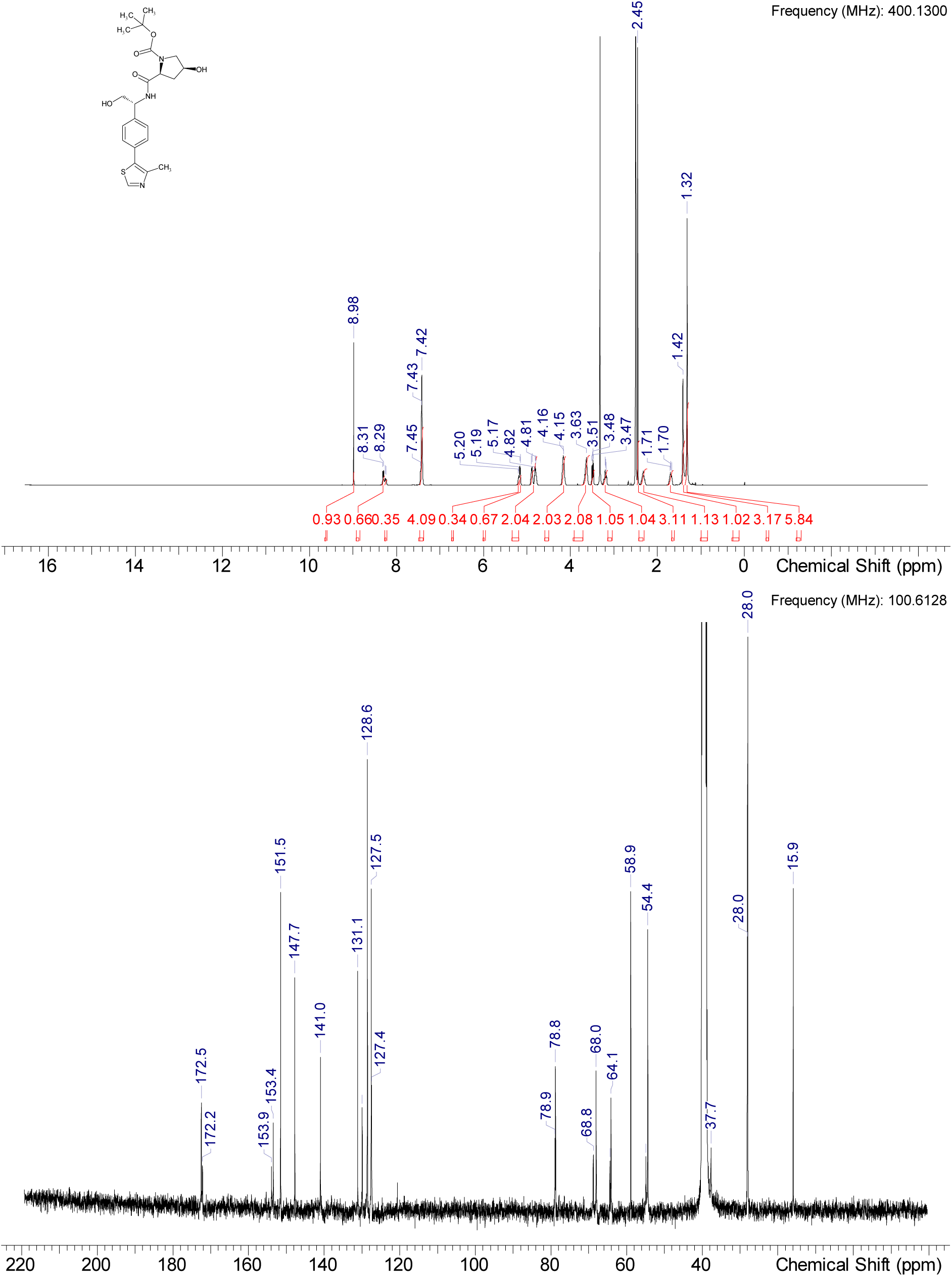

HRMS (m/z): [M+H]+ calcd. for C22H29N3O5S , 448.19007; found, 448.1904

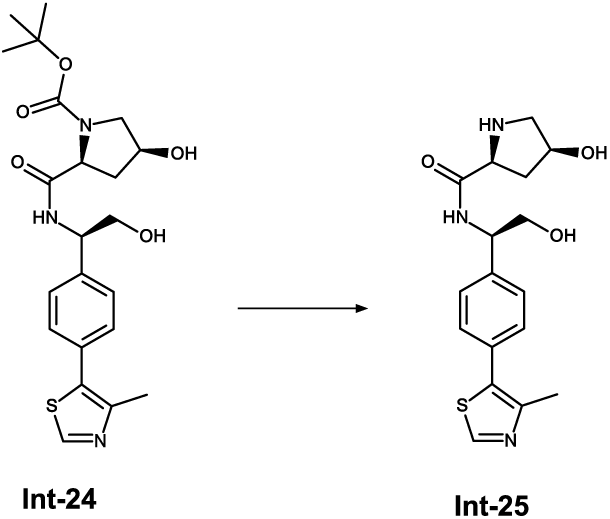

To a stirred solution of Int-24 (60 mg, 0.134 mmol, 1.0 eq.)in methanol (0.3 mL) is added a HCl solution in dioxane (0.101 mL, 0.402 mmol, 4 M, 3.0 eq.) at RT. The mixture is heated to 45°C and stirred for 90 min. The mixture is cooled the RT and the all volatiles are evaporated under reduced pressure to obtain the curde product Int-25 (51 mg, 0.134 mmol, 100% yield) as HCl salt which is used in the next step without further purification. An analytical sample of the free base was obtained by RP HPLC Method 3a) purification and lyophilization.

^1^H NMR (DMSO-d_6_) δ: 8.98 (s, 1H), 8.46 (br d, J=8.4 Hz, 1H), 7.33-7.54 (m, 4H), 4.81-5.23 (m, 2H), 4.73 (br s, 0.75H rotamer), 4.19 (br s, 0.25H rotamer), 4.13 (br s, 1H), 3.63 (br d, J=5.6 Hz, 2H), 3.47-3.58 (m, 1H), 3.19 (br s, 0.25H rotamer), 2.92 (dd, J=10.9, 5.3 Hz, 0.75H rotamer), 2.69 (dd, J=11.0, 3.9 Hz, 0.75H rotamer), 2.45 (s, 3H), 2.16 (br s, 1H), 1.79 (br d, J=11.4 Hz, 0.26H rotamer), 1.62 (dt, J=12.8, 5.0 Hz, 0.75H rotamer), one interchangeable missing

^13^C NMR (DMSO-d_6_) δ: 174.0, 151.5, 147.7, 141.1, 140.9, 131.1, 131.1, 129.9, 129.8, 128.6, 128.6, 127.4, 71.1, 64.6, 64.4, 59.3, 54.6, 54.5, 54.0, 15.9

More carbon peaks detected than present in the structure due to presence of rotamers

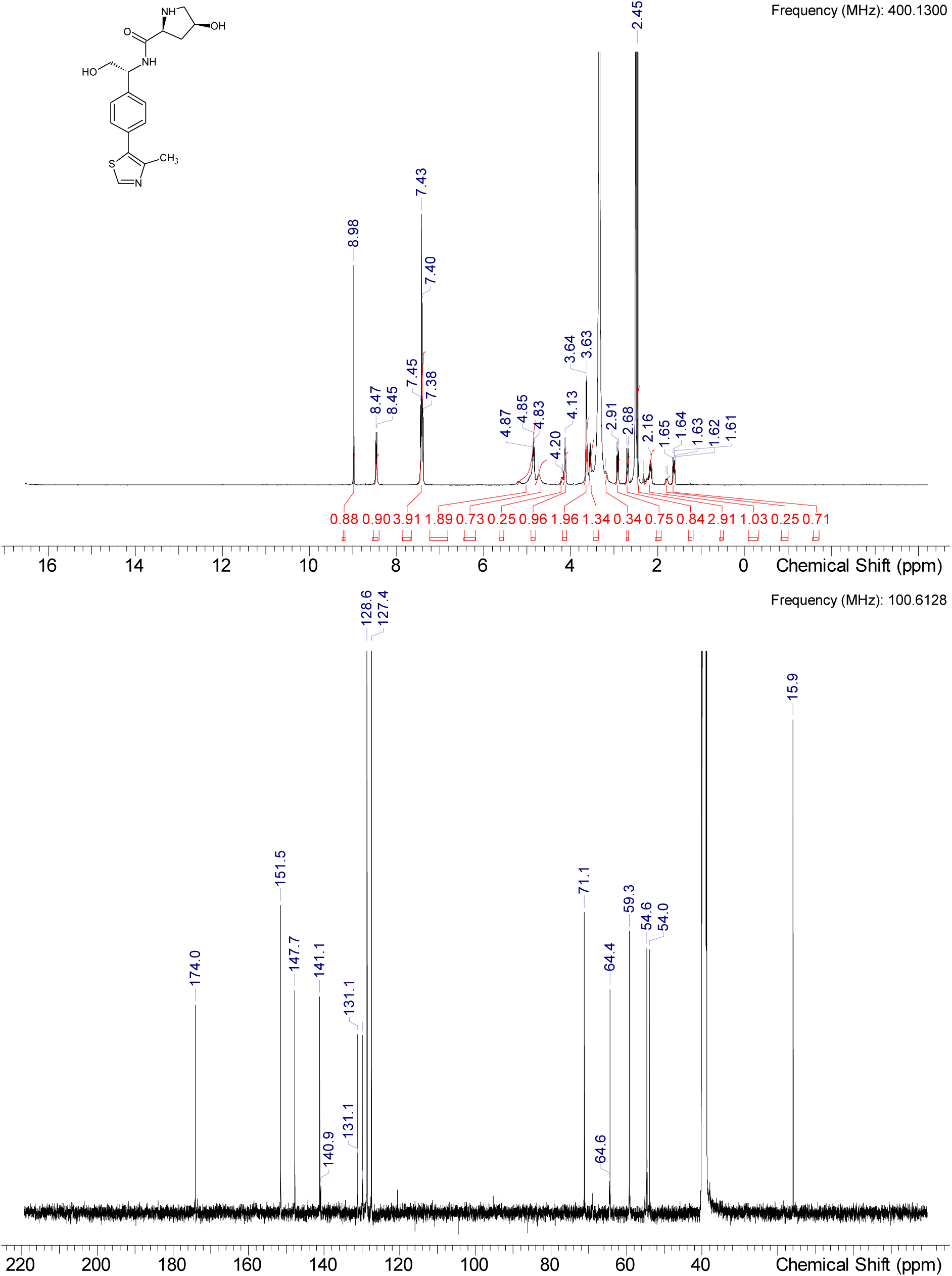

HRMS (m/z): [M+H]+ calcd. for C17H21N3O3S , 348.13788; found, 348.13788

#### Compound 8 (cis ACBI3)

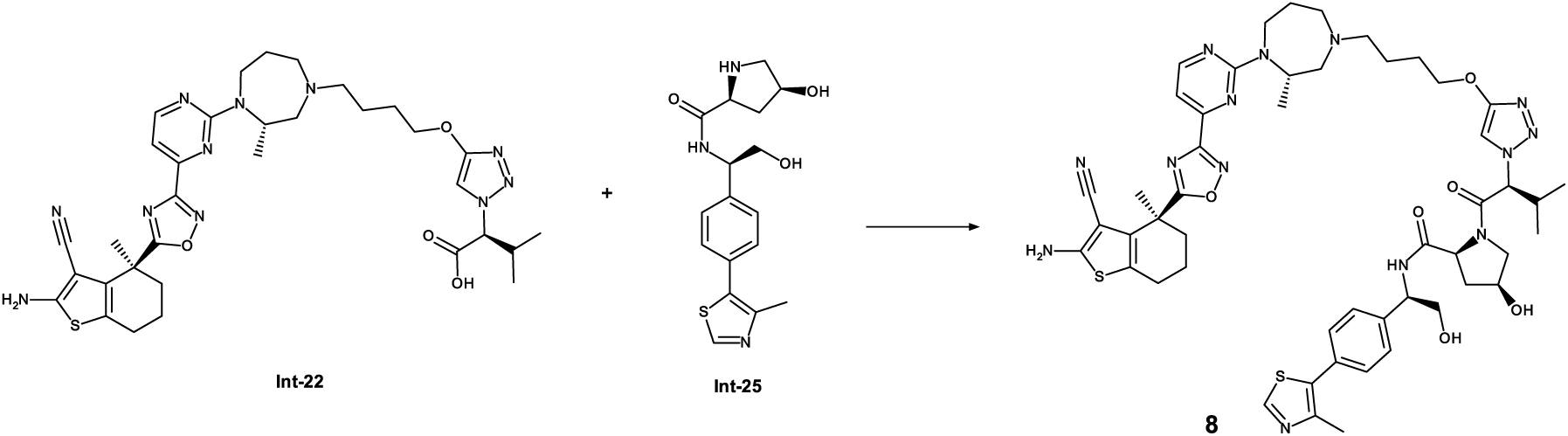

To a stirred solution of Int-22 (35 mg, 0.051 mmol, 1.0 eq.), Int-25 (50 mg, 0.130 mmol, 2.6 eq.) in NMP (1 ml) at 0°C is added HATU (28 mg, 0.071 mmol, 1.4 eq.) and DIPEA (0.044 mL, 0.254 mmol, 5.0 eq) and the reaction mixture is stirred at 0°C for 15 min. To the mixture is added water and acetonitrile and purified by RP chromatography (Method 4b) to obtain compound 8 (79 mg, 0.040 mmol, 79 %)

^1^H NMR (DMSO-d_6_) δ: 8.92 (s, 1H), 8.90 (s, 0.1H rotamer), 8.64 (d, J=7.6 Hz, 0.1H rotamer), 8.49 (d, J=4.6 Hz, 1H), 8.40 (d, J=7.6 Hz, 1H), 7.60 (s, 1H), 7.54 (d, J=18.5 Hz, 0.2H rotamer), 7.26-7.42 (m, 4H), 7.18-7.24 (m, 0.17H rotamer), 6.92-7.13 (m, 3H), 5.23 (d, J=6.8 Hz, 0.79H rotamer), 5.18-5.21 (m, 0.1H rotamer), 5.13 (s, 0.18H rotamer), 5.10 (br d, J=10.1 Hz, 1H), 4.89-4.94 (m, 0.1H rotamer), 4.79 (s, 2H), 4.55-4.70 (m, 1H), 4.09-4.38 (m, 3H), 3.95 (br s, 2H), 3.77 (br s, 1H), 3.63-3.72 (m, 0.24H rotamer), 3.41-3.62 (m, 3H), 3.12 (br s, 1H), 2.87-3.03 (m, 1H), 2.77 (br d, J=9.1 Hz, 1H), 2.47 (br s, 3H), 2.26-2.41 (m, 7H), 1.96-2.12 (m, 1H), 1.84 (s, 3H), 1.72 (s, 3H), 1.38-1.68 (m, 7H), 0.96 (br d, J=6.3 Hz, 6H), 0.87-0.92 (m, 0.52H rotamer), 0.61 (br d, J=6.6 Hz, 3H)

^13^C NMR (DMSO-d_6_) δ: 184.4, 171.1, 167.1, 166.2, 163.7, 160.7, 160.4, 160.1, 160.0, 153.5, 151.5, 147.8, 140.9, 131.3, 131.1, 129.9, 128.6, 127.4, 120.6, 119.6, 115.7, 107.2, 105.7, 81.3, 70.1, 68.8, 66.9, 64.4, 58.8, 56.2, 56.1, 55.1, 54.7, 50.0, 37.0, 36.7, 31.3, 26.4, 24.7, 23.6, 19.4, 18.2, 18.2, 16.0, 15.7, 15.6, 15.6, 1 C under DMSO

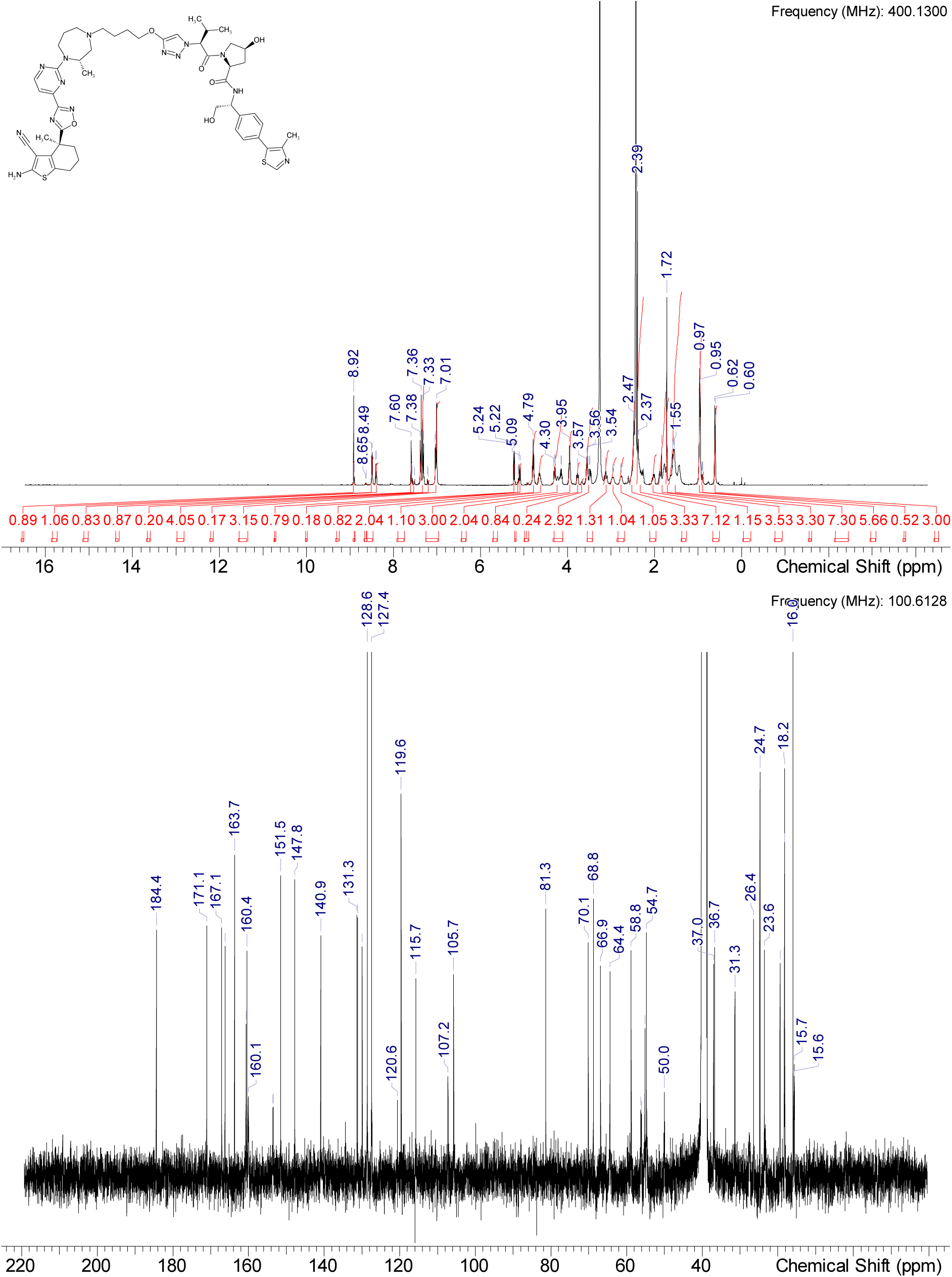

HRMS (m/z): [M+H]+ calcd. for C50H62N14O6S2 , 1019.44909; found, 1019.44897

## Notes

### Competing Interest Statement

Johannes Popow, Andreas Gollner, Christiane Kofink, Gerhard Fischer, Melanie Wurm, Nikolai Mischerikow, Carina Hasenoehrl, Heribert Arnhof, Silvia Arce-Solano, Sammy Bell, Georg Boeck, Jale Karolyi-Oezguer, Theresa Klawatsch, Manfred Koegl, Roland Kousek, Barbara Kratochvil, Katrin Kropatsch, Arnel A. Lauber, Sabine Olt, Daniel Peter, Oliver Petermann, Vanessa Roessler, Peggy Stolt-Bergner, Patrick Strack, Eva Strauss, Jens Quant, Harald Weinstabl, Peter Ettmayer are current or former employees of Boehringer Ingelheim. A.C. is a scientific founder, shareholder and advisor of Amphista Therapeutics, a company that is developing targeted protein degradation therapeutic platforms. The Ciulli laboratory receives or has received sponsored research support from Almirall, Amgen, Amphista Therapeutics, Boehringer Ingelheim, Eisai, Merck KaaG, Nurix Therapeutics, Ono Pharmaceuticals and Tocris-Biotechne.

### Summary of Updates

This is a revised version of the manuscript.

